# A 3-hydroxy-3-methylglutaryl-CoA synthase-based probe for the discovery of the acyltransferase-less type I polyketide synthases

**DOI:** 10.1101/509463

**Authors:** Haoyu Liang, Lin Jiang, Qiyun Jiang, Jie Shi, Jingxi Xiang, Xiaohui Yan, Xiangcheng Zhu, Lixing Zhao, Ben Shen, Yanwen Duan, Yong Huang

## Abstract

Acyltransferase (AT)-less type I polyketide synthases (PKSs) produce complex natural products due to the presence of many unique tailoring enzymes. The 3-hydroxy-3-methylglutaryl coenzyme A synthases (HCSs) are responsible for β-alkylation of the growing polyketide intermediates in AT-less type I PKSs. In this study, we discovered a large group of HCSs, closely associated with the characterized and orphan AT-less type I PKSs through *in silico* genome mining, sequence and genome neighborhood network analysis. Using HCS-based probes, the survey of 1207 inhouse strains and 18 soil samples from different geological locations revealed the vast diversity of HCS-containing AT-less type I PKSs. The presence of HCSs in many AT-less type I PKSs suggests their co-evolutionary relationship. Our study should inspire future efforts to discover new polyketide natural products from AT-less type I PKSs.

## INTRODUCTION

Many pharmacologically important natural products, such as antibiotic erythromycin, immunosuppressant rapamycin and anthelmintic avermectin, are biosynthesized by type I polyketide synthases (PKSs). Acyltransferase (AT)-less type I PKSs (also known as trans-AT PKSs) belong to a rapidly emerging family of novel PKSs, which are characterized by the absence of AT domain in each PKS module. The transfer of acyl CoA building block to the acyl carrier protein (ACP) domain in the elongation step is provided by one or more ATs *in trans.* Since the discovery and establishment of the biosynthetic logic for AT-less type I PKSs in leinamycin (LNM, 1) and pederin (2), the study of over 40 different polyketides have resulted in new insights into their biosynthesis beyond the canonical type I PKSs (Cheng et al., 2003; Helfrich and Piel, 2016; Piel, 2002, 2010). For example, the KSs from AT-less type I PKSs could be extremely useful for predicting the sub-structure of the corresponding genetic-coded natural products, based on their structural and evolutionary relationships (Lohman et al., 2015; Nguyen et al., 2008). In addition, 3-hydroxy-3-methylglutaryl coenzyme A synthases (HCSs) and the associated HCS cassette proteins are responsible for the β-branching in many AT-less type I PKSs-derived polyketides, including LNM, pederin, and bacillaenes (BAEs, 3 and 4) (Calderone, 2008; Calderone et al., 2006; Calderone et al., 2007; Liu et al., 2009) (Figures 1A and S1). Interestingly, HCSs are only observed in three canonical type I PKSs for curacin, jamaicamide and cylindrocyclophane biosynthesis, all of which were from photosynthetic bacteria (Chang et al., 2004; Edwards et al., 2004; Nakamura et al., 2012). The installation of various β-branches imposes an additional level of structure diversity in these natural products, critical for their conformation and individual biological activity (Haines et al., 2013).

**Figure 1.**
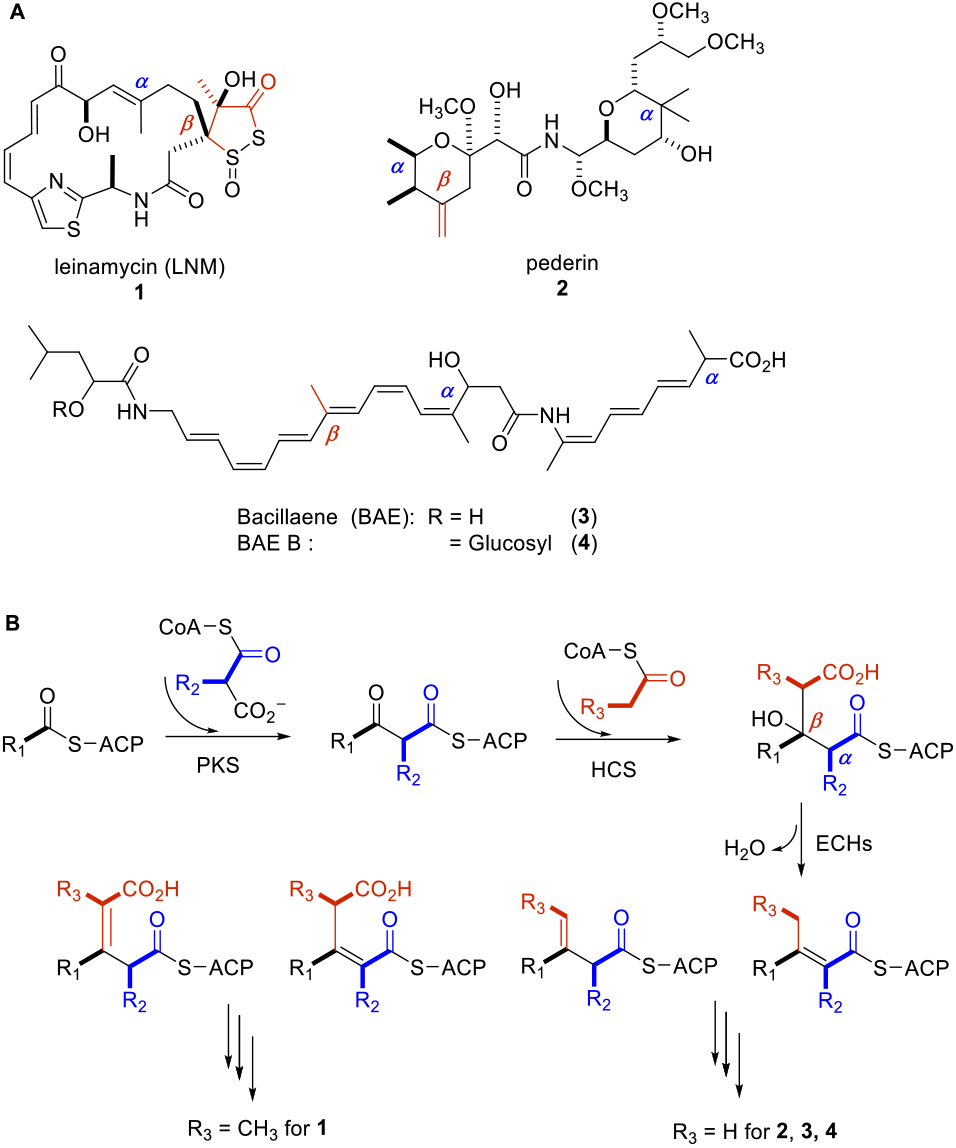
Selected polyketides biosynthesized by AT-less type I PKSs **(A)** and the mechanism of *β*-branch formation involving HCS cassette proteins **(B)**. The *β*-branches are highlighted in red.

The HCSs are responsible for generating structure diversity for polyketides, by attaching methyl, ethyl, or carboxylic acids to their *β*-carbons, in a similar manner with mevalonate biosynthesis (Calderone, 2008; Maloney et al., 2016). HCSs catalyze the Claisen condensation between acyl-*S*-ACP and the ACP-tethered polyketide to afford an HMG-*S*-ACP intermediate, followed by dehydration and/or decarboxylation by the action of enoyl-CoA hydratases (ECH1 and ECH2), resulting in the *β*-alkylated intermediates (Figure 1B). The enzymes involved in the above sequential catalysis are called HCS gene cassettes, which are integral parts of many AT-less type I PKS gene clusters.

A recent genome scale analysis suggested that AT-less type I PKSs may constitute about 38% of bacterial modular PKSs among the sequenced bacterial genomes (O’Brien et al., 2014). Our discovery of the LNM family of natural products through *in silico* and genome mining also revealed that there is huge potential for structure diversity from AT-less type I PKSs (Pan et al., 2017). Although KSs in either type I or II PKSs have been used to survey the structure diversity of PKSs in different geological locations (Charlop-Powers et al., 2015; Charlop-Powers et al., 2016), the diversity of AT-less type I PKSs in the environment, to our knowledge, has not been explored in a systematic manner. In this study, we surveyed the diversity of AT-less type I PKSs containing HCSs. Our studies identified 13 HCS-containing bacterial strains from 1207 in-house bacterial strain collection. Two of them have been confirmed to contain the respective AT-less type I PKS gene clusters. The survey of 18 soil samples revealed the presence of numerous HCSs in soil, which might be associated with AT-less type I PKSs. Our study suggests the wide-spread of AT-less type I PKSs and should inspire future efforts to discover more polyketide natural products from this important family of PKSs.

## RESULTS

### A specific type of HCSs mainly clustered with AT-less type I PKSs

In order to understand the evolution of HCSs in AT-less type I PKSs, a total of 6,712 HCS protein sequences from GenBank (as of September 27, 2018) with > 25% sequence identity of LnmM HCS from LNM biosynthetic gene cluster (BGC) were obtained. These HCSs were clustered into 2,250 representative sequences with 90% identity cutoff using CD-HIT (Huang et al., 2010). Sequence similarity network (SSN) analysis of the 2,250 HCSs were conducted using multiple *E*-value cutoffs ranging from 1 × 10^-10^ to 1 × 10^-150^ (Figures 2 and S2). An *E*-value threshold of 1 × 10^-135^ showed ~70% sequence identity for HCSs, resulting in 11 major clusters. The SSN analysis revealed that most HCSs are grouped to mevalonate biosynthetic pathways, which reflects their house-keeping roles in these organisms. In contrast, cluster C contains 251 HCSs, which belong to several bacterial phyla, including Actinobacteria, Firmicutes and Proteobacteria. Cluster C was identified as HCSs for *β*-branching in polyketide biosynthesis based on the following reasons: (1) this cluster contains all of the 32 HCSs from the 29 characterized AT-less type I PKSs, as well as JamH (AAS98779.1), CurD (AAT70099.1) and CylF (ARU81120.1) from canonical type I PKSs (Figures 2B and S1); (2) 200 out of 219 orphan HCSs are clustered with the 170 putative AT-less type I PKS gene clusters, based on the co-localization and similarity of their flanking open reading frames (ORFs) with proteins from characterized AT-less type I PKSs (Tables S1-S170). Interestingly, only two HCSs, AJC59428.1 and CP001344.1, both from *Cyanothece* sp. PCC 7425, are clustered with canonical type I PKSs (Tables S171 and S172), while seven HCSs may be associated with type II PKSs (Tables S173-S179). The role of the remaining 10 HCSs cannot be determined due to the incomplete genome sequencing (Tables S180-S189).

**Figure 2.**
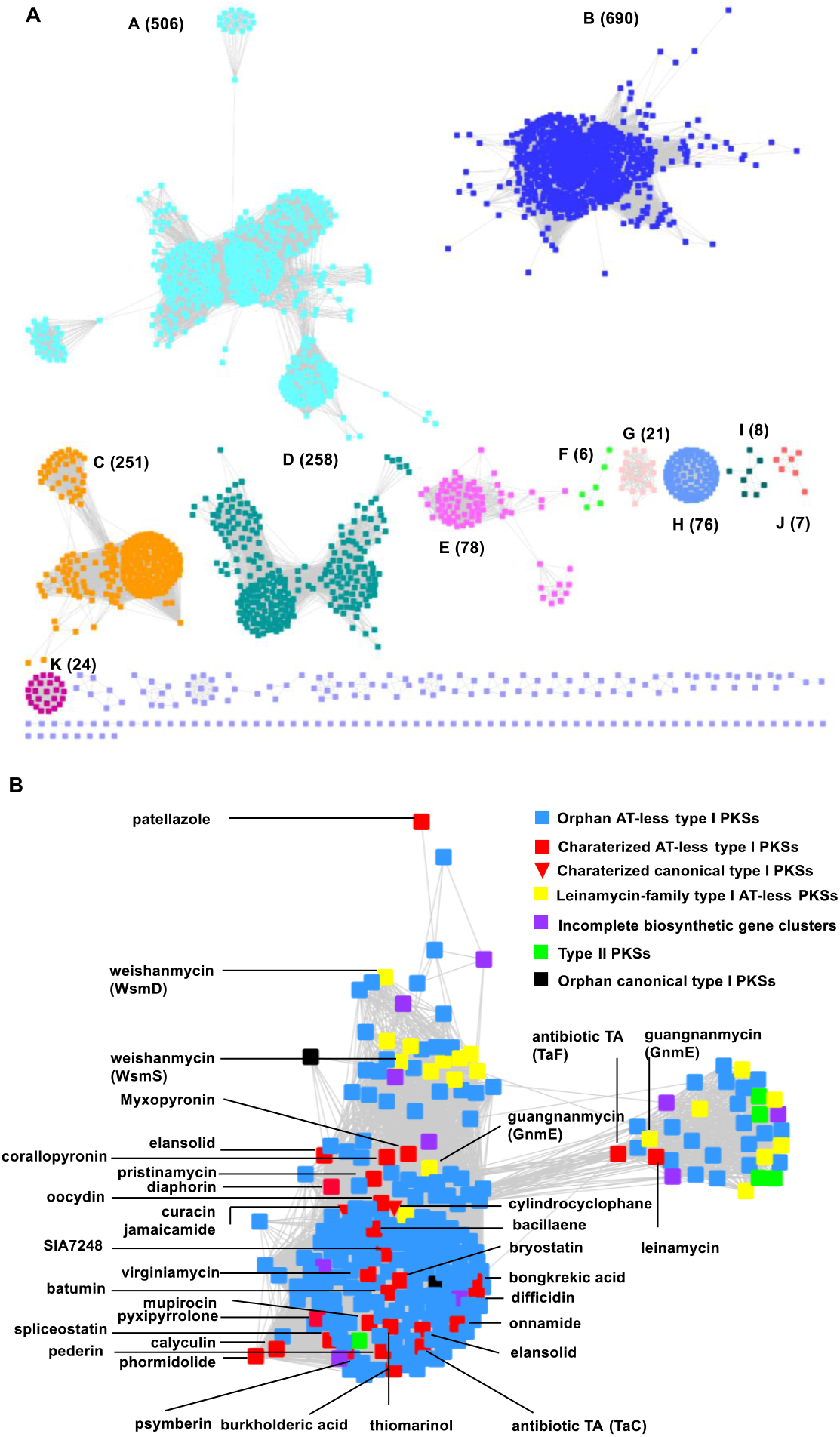
Sequence similarity network (SSN) analysis for HCSs. **(A)** SSN of the 2,215 HCS sequences. The SSN was displayed with an *E*-value threshold of 10^-135^. Colors represent different clusters of HCSs. **(B)** A sub-group of HCSs responsible for **β**-branching in polyketide biosynthesis. The corresponding polyketide products and their corresponding biosynthetic enzymes were embedded.

Since our mining of AT-less type I PKS using HCS probes *in silico* reveals 170 orphan AT-less type I PKSs, we performed their genome neighborhood network (GNN) analysis (Figure 3). In order to understand the evolutionary relationships of the HCS cassettes and AT-less type I PKSs, the characterized AT-less type I PKSs containing HCS cassettes were also included. Using *E*-value cutoff of 1 × 10^-6^, the GNN analysis revealed not only the clustering of HCSs in the context of the entire AT-less type I PKSs, but the close association of the other accessory enzymes in HCS cassettes with AT-less type I PKSs, including ACPs, ECHs, ATs, ketosynthases (KSs) and acyltransferase/decarboxylases (AT/DCs). Under the current *E*-value cutoff, HCSs in cluster C are likely grouped, based on their respective substrates propionyl-*S*-ACP or acetyl-*S*-ACP (Figures 1B and S1) (Pan et al., 2017). In addition, phylogenetic analysis suggested that the ACPs from HCS cassettes were also clustered into propionyl-*S*-ACP or acetyl-*S*-ACP specific groups (Figure S3). Therefore, the association of these proteins with the AT-less type I PKSs might reflect their co-evolutionary relationships, which is consistent with the recent observation of strong protein-protein interaction between CurD and its donor ACP in canonical type I PKSs, as well as the unique acceptor ACP domains present within the PKS modules for **β**-branching in mupirocin AT-less type I PKS (Haines et al., 2013; Maloney et al., 2016).

**Figure 3.**
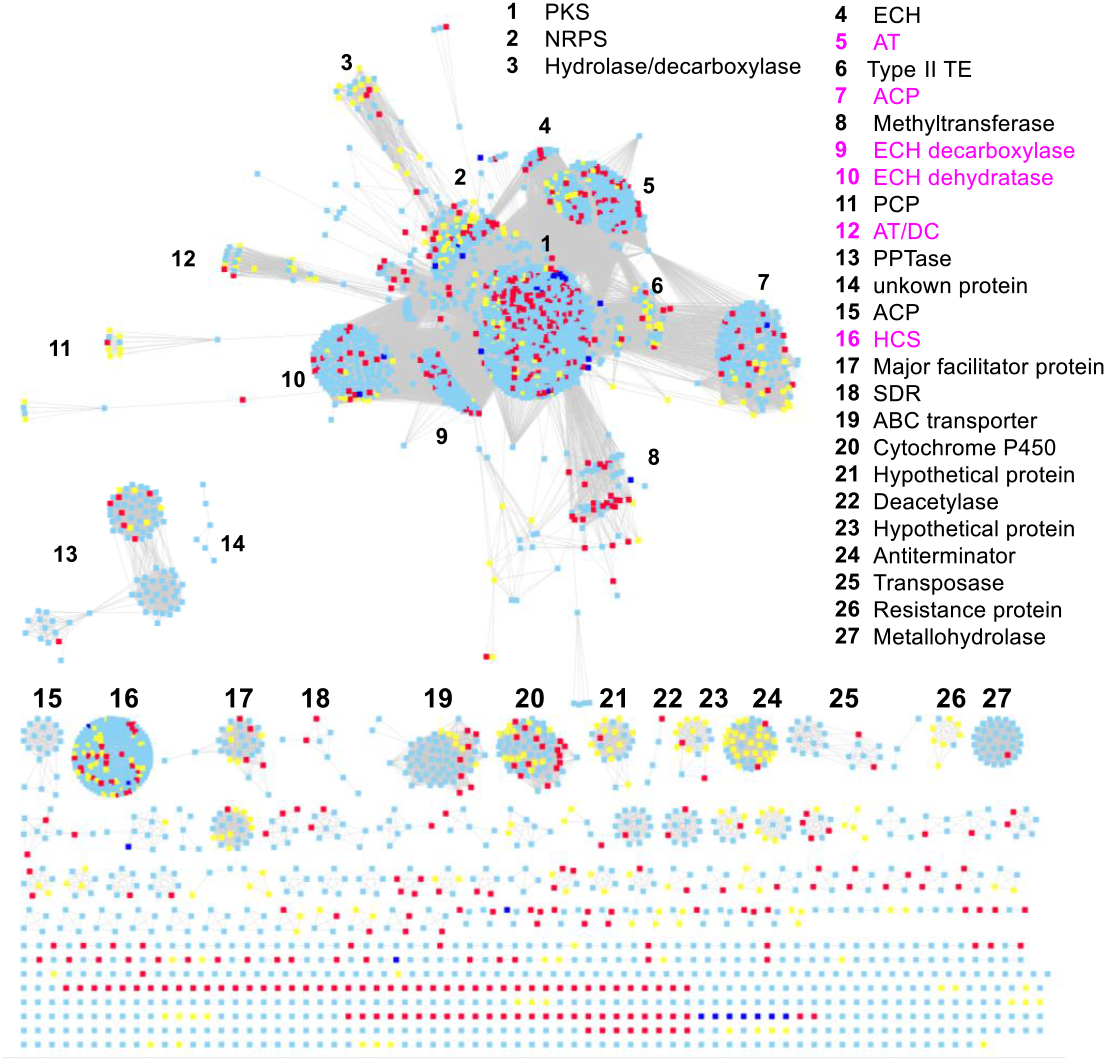
GNN analysis of 170 orphan AT-less type I PKSs and 29 characterized AT-less type I PKSs. Blue square 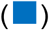: orphan AT-less type I PKSs; red square 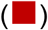: characterized AT-less type I PKSs; yellow square 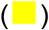: leinamycin-type AT-less type I PKSs. The inserted legends represent each protein annotation, in which the HCS cassette proteins are colored in pink.

### Survey 1207 bacterial strains using HCS-specific probes

HCSs have been used to identify and clone the BGCs of potent antitumor compound FR901464 in *Burkholderia*, cylindrocyclophane with in vitro activities towards 20S proteasome from cyanobacteria and protein phosphatase 1 inhibitor calyculin from a metagenomic library of a marine sponge (Nakamura et al., 2012; Wakimoto et al., 2014; Zhang et al., 2011). We therefore hypothesized that HCS-based PCR probe could be used to discover AT-less type I PKS BGCs in our strain collection (Hindra et al., 2014; Yan et al., 2016; Pan et al., 2017). Because these HCSs are highly conserved, degenerate primers were designed using their conserved region in characterized AT-less type I PKSs BGCs (Figures 4 and S4). The primers were first used to clone LnmM, the HCS gene from *lnm* gene cluster (Figure S4). They were subsequently used in the PCR detection of HCS genes from 1207 bacterial strains, resulting in 13 strains with the expected 0.2-to 0.3-kb DNA fragments. These DNA fragments were sequenced and confirmed to be partial HCS genes, which were used to construct an evolutionary tree with known HCSs associated with AT-less type I PKSs using Mega 7 (Figure 4C). The HCS genes in those 13 strains are clustered into six branches, using the HCS from the mevalonate biosynthesis pathway of *Staphylococcus aureus* as an out group. Six HCS genes from CB00144, CB00158, CB01229, CB01357, CB01456, and CB01509, are in the same cluster with SnaI (CBW45745.1) and VirC (BAF50725.1) involved in the biosynthesis of streptogramin A-type antibiotics. Three HCSs from CB01882, CB01883 and CB02980 are similar to LnmM, suggesting their involvement in the biosynthesis of the recently identified leinamycin-family of natural products, in which a novel polyketide weishanmycin was identified from CB01883 using an alternative probe of DUF-SH domain (Pan et al., 2017). The remaining HCSs from CB00111, CB01828 and CB02999 resemble CalT, CorE and BaeG, with sequence identities of 67%, 62% and 98%, respectively.

**Figure 4.**
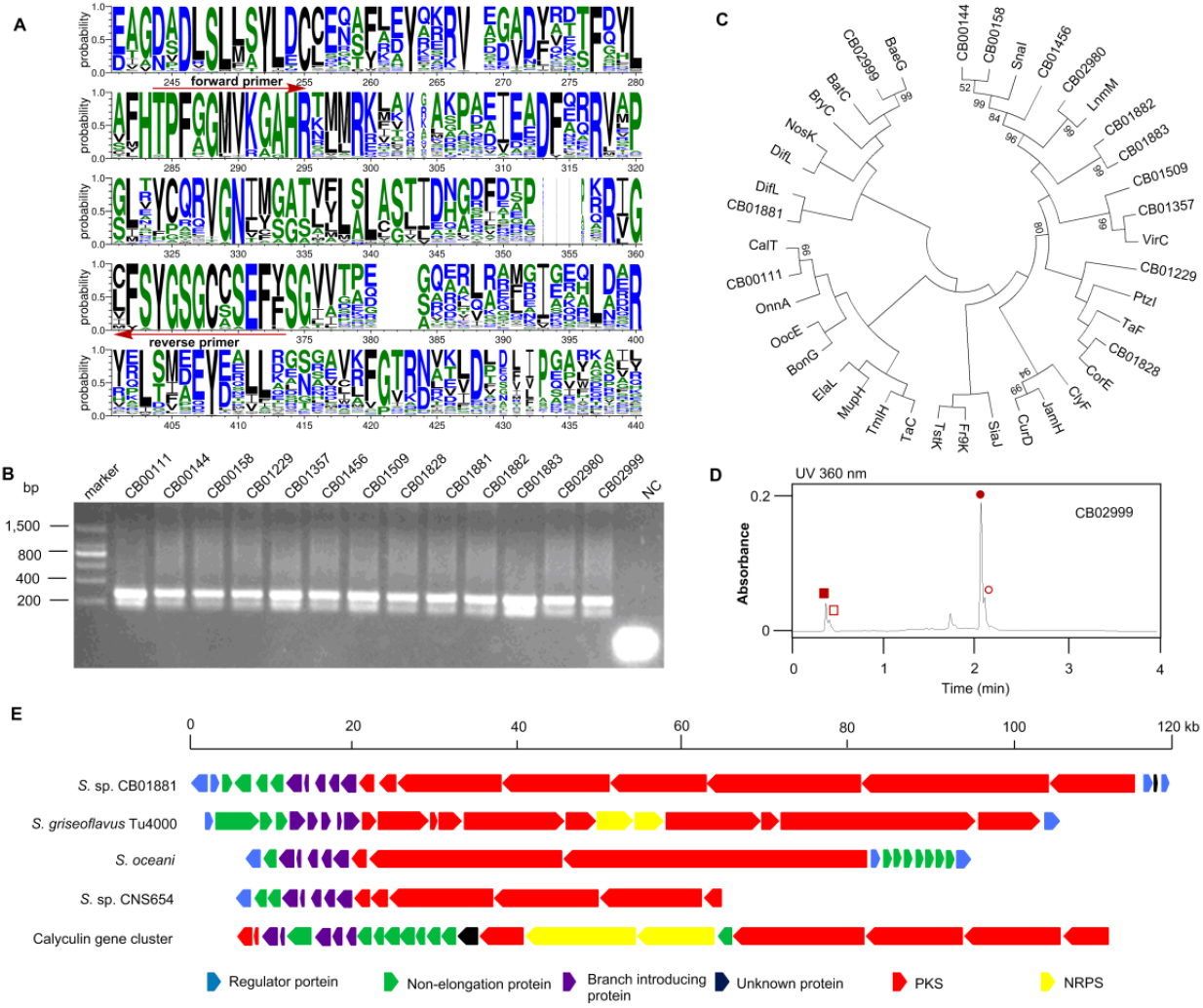
The survey of 1207 in-house strains using specific HCS probes. **(A)** Sequence alignment of selected HCS sequences from characterized AT-less type I PKS BGCs. Including BaeG (ABS74058.1), BatC (ADD82944.1), BonG (AFN27479.1), BryR (ABM63533.1), CalT (BAP05578.1), CorE (ADI59527.1), CorE (ADI59527.1), CylF (ARU81120.1), DifL (CAG23983.1), ElaL (AEC04358.1), Fr9K (AIC32697.1), LnmM (AAN85526.1), MupH (AAM12922.1), NosK (WP_094329473.1), OnnA (AAV97869.1), OocE (AFX60327.1), PedP (AAW33975.1), SnaI (CBW45745.1), TaC (CAB46502.1), TaF (CAB46505.1), TmlH (CBK62727.1) and TstK (AGN11885.1). Aligned amino acid residues are colored based on the conservation level. The amino acid sequences used to design degenerated primers to clone HCS genes using CODHOP strategy; **(B)** PCR amplification of specific HCS gene fragments from inhouse strains; **(C)** Phylogenic analysis of the cloned HCS gene fragments; **(D)** The HPLC profile of *B. amyloliquefaciens* CB02999 and the production of BAEs. 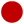, BAE (3); 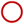, dihydro-BAE; 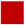, BAE B (4); 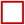, dihydro-BAE B. **(E)** The BGC of AT-less type I PKS in S. sp. CB01881.

Because GNN analysis suggests that a sub-group of HCSs are clustered with AT-less type I PKS BGCs, strain CB02999 and CB01881 were further selected to determine if they could produce specific polyketides biosynthesized by AT-less type I PKSs or contain their BGCs. The HCS from CB02999 shares high sequence similarity with the BaeG HCS, involved in the biosynthesis of BAEs in *Bacillus subtilis* and *Bacillus amyloliquefaciens* (Chen et al., 2006). CB02999 was first determined to resemble *Bacillus amyloliquefaciens* based on its house-keeping genes 16s rRNA and *rpoB* (NR_041455.1 and CP003332.1, 99% and 99% identity). The production of BAE, dihydro-BAE, and the glycosylated BAE derivative in CB02999 were confirmed by comparing their UV spectra and high resolution mass spectrometry (HRMS) with those previously reported (Figures 4D and S5) (Butcher et al., 2007; Moldenhauer et al., 2010; Qin et al., 2014). These studies suggested that *B. amyloliquefaciens* CB02999 contains the prototype AT-less type I PKS responsible for the biosynthesis of BAEs.

In order to identify the AT-less type I PKS BGC in CB01881, shotgun genome sequencing data were generated using the Illumina HiSeq 2000. Analysis of the genome sequence of CB01881 revealed the presence of an AT-less type I PKS BGC. The encoded PKS enzymes are similar to the PKSs for the biosynthesis of calyculin (~36% identity) and a few orphan AT-less type I PKSs from *S. griseoflavus* Tu4000 (GG657758.1), *S. oceani* DSM 16646 (CP002131.1) and S. sp. CNS654 (KL370899.1) (Figure 4E). The AT-less type I PKS in CB01881 contains multiple PKS modules without AT domains in six ORFs and four discrete ATs. Its HCS containing the previously cloned HCS gene fragment was also identified (Tables S190-191). Interestingly, four contiguous ACP di-domains were also present in the PKS modules, which were the putative sites to attach the *β*-branches (Haines et al., 2013).

### The survey of 18 soil samples using HCS probes revealed the presence of diverse HCSs in the environment

The discovery of more than 200 HCSs in AT-less type I PKS BGCs *in silico* and the identification of strains containing these PKSs in our in-house strain collection, encouraged us to study the diversity of HCSs and AT-less type I PKSs in the environment. The biosynthetic potential of polyketides in environmental samples is typically surveyed using conserved KS domains in type I or type II PKSs. One limitation is that a single bacteria strain in a typical soil sample may contain dozens of KS domains, due to the presence of several PKS gene clusters and multiple PKS modules in each type I PKS. Therefore the accurate identification of KSs in soil samples would rely on the deep-sequencing of environmental DNAs, in order to reduce the false negative results (Crits-Christoph et al., 2018). Because only one or two HCSs are present in AT-less type I PKS BGCs, we envisioned the potential to survey environmental HCS genes specific for AT-less type I PKSs.

We first determined if the current HCS probes could be used to survey soil samples. Eighteen soil samples from various places in China were collected and the environmental DNAs were extracted based on the reported protocol (Figure 5) (Charlop-Powers et al., 2015). Clear DNA fragments with the expected size of 0.2-to 0.3-kb were amplified from all of the soil samples, which were gel-purified and used for high-throughput sequencing using Illumina MiSeq platforms, along with the mixture of the 13 in-house strains containing HCSs (CB). The obtained DNA sequences were then clustered into operational taxonomic units (OTUs) at a sequence distance of 10% (> 90% sequence identity), and analyzed by BlastX. Based on the analysis of the HCS genes from the known AT-less polyketide gene clusters and the HCS genes cloned from the in-house strains, an *E*-value above 10^-28^ was deemed to be HCS homologues for *β*-alkylation (Figure 2).

**Figure 5.**
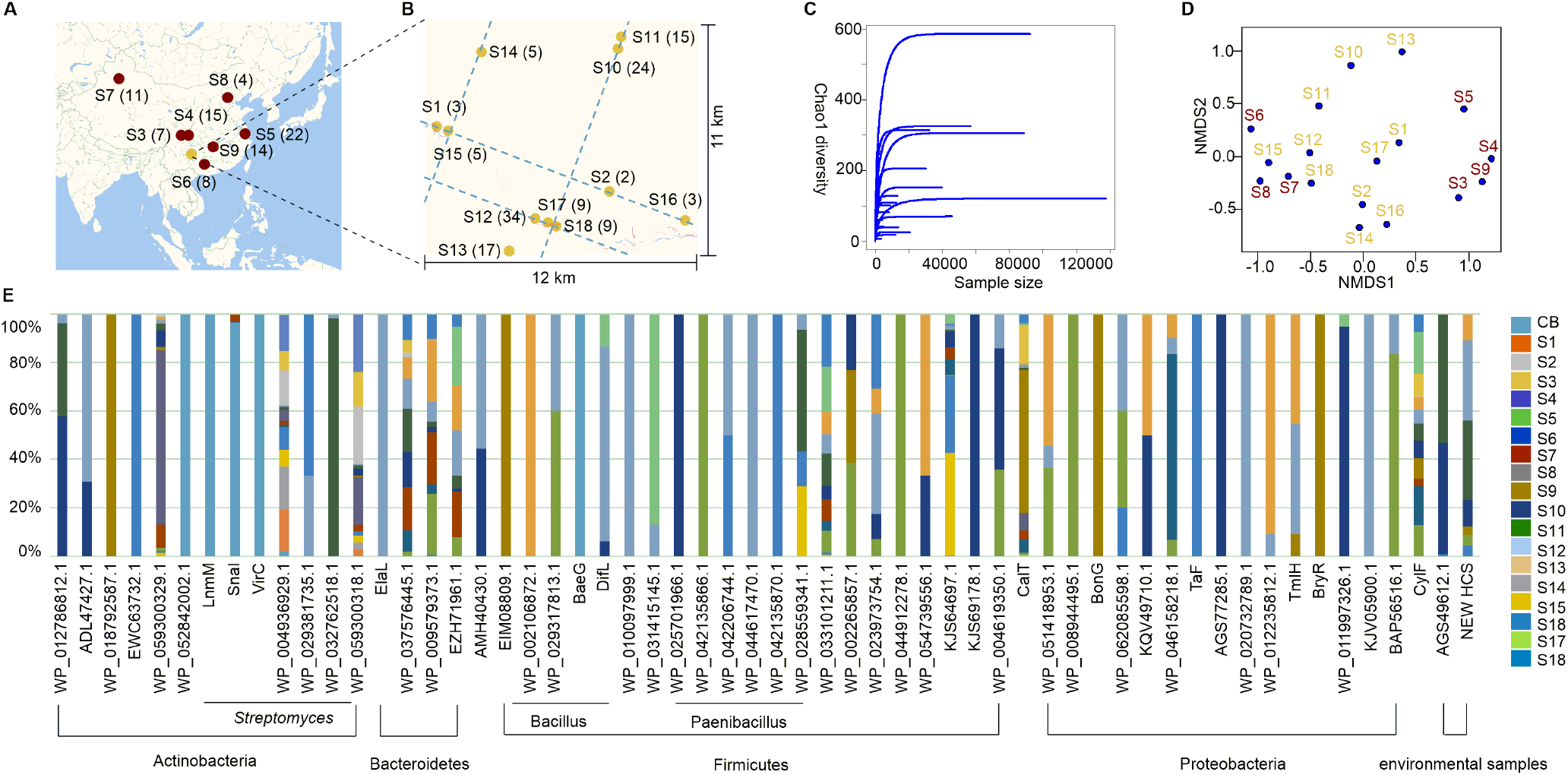
Profiling of HCS abundance in 18 soil samples. **(A)** Geological location of the 18 soil samples. The number of different HCS OTUs only with over 100 reads from high-throughput sequencing in each soil sample were included for the analysis and shown in parentheses; **(B)** The determination of the richness of HCSs using Chao1-based rarefraction analysis; **(C)** NMDS analysis of HCSs. **(D)** The distribution of HCS genes in soil samples S1-S18.

Rarefaction analysis suggested that the sequence space of these HCS genes was saturated (Figure 5C). Nonmetric multidimensional scaling (NMDS) ordination analysis revealed a tremendous beta diversity for the 18 soil samples (Figure 5D). For example, 12 out 18 soils have more than 6 different types of HCSs, and there are 34 different HCS genes in soil S12. Based on these pioneering studies to survey the secondary metabolite biosynthetic genes diversity, various factors of the soil samples, such as elevation, annual precipitation, temperature, pH, latitude and longitude of sampling sites, may contribute to the high beta diversity for soil samples S3-S9 (Charlop-Powers et al., 2015; Charlop-Powers et al., 2016). Interestingly, soil samples S1-S2 and S10-S18, collected from an area of around 120-km^2^ region within the Yunnan-Kweichow plateau, still possessed great beta diversity, in comparison to samples collected 1,000-km away. This was attributed to the microenvironment difference in the individual sampling sites (Charlop-Powers et al., 2016).

Interestingly, the HCS genes of streptogramin-A type compounds were not widely distributed, which were different from our in-house screening and previous reports (Genilloud et al., 2011). Several other HCS genes, including EZH71961.1, WP_004936929.1, WP_033101211.1, WP_037576445.1, and WP_059300318.1, are abundant in the surveyed soil samples (Figures 5E and S6). This result implied that there might be certain differences for the HCS diversity between our strain collection and the collected soil samples. The OTUs of the HCS genes in soil samples S3-S7, S9-S13 and S17-18 were significantly more than those in S1-S2, S8 and S14-S16 (Figure S7). Putative new HCSs were also identified in soil samples S4-S5 and S9-S13 (Figure 5E). The survey of the diversity of HCSs in soil samples and the presence of specific HCS genes in all the samples suggested the wide-spread of AT-less type I PKSs.

### The presence of diverse AT-less type I PKS specific KS domains in soil samples

Canonical type I PKSs share high homology and mainly clade with members of their biosynthetic pathways, while the KS domains from AT-less type I PKS possess substrate specificity and fall into phylogenetic clades that correlate with their substrates. These differences have been used to dissect these two types of PKSs (Piel et al., 2004; Jenke-Kodama et al., 2005; Nguyen et al., 2008; Lohman et al., 2015). In order to verify the presence and the diversity of AT-less type I PKSs in the soil samples, we next determined the presence of KSs belonging to AT-less type I PKSs, using 11 out of the previous 18 soil samples with specific KS-based PCR probes (Figure 6).

**Figure 6.**
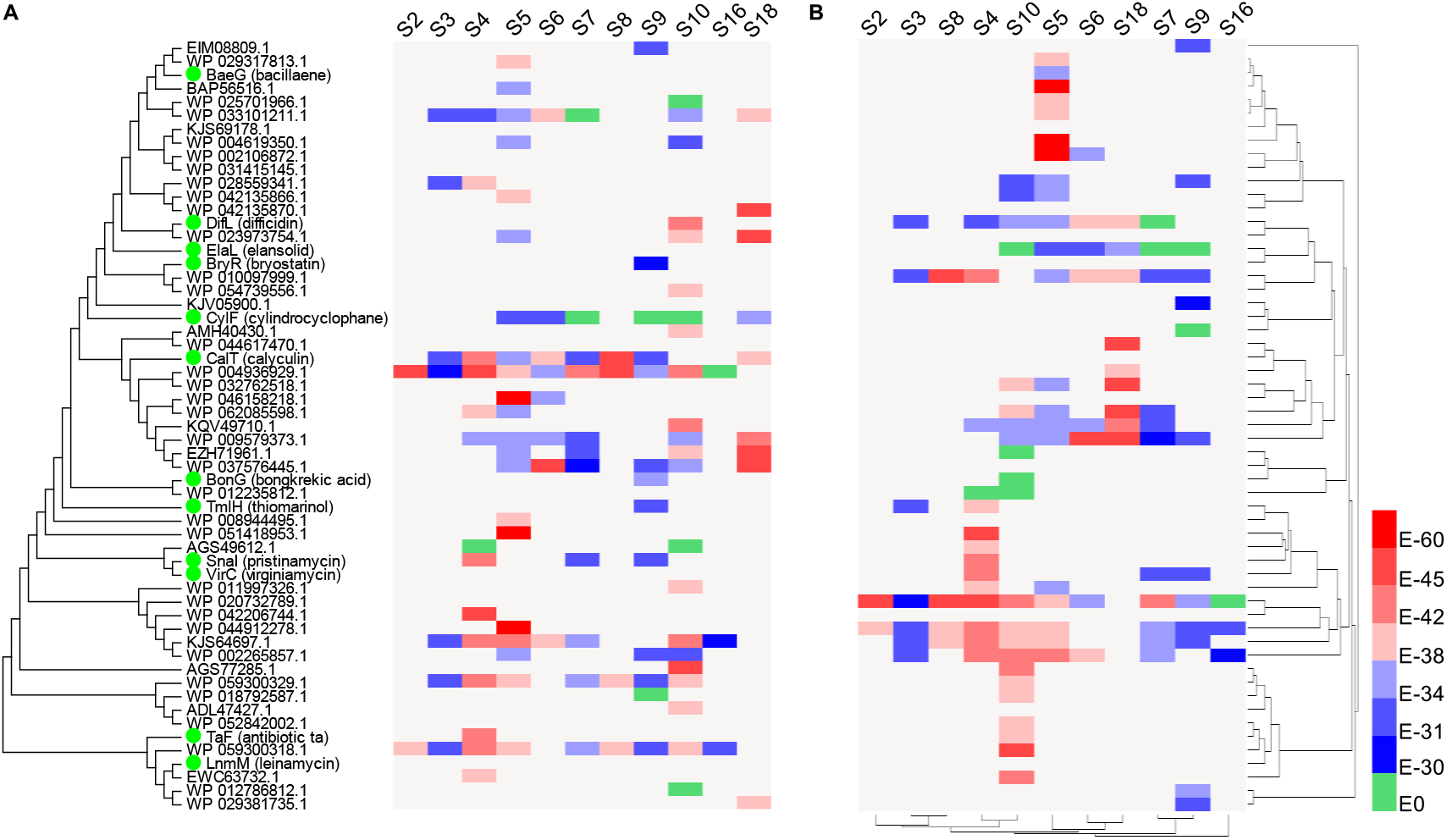
Profiling of KS domains of AT-less type I PKS in 11 soil samples. **(A)** The heatmap of KS domains from AT-less type I PKSs in 11 soil samples. The rows were arranged to match the phylogenetic tree of HCSs in the left. The color of the heatmap represents the similarity of the KS domains in each soil sample with the individual KS domains in HCS-containing AT-less type I PKS BGCs. The similarity was calculated using BlastX. **(B)** The heatmap of AT-less type I PKS KSs of each soil. The rows and columns were clustered based on similarity of the elements of the corresponding heatmap.

The expected KS gene fragments were amplified by PCR from the soil DNAs, using the previously reported primers to clone the conserved KS domains in type I PKSs (Figure S8) (Savic et al., 2006). The cloned KS DNA fragments were subjected to high-throughput sequencing, and then clustered into OTUs at a sequence distance of 10% (> 90% sequence identity), and compared to 224 KS domains from identified AT-less type I PKSs in GenBank by BlastX, using cylindrocyclophane as a control from canonical type I PKSs. The *E*-value ranging from 5.82 × 10^-31^ to 2.37 × 10^-53^ were deemed appropriate to assign the KS sequences from the soil DNA to each AT-less type I PKS. Our analysis indicated that there were abundant KS domains belonging to AT-less type I PKS gene clusters containing HCSs. Consistent with the presence of multiple HCSs in specific soil samples, various KS fragments belonging to AT-less type I PKSs were identified from the corresponding soils. For example, soil samples S5 and S10 contained KS fragments with high sequence similarity to more than 20 different types of AT-less type I PKSs, suggesting the huge biodiversity even in a single soil sample (Figure 6).

## DISCUSSION

Microbial natural products remain one of the most important sources of new small molecule drugs, while the discovery of novel scaffolds become increasingly difficult in well-studied bacterial phyla (Newman and Cragg, 2016; Cameron et al., 2017). Polyketides produced by AT-less type I PKSs represent a growing family of natural products with complex molecular structures and promising biological activities (Piel, 2010; Helfrich and Piel, 2016). Compared to canonical type I PKSs, AT-less type I PKSs possess not only KS domains specific for various alky substrates in individual modules, but various tailoring enzymes for online modification of the growing polyketide chains, such as the imbedded methyltransferase, DUF-SH domains and HCSs (Pan et al., 2017; Xie et al., 2017). HCSs play important roles to diversify polyketide structures for modular PKSs, especially AT-less type I PKSs (Calderone et al., 2006). The current study revealed that a unique cluster of HCSs of various bacteria phyla are closely associated with characterized and orphan AT-less type I PKSs. HCS-specific primers were then used to screen 1207 microbial strains and 18 soil samples collected in representative geological locations in China, which reveals the vast biodiversity of HCS and AT-less type I PKS genes in the environment (Figures 5 and 6).

The specific KS probes have been widely used to survey the biogeographic diversity of type I PKSs and type II PKSs, which reveal different biosynthetic richness of specific soil types in various geographic locations (Charlop-Powers et al., 2015; Charlop-Powers et al., 2016). In this study, specific HCS gene-based probe was used for the first time to survey the biogeographic distribution of HCSs and the AT-less type I PKSs. One unique feature of the HCS survey is that HCS is only responsible for the *β*-alkylation of the growing polyketide intermediates on ACPs, while distantly related to the overall polyketide structures, which could lead to discover new polyketides with diverse molecular scaffolds. In addition, there are only limited HCSs present in one bacterial strain, compared to dozens of KSs or KS domains in many bacteria (Crits-Christoph et al., 2018). Together with the KS survey in the 11 soil samples and BlastX analysis with KS domains from AT-less type I PKSs, our study revealed that soil samples are rich in HCSs and AT-less type I PKSs. For example, 57 unique HCS genes were distributed in 18 different soil samples, while one soil sample may contain dozens of HCS genes. This is in line with the recent survey of the biosynthetic potential in soils and therefore should be useful for the discovery of new polyketides (O’Brien et al., 2014).

The HCSs in AT-less type I PKSs may be acquired through horizontal gene transfer. Based on the phylogenetic and comparative genome analyses, the genes for mevalonate biosynthesis in gram-positive cocci were proposed to be horizontally transferred from a primitive eukaryotic cell (Wilding et al., 2000). Our results suggested that HCSs in “HCS cassette” would likely be recruited to AT-less type I PKSs from the mevalonate pathway, in a later evolution point. After the HCSs were incorporated into PKS BGCs, HCSs may further co-evolve with type I PKSs to facilitate the *β*-alkylation, with the presence of the acceptor ACPs containing the conserved Tyr residue in the PKS module (Haines et al., 2013). For example, the LnmKLM HCS cassette proteins are not only co-transcribed with the LNM PKSs in LNM biosynthesis, inactivation of either of the *lnmK, lnmL* or *lnmM* generated a series of shunt metabolites from the premature release of early biosynthetic intermediates, probably due to disruption of specific protein-protein interaction between HCS cassette proteins with LNM PKSs (Huang et al., 2011). There are several other discrete proteins in HCS cassette, including ACPs, ECHs and AT/DCs or ATs. How these proteins were gradually incorporated into the AT-less type I PKS remains a fascinating evolutionary question.

The AT-less type I PKSs have been proposed to evolve independently from canonical type I PKSs mainly because those KS domains only form certain clades based on their individual acyl-*S*-ACP substrates, while KSs from canonical type I PKSs form biosynthetic pathway-specific clades. Recent structural and evolutionary relationships study of KSs in AT-less type I PKSs captured several of their evolutionary traits from canonical type I PKSs, including the presence and transformation of KS-AT didomain OzmQ-KS1 towards a KS domain in AT-less type I PKSs for oxazolomycin biosynthesis (Lohman et al., 2015). Interestingly, only three HCS cassettes are present in known canonical type I PKSs, including curacin, jamaicamide and cylindrocyclophane, while two HCS cassettes are identified in orphan canonical type I PKSs (Tables S171 and S172). All of the five HCS cassettes for canonical type I PKSs are from cyanobacteria. In contrast, there are 32 HCSs in the characterized AT-less type I PKSs, and 217 orphan HCSs for AT-less type I PKSs. The preference of *β*-alkylation in AT-less PKSs might attribute to the presence of a standalone AT responsible for priming the ACP with the malonyl-CoA, while canonical type I PKSs typically lack this activity. In addition, *β*-alkylation in AT-less type I PKSs greatly expands their structural diversity and rigidity, due to the malonyl-CoA specific AT activity in most AT-less type I PKSs, comparing to the diverse acyl-CoA substrates of ATs in canonical type I PKSs (Chan et al., 2006). Interestingly, a few HCSs associated with orphan type II PKS gene clusters were also identified and their role in morphing aromatic polyketides remains to be deciphered.

## MATERIALS AND METHODS

### Bacterial strains, culture conditions and general methods

All chemical and biological reagents, including antibiotics, inorganic salts, carbon and nitrogen sources used in this study were obtained from common commercial sources, unless otherwise specified. Tryptic soy broth (TSB) with 0.2% glycine were used for mycelium growth, containing (per liter): glycine 2 g, tryptone 17 g, soy peptone 3 g, NaCl 5 g, K_2_HPO_4_ 2.5 g, dextrose 2.5 g, pH 7.3 ± 0.2. S. sp. CB01881 was grown at 30 °C on R2A agar for sporulation. *B. amyloliquefaciens* CB02999 was cultured in 50 mL TSB in 250-mL shaking flask and grown for 24 h at 30 °C and 220 rpm. Then 5 mL of seed culture was inoculated to Landy medium, containing (per liter): glucose 4 g, yeast extract 4 g, malt extract 10 g, and 0.5 ml of 1 M NaOH. It was then cultured for 3 days at 30 °C and 220 rpm.

BAE and its analogues were analyzed on a Waters Ultra Performance Liquid Chromatography (UPLC) system equipped with a photo-diode array (PDA) detector, using a C-18 column (2.7 μm, 4.6 mm × 50 mm, Waters). The mobile phase consisted of buffer A (ultrapure H_2_O containing 0.1 % HCOOH and 0.1 % CH_3_CN) and buffer B (chromatographic grade CH_3_CN containing 0.1 % HCOOH) was applied at a flow rate of 0.4 mL min^-1^. A linear gradient program (60 % buffer A and 40 % buffer B to 50 % buffer A and 50 % buffer B for 1 min, 50 % buffer A and 50 % buffer B to 0 % buffer A and 100 % buffer B for 2 mins, followed by 60 % buffer A and 40 % buffer B for 1 min) was applied to detect BAEs at 360 nm.

### Total DNA extraction

Our strain collection consists of strains isolated from various unexplored and underexplored ecological niches. The cultivation of bacterial strains and genomic DNA preparation followed standard protocols (Kieser et al., 2000). The concentrations of genomic DNA samples were estimated using a micro spectrophotometer (Analytic Jena).

### Sequence similarity network analysis of HCS genes in GenBank

#### Data set source

The HCS sequences were downloaded from the NCBI GenBank protein database on September 27, 2018. LnmM (AAN85526.1) was used as a query ID for BlastP search in non-redundant protein sequences database and 6, 712 sequences with > 25% sequence identity with LnmM were obtained.

#### Data set curation

Because some of the 6, 712 HCS sequences are duplicates or share very high sequence similarity, it is necessary to remove those sequences for further bioinformatic analysis. Using HCS sequences from characterized AT-less type I PKSs as a training set, 90% sequence identity was considered as an appropriate threshold. If two HCSs have > 90% sequence identity, they were treated as a single input HCS seqeunce. Using this strategy, the 6,712 HCS sequences were winnowed into 2,250 representative sequences by filtering to a maximum identity of 90% using CD-HIT (Huang et al., 2010).

#### Construction of networks

A database containing the 2,250 representative HCS sequences was built using formatdb command in Blast+. To construct the network, each individual sequence was used as a query to search the whole database using BlastP. The Blast *E*-value was used to define similarity between the query HCS and the hits. An *E*-value threshold of 1.0 × 10^-135^ was established based on the lowest sequence similarity that all of the characterized HCS sequences were grouped together. The edges formed from self-loops between each node itself, and those from duplicates between two nodes were deleted from the network. Finally, the sequence similarity network was visualized using the organic layout in Cytoscape 4.1 (Kohl et al., 2011).

### High-throughput PCR screening of HCS genes from in-house strains

#### Design of the PCR primers

Two conserved regions of amino acids of HCSs were chosen and a pair of degenerated primers were designed according to CODEHOP strategy (Rose et al., 2003). Forward primer: 5’-CCGTTCGGCGGCATG GTCAAGGGNGCNCAYCG-3’, Reverse primer: 5’- GAAGAACTCGGAGGARCANCC GGTNCCRTA-3’ (R=A/G, M=A/C, N=A/T/G/C). PCR reactions were performed in 20 μl reaction mixtures consisting of 200–500 ng of genomic DNA, 10 pmol of each primer, 4 mmol of dNTP, 1 × GC buffer II and 1 U of LA Taq DNA polymerase (TaKaRa). The PCR cycles consisted of 95 °C for 5 min, followed by 10 cycles of denaturation (95 °C for 0.5 min), annealing (65 °C to 55 °C for 0.5 min), and extension (72 °C for 0.3 min), and 25 cycles of denaturation (95 °C for 0.5 min), annealing (55 °C for 0.5 min), and extension (72 °C for 0.3 min), ended after a final extension (72 °C for 10 min). The resultant PCR products were cloned and sequenced.

#### Phylogenetic analysis

Phylogenetic analysis was performed using MEGA 7 software. The phylogenetic tree was constructed using the neighbor-joining method and 1000 bootstrap replications.

### Genome scanning to identify an AT-less type I PKS gene cluster from S. sp. CB01881

The shotgun genome sequence of S. sp. CB01881 was generated on an Illumina HiSeq 2000 instrument (BGI, Shengzhen, China). The data sets consisted of 11,420,954 single 125-bp reads. The assembly revealed 478 sequence contigs, which were combined into 41 scaffolds. Secondary metabolite biosynthetic gene cluster prediction of S. sp. CB01881 was performed using antiSMASH 4.0 (Blin et al., 2017). The identified AT-less type I PKS gene cluster was further analyzed using Blast. The sequencing data of S. sp. CB01881 can be accessed at GenBank under accession number RSFG00000000.

### Survey AT-less type I PKSs in soil

#### Soil DNA isolation

The 18 soil samples at 20 cm depth of soil were collected in various places in China. The geological location for all the samples was listed in Table S192. DNA was extracted from soil using previously published DNA isolation protocol with modifications (Charlop-Powers et al., 2015). Crude environmental DNA was passed through one round of column purification using the Ezup Column Soil DNA Purification Kit (Sangon Biotech, Shanghai, China).

#### PCR amplification and high-throughput sequencing

The HCS genes in the eDNA were amplified using the same PCR condition in PCR screening of HCS genes from in-house strains. The KS gene fragments of the AT-less type I PKS KSs were amplified from the same eDNA samples, using primers MAK1 (5’-GACACSGCSTGY TCBTCGTCG-3’) and MAK3 (5’-CASTTGGTCCTGCCRCGCAGSTTGCC-3’). The following PCR conditions were used (Analytik Jena FlexCycler2): 95 °C for 5 min, followed by 10 cycles of denaturation (95 °C for 0.5 min), annealing (65 °C to 55 °C for 0.5 min), and extension (72 °C for 0.3 min), and 25 cycles of denaturation (95 °C for 0.5 min), annealing (55 °C for 0.5 min), and extension (72 °C for 0.3 min), and a final extension step at 72 °C for 10 min. The primers for HCS and KS were designed to contain a unique 12 bp barcode that allowed simultaneous sequencing different samples in a single run. PCR reactions were examined by agarose gel electrophoresis to determine the concentration and purity of each amplicon. Amplicons of the appropriate size were recovered using SanPrep Column DNA Gel Extraction Kit (Sangon Biotech, Shanghai, China). The purified HCS and KS amplicons were pooled in equal molar ratios and were then sequenced using Illumina MiSeq platform. The resulting fasta files with demultiplexed and quality-filtered sequences for each soil sample were used for downstream analysis.

#### Data analysis

Operational taxonomic units (OTUs) were generated using CD-HIT based on the following thresholds: the HCS nucleotide sequences were clustered with sequence identity cut-off of 90%. The KS nucleotide sequences were first clustered at 97% sequence identity, and the non-redundant sequences were further clustered at 95% sequence identity. The formed OTUs of HCS sequences containing less than 10 reads were removed. The remaining OTUs of HCS sequences were used to do rarefaction analysis based on Chao1 diversity. The OTUs of HCS sequences containing at least 100 reads were selected to analyze high abundance HCS genes in soil samples.

HCS and KS centroid sequences were assigned to AT-less type I PKS gene clusters using BlastX command in Blast+ at *E*-value cutoff of 10^-28^ and 10^-30^, respectively. Based on the BlastX results, the ratio of HCS gene hits in the AT-less type I PKS gene clusters in each soil sample were manually counted and calculated. To compare HCS gene beta diversity of the soil samples, nonmetric multidimensional scaling ordination analysis was performed using R package – vegan. Phylogenetic analysis of HCS gene hits was performed using MEGA 7. The heat map of HCSs or AT-less type I PKS KSs of each soil sample was generated using software HemI (Deng et al., 2014).

## ASSOCIATED CONTENT

### SUPPORTING INFORMATION

The supporting information is available free of charge on the Publication website.

### MAJOR DATASET

The 16S rRNA sequence of *B. amyloliquefaciens* CB02999 (MK351256), The shotgun genome sequence of S. sp. CB01881 (RSFG00000000), The HCS gene fragments of the 13 in-house strains (Bioproject NO: PRJNA506854), and the high-throughput sequencing dataset of HCS and KS domains amplified from the soil DNA (Biorproject NO: PRJNA511514) were submitted to GenBank and may be accessed.

### NOTES

The authors declare no competing financial interest.

## ACKNOWLEDGMENTS

This work was supported by NSFC grants 81473124 (to YH.) and the Chinese Ministry of Education 111 Project B0803420 (to Y.D.).

