## Supplemental data for "A 3-hydroxy-3-methylglutaryl-CoA synthase-based probe for the discovery of the acyltransferase-less type I polyketide synthases"

### Table of Contents


**Fig. S1.** The structures of selected polyketides with $\beta$-branches.................. **S3**

**Fig. S2.** Sequence similarity network analysis of HCSs............................ **S4**

**Fig. S3.** Phylogenetic analysis of ACPs in HCS cassettes ......................... **S5**

**Fig. S4.** Degenerate primers design and cloning of *lnmM* HCS.................. **S6**

**Fig. S5.** The heatmap of HCS genes in soil sample and 13 in-house strains containing HCS genes (CB)...................................................... **S7**

**Fig. S6.** The CB02999 fermentation profiles of BAEs.................................. **S8**

**Fig. S7.** The numbers of HCS groups in soil samples................................. **S9**

**Fig. S8.** The alignment of KS domains of AT-less type I PKSs..................... **S10**

**Tables S1~S10.** Orphan leinamycin-type AT-less type I PKS BGCs with propionyl-*S*-ACP specific HCSs.................................... **S11~S20**

**Tables S11~S27.** Orphan AT-less type I PKS BGCs with propionyl-*S*-ACP specific HCSs.................................................................. **S21~S38**

**Tables S28~S30.** Orphan leinamycin-type AT-less type I PKS BGCs with acetyl-*S*-ACP specific HCSs .......................................... **S39~S41**

**Tables S31~S170.** Orphan AT-less type I PKS BGCs with acetyl-*S*-ACPspecific HCSs....................................... **S42~S182**

**Tables S171~S172.** Orphan canonical type I PKS BGCs............................. **S183~S184**

**Tables S173~S179.** Orphan type II PKS BGCs with HCSs .......................... **S185~S191**

**Tables S180~S189.** Incompele BGCs with HCSs ........................................ **S192~S201**

**Table S190.** Domain and module organization of the *S.* sp. CB01881 AT-less type I PKS BGCs..................................................... **S202**

**Table S191.** Annotation of *S.* sp. CB01881 AT-less gene clusters in comparison of characterized AT-less PKS BGCs..................... **S203**

**Table S192.** The location data of 18 soil samples......................................... **S204**

**Fig. S1.** The structures of selected polyketides with $\beta$-branches. (**A**) polyketides from AT-less type I PKSs: bacillaene, batumin, bongkrekic acid, bryostatin, calyculin, corallopyronin, diaphorin, difficidin, elansolids, etnsngien, guangnanmycin, leinamycin, myxopyronin, myxovirescin, mupirocin, nosperin, onnamide A, oocydin A, patellazole, pederin, phormidolide, pristinamycin, psymberin, SIA7248, thailandamide, thailanstatin, thiomariinol, weishanmycin; (B) polyketides from canonical type I PKSs: curacin, jamacamide, cylindrocyclophane. The $\beta$-branches are highlighted in red.

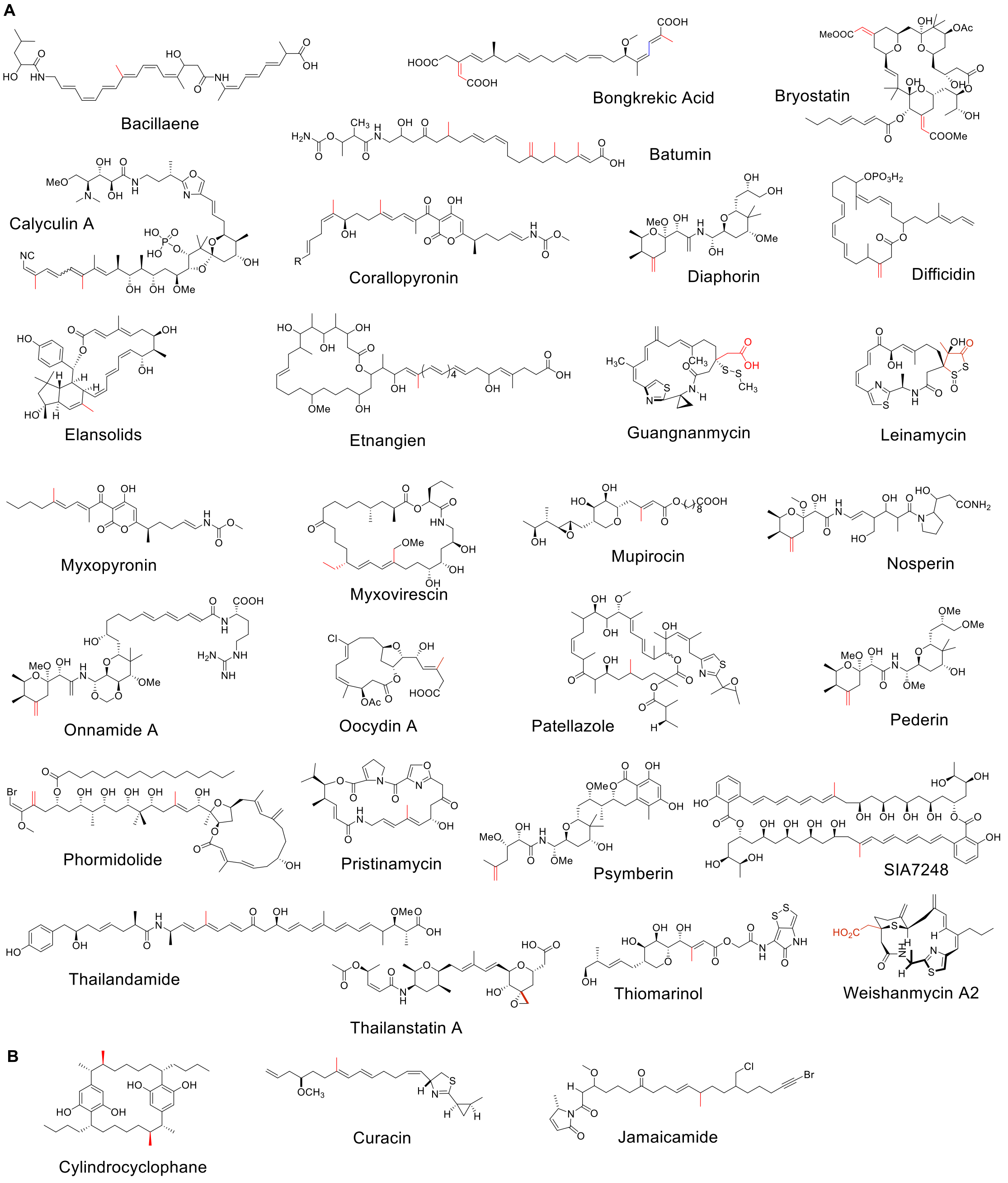

**Fig. S2** Sequence similarity network (SSN) for HCSs. (**a**) SSN of HCS sequences with an *E* value threshold of $10^{-130}$. Eleven HCS sequences, including WP_009943563.1, WP_020673005.1, WP_042180998.1, WP_067233257.1, WP_073838783.1, WP_083822270.1, WP_086782851.1, WP_089099957.1, WP_090099145.1, WP_092536194.1, WP_100840950.1, which were also included into cluster D (The cluster C in the main text). (**b**) SSN of HCSs with an *E* value threshold of $10^{-150}$. Six HCSs AHC73477.1 [PtzI], CDF01048.1, WP_044562062.1, WP_048463408.1, WP_051418953.1 and WP_085216372.1, belonging to AT-less type I PKS BGCs or incomplete gene cluster (WP_048463408.1), were excluded from cluster D.

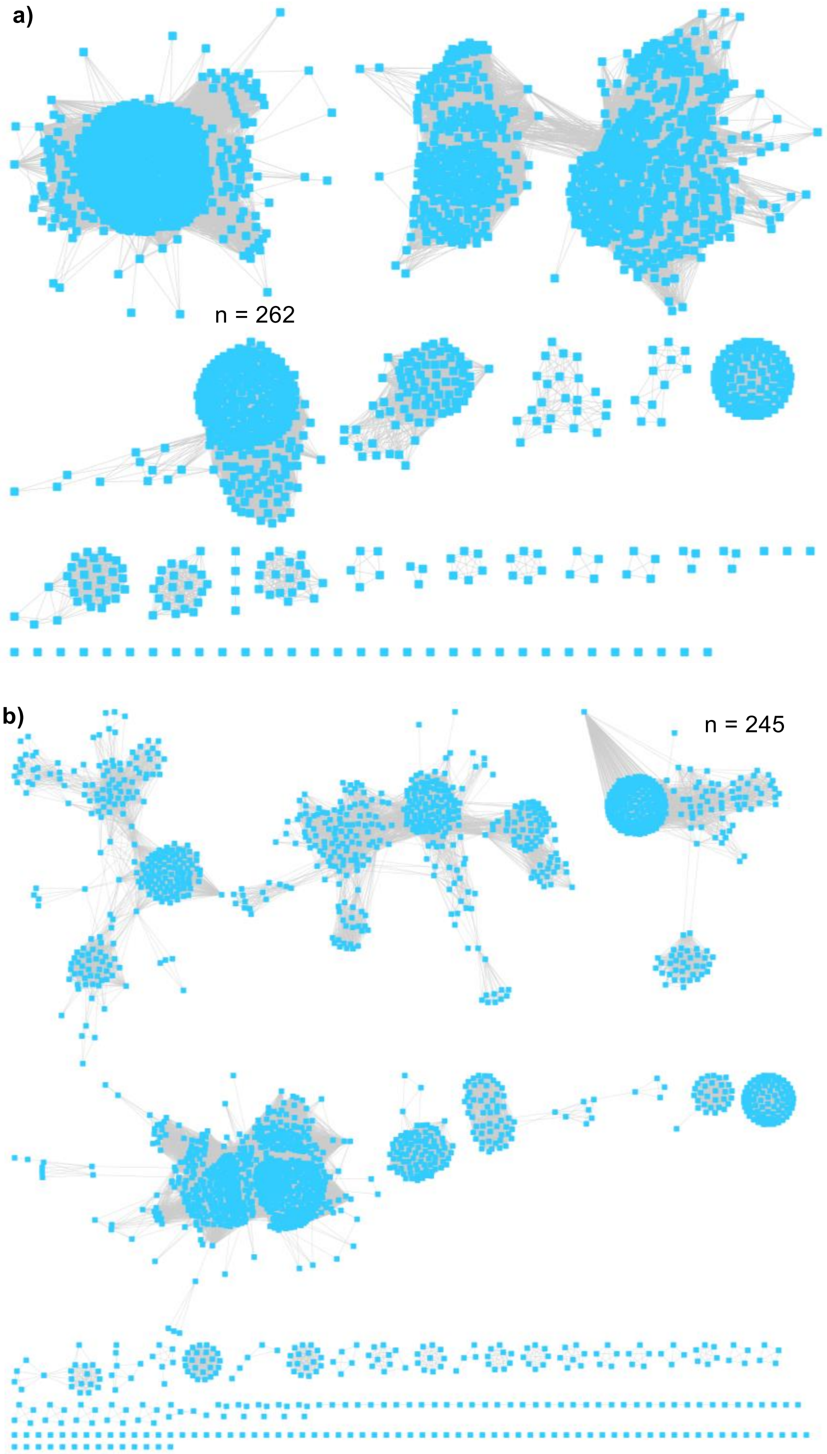

**Fig. S3.** Phylogenetic analysis of the ACP sequences in HCS cassettes. ACPs involved in fatty acid biosynthesis (3EJB_A, 2KOO_A), Type II PKS (Q02054.1, P12884.1), canonical type I PKS (Q03132.3, Q03131.1, CAA60459.1, CAA60460.1, CAA60462.1), acceptor ACPs in AT-less type I PKSs (ABM63537.1, ABF5931.1, AAM12909.2) and BlmI PCP in nonribosomal peptide biosynthesis (EME99231.1) were included as controls. The analysis revealed that the ACPs in HCS cassettes have an independent evolutionary path, compared to those ACPs involving fatty acids synthesis, type I and type II PKSs.

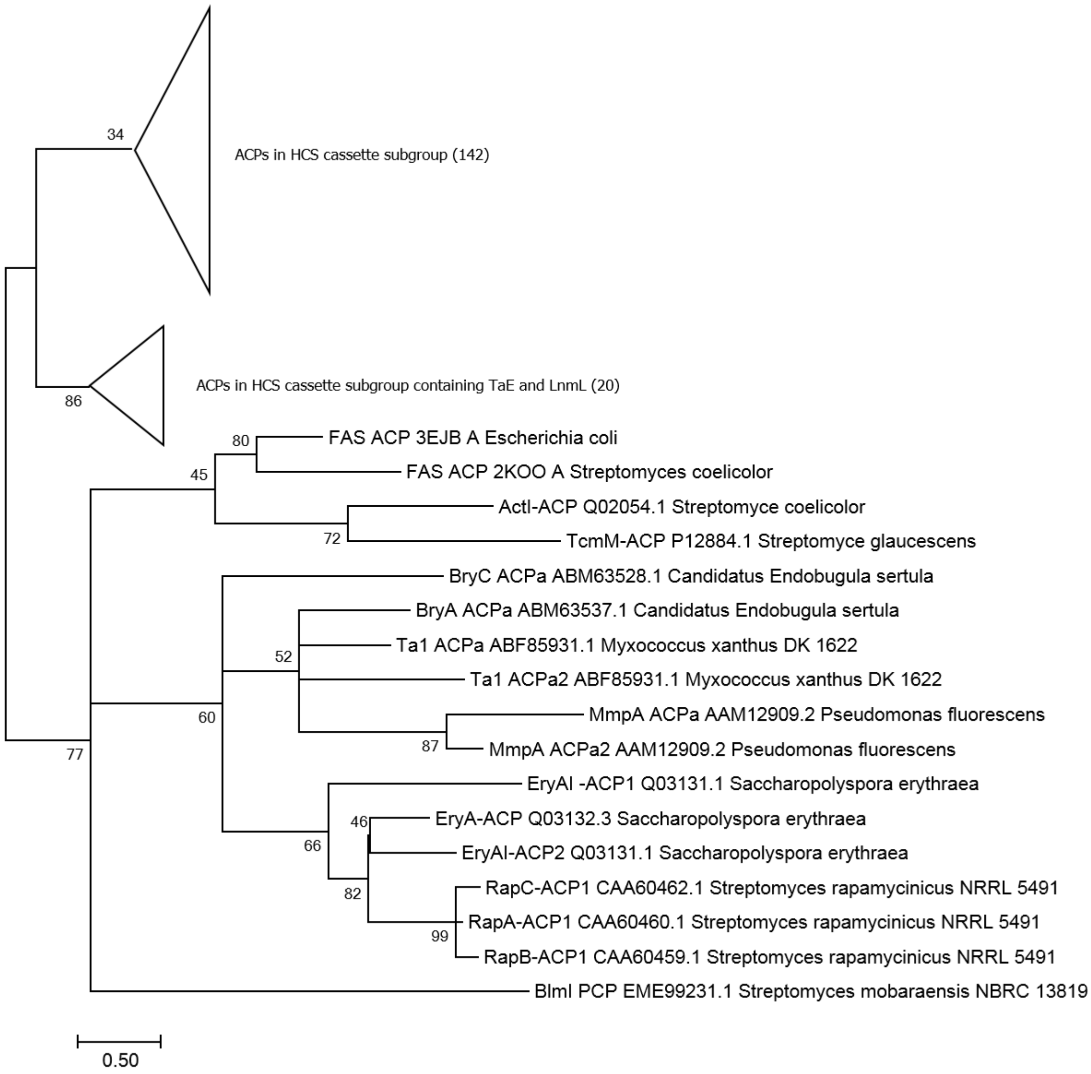

**Fig. S4.** Degenerate primers design and cloning of *lnmM* HCS. (**A**) Sequence alignment of selected HCSs from the characterized AT-less type I PKS BGCs, including BaeG (ABS74058.1), BatC (ADD82944.1), BonG (AFN27479.1), BryR (ABM63533.1), CalT (BAP05578.1), CorE (ADI59527.1), CorE (ADI59527.1), CylF (ARU81120.1), DifL (CAG23983.1), ElaL (AEC04358.1), Fr9K (AIC32697.1), LnmM (AAN85526.1), MupH (AAM12922.1), NosK (WP_094329473.1), OnnA (AAV97869.1), OocE (AFX60327.1), PedP (AAW33975.1), SnaI (CBW45745.1), TaC (CAB46502.1), TaF (CAB46505.1), TmlH (CBK62727.1) and TstK (AGN11885.1). Aligned residues are colored based on the conservation level (grey box shows strict identity and white shows low identity). Primer design of HCS genes using CODHOP strategy for AT-less type I PKS scanning. (**B**) PCR primers designed according to CODEHOPE strategy. (**C**)The amplification of *lnmM* HCS gene from *S. atroolivaceous,* the leinamycin producer. Lane 1, DNA marker; lane 2, the HCS gene PCR product from *S. atroolivaceous*; lane 3, negative control.

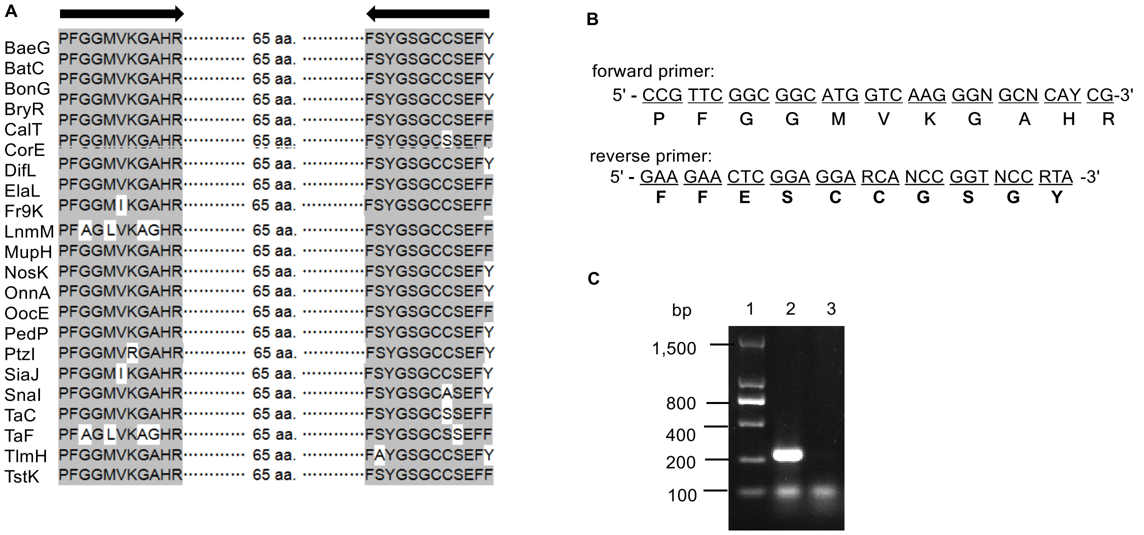

**Fig. S5.** The heatmap of HCS genes in soil samples S1 -S18 and the thirteen in-house strains containing HCS genes (CB). The rows and columns were clustered based on sequence similarity. The colors of the heatmap were determined by the abundance of each HCS genes.

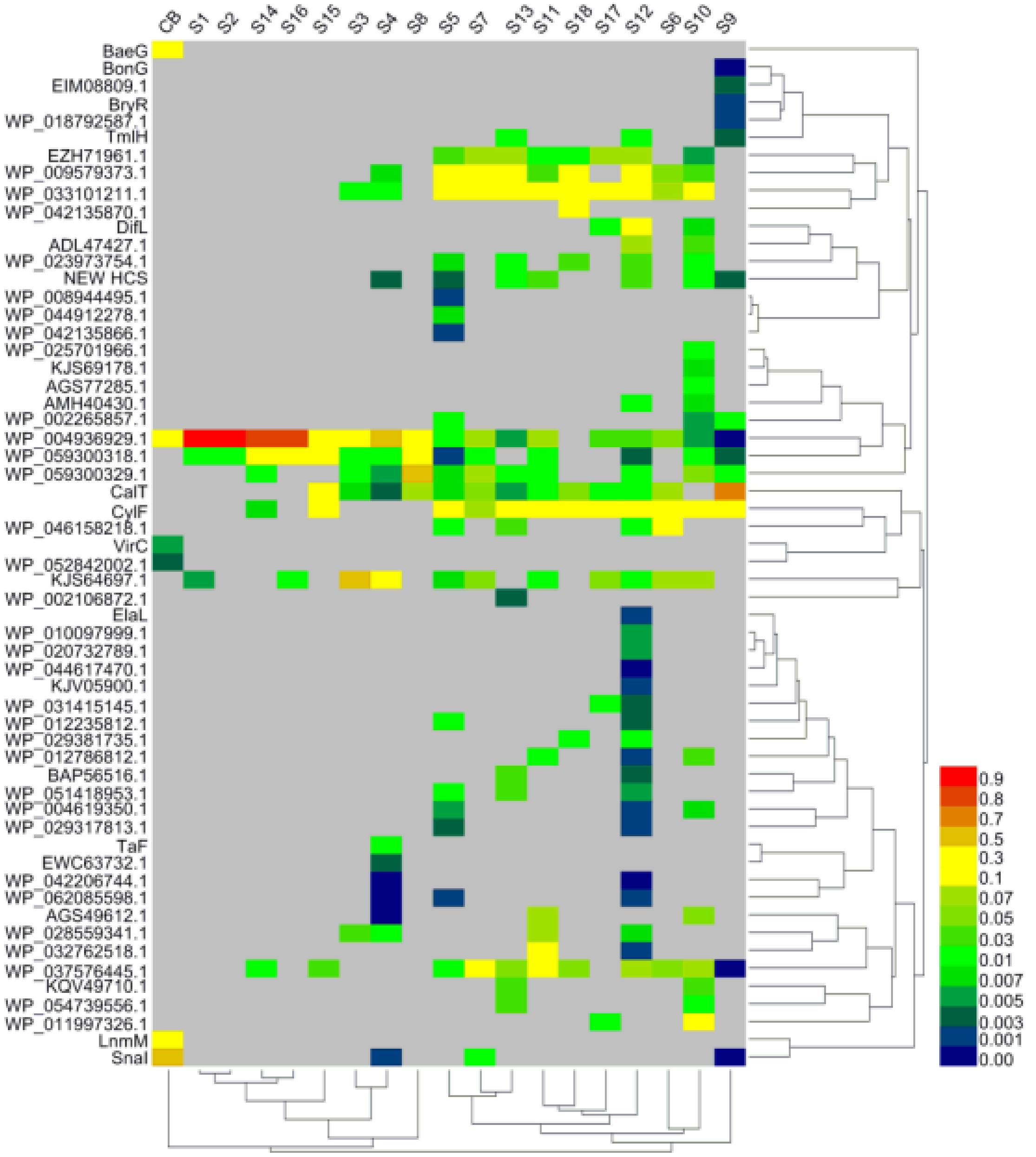

**Fig. S6.** The CB2999 fermentation profiles of BAEs. (**a)** Extracted ion chromatogram of BAE in LCMS. **b)** UV spectrum of BAE. **c)** The HRMS spectrum of BAE. **d)** The MS/MS spectrum of BAE. **e)** Extracted ion chromatogram of dehydro-BAE in LCMS. **f)** UV spectrum of dehydro-BAE. **g)** The HRMS spectrum of dehydro-BAE. **h)** The MS/MS spectrum of dehydro-BAE. **i)** Extracted ion chromatogram of BAE B in LCMS. **j)** UV spectrum of BAE B. **k)** The HRMS spectrum of BAE B. **l)** The MS/MS spectrum of BAE B. **m)** Extracted ion chromatogram of dehydro-BAE B in LCMS. **n)** UV spectrum of dehydro-BAE B. **o)** The HRMS spectrum of dehydro-BAE B.

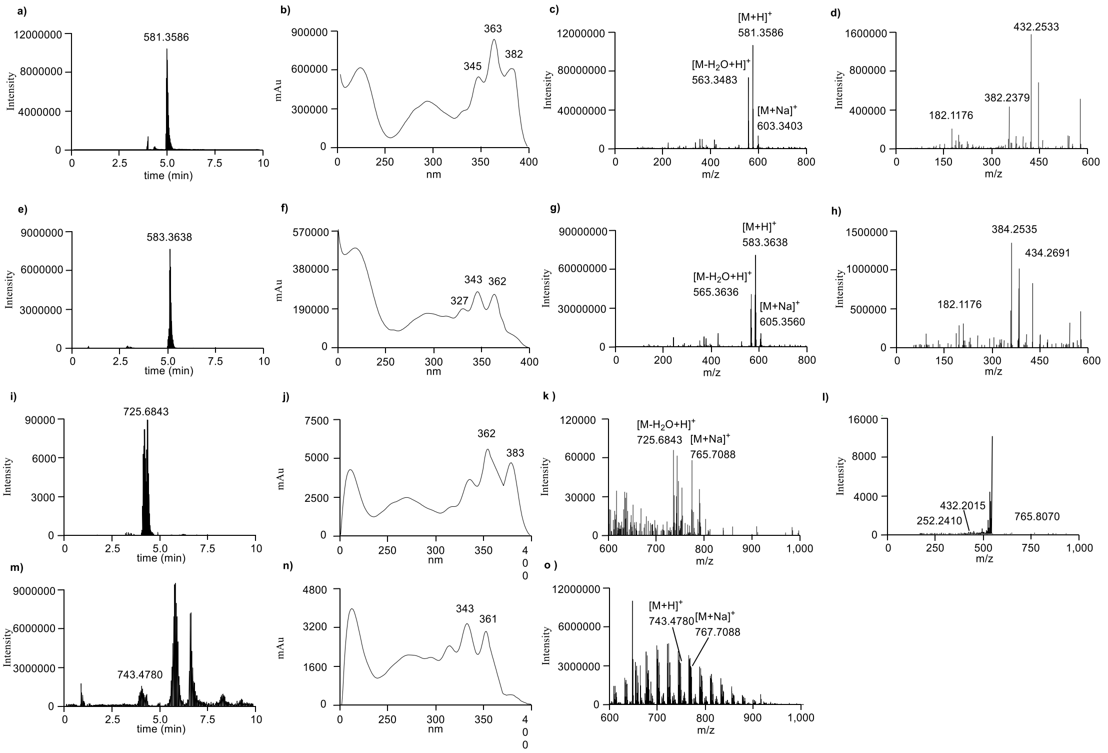

**Fig. S7.** The numbers of HCSs in each soil sample. The HCSs were clustered from HCS OTUs containing over 100 reads of each soil sample (S1-S18) and the mixture of genomic DNA from the thirteen in-house strains (CB).

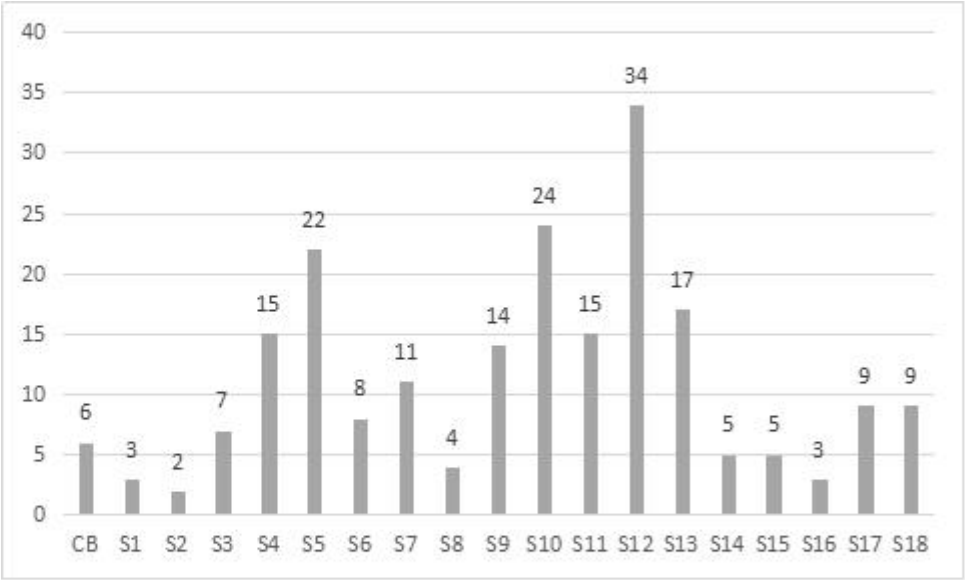

**Fig. S8.** The alignment of KS domains of AT-less type I PKSs. The primers were designed based on the conserved regions “DTACSSSLVALH” and “SAVNQDGASNG” (See main text for the reference).

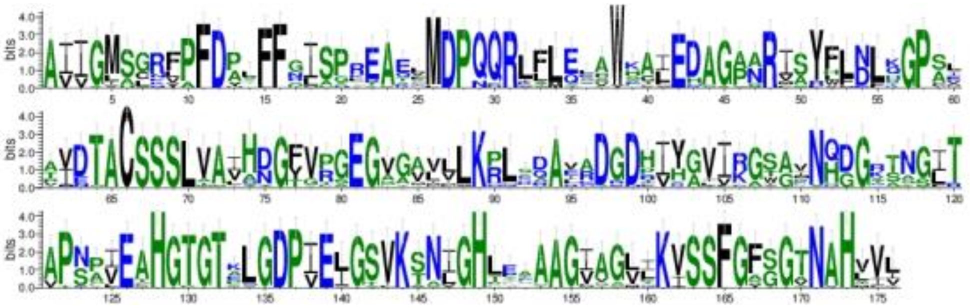

**Table S1.** Predicted functions of ORFs in the CP002162.1 containing ADL47416.1 and ADL47427.1

| gene | aa[a] | putative function | Protein homologue | %identity/ %similarity |
|---|---|---|---|---|
| ADL47392.1 | 644 | radical SAM domain protein | / | / |
| ADL47393.1 | 170 | methyltransferase type 11 | / | / |
| ADL47394.1 | 121 | glyoxalase/bleomycin resistance protein | JamB [AAS98775.1] | 45/64 |
| ADL47395.1 | 136 | hypothetical protein | JamA [AAS98774.1] | 45/64 |
| ADL47396.1 | 518 | ABC transporter | JamC [AAS98798.1] | 31/62 |
| ADL47397.1 | 263 | transport systems | LnmS [AAN85532.1] | 60/71 |
| ADL47398.1 | 318 | transport systems | LnmT [AAN85533.1] | 59/72 |
| ADL47399.1 | 519 | extracellular solute-binding protein family | LnmU [AAN85534.1] | 53/65 |
| ADL47400.1 | 394 | monooxygenase | / | / |
| ADL47401.1 | 412 | beta-ketoacyl synthase | CorD [ADI59526.1] | 48/61 |
| ADL47402.1 | 76 | acyl carrier protein | PsyL [ADA82592.1] | 35/67 |
| ADL47403.1 | 321 | chlorinating enzyme | / | / |
| ADL47404.1 | 621 | NRPS | kirromycin [CAN89656.1] | 46/55 |
| ADL47405.1 | 313 | hypothetical protein | LnmH [AAN85521.1] | 44/58 |
| ADL47406.1 | 767 | AT/Ox | DifA [CAG23974.1] | 45/63 |
| ADL47407.1 | 2174 | NRPS | LnmI [AAN85522.1] | 48/58 |
| ADL47408.1 | 246 | oleoyl-(acyl carrier protein) hydrolase | LnmN [AAN85527.1] | 55/63 |
| ADL47409.1 | 453 | aminotransferase class-III | Malleilactone [ABC35627.1] | 33/51 |
| ADL47410.1 | 412 | cytochrome P450 | BaeS [CAG23962.1] | 38/58 |
| ADL47411.1 | 328 | ABC transporter ATPase subunit | Rhizopodin [CCA89322.1] | 31/49 |
| ADL47412.1 | 266 | ABC-2 type transporter | / | / |
| ADL47413.1 | 418 | hypothetical protein | / | / |
| ADL47414.1 | 32 | hypothetical protein | / | / |
| ADL47415.1 | 397 | alpha/beta hydrolase fold | MupV [AAM12938.1] | 31/47 |
| ADL47416.1 | 411 | HMG-CoA synthase | TaF [ABF92623.1] | 62/75 |
| ADL47417.1 | 82 | acyl carrier protein | LnmL [AAN85525.1] | 59/71 |
| ADL47418.1 | 311 | AT/DC | LnmK [AAN85524.1] | 55/67 |
| ADL47419.1 | 6727 | AT-less type I PKS | CorL [ADI59534.1] | 41/51 |
| ADL47420.1 | 2005 | AT-less type I PKS | ChiD [AAY89051.1] | 41/52 |
| ADL47421.1 | 292 | hypothetical protein | LnmE [AAN85518.1] | 43/60 |
| ADL47422.1 | 252 | enoyl-CoA hydratase | JamI [AAS98780.1] | 40/58 |
| ADL47423.1 | 735 | AT/Ox | ChiA [AAY89048.1] | 37/50 |
| ADL47424.1 | 253 | cyclic nucleotide-binding | LnmO [AAN85528.1] | 46/59 |
| ADL47425.1 | 426 | sodium/hydrogen exchanger | / | / |
| ADL47426.1 | 179 | protein of unknown function DUF1697 | Leinamycin [AAN85541.1] | 32/47 |
| ADL47427.1 | 413 | HMG-CoA synthase | CorE [ADI59527.1] | 55/68 |
| ADL47428.1 | 494 | long-chain fatty acid--CoA ligase | LnmW [AAN85536.1] | 42/57 |

[a] Number of amino acids

**Table S2.** Predicted functions of ORFs in the FMCQ01000002.1 containing SCE73423.1

| gene | aa [a] | putative function | Protein homologue | %identity/ %similarity |
|---|---|---|---|---|
| SCE73331.1 | 379 | alpha/beta fold hydrolase | MupV [AAM12938.1] | 27/40 |
| SCE73340.1 | 244 | cAMP-dependent protein kinases | LnmO [AAN85528.1] | 50/66 |
| SCE73351.1 | 306 | hypothetical protein | LnmE [AAN85518.1] | 49/61 |
| SCE73357.1 | 257 | enoyl-CoA hydratase | LnmF [AAN85519.1] | 53/64 |
| SCE73369.1 | 778 | AT/Ox | DifA [CAG23974.1] | 48/66 |
| SCE73373.1 | 314 | hypothetical protein | LnmH [AAN85521.1] | 44/61 |
| SCE73388.1 | 4285 | hybrid PKS/NRPS | LnmI [AAN85522.1] | 50/60 |
| SCE73393.1 | 7078 | AT-less type I PKS | LnmJ [AAN85523.1] | 50/60 |
| SCE73405.1 | 329 | AT/DC | LnmK [AAN85524.1] | 55/70 |
| SCE73412.1 | 86 | acyl carrier protein | LnmL [AAN85525.1] | 58/73 |
| SCE73423.1 | 413 | HMG-CoA synthase | LnmM [AAN85526.1] | 65/79 |
| SCE73435.1 | 250 | thioesterase | LnmN [AAN85527.1] | 55/65 |
| SCE73448.1 | 134 | hypothetical protein | / | / |
| SCE73453.1 | 476 | oxidase EvaA | / | / |
| SCE73466.1 | 333 | dehydrogenase | / | / |

[a] Number of amino acids

**Table S3**. Predicted functions of ORFs in the NZ_LN831790.1 containing WP_029381735.1 and WP_029382854.1

| gene | aa[a] | putative function | Protein homologue | %identity/ %similarity |
|---|---|---|---|---|
| WP_029381741.1 | 396 | cytochrome P450 | Dor11 [ACY01396.1] | 45/61 |
| WP_078648056.1 | 144 | DUF3224 domain | LnmZ [AAN85539.1] | 34/48 |
| WP_029381739.1 | 139 | nuclear transport factor 2 | LnmV [AAN85535.1] | 43/61 |
| WP_029381738.1 | 260 | thioesterase | LnmN [AAN85527.1] | 52/64 |
| WP_029381737.1 | 424 | beta-ketoacyl synthase | CorD [ADI59526.1] | 47/60 |
| WP_078648055.1 | 99 | acyl carrier protein | PsyL [ADA82592.1] | 43/66 |
| WP_029381735.1 | 410 | HMG-CoA synthase | LnmM [AAN85526.1] | 64/76 |
| WP_029381734.1 | 86 | acyl carrier protein | LnmL [AAN85525.1] | 50/67 |
| WP_047121549.1 | 327 | AT/DC | LnmK [AAN85524.1] | 52/64 |
| WP_047122366.1 | 6540 | AT-less type I PKS | LnmJ [AAN85523.1] | 50/61 |
| WP_049976752.1 | 4258 | NRPS | LnmI [AAN85522.1] | 49/59 |
| WP_049976753.1 | 387 | alpha/beta fold hydrolase | MupV [AAM12938.1] | 28/40 |
| WP_029382850.1 | 316 | beta-ketoacyl synthase | MxnB [AGS77282.1] | 27/37 |
| WP_047121551.1 | 626 | NRPS | Kirromycin [CAN89656.1] | 48/59 |
| WP_047121552.1 | 337 | hypothetical protein | LnmH [AAN85521.1] | 44/60 |
| WP_047121553.1 | 823 | AT/Ox | MmpC [AAM12912.1] | 48/64 |
| WP_047122367.1 | 261 | enoyl-CoA hydratase | LnmF [AAN85519.1] | 50/63 |
| WP_029382854.1 | 411 | HMG-CoA synthase | MxnE [AGS77285.1] | 53/68 |
| WP_078648135.1 | 563 | class I SAM-dependent methyltransferase | KirM [CAN89643.1] | 32/44 |
| WP_049976754.1 | 572 | long-chain fatty acid--CoA ligase | LnmW [AAN85536.1] | 41/56 |
| WP_047121555.1 | 230 | TetR/AcrR family transcriptional regulator | Leinamycin [AAN85486.1] | 37/59 |
| WP_078648140.1 | 465 | nuclear transport factor 2 | LnmV [AAN85535.1] | 42/59 |
| WP_047121556.1 | 404 | glycosyltransferase | CylN [ARU81128.1] | 25/42 |
| WP_078648136.1 | 382 | winged helix DNA-binding domain | BryD [ABM63530.1] | 36/68 |

[a] Number of amino acids

**Table S4.** Predicted functions of ORFs in the NZ_JNWQ01000013.1 containing WP_030056132.1 and WP_030056101.1

| gene | aa[a] | putative function | Protein homologue | %identity/ %similarity |
|---|---|---|---|---|
| WP_030056128.1 | 314 | daunorubicin resistance protein | LnmR [AAN85531.1] | 33/48 |
| WP_030056129.1 | 266 | hypothetical protein | / | / |
| WP_051829787.1 | 258 | thioesterase | LnmN [AAN85527.1] | 48/60 |
| WP_030257036.1 | 405 | beta-ketoacyl synthase | CorD [ADI59526.1] | 48/60 |
| WP_051829788.1 | 82 | acyl carrier protein | acpK [CAG23953.1] | 45/71 |
| WP_030056132.1 | 408 | HMG-CoA synthase | TaF [ABF92623.1] | 56/69 |
| WP_033331212.1 | 7724 | AT-less type I PKS | MxnK [AGS77291.1] | 40/50 |
| WP_078588637.1 | 2878 | NRPS | LnmI [AAN85522.1] | 48/57 |
| WP_051829790.1 | 596 | FAD-dependent oxidoreductase | Ena5928 [ABI91473.1] | 31/42 |
| WP_051829791.1 | 803 | class-III aminotransferase | BatP [ADD82957.1] | 33/47 |
| WP_051829792.1 | 798 | AT-less type I PKS | JamK [AAS98782.1] | 39/52 |
| WP_051829793.1 | 973 | AT-less type I PKS | JamL [AAS98783.1] | 38/51 |
| WP_051829794.1 | 82 | acyl carrier protein | SorA [ADN68476.1] | 43/58 |
| WP_051829795.1 | 564 | AT-less type I PKS | JamA [AAS98774.1] | 38/52 |
| WP_078588640.1 | 520 | AT-less type I PKS | / | / |
| WP_030056087.1 | 341 | hypothetical protein | / | / |
| WP_030056088.1 | 495 | AMP-dependent synthetase | SgvE4 [AGN74895.1] | 31/45 |
| WP_030056089.1 | 406 | decarboxylase | Leinamycin [AAN85498.1] | 26/41 |
| WP_030257054.1 | 83 | peptidyl carrier protein | LnmP [AAN85529.1] | 37/50 |
| WP_030257056.1 | 542 | alpha/beta hydrolase | LnmD [AAN85517.1] | 40/52 |
| WP_051829798.1 | 223 | Crp/Fnr family transcriptional regulator | LnmO [AAN85528.1] | 41/59 |
| WP_030056093.1 | 70 | ferredoxin | / | / |
| WP_030056094.1 | 402 | cytochrome P450 | Dor11 [ACY01396.1] | 43/59 |
| WP_030056095.1 | 404 | cytochrome P450 | ChxI [AFO59870.1] | 47/61 |
| WP_051829799.1 | 150 | hypothetical protein | / | / |
| WP_030056097.1 | 193 | hypothetical protein | / | / |
| WP_051829800.1 | 281 | hypothetical protein | LnmH [AAN85521.1] | 41/58 |
| WP_030056099.1 | 764 | AT/Ox | MmpC [AAM12912.1] | 49/66 |
| WP_106973353.1 | 262 | enoyl-CoA hydratase | JamI [AAS98780.1] | 38/57 |
| WP_030056101.1 | 408 | HMG-CoA synthase | ThaK [ABC34601.1] | 49/63 |

[a] Number of amino acids

**Table S5.** Predicted functions of ORFs in the NZ_BBNN01000015.1 containing WP_042163761.1 and WP_042163784.1

| gene | aa[a] | putative function | Protein homologue | %identity/ %similarity |
|---|---|---|---|---|
| WP_052479722.1 | 237 | Crp/Fnr family transcriptional regulator | LnmO [AAN85528.1] | 37/58 |
| WP_042163759.1 | 189 | hypothetical protein | / | / |
| WP_052479723.1 | 244 | PPTase | BatI [ADD82950.1] | 38/50 |
| WP_042163761.1 | 409 | HMG-CoA synthase | JamH [AAS98779.1] | 45/65 |
| WP_042163967.1 | 255 | enoyl-CoA hydratase | MxnF [AGS77286.1] | 39/52 |
| WP_052479724.1 | 290 | ACP S-malonyltransferase | MisG [AKQ22695.1] | 35/54 |
| WP_052479725.1 | 1993 | AT-less type I PKS | LnmI [AAN85522.1] | 47/56 |
| WP_042163764.1 | 274 | transposase | / | / |
| WP_042163766.1 | 131 | VOC family protein | / | / |
| WP_042163769.1 | 524 | MFS transporter | SnbR [CBW45761.1] | 31/44 |
| WP_052479727.1 | 233 | hypothetical protein | LnmE [AAN85518.1] | 38+54 |
| WP_052479728.1 | 776 | AT/Ox | DifA [CAG23974.1] | 46/64 |
| WP_042163770.1 | 307 | hypothetical protein | LnmH [AAN85521.1] | 35/52 |
| WP_042163773.1 | 120 | VOC family protein | / | / |
| WP_042163774.1 | 236 | N-acetylglucosaminyl deacetylase | LnmX [AAN85537.1] | 50/62 |
| WP_042163775.1 | 249 | methyltransferase | KirM [CAN89642.1] | 42/53 |
| WP_042163776.1 | 418 | alpha/beta hydrolase | LnmD [AAN85517.1] | 36/46 |
| WP_042163778.1 | 81 | peptidyl carrier protein | LnmP [AAN85529.1] | 45/60 |
| WP_042163780.1 | 525 | discrete adenylation domain | LnmQ [AAN85530.1] | 52/62 |
| WP_042163782.1 | 7438 | AT-less type I PKS | LnmJ [AAN85523.1] | 43/54 |
| WP_042163784.1 | 416 | HMG-CoA synthase | LnmM [AAN85526.1] | 56/69 |
| WP_042163787.1 | 81 | acyl carrier protein | ElaE [AEC04351.1] | 37/53 |
| WP_042163976.1 | 403 | beta-ketoacyl synthase | CorD [ADI59526.1] | 45/55 |
| WP_042163789.1 | 248 | thioesterase | LnmN [AAN85527.1] | 55/64 |
| WP_052479730.1 | 2337 | NRPS | LnmI [AAN85522.1] | 45/54 |
| WP_052479731.1 | 143 | hypothetical protein | / | / |
| WP_042163792.1 | 408 | cytochrome P450 | Dor11 [ACY01396.1] | 41/57 |
| WP_042163795.1 | 74 | ferredoxin | LnmB [AAN85516.1] | 31/51 |
| WP_042163797.1 | 453 | crotonyl-CoA carboxylase/reductase | kirN [CAN89653.1] | 51/63 |
| WP_042163798.1 | 330 | beta-ketoacyl synthase | MxnB [AGS77282.1] | 29/43 |
| WP_042163799.1 | 289 | 3-hydroxybutyryl-CoA dehydrogenase | OzmG [ABS90469.1] | 30/56 |
| WP_042163800.1 | 815 | ABC transporter permease | / | / |

[a] Number of amino acids

**Table S6.** Predicted functions of ORFs in the NZ_NNBP01000008.1 containing WP_100602376.1

| gene | aa [a] | putative function | Protein homologue | %identity/ %similarity |
|---|---|---|---|---|
| WP_100602431.1 | 272 | hypothetical protein | / | / |
| WP_100602432.1 | 152 | hypothetical protein | / | / |
| WP_100602366.1 | 402 | cytochrome P450 | SgvP [AGN74891.1] | 51/64 |
| WP_100602367.1 | 247 | N-acetylglucosaminyl deacetylase | LnmX [AAN85537.1] | 54/66 |
| WP_100602368.1 | 446 | alpha/beta hydrolase | LnmD [AAN85517.1] | 62/68 |
| WP_100602369.1 | 271 | enoyl-CoA hydratase | LnmF [AAN85519.1] | 82/88 |
| WP_100602370.1 | 762 | AT/Ox | DifA [CAG23974.1] | 49/67 |
| WP_100602371.1 | 280 | hypothetical protein | LnmH [AAN85521.1] | 83/89 |
| WP_100602372.1 | 4391 | hybrid PKS/NRPS | LnmI [AAN85522.1] | 68/74 |
| WP_100602373.1 | 7490 | AT-less type I PKS | LnmJ [AAN85523.1] | 68/75 |
| WP_100602374.1 | 328 | AT/DC | LnmK [AAN85524.1] | 74/83 |
| WP_100602375.1 | 86 | acyl carrier protein | LnmL [AAN85525.1] | 76/81 |
| WP_100602376.1 | 419 | HMG-CoA synthase | LnmM [AAN85526.1] | 83/88 |
| WP_100602377.1 | 264 | thioesterase | LnmN [AAN85527.1] | 68/77 |
| WP_100602378.1 | 228 | Crp/Fnr family transcriptional regulator | LnmO [AAN85528.1] | 76/85 |
| WP_100602379.1 | 377 | DUF1205 domain-containing protein | / | / |
| WP_100602380.1 | 87 | peptidyl carrier protein | LnmP [AAN85529.1] | 61/68 |
| WP_100602381.1 | 517 | discrete adenylation domain | LnmQ [AAN85530.1] | 76/81 |
| WP_100602433.1 | 538 | ABC transporter ATP-binding protein | LnmR [AAN85531.1] | 70/78 |
| WP_100602382.1 | 289 | ABC transporter permease | LnmS [AAN85532.1] | 85/90 |
| WP_100602383.1 | 331 | ABC transporter permease | LnmT [AAN85533.1] | 81/87 |
| WP_100602384.1 | 514 | ABC transporter binding protein | LnmU [AAN85534.1] | 75/84 |
| WP_100602385.1 | 305 | thymidylyltransferase | marinomycin [BAG50457.1] | 56/73 |
| WP_100602434.1 | 307 | mycothiol conjugate amidase | leinamycin [AAN85543.1] | 69/77 |
| WP_100602386.1 | 634 | HAD-IIIC family phosphatase | tartrolon [ACR13997.1] | 35/50 |
| WP_100602387.1 | 524 | polyketide oxidase | / | / |

[a] Number of amino acids

**Table S7.** Predicted functions of ORFs in the NZ_BEWA01000011.1 containing WP_109002541.1 and WP_109002556.1

| gene | aa [a] | putative function | Protein homologue | %identity/ %similarity |
|---|---|---|---|---|
| WP_109002533.1 | 187 | hemerythrin | / | / |
| WP_109002534.1 | 530 | FAD-binding monooxygenase | / | / |
| WP_109002535.1 | 208 | TetR/AcrR family transcriptional regulator | / | / |
| WP_109002536.1 | 423 | hypothetical protein | / | / |
| WP_109002537.1 | 229 | Crp/Fnr family transcriptional regulator | LnmO [AAN85528.1] | 44/62 |
| WP_109002538.1 | 435 | aminotransferase | / | / |
| WP_109002539.1 | 413 | cytochrome P450 | Dor11 [ACY01396.1] | 44/58 |
| WP_109002575.1 | 423 | beta-ketoacyl synthase | CorD [ADI59526.1] | 47/58 |
| WP_109002540.1 | 91 | PPTase | PedN [AAW33973.1] | 44/74 |
| WP_109002541.1 | 416 | HMG-CoA synthase | LnmM [AAN85526.1] | 60/75 |
| WP_109002542.1 | 7216 | AT-less type I PKS | CorL [ADI59534.1] | 40/52 |
| WP_109002543.1 | 542 | discrete adenylation domain | LnmQ [AAN85530.1] | 54/64 |
| WP_109002544.1 | 153 | peptidyl carrier protein | LnmP [AAN85529.1] | 47/59 |
| WP_109002545.1 | 792 | alpha/beta hydrolase | LnmD [AAN85517.1] | 44/56 |
| WP_109002546.1 | 248 | SAM-dependent methyltransferase | KirM [CAN89643.1] | 44/52 |
| WP_109002547.1 | 238 | N-acetylglucosaminyl deacetylase | LnmX [AAN85537.1] | 54/68 |
| WP_109002548.1 | 135 | DUF3224 family protein | / | / |
| WP_109002549.1 | 121 | VOC family protein | / | / |
| WP_109002550.1 | 310 | hypothetical protein | LnmH [AAN85521.1] | 39/55 |
| WP_109002551.1 | 796 | AT/ER | MxnA [AGS77281.1] | 48/63 |
| WP_109002552.1 | 299 | hypothetical protein | LnmE [AAN85518.1] | 45/60 |
| WP_109002576.1 | 489 | DHA2 family efflux MFS transporter permease | LnmY [AAN85538.1] | 31/50 |
| WP_109002553.1 | 2096 | AT-less type I PKS | LnmI [AAN85522.1] | 52/61 |
| WP_109002554.1 | 803 | AT/ER | MxnA [AGS77281.1] | 34/45 |
| WP_109002555.1 | 263 | enoyl-CoA hydratase | JamI [AAS98780.1] | 36/56 |
| WP_109002556.1 | 410 | HMG-CoA synthase | MxnE [AGS77285.1] | 56/67 |
| WP_109002577.1 | 314 | PPTase | MupN [AAM12928.1] | 37/47 |
| WP_109002557.1 | 252 | thioesterase | LnmN [AAN85527.1] | 52/64 |
| WP_109002578.1 | 2346 | NRPS | LnmI [AAN85522.1] | 47/56 |
| WP_109002558.1 | 145 | globin | / | / |
| WP_109002559.1 | 195 | hypothetical protein | / | / |
| WP_109002579.1 | 125 | hypothetical protein | / | / |

[a] Number of amino acids

**Table S8.** Predicted functions of ORFs in the NZ_CP011664.1 containing WP_008743704.1 and WP_037796927.1

| gene | aa[a] | putative function | Protein homologue | %identity/ %similarity |
|---|---|---|---|---|
| WP_047960902.1 | 248 | transcriptional regulator | / | / |
| WP_037796918.1 | 1306 | NRPS | kirromycin [CAN89656.1] | 37/49 |
| WP_037796920.1 | 475 | M1 family peptidase | / | / |
| WP_106433445.1 | 294 | 3-hydroxybutyryl-CoA dehydrogenase | OzmG [ABS90469.1] | 35/50 |
| WP_008744144.1 | 330 | beta-ketoacyl synthase | MxnB [AGS77282.1] | 28/44 |
| WP_008744143.1 | 452 | crotonyl-CoA carboxylase/reductase | kirN [CAN89653.1] | 49/62 |
| WP_037795615.1 | 65 | ferredoxin | LnmB [AAN85516.1] | 36/54 |
| WP_008744141.1 | 408 | cytochrome P450 | Dor11 [ACY01396.1] | 41/56 |
| WP_008744140.1 | 143 | globin | / | / |
| WP_053065722.1 | 2250 | hybrid AT-less type I PKS/NRPS | LnmI [AAN85522.1] | 45/54 |
| WP_047960426.1 | 251 | thioesterase | LnmN [AAN85527.1] | 55/62 |
| WP_053065723.1 | 400 | beta-ketoacyl synthase | CorD [ADI59526.1] | 43/55 |
| WP_037796925.1 | 81 | acyl carrier protein | OocG [AFX60329.1] | 36/63 |
| WP_037796927.1 | 414 | HMG-CoA synthase | LnmM [AAN85526.1] | 58/71 |
| WP_047960427.1 | 7406 | AT-less type I PKS | LnmJ [AAN85523.1] | 44/54 |
| WP_053065724.1 | 532 | discrete adenylation domain | LnmQ [AAN85530.1] | 52/62 |
| WP_107082496.1 | 130 | peptidyl carrier protein | LnmP [AAN85529.1] | 46/68 |
| WP_107082497.1 | 420 | alpha/beta hydrolase | LnmD [AAN85517.1] | 38/49 |
| WP_047960428.1 | 425 | class I SAM-dependent methyltransferase | KirM [CAN89642.1] | 43/55 |
| WP_073794140.1 | 236 | N-acetylglucosaminyl deacetylase | LnmX [AAN85537.1] | 50/60 |
| WP_008744093.1 | 121 | VOC family protein | / | / |
| WP_047960429.1 | 302 | hypothetical protein | LnmH [AAN85521.1] | 38/54 |
| WP_008744095.1 | 775 | AT/Ox | DifA [CAG23974.1] | 46/62 |
| WP_050779435.1 | 233 | hypothetical protein | LnmE [AAN85518.1] | 37/52 |
| WP_037795575.1 | 521 | MFS transporter | AlbXIV [CAE52325.1] | 31/47 |
| WP_053065726.1 | 2155 | AT-less type I PKS | LnmI [AAN85522.1] | 45/54 |
| WP_008743702.1 | 290 | acyltransferase/oxidoreductase | LnmG [AAN85520.1] | 42/54 |
| WP_008743703.1 | 265 | enoyl-CoA hydratase | CorF [ADI59528.1] | 39/54 |
| WP_008743704.1 | 409 | HMG-CoA synthase | MxnE [AGS77285.1] | 51/65 |
| WP_078489253.1 | 275 | PPTase | BatI [ADD82950.1] | 36/50 |
| WP_037795280.1 | 190 | hypothetical protein | / | / |
| WP_078489254.1 | 307 | Crp/Fnr family transcriptional regulator | LnmO [AAN85528.1] | 38/57 |
| WP_008743708.1 | 294 | methyltransferase domain | DipM [AGS06826.1] | 25/42 |
| WP_047960430.1 | 426 | 2-methylisoborneol synthase | / | / |

[a] Number of amino acids

**Table S9.** Predicted functions of ORFs in the NC_013131.1 containing WP_012786806.1 and WP_012786812.1

| gene[a] | aa[b] | putative function | Protein homologue | %identity/ %similarity |
|---|---|---|---|---|
| WP_085953917.1 | 399 | monooxygenase FAD-binding | / | / |
| WP_012786789.1 | 483 | amidase | SorP [ADN68490.1] | 35/48 |
| WP_012786790.1 | 503 | long-chain fatty acid--CoA ligase | LnmW [AAN85536.1] | 33/45 |
| WP_012786791.1 | 539 | MFS transporter | Leinamycin [AAN85487.1] | 29/47 |
| WP_012786792.1 | 360 | FAD-dependent pyridine nucleotide-disulfide oxidoreductase | / | / |
| WP_083795694.1 | 450 | cytochrome P450 | Dor11 [ACY01396.1] | 43/59 |
| WP_012786794.1 | 308 | hypothetical protein | LnmH [AAN85521.1] | 43/60 |
| WP_012786795.1 | 132 | DUF3224 domain | LnmZ' [AAN85540.1] | 34/49 |
| WP_012786796.1 | 179 | DUF1697 domain | / | / |
| WP_012786797.1 | 80 | ferredoxin | / | / |
| WP_012786798.1 | 865 | AT/Ox | ChiA [AAY89048.1] | 46/62 |
| WP_041541654.1 | 605 | NRPS | Kirromycin [CAN89656.1] | 48/58 |
| WP_012786800.1 | 332 | chlorinating enzyme | / | / |
| WP_012786801.1 | 397 | alpha/beta fold hydrolase | MupV [AAM12938.1] | 30/45 |
| WP_012786802.1 | 221 | Crp/Fnr family transcriptional regulator | LnmO [AAN85528.1] | 46/66 |
| WP_049871560.1 | 2445 | hybrid PKS/NRPS | LnmI [AAN85522.1] | 46/58 |
| WP_012786804.1 | 264 | thioesterase | LnmN [AAN85527.1] | 56/67 |
| WP_012786805.1 | 325 | hypothetical protein | LnmE [AAN85518.1] | 44/63 |
| WP_012786806.1 | 413 | HMG-CoA synthase | MxnE [AGS77285.1] | 53/66 |
| WP_085953918.1 | 235 | enoyl-CoA hydratase | MxnF [AGS77286.1] | 41/52 |
| ORF1 | 4065 | AT-less type I PKS | LnmI [AAN85522.1] | 49/58 |
| WP_083796070.1 | 408 | alpha/beta fold hydrolase | LnmJ [AAN85523.1] | 56/70 |
| WP_012786810.1 | 324 | AT/DC | LnmK [AAN85524.1] | 50/68 |
| WP_012786812.1 | 424 | HMG-CoA synthase | LnmM [AAN85526.1] | 62/74 |
| WP_012786813.1 | 83 | acyl carrier protein | AcpK [CAG23953.1] | 40/66 |
| WP_012786814.1 | 428 | beta-ketoacyl synthase | MxnD [AGS77284.1] | 43/56 |
| WP_012786815.1 | 241 | N-acetylglucosaminyl deacetylase | LnmX [AAN85537.1] | 54/64 |
| WP_041540194.1 | 67 | hypothetical protein | / | / |
| WP_012786816.1 | 267 | class IV aminotransferase | / | / |

[a] ORFs are proteins without protein ID provided by NCBI [b] Number of amino acids

**Table S10.** Predicted functions of ORFs in the NC_019673.1 containing WP_015102005.1 and WP_015102011.1

| gene | aa[a] | putative function | Protein homologue | %identity/ %similarity |
|---|---|---|---|---|
| WP_015102000.1 | 412 | cytochrome P450 | ElaG [AEC04353.1] | 37/59 |
| WP_041313644.1 | 416 | ABC transporter ATP-binding protein | / | / |
| WP_015102002.1 | 267 | ABC transporter permease | / | / |
| WP_015102003.1 | 416 | hypothetical protein | / | / |
| WP_015102004.1 | 406 | alpha/beta hydrolase | / | / |
| WP_015102005.1 | 412 | HMG-CoA synthase | LnmM [AAN85526.1] | 63/77 |
| WP_015102006.1 | 92 | acyl carrier protein | LnmL [AAN85525.1] | 50/66 |
| WP_015102007.1 | 312 | AT/DC | LnmK [AAN85524.1] | 51/63 |
| WP_015102008.1 | 6076 | AT-less type I PKS | LnmJ [AAN85523.1] | 46/57 |
| WP_015102009.1 | 1999 | AT-less type I PKS | LnmI [AAN85522.1] | 49/57 |
| WP_041317703.1 | 297 | hypothetical protein | LnmE [AAN85518.1] | 44/60 |
| WP_015102011.1 | 412 | HMG-CoA synthase | SnaI [CBW45745.1] | 52/66 |
| WP_041317707.1 | 251 | enoyl-CoA hydratase | JamI [AAS98780.1] | 38/58 |
| WP_015102013.1 | 741 | AT/Ox | DifA [CAG23974.1] | 36/53 |
| WP_041313647.1 | 179 | DUF1697 domain-containing protein | / | / |
| WP_015102015.1 | 500 | long-chain fatty acid--CoA ligase | LnmW [AAN85536.1] | 40/56 |
| WP_084672726.1 | 253 | Crp/Fnr family transcriptional regulator | LnmO [AAN85528.1] | 44/64 |
| WP_015102017.1 | 121 | VOC family protein | MxnI [AGS77289.1] | 30/47 |
| WP_015102018.1 | 136 | DUF3224 domain-containing protein | LnmZ’ [AAN85540.1] | 35/56 |

[a] Number of amino acids

**Table S11.** Predicted functions of ORFs in the KY560358.1 containing ASV46894.1

| gene | aa [a] | putative function | Protein homologue | %identity/ %similarity |
|---|---|---|---|---|
| ASV46884.1 | 229 | PPTase | TmlN [CBK62709.1] | 32/49 |
| ASV46885.1 | 317 | hypothetical protein | LnmH [AAN85521.1] | 46/63 |
| ASV46886.1 | 177 | hypothetical protein | / | / |
| ASV46887.1 | 297 | hypothetical protein | LnmE [AAN85518.1] | 39/55 |
| ASV46888.1 | 133 | hypothetical protein | LnmZ' [AAN85540.1] | 42/55 |
| ASV46889.1 | 515 | long-chain fatty acid--CoA ligase | LnmW [AAN85536.1] | 37/53 |
| ASV46890.1 | 142 | ABC transporter component | LnmT [AAN85533.1] | 23/44 |
| ASV46891.1 | 253 | thioesterase | LnmN [AAN85527.1] | 50/63 |
| ASV46892.1 | 419 | beta-ketoacyl synthase | MxnD [AGS77284.1] | 46/59 |
| ASV46893.1 | 87 | hypothetical protein | AcpK [CAG23953.1] | 36/65 |
| ASV46894.1 | 408 | HMG-CoA synthase | LnmM [AAN85526.1] | 61/76 |
| ASV46895.1 | 87 | acyl carrier protein | LnmL [AAN85525.1] | 58/71 |
| ASV46896.1 | 318 | AT/DC | LnmK [AAN85524.1] | 46/60 |
| ASV46897.1 | 1266 | AT-less type I PKS | LnmJ [AAN85523.1] | 50/64 |
| ASV46898.1 | 4602 | AT-less type I PKS | BaeN [CAG23960.2] | 32/48 |
| ASV46899.1 | 1339 | AT-less type I PKS | ChiE [AAY89052.1] | 36/48 |
| ASV46900.1 | 1014 | AT-less type I PKS | kirAIV [CAN89634.1] | 41/53 |

[a] Number of amino acids

**Table S12.** Predicted functions of ORFs in the AYXG01000037.1 containing EWC63732.1

| gene | aa[a] | putative function | Protein homologue | %identity/ %similarity |
|---|---|---|---|---|
| EWC63728.1 | 208 | hypothetical protein UO65_0923 | / | / |
| EWC63729.1 | 2205 | AT-less type I PKS | LnmJ [AAN85523.1] | 65/73 |
| EWC63730.1 | 346 | hypothetical protein UO65_0925 | LnmK [AAN85524.1] | 62/75 |
| EWC63731.1 | 84 | acyl carrier protein | LnmL [AAN85525.1] | 58/72 |
| EWC63732.1 | 409 | HMG-CoA synthase | LnmM [AAN85526.1] | 71/80 |
| EWC63733.1 | 255 | thioesterase | LnmN [AAN85527.1] | 60/70 |
| EWC63734.1 | 375 | glycosyltransferase | / | / |
| EWC63735.1 | 89 | peptidyl carrier protein | LnmP [AAN85529.1] | 53/66 |
| EWC63736.1 | 518 | discrete adenylation domain | LnmQ [AAN85530.1] | 63/69 |
| EWC63737.1 | 373 | glycosyltransferase | SorF [ADN68481.1] | 25/42 |
| EWC63738.1 | 122 | nuclear transport factor 2 | LnmV [AAN85535.1] | 51/69 |
| EWC63739.1 | 543 | hypothetical protein | Nosperin [ADA69237.1] | 35/55 |
| EWC63740.1 | 364 | acyltransferase | / | / |
| EWC63741.1 | 525 | putative ABC transporter ATP-binding protein | LnmR [AAN85531.1] | 66/75 |
| EWC63742.1 | 285 | dipeptide transport system permease protein | LnmS [AAN85532.1] | 69/79 |
| EWC63743.1 | 312 | Peptide ABC transporter, permease protein | LnmT [AAN85533.1] | 62/76 |
| EWC63744.1 | 507 | ABC transporter component | LnmU [AAN85534.1] | 56/70 |
| EWC63745.1 | 487 | drug resistance transporter | LnmV [AAN85535.1] | 55/70 |
| EWC63746.1 | 132 | DUF3224 family protein | LnmZ' [AAN85540.1] | 34/52 |
| EWC63747.1 | 403 | transcriptional regulator, XRE family | / | / |
| EWC63748.1 | 291 | transcriptional regulator, Crp/Fnr family | LnmO [AAN85528.1] | 53/69 |
| EWC63749.1 | 311 | C-5 sterol desaturase | / | / |
| EWC63750.1 | 198 | transcriptional regulator, TetR family | PapR3 [CBW45765.1] | 42/62 |

[a] Number of amino acids

**Table S13.** Predicted functions of ORFs in the JYJF01000008.1 containing KJK59190.1 and KJK59203.1

| gene | aa[a] | putative function | Protein homologue | %identity/ %similarity |
|---|---|---|---|---|
| KJK59185.1 | 75 | hypothetical protein | / | / |
| KJK59186.1 | 126 | glyoxalase | / | / |
| KJK59187.1 | 411 | cytochrome P450 | Dor11 [ACY01396.1] | 35/55 |
| KJK59188.1 | 464 | ferredoxin reductase | / | / |
| KJK59219.1 | 78 | ferredoxin | / | / |
| KJK59189.1 | 144 | hypothetical protein | / | / |
| KJK59220.1 | 251 | thioesterase | LnmN [AAN85527.1] | 56/67 |
| KJK59190.1 | 410 | HMG-CoA synthase | PyxM | 55/69 |
| KJK59191.1 | 260 | enoyl-CoA hydratase | MxnF [AGS77286.1] | 43/53 |
| KJK59192.1 | 750 | AT/Ox | ChiA [AAY89048.1] | 37/51 |
| KJK59221.1 | 503 | multidrug MFS transporter | SnbR [CBW45761.1] | 31/44 |
| KJK59222.1 | 263 | hypothetical protein | LnmE [AAN85518.1] | 44/62 |
| KJK59193.1 | 764 | AT/Ox | MmpC [AAM12912.1] | 51/66 |
| KJK59194.1 | 306 | hypothetical protein | LnmH [AAN85521.1] | 43/59 |
| KJK59195.1 | 375 | hypothetical protein | / | / |
| KJK59196.1 | 119 | glyoxalase | / | / |
| KJK59197.1 | 238 | N-acetylglucosaminyl deacetylase | LnmX [AAN85537.1] | 55/68 |
| KJK59198.1 | 248 | class I SAM-dependent methyltransferase | Dor10 [ACY01395.1] | 31/46 |
| KJK59199.1 | 424 | hydrolase/decarboxylase | LnmD [AAN85517.1] | 46/55 |
| KJK59200.1 | 193 | hypothetical protein | / | / |
| KJK59201.1 | 82 | peptidyl carrier protein | LnmP [AAN85529.1] | 43/59 |
| KJK59202.1 | 7401 | AT-less type I PKS | SorA [ADN68476.1] | 36/48 |
| KJK59203.1 | 411 | HMG-CoA synthase | LnmM [AAN85526.1] | 63/76 |
| KJK59204.1 | 82 | enoyl-CoA hydratase | DipF [AGS06833.1] | 33/64 |
| KJK59224.1 | 64 | enoyl-CoA isomerase | / | / |
| KJK59205.1 | 210 | NACHT domain protein | LnmO [AAN85528.1] | 49/67 |
| KJK59225.1 | 179 | hypothetical protein | / | / |
| KJK59206.1 | 86 | amidase family protein | / | / |
| KJK59207.1 | 305 | aldo/keto reductase | / | / |

[a] Number of amino acids

**Table S14.** Predicted functions of ORFs in the LDST01000044.1 containing KUZ73249.1 and KUZ73260.1

| gene | aa[a] | putative function | Protein homologue | %identity/ %similarity |
|---|---|---|---|---|
| KUZ73239.1 | 199 | lysine transporter LysE | OocC [AFX60325.1] | 32/53 |
| KUZ73302.1 | 371 | efflux RND transporter periplasmic adaptor | / | / |
| KUZ73240.1 | 503 | OprM | / | / |
| KUZ73241.1 | 330 | hypothetical protein | LnmH [AAN85521.1] | 43/60 |
| KUZ73242.1 | 396 | NAD(P)/FAD-dependent oxidoreductase | / | / |
| KUZ73243.1 | 510 | long-chain fatty acid--CoA ligase | LnmW [AAN85536.1] | 34/51 |
| KUZ73244.1 | 130 | DUF3224 domain-containing protein | / | / |
| KUZ73245.1 | 271 | hypothetical protein | LnmE [AAN85518.1] | 37/55 |
| KUZ73246.1 | 157 | DUF1697 domain-containing protein | / | / |
| KUZ73247.1 | 134 | DUF3224 family protein | / | / |
| KUZ73248.1 | 267 | PPTase | SnaN [CBW45738.1] | 39/46 |
| KUZ73249.1 | 410 | HMG-CoA synthase | JamH [AAS98779.1] | 51/66 |
| KUZ73251.1 | 295 | S-malonyltransferase | PsyH [ADA82589.1] | 47/65 |
| KUZ73252.1 | 769 | AT/Ox | DifA [CAG23974.1] | 44/61 |
| KUZ73253.1 | 609 | NRPS | SgvD2 [AGN74876.1] | 46/57 |
| KUZ73254.1 | 313 | chemotaxis protein CheX | / | / |
| KUZ73256.1 | 4169 | AT-less type I PKS | LnmI [AAN85522.1] | 39/49 |
| KUZ73257.1 | 6923 | AT-less type I PKS | LnmJ [AAN85523.1] | 42/54 |
| KUZ73258.1 | 315 | AT/DC | LnmK [AAN85524.1] | 44/58 |
| KUZ73260.1 | 409 | HMG-CoA synthase | LnmM [AAN85526.1] | 58/72 |
| KUZ73261.1 | 92 | acyl carrier protein | NspG [ADA71314.1] | 31/52 |
| KUZ73262.1 | 420 | beta-ketoacyl synthase | CorD [ADI59526.1] | 44/56 |
| KUZ73303.1 | 174 | VOC family protein | / | / |
| KUZ73263.1 | 734 | methylmalonyl-CoA mutase | / | / |

[a] Number of amino acids

**Table S15.** Predicted functions of ORFs in the NZ_JODU01000021.1 containing WP_052841989.1 and WP_052842002.1

| gene | aa[a] | putative function | Protein homologue | %identity/ %similarity |
|---|---|---|---|---|
| WP_052841978.1 | 330 | LysR family transcriptional regulator | SmdB [CCC21116.1] | 29/43 |
| WP_052841979.1 | 397 | glycine C-acetyltransferase | / | / |
| WP_052841980.1 | 342 | L-threonine 3-dehydrogenase | MupE [AAM12917.1] | 28/43 |
| WP_052841981.1 | 191 | GAF domain-containing protein | / | / |
| WP_052841982.1 | 209 | ATP-binding protein | / | / |
| WP_030189437.1 | 144 | roadblock/LC7 domain | / | / |
| WP_052841984.1 | 605 | sensor histidine kinase | / | / |
| WP_052841985.1 | 193 | Crp/Fnr family transcriptional regulator | LnmO [AAN85528.1] | 44/61 |
| WP_052841986.1 | 433 | D-alanine--poly(phosphoribitol) ligase | / | / |
| WP_052841987.1 | 413 | cytochrome P450 | Dor11 [ACY01396.1] | 44/58 |
| WP_052841988.1 | 433 | beta-ketoacyl synthase | CorD [ADI59526.1] | 47/60 |
| WP_052842019.1 | 73 | acyl carrier protein | Fr9M [AIC32699.1] | 36/62 |
| WP_052841989.1 | 411 | HMG-CoA synthase | LnmM [AAN85526.1] | 59/75 |
| WP_052841990.1 | 7020 | AT-less type I PKS | CorL [ADI59534.1] | 40/51 |
| WP_052841991.1 | 542 | discrete adenylation domain | LnmQ [AAN85530.1] | 54/64 |
| WP_052841992.1 | 84 | peptidyl carrier protein | LnmP [AAN85529.1] | 46/58 |
| WP_052841993.1 | 684 | alpha/beta hydrolase | LnmD [AAN85517.1] | 44/56 |
| WP_052841994.1 | 248 | methyltransferase domain | KirM [CAN89643.1] | 45/54 |
| WP_052841995.1 | 238 | N-acetylglucosaminyl deacetylase | LnmX [AAN85537.1] | 54/67 |
| WP_052841996.1 | 133 | DUF3224 domain | / | / |
| WP_052841997.1 | 121 | VOC family protein | / | / |
| WP_052841998.1 | 310 | hypothetical protein | LnmH [AAN85521.1] | 40/56 |
| WP_052842020.1 | 264 | hypothetical protein | LnmE [AAN85518.1] | 48/63 |
| WP_052842021.1 | 489 | MFS transporter | SnbR [CBW45761.1] | 32/46 |
| WP_052841999.1 | 2042 | AT-less type I PKS | LnmI [AAN85522.1] | 49/58 |
| WP_052842000.1 | 738 | AT/ER | MxnA [AGS77281.1] | 36/48 |
| ORF1 | 264 | enoyl-CoA hydratase | MxnF [AGS77286.1] | 42/56 |
| WP_052842002.1 | 410 | HMG-CoA synthase | MxnE [AGS77285.1] | 56/67 |
| WP_052842003.1 | 245 | PPTase | BatI [ADD82950.1] | 32/45 |
| WP_052842004.1 | 252 | thioesterase | LnmN [AAN85527.1] | 54/63 |
| WP_052842005.1 | 145 | hypothetical protein | / | / |

[a] Number of amino acids

**Table S16.** Predicted functions of ORFs in the NZ_LQCG01000010.1 containing WP_059300318.1 and WP_059300329.1

| gene | aa[a] | putative function | Protein homologue | %identity/ %similarity |
|---|---|---|---|---|
| WP_059300313.1 | 488 | long-chain fatty acid--CoA ligase | LnmW [AAN85536.1] | 44/57 |
| WP_079033002.1 | 175 | DUF3224 family protein | LnmZ' [AAN85540.1] | 34/55 |
| WP_059300314.1 | 239 | N-acetylglucosaminyl deacetylase | LnmX [AAN85537.1] | 55/65 |
| WP_059300315.1 | 253 | thioesterase | LnmN [AAN85527.1] | 53/65 |
| WP_059300316.1 | 422 | beta-ketoacyl synthase | MxnD [AGS77284.1] | 44/58 |
| WP_059300317.1 | 80 | acyl carrier protein | ElaE [AEC04351.1] | 36/65 |
| WP_059300318.1 | 412 | HMG-CoA synthase | LnmM [AAN85526.1] | 63/77 |
| WP_059300319.1 | 87 | acyl carrier protein | LnmL [AAN85525.1] | 60/69 |
| WP_059300320.1 | 329 | AT/DC | LnmK [AAN85524.1] | 55/67 |
| WP_059300321.1 | 7157 | AT-less type I PKS | LnmJ [AAN85523.1] | 49/61 |
| WP_059300322.1 | 4242 | NRPS | LnmI [AAN85522.1] | 51/61 |
| WP_059300323.1 | 391 | alpha/beta fold hydrolase | MupV [AAM12938.1] | 31/43 |
| WP_059300324.1 | 311 | chlorinating enzyme | / | / |
| WP_059300325.1 | 599 | NRPS | Fr9DEF [AIC32693.1] | 47/60 |
| WP_059300326.1 | 316 | hypothetical protein | LnmH [AAN85521.1] | 43/59 |
| WP_059300327.1 | 867 | AT/Ox | ChiA [AAY89048.1] | 45/59 |
| WP_059300328.1 | 263 | enoyl-CoA hydratase | LnmF [AAN85519.1] | 51/65 |
| WP_059300329.1 | 410 | HMG-CoA synthase | MxnE [AGS77285.1] | 53/67 |
| WP_059300330.1 | 275 | hypothetical protein | LnmE [AAN85518.1] | 46/63 |
| WP_059300331.1 | 437 | proton exchanger | / | / |
| WP_079033003.1 | 296 | Crp/Fnr family transcriptional regulator | LnmO [AAN85528.1] | 43/62 |
| WP_079033004.1 | 433 | cytochrome P450 | ElaG [AEC04353.1] | 37/56 |
| WP_079033005.1 | 250 | PPTase | BatI [ADD82950.1] | 35/48 |
| WP_059300334.1 | 228 | Crp/Fnr family transcriptional regulator | LnmO [AAN85528.1] | 42/59 |
| WP_063895741.1 | 454 | cytochrome P450 | CorO [ADI59538.1] | 28/47 |

[a] Number of amino acids

**Table S17.** Predicted functions of ORFs in the NZ_FQVU01000002.1 containing WP_073388681.1 and WP_073388691.1

| gene | aa [a] | putative function | Protein homologue | %identity/ %similarity |
|---|---|---|---|---|
| WP_073389586.1 | 248 | N-acetylglucosaminyl deacetylase | LnmX [AAN85537.1] | 50/62 |
| WP_073388648.1 | 541 | MFS transporter | SnbR [CBW45761.1] | 28/45 |
| WP_073388651.1 | 514 | long-chain fatty acid--CoA ligase | LnmW [AAN85536.1] | 32/44 |
| WP_073388654.1 | 223 | hypothetical protein | / | / |
| WP_073388656.1 | 755 | AT/Ox | MmpC [AAM12912.1] | 49/64 |
| WP_073388658.1 | 316 | chlorinating enzyme | / | / |
| WP_073388661.1 | 379 | alpha/beta fold hydrolase | MupV [AAM12938.1] | 25/39 |
| WP_073388672.1 | 2302 | NRPS | LnmI [AAN85522.1] | 45/55 |
| WP_073388676.1 | 251 | thioesterase | LnmN [AAN85527.1] | 52/61 |
| WP_073388681.1 | 416 | HMG-CoA synthase | MxnE [AGS77285.1] | 53/66 |
| WP_084180858.1 | 250 | enoyl-CoA hydratase | CorF [ADI59528.1] | 40/58 |
| WP_073388683.1 | 747 | S-malonyltransferase | BaeE [CAG23952.1] | 34/52 |
| WP_073388684.1 | 2014 | AT-less type I PKS | LnmI [AAN85522.1] | 48/57 |
| ORF1 | 7281 | AT-less type I PKS | MisF [AKQ22696.1] | 31/45 |
| WP_084180926.1 | 353 | alpha/beta hydrolase | LnmJ [AAN85523.1] | 50/64 |
| WP_084180927.1 | 287 | AT/DC | LnmK [AAN85524.1] | 45/60 |
| WP_073388688.1 | 80 | acyl carrier protein | LnmL [AAN85525.1] | 59//72 |
| WP_073388691.1 | 418 | HMG-CoA synthase | LnmM [AAN85526.1] | 66/79 |
| WP_073388694.1 | 87 | acyl carrier protein | PedN [AAW33973.1] | 45/65 |
| WP_073388697.1 | 420 | beta-ketoacyl synthase | MxnD [AGS77284.1] | 45/58 |
| WP_084180859.1 | 470 | propionyl-CoA carboxylase | SnbS [CBW45762.1] | 41/56 |
| WP_073388703.1 | 369 | flavin-dependent monooxygenase | OocK [AFX60333.1] | 24/38 |
| WP_084180860.1 | 400 | cytochrome P450 | Dor11 [ACY01396.1] | 38/55 |
| WP_073388706.1 | 307 | hypothetical protein | LnmH [AAN85521.1] | 45/63 |
| WP_084180861.1 | 428 | hypothetical protein | / | / |
| WP_073388712.1 | 175 | DUF1697 domain-containing protein | / | / |

[a] Number of amino acids

**Table S18.** Predicted functions of ORFs in the NZ_MKQR01000006.1 containing WP_075973383.1 and WP_075973463.1

| gene | aa [a] | putative function | Protein homologue | %identity/ %similarity |
|---|---|---|---|---|
| WP_075973378.1 | 137 | DUF3224 domain-containing protein | LnmZ' [AAN85540.1] | 39/59 |
| WP_075973379.1 | 264 | Crp/Fnr family transcriptional regulator | LnmO [AAN85528.1] | 42/58 |
| WP_075973380.1 | 505 | long-chain fatty acid--CoA ligase | LnmW [AAN85536.1] | 41/55 |
| WP_075973381.1 | 180 | DUF1697 domain-containing protein | leinamycin [AAN85541.1] | 45/61 |
| WP_075973382.1 | 671 | AT/Ox | DifA [CAG23974.1] | 36/52 |
| WP_075973462.1 | 245 | enoyl-CoA hydratase | MxnF [AGS77286.1] | 44/55 |
| WP_075973383.1 | 412 | HMG-CoA synthase | CorE [ADI59527.1] | 53/65 |
| WP_075973384.1 | 293 | hypothetical protein | LnmE [AAN85518.1] | 41/58 |
| WP_084793662.1 | 1875 | AT-less type I PKS | LnmI [AAN85522.1] | 48/75 |
| WP_084793673.1 | 6566 | AT-less type I PKS | CorL [ADI59534.1] | 38/49 |
| WP_075973387.1 | 311 | AT/DC | LnmK [AAN85524.1] | 51/63 |
| WP_075973388.1 | 84 | acyl carrier protein | LnmL [AAN85525.1] | 52/69 |
| WP_075973463.1 | 410 | HMG-CoA synthase | LnmM [AAN85526.1] | 61/73 |
| WP_075973389.1 | 382 | alpha/beta hydrolase | MupV [AAM12938.1] | 25/37 |
| WP_075973390.1 | 416 | hypothetical protein | / | / |

[a] Number of amino acids

**Table S19.** Predicted functions of ORFs in the NZ_KI912588.1 containing WP_080644569.1 and WP_018792600.1

| gene[a] | aa[b] | putative function | Protein homologue | %identity/ %similarity |
|---|---|---|---|---|
| WP_018792612.1 | 354 | cysteine synthase | Kirromycin [CAN89659.1] | 42/56 |
| WP_018792611.1 | 485 | argininosuccinate lyase | Kirromycin [CAN89657.1] | 37/49 |
| WP_018792610.1 | 407 | ATP-grasp domain | Kirromycin [CAN89659.1] | 42/53 |
| WP_018792609.1 | 499 | long-chain fatty acid--CoA ligase | LnmW [AAN85536.1] | 41/54 |
| WP_018792608.1 | 297 | hypothetical protein | LnmE [AAN85518.1] | 44/59 |
| WP_018792607.1 | 547 | ABC transporter ATP-binding protein | LnmR [AAN85531.1] | 57/69 |
| WP_018792606.1 | 263 | ABC transporter permease | LnmS [AAN85532.1] | 62/72 |
| WP_026269204.1 | 318 | ABC transporter permease | LnmT [AAN85533.1] | 55/72 |
| WP_018792604.1 | 511 | ABC transporter substrate-binding protein | LnmU [AAN85534.1] | 58/74 |
| WP_018792603.1 | 270 | thioesterase | LnmN [AAN85527.1] | 56/69 |
| WP_018792602.1 | 433 | beta-ketoacyl synthase | MxnD [AGS77284.1] | 48/62 |
| WP_018792601.1 | 86 | acyl carrier protein | SnaG [CBW45747.1] | 46/63 |
| WP_018792600.1 | 407 | HMG-CoA synthase | LnmM [AAN85526.1] | 64/76 |
| WP_018792599.1 | 90 | acyl carrier protein | LnmL [AAN85525.1] | 57/66 |
| WP_018792598.1 | 320 | AT/DC | LnmK [AAN85524.1] | 56/67 |
| ORF1 | 7105 | AT-less type I PKS (DH-Acp-KR-KS-DH-ECH-Acp-KS-DH-KR-MT-Acp-KS-DH-KR-Acp-KS-DH-Acp-Acp) | CorL [ADI59534.1] | 40/52 |
| WP_018792597.1 | 381 | alpha/beta fold hydrolase | MupV [AAM12938.1] | 29/44 |
| WP_018792596.1 | 317 | chlorinating enzyme | / | / |
| WP_028674776.1 | 81 | acyl carrier protein | BaeJ [CAG23957.2] | 47/56 |
| WP_018792594.1 | 1086 | NRPS (A-PCP-TE) | CalI [BAP05597.1] | 37/52 |
| WP_018792593.1 | 311 | kinase | Kirromycin [CAN89658.1] | 52/63 |
| WP_018792592.1 | 76 | hypothetical protein | / | / |
| WP_018792591.1 | 336 | TauD/TfdA family dioxygenase | Leinamycin [AAN85495.1] | 35/50 |
| WP_018792590.1 | 4311 | hybrid NRPS/PKS | LnmI [AAN85522.1] | 49/59 |
| WP_018792589.1 | 783 | AT/Ox | DifA [CAG23974.1] | 46/66 |
| WP_018792588.1 | 262 | enoyl-CoA hydratase | JamI [AAS98780.1] | 37/59 |
| WP_080644569.1 | 381 | HMG-CoA synthase | SnaI [CBW45745.1] | 55/70 |
| WP_018792586.1 | 287 | hypothetical protein | LnmE [AAN85518.1] | 37/52 |
| WP_018792585.1 | 291 | PPTase | BatI [ADD82950.1] | 37/52 |
| WP_018792584.1 | 133 | DUF3224 domain | LnmZ' [AAN85540.1]' | 39/56 |
| WP_032717735.1 | 115 | VOC family protein | / | / |
| WP_018792582.1 | 405 | hypothetical protein | / | / |
| WP_018792581.1 | 308 | hypothetical protein | LnmH [AAN85521.1] | 49/58 |

[a] ORFs are proteins without protein ID provided by NCBI [b] Number of amino acids

**Table S20.** Predicted functions of ORFs in the NZ_FMZZ01000014.1 containing WP_091455311.1

| gene | aa [a] | putative function | Protein homologue | %identity/ %similarity |
|---|---|---|---|---|
| WP_091455300.1 | 171 | DNA-binding protein | / | / |
| WP_091455302.1 | 368 | hypothetical protein | / | / |
| WP_091455304.1 | 4197 | hybrid PKS/NRPS | LnmI [AAN85522.1] | 54/62 |
| WP_091455305.1 | 6588 | AT-less type I PKS | LnmJ [AAN85523.1] | 55/65 |
| WP_091455307.1 | 319 | AT/DC | LnmK [AAN85524.1] | 60/72 |
| WP_091455309.1 | 86 | acyl carrier protein | LnmL [AAN85525.1] | 62/76 |
| WP_091455311.1 | 409 | HMG-CoA synthase | LnmM [AAN85526.1] | 69/79 |
| WP_091455426.1 | 246 | thioesterase | LnmN [AAN85527.1] | 58/70 |
| WP_091455313.1 | 1026 | PKD domain-containing protein | / | / |
| WP_091455314.1 | 373 | DUF1205 domain-containing protein | / | / |

[a] Number of amino acids

**Table S21.** Predicted functions of ORFs in the NZ_CVPB01000001.1 containing WP_093802472.1 and WP_093802479.1

| gene | aa [a] | putative function | Protein homologue | %identity/ %similarity |
|---|---|---|---|---|
| WP_093802448.1 | 531 | ABC transporter ATP-binding protein | LnmR [AAN85531.1] | 56/69 |
| WP_093802449.1 | 194 | cytochrome P450 | ElaG [AEC04353.1] | 37/55 |
| WP_093802450.1 | 233 | cytochrome P450 | ElaG [AEC04353.1] | 45/61 |
| WP_093802451.1 | 392 | class I SAM-dependent methyltransferase | OzmL [ABS90473.1] | 38/52 |
| WP_093802452.1 | 289 | condensation domain | kirHI [CAN89638.1] | 35/48 |
| WP_093802453.1 | 272 | hypothetical protein | LnmE [AAN85518.1] | 33/49 |
| WP_093802454.1 | 791 | AT/Ox | MmpC [AAM12912.1] | 49/65 |
| WP_093802456.1 | 4275 | NRPS | LnmI [AAN85522.1] | 51/59 |
| WP_093802458.1 | 316 | dioxygenase | AlbVIII [CAE52335.1] | 31/49 |
| WP_093803771.1 | 305 | kinase | kirromycin [CAN89658.1] | 51/62 |
| WP_093802459.1 | 1061 | NRPS | CalI [BAP05597.1] | 34/48 |
| WP_093802460.1 | 400 | ATP-grasp domain | kirromycin [CAN89659.1] | 42/53 |
| WP_093802461.1 | 486 | argininosuccinate lyase | kirromycin [CAN89657.1] | 39/52 |
| WP_093802463.1 | 355 | cysteine synthase | kirromycin [CAN89659.1] | 46/57 |
| WP_093802464.1 | 165 | acyl carrier protein | kirromycin [CAN89663.1] | 39/51 |
| WP_093802465.1 | 317 | chlorinating enzyme | / | / |
| WP_093802466.1 | 285 | alpha/beta fold hydrolase | AlbXI [CAE52328.1] | 26/39 |
| WP_093802467.1 | 1060 | AT-less type I PKS | LnmJ [AAN85523.1] | 44/54 |
| WP_093802468.1 | 543 | AT-less type I PKS | LnmJ [AAN85523.1] | 71/80 |
| ORF1 | 5854 | AT-less type I PKS | LnmJ [AAN85523.1] | 49/59 |
| WP_107081135.1 | 282 | alpha/beta fold hydrolase | LnmJ [AAN85523.1] | 61/73 |
| WP_093802470.1 | 299 | AT/DC | LnmK [AAN85524.1] | 54/65 |
| WP_093802471.1 | 86 | acyl carrier protein | LnmL [AAN85525.1] | 59/71 |
| WP_093802472.1 | 406 | HMG-CoA synthase | LnmM [AAN85526.1] | 66/76 |
| WP_093803772.1 | 79 | PPTase | PedN [AAW33973.1] | 36/64 |
| WP_093802473.1 | 419 | beta-ketoacyl synthase | MxnD [AGS77284.1] | 46/59 |
| WP_093802474.1 | 266 | thioesterase | LnmN [AAN85527.1] | 51/63 |
| WP_093802476.1 | 212 | YbhB/YbcL family Raf kinase inhibitor | / | / |
| WP_093802477.1 | 127 | DUF3224 domain-containing protein | LnmZ' [AAN85540.1] | 34/50 |
| WP_093803774.1 | 220 | PPTase | LtmL [ACY01405.1] | 48/60 |
| WP_093802478.1 | 261 | enoyl-CoA hydratase | LnmF [AAN85519.1] | 52/66 |
| WP_093802479.1 | 410 | HMG-CoA synthase | SnaI [CBW45745.1] | 55/67 |
| WP_093802480.1 | 442 | cation/H(+) antiporter | / | / |
| WP_093803775.1 | 238 | Crp/Fnr family transcriptional regulator | LnmO [AAN85528.1] | 43/60 |
| WP_093802482.1 | 134 | DUF3224 domain-containing protein | LnmZ' [AAN85540.1] | 42/58 |
| WP_093802483.1 | 314 | hypothetical protein | LnmH [AAN85521.1] | 43/57 |
| WP_093802484.1 | 522 | long-chain fatty acid--CoA ligase | LnmW [AAN85536.1] | 40/55 |
| WP_093802485.1 | 236 | N-acetylglucosaminyl deacetylase | LnmX [AAN85537.1] | 52/65 |
| WP_093802486.1 | 273 | hypothetical protein | LnmE [AAN85518.1] | 45/63 |
| WP_093802487.1 | 93 | acyl carrier protein | kirB [CAN89639.1] | 41/56 |

| | | | | |
|---|---|---|---|---|
| WP_093802488.1 | 276 | hypothetical protein | kirHI [CAN89638.1] | 31/46 |
| WP_093802489.1 | 500 | long-chain fatty acid--CoA ligase | LnmW [AAN85536.1] | 43/57 |
| WP_093802490.1 | 451 | serine hydroxymethyltransferase | / | / |
| WP_093802491.1 | 762 | NRPS | JamO [AAS98786.1] | 34/50 |
| WP_093802492.1 | 137 | DUF427 domain-containing protein | / | / |
| WP_093802493.1 | 364 | hypothetical protein | / | / |

[a] Number of amino acids

**Table S22.** Predicted functions of ORFs in the NZ_OBMJ01000001.1 containing WP_097240137.1 and WP_097237235.1

| gene | aa [a] | putative function | Protein homologue | %identity/ %similarity |
|---|---|---|---|---|
| WP_097237206.1 | 130 | RidA family protein | / | / |
| WP_097237207.1 | 399 | class I SAM-dependent methyltransferase | / | / |
| WP_097237209.1 | 119 | nuclear transport factor 2 | LnmV [AAN85535.1] | 42/63 |
| WP_097237210.1 | 406 | DUF1205 domain-containing protein | / | / |
| WP_097237211.1 | 283 | hypothetical protein | LnmE [AAN85518.1] | 45/64 |
| WP_097240137.1 | 413 | HMG-CoA synthase | SnaI [CBW45745.1] | 54/67 |
| WP_097237212.1 | 270 | enoyl-CoA hydratase | LnmF [AAN85519.1] | 49/66 |
| WP_097237214.1 | 417 | cytochrome P450 | ElaG [AEC04353.1] | 36/53 |
| WP_097237215.1 | 127 | DUF3224 domain-containing protein | LnmZ' [AAN85540.1] | 38/57 |
| WP_097237216.1 | 249 | macrocin O-methyltransferase | / | / |
| WP_097237217.1 | 289 | dTDP-4-dehydrorhamnose reductase | / | / |
| WP_097237218.1 | 287 | hypothetical protein | LnmH [AAN85521.1] | 43/63 |
| WP_097237219.1 | 262 | hypothetical protein | LnmE [AAN85518.1] | 32/49 |
| WP_097237220.1 | 760 | AT/Ox | MmpC [AAM12912.1] | 52/68 |
| WP_097237221.1 | 4209 | hybrid PKS/NRPS | LnmI [AAN85522.1] | 50/60 |
| WP_097237222.1 | 316 | regulatory protein | leinamycin [AAN85494.1] | 41/57 |
| WP_097240138.1 | 309 | kinase | kirromycin [CAN89658.1] | 53/63 |
| WP_097240139.1 | 1054 | NRPS | CalI [BAP05597.1] | 34/48 |
| WP_097237223.1 | 403 | ATP-grasp domain-containing protein | kirromycin [CAN89659.1] | 44/54 |
| WP_097237224.1 | 483 | argininosuccinate lyase | kirromycin [CAN89657.1] | 37/52 |
| WP_097237225.1 | 348 | pyridoxal-phosphate dependent enzyme | kirromycin [CAN89659.1] | 46/58 |
| WP_097237226.1 | 84 | acyl carrier protein | kirromycin [CAN89663.1] | 42/53 |
| WP_097237228.1 | 318 | chlorinating enzyme | / | / |
| WP_097237230.1 | 281 | alpha/beta fold hydrolase | MupV [AAM12938.1] | 35/50 |
| WP_097237232.1 | 6910 | AT-less type I PKS | LnmJ [AAN85523.1] | 49/60 |
| WP_097237234.1 | 86 | acyl carrier protein | LnmL [AAN85525.1] | 71/80 |
| WP_097237235.1 | 410 | HMG-CoA synthase | LnmM [AAN85526.1] | 65/76 |
| WP_097240140.1 | 80 | acyl carrier protein | PedN [AAW33973.1] | 36/66 |
| WP_097237236.1 | 421 | beta-ketoacyl synthase | CorD [ADI59526.1] | 48/61 |
| WP_097237237.1 | 264 | thioesterase | LnmN [AAN85527.1] | 52/63 |
| WP_097237239.1 | 324 | LuxR family regulator | kirromycin [CAN89667.1] | 30/44 |
| WP_097237240.1 | 100 | hypothetical protein | / | / |

[a] Number of amino acids

**Table S23.** Predicted functions of ORFs in the NZ_OBDY01000002.1 containing WP_097318953.1 and WP_097318960.1

| gene | aa [a] | putative function | Protein homologue | %identity/ %similarity |
|---|---|---|---|---|
| WP_097318941.1 | 444 | cation/H (+) antiporter | / | / |
| WP_097318942.1 | 556 | MFS transporter | leinamycin [AAN85487.1] | 30/46 |
| WP_097319226.1 | 397 | cytochrome P450 | Dor11 [ACY01396.1] | 44/59 |
| WP_097318943.1 | 303 | hypothetical protein | LnmH [AAN85521.1] | 45/61 |
| WP_097318944.1 | 185 | DUF3224 domain-containing protein | LnmZ' [AAN85540.1] | 30/52 |
| WP_097318945.1 | 766 | AT/Ox | MmpC [AAM12912.1] | 51/66 |
| WP_097318946.1 | 601 | NRPS | fr9DEF [AIC32693.1] | 47/61 |
| WP_097318947.1 | 313 | chlorinating enzyme | / | / |
| WP_097318948.1 | 397 | alpha/beta hydrolase | MupV [AAM12938.1] | 30/45 |
| WP_097318949.1 | 208 | Crp/Fnr family transcriptional regulator | LnmO [AAN85528.1] | 49/69 |
| WP_097318950.1 | 2200 | hybrid PKS/NRPS | LnmI [AAN85522.1] | 49/69 |
| WP_097318951.1 | 275 | thioesterase | LnmN [AAN85527.1] | 56/66 |
| WP_097318952.1 | 300 | hypothetical protein | LnmE [AAN85518.1] | 48/62 |
| WP_097318953.1 | 407 | HMG-CoA synthase | MxnE [AGS77285.1] | 56/69 |
| WP_097318954.1 | 248 | enoyl-CoA hydratase | CylG [ARU81121.1] | 36/58 |
| WP_097318955.1 | 690 | S-malonyltransferase | ElaB [AEC04348.1] | 38/51 |
| WP_097318956.1 | 1906 | AT-less type I PKS | LnmI [AAN85522.1] | 53/62 |
| WP_097318957.1 | 7153 | AT-less type I PKS | LnmJ [AAN85523.1] | 47/57 |
| WP_097318958.1 | 318 | AT/DC | LnmK [AAN85524.1] | 50/63 |
| WP_097318959.1 | 81 | acyl carrier protein | LnmL [AAN85525.1] | 57/75 |
| WP_097318960.1 | 417 | HMG-CoA synthase | LnmM [AAN85526.1] | 62/76 |
| WP_097318961.1 | 79 | acyl carrier protein | PedN [AAW33973.1] | 41/67 |
| WP_097318962.1 | 409 | beta-ketoacyl synthase | MxnD [AGS77284.1] | 48/58 |
| WP_097319227.1 | 238 | N-acetylglucosaminyl deacetylase | LnmX [AAN85537.1] | 56/64 |
| WP_097318963.1 | 191 | hypothetical protein | / | / |
| WP_097318964.1 | 504 | long-chain fatty acid--CoA ligase | LnmW [AAN85536.1] | 34/45 |
| WP_097318965.1 | 132 | DUF3224 domain-containing protein | LnmZ' [AAN85540.1] | 31/53 |
| WP_097318966.1 | 356 | hypothetical protein | / | / |
| WP_097318967.1 | 177 | DUF1697 domain-containing protein | / | / |
| WP_097318968.1 | 256 | thioesterase | CylH [ARU81122.1] | 42/59 |
| WP_097318969.1 | 1239 | AT-less type I PKS | BaeJ [CAG23957.2] | 37/54 |
| WP_097318970.1 | 442 | hypothetical protein | OzmL [ABS90473.1] | 33/46 |
| WP_097319228.1 | 420 | salicylate synthase | / | / |

[a] Number of amino acids

**Table S24.** Predicted functions of ORFs in the NZ_NNBJ01000007.1 containing WP_100575958.1 and WP_100575972.1

| gene | aa [a] | putative function | Protein homologue | %identity/ %similarity |
|---|---|---|---|---|
| WP_100575950.1 | 889 | GIY-YIG nuclease | / | / |
| WP_100575951.1 | 68 | hypothetical protein | / | / |
| WP_100575952.1 | 61 | hypothetical protein | / | / |
| WP_100575953.1 | 367 | winged helix DNA-binding domain | / | / |
| WP_100575954.1 | 145 | globin | / | / |
| WP_100575955.1 | 2338 | hybrid PKS/NRPS | LnmI [AAN85522.1] | 46/57 |
| WP_100575956.1 | 252 | thioesterase | LnmN [AAN85527.1] | 53/62 |
| WP_100575957.1 | 290 | PPTase | TmlN [CBK62709.1] | 30/46 |
| WP_100575958.1 | 410 | HMG-CoA synthase | MxnE [AGS77285.1] | 52/66 |
| WP_100575959.1 | 257 | enoyl-CoA hydratase | LnmF [AAN85519.1] | 50/62 |
| WP_100575960.1 | 743 | AT/ER | MxnA [AGS77281.1] | 41/53 |
| WP_100575961.1 | 2063 | AT-less type I PKS | LnmI [AAN85522.1] | 51/60 |
| WP_100576080.1 | 486 | DHA2 family efflux MFS transporter | LnmY [AAN85538.1] | 32/49 |
| WP_100575962.1 | 299 | hypothetical protein | LnmE [AAN85518.1] | 43/59 |
| WP_100575963.1 | 756 | AT/Ox | MmpC [AAM12912.1] | 49/67 |
| WP_100575964.1 | 310 | hypothetical protein | LnmH [AAN85521.1] | 42/58 |
| WP_100575965.1 | 121 | VOC family protein | / | / |
| WP_100575966.1 | 135 | hypothetical protein | LnmZ' [AAN85540.1] | 35/49 |
| WP_100575967.1 | 238 | N-acetylglucosaminyl deacetylase | LnmX [AAN85537.1] | 52/65 |
| WP_100575968.1 | 248 | methyltransferase | KirM [CAN89643.1] | 41/53 |
| WP_100575969.1 | 664 | alpha/beta hydrolase | LnmD [AAN85517.1] | 36/47 |
| WP_100576081.1 | 74 | peptidyl carrier protein | LnmP [AAN85529.1] | 41/60 |
| WP_100575970.1 | 528 | discrete adenylation domain | LnmQ [AAN85530.1] | 55/66 |
| WP_100575971.1 | 7016 | AT-less type I PKS | SorA [ADN68476.1] | 36/49 |
| WP_100575972.1 | 418 | HMG-CoA synthase | LnmM [AAN85526.1] | 62/78 |
| WP_100576082.1 | 73 | acyl carrier protein | OocG [AFX60329.1] | 42/61 |
| WP_100576083.1 | 427 | beta-ketoacyl synthase | CorD [ADI59526.1] | 49/62 |
| WP_100575973.1 | 412 | cytochrome P450 | Dor11 [ACY01396.1] | 45/60 |
| WP_100575974.1 | 438 | aminotransferase | / | / |
| WP_100575975.1 | 229 | Crp/Fnr family transcriptional regulator | LnmO [AAN85528.1] | 42/60 |
| WP_100576084.1 | 569 | S-methyl-5-thioribose-1-phosphate isomerase | / | / |

[a] Number of amino acids

**Table S25.** Predicted functions of ORFs in the NZ_NNBO01000005.1 containing WP_100586607.1

| gene | aa [a] | putative function | Protein homologue | %identity/ %similarity |
|---|---|---|---|---|
| WP_100586595.1 | 543 | ABC transporter ATP-binding protein | LnmR [AAN85531.1] | 65/73 |
| WP_100586732.1 | 517 | discrete adenylation domain | LnmQ [AAN85530.1] | 63/72 |
| WP_100586596.1 | 91 | peptidyl carrier protein | LnmP [AAN85529.1] | 55/70 |
| WP_100586597.1 | 228 | Crp/Fnr family transcriptional regulator | LnmO [AAN85528.1] | 60/73 |
| WP_100586598.1 | 76 | ferredoxin | LnmB [AAN85515.1] | 71/80 |
| WP_100586599.1 | 399 | cytochrome P450 | LnmA [AAN85514.1] | 82/90 |
| WP_100586600.1 | 449 | alpha/beta hydrolase | LnmD [AAN85517.1] | 53/63 |
| WP_100586733.1 | 261 | enoyl-CoA hydratase | LnmF [AAN85519.1] | 63/70 |
| WP_100586601.1 | 757 | AT/Ox | DifA [CAG23974.1] | 49/67 |
| WP_100586602.1 | 290 | hypothetical protein | LnmH [AAN85521.1] | 73/83 |
| WP_100586603.1 | 4552 | hybrid PKS/NRPS | LnmI [AAN85522.1] | 62/70 |
| WP_100586604.1 | 7573 | AT-less type I PKS | LnmJ [AAN85523.1] | 61/69 |
| WP_100586605.1 | 328 | AT/DC | LnmK [AAN85524.1] | 67/77 |
| WP_100586606.1 | 87 | acyl carrier protein | LnmL [AAN85525.1] | 66/78 |
| WP_100586607.1 | 418 | HMG-CoA synthase | LnmM [AAN85526.1] | 77/85 |
| WP_100586734.1 | 251 | thioesterase | LnmN [AAN85527.1] | 62/71 |
| WP_100586608.1 | 266 | PPTase | snaN [CBW45738.1] | 52/61 |
| WP_100586609.1 | 136 | DUF3224 domain-containing protein | LnmZ' [AAN85540.1] | 82/85 |
| WP_100586735.1 | 396 | cytochrome P450 | LnmZ [AAN85539.1] | 76/86 |
| WP_100586610.1 | 130 | DUF3224 domain-containing protein | / | / |
| WP_100586611.1 | 475 | MFS transporter | LnmY [AAN85538.1] | 77/87 |
| WP_100586612.1 | 242 | N-acetylglucosaminyl deacetylase | LnmX [AAN85537.1] | 79/88 |
| WP_100586613.1 | 512 | long-chain fatty acid--CoA ligase | LnmW [AAN85536.1] | 74/80 |
| WP_100586614.1 | 120 | nuclear transport factor 2 family protein | LnmV [AAN85535.1] | 83/93 |
| WP_100586615.1 | 367 | alcohol dehydrogenase | enacyloxin [ABI91459.1] | 28/41 |
| WP_100586616.1 | 305 | fructokinase | / | / |
| WP_100586617.1 | 479 | hypothetical protein | / | / |
| WP_100586618.1 | 174 | intradiol ring-cleavage dioxygenase | / | / |

[a] Number of amino acids

**Table S26.** Predicted functions of ORFs in the NZ_PNRA02000003.1 containing WP_102642253.1 and WP_102637318.1

| gene | aa [a] | putative function | Protein homologue | %identity/ %similarity |
|---|---|---|---|---|
| WP_102641881.1 | 264 | thioesterase | LnmN [AAN85527.1] | 53/62 |
| WP_109288566.1 | 408 | beta-ketoacyl synthase | CorD [ADI59526.1] | 46/55 |
| WP_102642254.1 | 97 | acyl carrier protein | DipF [AGS06833.1] | 32/56 |
| WP_102642253.1 | 416 | HMG-CoA synthase | TaF [ABF92623.1] | 55/69 |
| WP_109288567.1 | 7532 | AT-less type I PKS | LnmJ [AAN85523.1] | 44/53 |
| WP_109288568.1 | 514 | discrete adenylation domain | LnmQ [AAN85530.1] | 53/62 |
| WP_107503768.1 | 114 | peptidyl carrier protein | LnmP [AAN85529.1] | 44/65 |
| WP_102642172.1 | 423 | alpha/beta hydrolase | LnmD [AAN85517.1] | 37/47 |
| WP_102642173.1 | 305 | methyltransferase | KirM [CAN89643.1] | 42/56 |
| WP_102642174.1 | 236 | N-acetylglucosaminyl deacetylase | LnmX [AAN85537.1] | 49/61 |
| WP_102642175.1 | 120 | VOC family protein | CalO [BAP05583.1] | 26/39 |
| WP_102641762.1 | 302 | hypothetical protein | LnmH [AAN85521.1] | 36/55 |
| WP_102641763.1 | 839 | AT/Ox | MmpC [AAM12912.1] | 46/60 |
| WP_102641764.1 | 240 | hypothetical protein | LnmE [AAN85518.1] | 38/54 |
| WP_102641765.1 | 521 | DHA2 family efflux MFS transporter permease | LnmY [AAN85538.1] | 32/49 |
| WP_109288569.1 | 2231 | AT-less type I PKS | ElaP [AEC04362.1] | 30/45 |
| WP_102637319.1 | 284 | acyltransferase | LnmG [AAN85520.1] | 42/57 |
| WP_102637328.1 | 255 | enoyl-CoA hydratase | MxnF [AGS77286.1] | 41/54 |
| WP_102637318.1 | 409 | HMG-CoA synthase | JamH [AAS98779.1] | 45/64 |
| WP_102637317.1 | 256 | PPTase | TmlN [CBK62709.1] | 31/44 |
| WP_102637316.1 | 264 | hypothetical protein | / | / |
| WP_102637315.1 | 236 | Crp/Fnr family transcriptional regulator | LnmO [AAN85528.1] | 34/56 |
| WP_102637314.1 | 164 | stress response protein | / | / |
| WP_102637313.1 | 551 | multicopper oxidase family protein | / | / |
| WP_102637312.1 | 339 | LacI family transcriptional regulator | / | / |

[a] Number of amino acids

**Table S27.** Predicted functions of ORFs in the NZ_QOIL01000017.1 containing WP_114031836.1 and WP_114031842.1

| gene [a] | aa [b] | putative function | Protein homologue | %identity/ %similarity |
|---|---|---|---|---|
| WP_114031831.1 | 477 | long-chain fatty acid--CoA ligase | LnmW [AAN85536.1] | 34/48 |
| WP_114031832.1 | 179 | DUF1697 domain-containing protein | / | / |
| WP_114031833.1 | 367 | flavin-dependent monooxygenase | OocK [AFX60333.1] | 32/47 |
| WP_114031834.1 | 147 | DUF3224 family protein | LnmZ' [AAN85540.1] | 30/47 |
| WP_114031835.1 | 79 | acyl carrier protein | PedN [AAW33973.1] | 44/70 |
| WP_114031836.1 | 419 | HMG-CoA synthase | LnmM [AAN85526.1] | 64/76 |
| WP_114031837.1 | 78 | acyl carrier protein | TaE [ABF88942.1] | 55/67 |
| WP_114031838.1 | 319 | AT/DC | LnmK [AAN85524.1] | 56/70 |
| WP_114031885.1 | 232 | alpha/beta fold hydrolase | LnmJ [AAN85523.1] | 64/75 |
| ORF1 | 8079 | AT-less type I PKS | CalG [BAP05595.1] | 33/47 |
| ORF2 | 4563 | AT-less type I PKS | LnmI [AAN85522.1] | 48/57 |
| WP_114031839.1 | 261 | alpha/beta fold hydrolase | LnmN [AAN85527.1] | 53/63 |
| WP_114031887.1 | 329 | SMP-30/gluconolactonase/LRE | / | / |
| WP_114031840.1 | 785 | AT/Ox | MmpC [AAM12912.1] | 50/65 |
| WP_114031841.1 | 241 | enoyl-CoA hydratase | LnmF [AAN85519.1] | 59/70 |
| WP_114031842.1 | 407 | HMG-CoA synthase | MxnE [AGS77285.1] | 54/68 |
| WP_114031843.1 | 300 | hypothetical protein | LnmE [AAN85518.1] | 44/63 |
| WP_114031844.1 | 480 | cation:proton antiporter | / | / |
| WP_114031845.1 | 1221 | choice-of-anchor D domain-containing protein | / | / |

[a] ORFs are proteins without protein ID provided by NCBI [b] Number of amino acids

**Table S28.** Predicted functions of ORFs in the CP002162.1 containing ADL47416.1 and ADL47427.1

| gene | aa[a] | putative function | protein homologue | %identity/ %similarity |
|---|---|---|---|---|
| ADL47410.1 | 412 | cytochrome P450 | BaeS [CAG23962.1] | 38/58 |
| ADL47411.1 | 328 | daunorubicin resistance ABC transporter ATPase subunit | Rhizopodin [CCA89322.1] | 31/49 |
| ADL47412.1 | 266 | hypothetical protein | / | / |
| ADL47413.1 | 418 | hypothetical protein | / | / |
| ADL47414.1 | 32 | hypothetical protein | / | / |
| ADL47415.1 | 397 | alpha/beta hydrolase fold | MupV [AAM12938.1] | 31/47 |
| ADL47416.1 | 411 | HMG-CoA synthase | Myxovirescins [ABF92623.1] | 62/75 |
| ADL47417.1 | 82 | acyl carrier protein | LnmL [AAN85525.1] | 59/71 |
| ADL47418.1 | 311 | AT/DC | LnmK [AAN85524.1] | 55/67 |
| ADL47419.1 | 6727 | AT-less type I PKS | CorL [ADI59534.1] | 41/51 |
| ADL47420.1 | 2005 | AT-less type I PKS | ChiD [AAY89051.1] | 41/52 |
| ADL47421.1 | 292 | hypothetical protein | LnmE [AAN85518.1] | 43/60 |
| ADL47422.1 | 252 | enoyl-CoA hydratase/isomerase | JamI [AAS98780.1] | 40/58 |
| ADL47423.1 | 735 | AT/Ox | ChiA [AAY89048.1] | 37/50 |
| ADL47424.1 | 253 | cyclic nucleotide-binding | LnmO [AAN85528.1] | 46/59 |
| ADL47425.1 | 426 | sodium/hydrogen exchanger | PsyD [ADA82585.1] | 36/54 |
| ADL47426.1 | 179 | protein of unknown function DUF1697 | Leinamycin [AAN85541.1] | 32/47 |
| ADL47427.1 | 413 | HMG-CoA synthase | CorE [ADI59527.1] | 55/68 |
| ADL47428.1 | 494 | long-chain fatty acid--CoA ligase | LnmW [AAN85536.1] | 42/57 |

[a] Number of amino acids

**Table S29.** Predicted functions of ORFs in the NZ_AWXG01000021.1 containing WP_029317813.1

| gene | aa[a] | putative function | Protein homologue | %identity/ %similarity |
|---|---|---|---|---|
| WP_029317807.1 | 205 | TetR family transcriptional regulator | SpbR [CBW45769.1] | 36/55 |
| WP_003231824.1 | 225 | MBL fold metallo-hydrolase | BaeB [CAG23949.2] | 65/80 |
| WP_029317808.1 | 83 | acyl carrier protein | ChxE [AFO59866.1] | 43/51 |
| WP_029317809.1 | 288 | malonyl-CoA-acyltransferase | BaeC [CAG23950.2] | 73/87 |
| WP_029317810.1 | 324 | acyltransferase | BaeD [CAG23951.1] | 59/73 |
| WP_029317811.1 | 767 | ACP-S-malonyltransferase | BaeE [CAG23952.1] | 69/80 |
| WP_003231809.1 | 82 | acyl carrier protein | AcpK [CAG23953.1] | 69/86 |
| WP_029317812.1 | 415 | beta-ketoacyl synthase | CalW [BAP05575.1] | 61/78 |
| WP_029317813.1 | 420 | HMG-CoA synthase | BaeG [CAG23954.2] | 83/91 |
| WP_080265352.1 | 259 | enoyl-CoA hydratase | BaeH [CAG23956.1] | 66/80 |
| WP_029317815.1 | 249 | enoyl-CoA hydratase | BaeI [CAG23956.1] | 75/87 |
| WP_029317816.1 | 5042 | NRPS | BaeJ [CAG23957.2] | 63/76 |
| WP_080265353.1 | 4540 | AT-less type I PKS | BaeL [CAG23958.2] | 63/77 |
| WP_029317818.1 | 4262 | AT-less type I PKS | BaeM [CAG23959.2] | 62/76 |
| WP_080265366.1 | 5488 | NRPS | BaeN [CAG23960.2] | 27/28 |
| WP_029317820.1 | 2543 | methyltransferase | BaeR [CAG23961.2] | 58/73 |
| WP_029317821.1 | 405 | cytochrome P450 | BaeS [CAG23962.1] | 75/86 |
| WP_003231786.1 | 118 | hypothetical protein | / | / |
| WP_029317822.1 | 274 | MBL fold metallo-hydrolase | Nosperin [ADA69237.1] | 30/41 |
| WP_029317823.1 | 442 | serine protease | Leinamycin [AAN85481.1] | 39/55 |
| WP_014476869.1 | 77 | hypothetical protein | / | / |

[a] Number of amino acids

**Table S30.** Predicted functions of ORFs in the NZ_LPJL01000073.1 containing WP_060317160.1

| gene | aa[a] | putative function | Protein homologue | %identity/ %similarity |
|---|---|---|---|---|
| WP_060131219.1 | 177 | VOC family protein | / | / |
| WP_080432263.1 | 428 | beta-ketoacyl synthase | CorD [ADI59526.1] | 43/56 |
| WP_060317143.1 | 77 | acyl carrier protein | ElaE [AEC04351.1] | 30/56 |
| WP_060169178.1 | 409 | HMG-CoA synthase | LnmM [AAN85526.1] | 58/73 |
| WP_059896899.1 | 89 | acyl carrier protein | LnmL [AAN85525.1] | 49/67 |
| WP_063900705.1 | 318 | AT/DC | LnmK [AAN85524.1] | 44/58 |
| WP_060317146.1 | 6915 | AT-less type I PKS | LnmJ [AAN85523.1] | 42/55 |
| WP_060317149.1 | 4171 | NRPS | LnmI [AAN85522.1] | 38/49 |
| WP_060317152.1 | 365 | alpha/beta hydrolase | MupV [AAM12938.1] | 25/40 |
| WP_059998239.1 | 313 | chlorinating enzyme | / | / |
| WP_060317154.1 | 609 | NRPS | SgvD2 [AGN74876.1] | 47/58 |
| WP_060317157.1 | 769 | AT/Ox | DifA [CAG23974.1] | 44/61 |
| WP_059896885.1 | 295 | S-malonyltransferase | PsyH [ADA82589.1] | 46/65 |
| WP_060169188.1 | 268 | enoyl-CoA hydratase | JamI [AAS98780.1] | 33/52 |
| WP_060317160.1 | 410 | HMG-CoA synthase | JamH [AAS98779.1] | 50/66 |
| WP_060317163.1 | 267 | PPTase | SnaN [CBW45738.1] | 37/44 |
| WP_080432264.1 | 134 | DUF3224 family protein | / | / |

[a] Number of amino acids

**Table S31.** Predicted functions of ORFs in the NZ_UETD01000006.1 containing WP_109726276.1 and WP_109726283.1

| gene | aa [a] | putative function | Protein homologue | %identity/ %similarity |
|---|---|---|---|---|
| WP_109726271.1 | 346 | 3-dehydroquinate synthase | / | / |
| WP_109726272.1 | 560 | GGDEF domain-containing protein | / | / |
| WP_109726273.1 | 1674 | AT-less type I PKS | MisC [AKQ22699.1] | 32/51 |
| WP_109726274.1 | 1947 | AT-less type I PKS | BasE [ERM18797.1] | 34/51 |
| WP_109726275.1 | 2820 | AT-less type I PKS | DifF [CAG23977.1] | 28/45 |
| WP_109726276.1 | 409 | HMG-CoA synthase | CorE [ADI59527.1] | 47/67 |
| WP_109726277.1 | 457 | acyltransferase | BaeE [CAG23952.1] | 48/71 |
| WP_109726278.1 | 1704 | methyltransferase | CorL [ADI59534.1] | 38/56 |
| WP_109726279.1 | 1722 | AT-less type I PKS | MisD [AKQ22698.1] | 35/52 |
| WP_109726280.1 | 3086 | AT-less type I PKS | LglD [AIU36100.1] | 35/54 |
| WP_109726281.1 | 1470 | AT-less type I PKS | DifJ [CAJ57410.1] | 31/50 |
| WP_109726282.1 | 2316 | AT-less type I PKS | SorA [ADN68476.1] | 32/50 |
| WP_022495056.1 | 252 | enoyl-CoA hydratase | MxnF [AGS77286.1] | 37/62 |
| WP_109726283.1 | 410 | HMG-CoA synthase | CorE [ADI59527.1] | 53/72 |
| WP_109726284.1 | 423 | beta-ketoacyl synthase | CorD [ADI59526.1] | 44/61 |
| WP_022495055.1 | 80 | magnesium and cobalt efflux protein | CorC [ADI59525.1] | 33/63 |
| WP_109726285.1 | 1096 | AT/Ox | MmpC [AAM12912.1] | 44/62 |
| WP_109726286.1 | 321 | electron transfer flavoprotein | / | / |
| WP_109726287.1 | 249 | electron transfer flavoprotein | / | / |
| WP_109726288.1 | 377 | acyl-CoA dehydrogenase | OzmD [ABA39084.2] | 24/43 |
| WP_109726289.1 | 342 | glyceroyl transferase/phosphatase | OzmB [ABA39082.2] | 46/64 |
| WP_022495050.1 | 282 | 3-hydroxyacyl-CoA dehydrogenase | OzmG [ABS90469.1] | 33/54 |
| WP_109726290.1 | 228 | MBL fold metallo-hydrolase | BaeB [CAG23949.2] | 41/56 |
| WP_109726291.1 | 756 | ABC transporter permease | / | / |
| WP_109726399.1 | 263 | ABC transporter ATP-binding protein | LnmR [AAN85531.1] | 31/50 |
| WP_109726292.1 | 1985 | NRPS | OzmL [ABS90473.1] | 34/54 |
| WP_109726293.1 | 700 | AT-less type I PKS | NspD [ADA69241.1] | 32/49 |
| WP_109726294.1 | 1387 | AT-less type I PKS | LglD [AIU36100.1] | 36/54 |
| WP_109726295.1 | 1847 | AT-less type I PKS | CorL [ADI59534.1] | 33/50 |
| WP_109726296.1 | 406 | acyltransferase | FenF [AAF08794.1] | 32/52 |
| WP_109726400.1 | 79 | acyl carrier protein | OzmE [ABS90467.1] | 37/61 |
| WP_109726297.1 | 228 | PPTase | snaN [CBW45738.1] | 31/47 |
| WP_109726298.1 | 245 | glycosyltransferase | marinomycin [BAG50452.1] | 29/47 |
| WP_109726299.1 | 723 | STAS domain-containing protein | / | / |
| WP_109726300.1 | 264 | AraC family transcriptional regulator | / | / |

[a] Number of amino acids

**Table S32.** Predicted functions of ORFs in the KF264550.1 containing AGS49612.1

| gene | aa[a] | putative function | protein homologue | %identity/ %similarity |
|---|---|---|---|---|
| AGS49592.1 | 114 | putative non-ribosomal peptide synthetase | SnbC [CBW45648.1] | 44/60 |
| AGS49593.1 | 399 | putative cytochrome P450 hydroxylase | SnbF [CBW45756.1] | 61/75 |
| AGS49594.1 | 498 | putative transporter | SnbR [CBW45761.1] | 55/70 |
| AGS49595.1 | 229 | lysine cyclodeaminase | PipA [CBW45757.1] | 44/54 |
| AGS49596.1 | 77 | isochorismatase [enterobactin] siderophore | SgvU [AGN74879.1] | 43/60 |
| AGS49597.1 | 360 | luciferase-like protein | VirN [BAF50713.1] | 65/78 |
| AGS49598.1 | 381 | sarcosine oxidase | SgvS [AGN74901.1] | 57/68 |
| AGS49599.1 | 298 | putative regulatory protein | PapR2 [CBW45736.1] | 54/64 |
| AGS49600.1 | 524 | long-chain-fatty-acid--CoA ligase | SgvD1 [AGN74880.1] | 62/72 |
| AGS49601.1 | 406 | aminotransferase | HPAA [CBW45759.1] | 65/78 |
| AGS49602.1 | 60 | 4-oxalocrotonate tautomerase | SnbT [CBW45760.1] | 59/82 |
| AGS49603.1 | 457 | drug resistance transporter | SgvT3 [AGN74899.1] | 50/63 |
| AGS49604.1 | 277 | TTPase | VirK [BAF50717.1] | 55/64 |
| AGS49605.1 | 262 | thioesterase | VirJ [BAF50718.1] | 60/69 |
| AGS49606.1 | 284 | malonyl CoA-ACP transacylase | SnaM [CBW45739.1] | 66/79 |
| AGS49607.1 | 2195 | NRPS (C-A-PCP-E-C-PCP-TE) | SnaD [CBW45640.1] | 44/56 |
| AGS49608.1 | 2371 | NRPS (C-C-A-PCP-PCP-KS) | VirH [BAF50720.1] | 54/63 |
| AGS49609.1 | 1706 | AT-less type I PKS | SnaE3 [CBW45741.1] | 53/62 |
| AGS49610.1 | 248 | enoyl-CoA hydratase | VirE [BAF50723.1] | 73/80 |
| AGS49611.1 | 244 | enoyl-CoA hydratase/isomerase | VirD [BAF50724.1] | 59/68 |
| AGS49612.1 | 419 | HMG-CoA synthase | VirC [BAF50725.1] | 81/87 |
| AGS49613.1 | 384 | beta-ketoacyl synthase | SnaH [CBW45746.1] | 62/69 |
| AGS49614.1 | 79 | acyl carrier protein | SnaG [CBW45747.1] | 62/74 |
| AGS49615.1 | 68 | acyl carrier protein | VirA [BAF50727.1] | 45/61 |

[a] Number of amino acids

**Table S32.** Predicted functions of ORFs in the CP009283.1 containing AIQ47425.1

| gene | aa[a] | putative function | Protein homologue | %identity/ %similarity |
|---|---|---|---|---|
| AIQ47414.1 | 583 | indolepyruvate ferredoxin oxidoreductase | ThaR [ABC36202.1] | 21/37 |
| AIQ47415.1 | 196 | hypothetical protein | / | / |
| AIQ47416.1 | 2079 | AT-less type I PKS | DifJ [CAJ57410.1] | 35/52 |
| AIQ47417.1 | 727 | hypothetical protein | OocJ [AFX60332.1] | 40/56 |
| AIQ47418.1 | 4066 | AT-less type I PKS | OzmN [ABS90475.1] | 34/49 |
| AIQ47419.1 | 3000 | AT-less type I PKS | SorH [ADN68483.1] | 35/50 |
| AIQ47420.1 | 870 | alpha/beta fold hydrolase | JamJ [AAS98781.1] | 32/49 |
| AIQ47421.1 | 225 | MBL fold metallo-hydrolase | BaeB [CAG23949.2] | 47/66 |
| AIQ47422.1 | 844 | AT/Ox | ChiA [AAY89048.1] | 45/62 |
| AIQ47423.1 | 79 | acyl carrier protein | OocG [AFX60329.1] | 52/73 |
| AIQ47424.1 | 412 | beta-ketoacyl synthase | DipR [AGS06821.1] | 42/61 |
| AIQ47425.1 | 410 | HMG-CoA synthase | CorE [ADI59527.1] | 57/72 |
| AIQ47426.1 | 259 | enoyl-CoA hydratase | JamI [AAS98780.1] | 39/59 |
| AIQ47427.1 | 170 | hypothetical protein | / | / |
| AIQ47428.1 | 319 | electron transfer flavoprotein subunit | / | / |
| AIQ47429.1 | 89 | acyl carrier protein | / | / |
| AIQ47430.1 | 349 | beta-ketoacyl synthase | MxnB [AGS77282.1] | 25/41 |
| AIQ47431.1 | 346 | hypothetical protein | / | / |
| AIQ47432.1 | 542 | AMP-binding protein | SgvD1 [AGN74880.1] | 28/48 |
| AIQ47433.1 | 88 | acyl carrier protein | / | / |
| AIQ47434.1 | 460 | coproporphyrinogen dehydrogenase | / | / |
| AIQ47435.1 | 356 | hypothetical protein | / | / |

[a] Number of amino acids

**Table S33.** Predicted functions of ORFs in the CP012201.1 containing AKU21008.1

| gene | aa[a] | putative function | Protein homologue | %identity/ %similarity |
|---|---|---|---|---|
| AKU24700.1 | 4943 | AT-less type I PKS | ThaO [ABC34675.1] | 51/61 |
| AKU24701.1 | 252 | enoyl-CoA hydratase | ThaM [ABC35867.1] | 67/78 |
| AKU21007.1 | 255 | enoyl-CoA hydratase | ThaM [ABC35867.1] | 60/69 |
| AKU21008.1 | 419 | HMG-CoA synthase | ThaK [ABC34601.1] | 80/90 |
| AKU21009.1 | 412 | hypothetical protein | ThaJ [ABC34518.1] | 63/74 |
| AKU21010.1 | 84 | acyl carrier protein | ThaI [ABC35804.1] | 58/74 |
| AKU21011.1 | 3503 | NRPS | ThaH [ABC35522.1] | 50/61 |
| AKU21012.1 | 5153 | AT-less type I PKS | MmpD [AAM12913.1] | 38/51 |
| AKU24702.1 | 1134 | S-malonyltransferase | BasH [ERM18800.1] | 43/62 |
| AKU21013.1 | 1163 | indolepyruvate ferredoxin oxidoreductase | ThaR [ABC36202.1] | 59/71 |
| AKU21014.1 | 178 | carboxylic acid reductase | ChxJ [ [AFO59871.1]] | 71/81 |
| AKU21015.1 | 394 | acetyl-CoA acetyltransferase | / | / |

[a] Number of amino acids

**Table S34.** Predicted functions of ORFs in the AP014633.1 containing BAP56516.1

| gene | aa[a] | putative function | Protein homologue | %identity/ %similarity |
|---|---|---|---|---|
| BAP56496.1 | 222 | transposase, IS1 family | / | / |
| BAP56497.1 | 429 | phosphoserine phosphatase | Misakinolide [AKQ22694.1] | 52/66 |
| BAP56498.1 | 4490 | AT-less type I PKS | OocJ [AFX60332.1] | 41/57 |
| BAP56499.1 | 6326 | AT-less type I PKS | BaeN [CAG23960.2] | 42/60 |
| BAP56500.1 | 263 | O-methyltransferase | Nosperin [ADA69243.1] | 59/74 |
| BAP56501.1 | 284 | 3-hydroxyacyl-CoA dehydrogenase | OzmG [ABS90469.1] | 33/52 |
| BAP56502.1 | 80 | acyl carrier protein | OzmE [ABS90467.1] | 41/66 |
| BAP56503.1 | 387 | acyl-CoA dehydrogenase | OzmD [ABA39084.2] | 31/49 |
| BAP56504.1 | 342 | glyceroyl transferase/phosphatase | OzmB [ABA39082.2] | 42/60 |
| BAP56505.1 | 2925 | AT-less type I PKS | BryA [ABM63537.1] | 43/60 |
| BAP56506.1 | 6150 | AT-less type I PKS | TaP [ABF88102.1] | 39/56 |
| BAP56507.1 | 128 | transposase family protein | / | / |
| BAP56508.1 | 138 | transposase IS4 | Misakinolide [AKQ22706.1] | 41/58 |
| BAP56509.1 | 210 | transposase family protein | SgvE1 [AGN74892.1] | 32/50 |
| BAP56510.1 | 103 | transposase, IS1 family | SorG [ADN68482.1] | 30/37 |
| BAP56511.1 | 73 | hypothetical protein | / | / |
| BAP56512.1 | 761 | AT/Ox | DifA [CAG23974.1] | 58/75 |
| BAP56513.1 | 82 | transposase | Misakinolide [AKQ22709.1] | 40/55 |
| BAP56514.1 | 422 | beta-ketoacyl synthase | DipR [AGS06821.1] | 50/68 |
| BAP56515.1 | 100 | Transposase | CalK [BAP05587.1] | 43/59 |
| BAP56516.1 | 419 | HMG-CoA synthase | BaeG [CAG23954.2] | 77/85 |
| BAP56517.1 | 251 | enoyl-CoA hydratase | BaeH [CAG23956.1] | 59/75 |
| BAP56518.1 | 63 | Transposase | / | / |

[a] Number of amino acids

**Table S35.** Predicted functions of ORFs in the AP018449.1 containing BBB92534.1

| gene | aa [a] | putative function | Protein homologue | %identity/ %similarity |
|---|---|---|---|---|
| BBB92531.1 | 361 | HTH-type transcriptional activator RhaS | / | / |
| BBB92532.1 | 145 | transcription antitermination protein RfaH | / | / |
| BBB92533.1 | 783 | AT/Ox | DifA [CAG23974.1] | 59/75 |
| BBB92534.1 | 420 | HMG-CoA synthase | BaeG [CAG23954.2] | 76/84 |
| BBB92535.1 | 254 | enoyl-CoA hydratase | BaeH [CAG23955.1] | 59/75 |
| BBB92536.1 | 2902 | AT-less type I PKS | NspC [ADA69239.2] | 4361 |
| BBB92537.1 | 3747 | AT-less type I PKS | BryB [ABM63527.1] | 45/62 |
| BBB92538.1 | 4444 | AT-less type I PKS | BaeN [CAG23960.2] | 39/57 |
| BBB92539.1 | 1877 | AT-less type I PKS | SorH [ADN68483.1] | 53/68 |
| BBB92540.1 | 1183 | AT-less type I PKS | SorI [ADN68484.1] | 43/61 |
| BBB92541.1 | 3680 | AT-less type I PKS | BaeN [CAG23960.2] | 51/68 |
| BBB92542.1 | 1281 | AT-less type I PKS | DifL [CAG23983.1] | 42/59 |
| BBB92543.1 | 249 | enoyl-CoA isomerase | BatE [ADD82946.1] | 68/84 |
| BBB92544.1 | 330 | hypothetical protein | / | / |
| BBB92545.1 | 166 | hypothetical protein | / | / |

[a] Number of amino acids

**Table S36.** Predicted functions of ORFs in the AP018449.1 containing BBB93452.1

| gene | aa [a] | putative function | Protein homologue | %identity/ %similarity |
|---|---|---|---|---|
| BBB93436.1 | 340 | methyltransferase | / | / |
| BBB93437.1 | 350 | uroporphyrinogen decarboxylase | / | / |
| BBB93438.1 | 7951 | AT-less type I PKS | BaeN [CAG23960.2] | 43/59 |
| BBB93439.1 | 5036 | AT-less type I PKS | bonA [AFN27480.1] | 47/61 |
| BBB93440.1 | 3421 | AT-less type I PKS | BaeN [CAG23960.2] | 47/64 |
| BBB93441.1 | 4417 | AT-less type I PKS | BaeN [CAG23960.2] | 37/54 |
| BBB93442.1 | 80 | chorismate lyase | ElaI [AEC04355.1] | 50/71 |
| BBB93443.1 | 352 | hypothetical protein | / | / |
| BBB93444.1 | 302 | hypothetical protein | / | / |
| BBB93445.1 | 124 | hypothetical protein | / | / |
| BBB93446.1 | 108 | hypothetical protein | / | / |
| BBB93447.1 | 140 | hypothetical protein | / | / |
| BBB93448.1 | 127 | hypothetical protein | / | / |
| BBB93449.1 | 114 | carboxymuconolactone decarboxylase | / | / |
| BBB93450.1 | 1138 | AT-less type I PKS | Bat3 [ADD82941.1] | 49/56 |
| BBB93451.1 | 254 | enoyl-CoA hydratase | BaeH [CAG23955.1] | 60/76 |
| BBB93452.1 | 418 | HMG-CoA synthase | BatC [ADD82944.1] | 77/87 |
| BBB93453.1 | 3806 | AT-less type I PKS | BaeN [CAG23960.2] | 42/60 |
| BBB93454.1 | 1108 | AT-less type I PKS | BaeN [CAG23960.2] | 45/63 |
| BBB93455.1 | 1189 | AT-less type I PKS | MisF [AKQ22696.1] | 51/67 |
| BBB93456.1 | 9197 | AT-less type I PKS | TaO [ABF92489.1] | 4057 |
| BBB93457.1 | 5126 | AT-less type I PKS | TaO [ABF92489.1] | 37/57 |
| BBB93458.1 | 137 | transcription antitermination protein RfaH | / | / |
| BBB93459.1 | 412 | malonyl-ACP decarboxylase | CalW [BAP05575.1] | 63/80 |
| BBB93460.1 | 81 | acyl carrier protein | AcpK [CAG23953.1] | 60/75 |
| BBB93461.1 | 285 | malonyl-CoA-acyltransferase | BaeC [CAG23950.2] | 66/80 |
| BBB93462.1 | 466 | AT/Ox | DifA [CAG23974.1] | 67/82 |
| BBB93463.1 | 323 | acyltransferase | BaeD [CAG23951.1] | 46/65 |
| BBB93464.1 | 256 | PPTase | Mis12 [AKQ22701.1] | 37/57 |
| BBB93465.1 | 224 | zinc-dependent hydrolase | BaeB [CAG23949.2] | 50/68 |
| BBB93466.1 | 260 | Ser/Thr protein phosphatase | ThaC [ABC35295.1] | 43/59 |
| BBB93467.1 | 304 | macrolide export protein MacA | / | / |
| BBB93468.1 | 810 | macrolide export ATP-binding protein MacB | / | / |

[a] Number of amino acids

**Table S37.** Predicted functions of ORFs in the FR902444.1 containing CDF01048.1

| gene | aa [a] | putative function | Protein homologue | %identity/ %similarity |
|---|---|---|---|---|
| CDF01045.1 | 805 | AT-less type I PKS | RhiB [CAL69889.1] | 23/43 |
| CDF01046.1 | 1704 | AT-less type I PKS | CorL [ADI59534.1] | 38/56 |
| CDF01047.1 | 457 | S-malonyltransferase | BaeE [CAG23952.1] | 48/71 |
| CDF01048.1 | 417 | HMG-CoA synthase | CorE [ADI59527.1] | 46/66 |

[a] Number of amino acids

**Table S38.** Predicted functions of ORFs in the CXWA01000005.1 containing CTQ60454.1

| gene | aa[a] | putative function | Protein homologue | %identity/ %similarity |
|---|---|---|---|---|
| CTQ60441.1 | 468 | Leucine-specific-binding protein precursor | / | / |
| CTQ60442.1 | 288 | HflK protein | / | / |
| CTQ60443.1 | 6838 | AT-less type I PKS | Myxovirescin [ABF92489.1] | 39/55 |
| CTQ60444.1 | 444 | oxidoreductase | PedG [AAS47561.1] | 64/77 |
| CTQ60445.1 | 6892 | AT-less type I PKS | Dor5 [ACY01390.1] | 44/57 |
| CTQ60446.1 | 778 | AT/Ox | CalY [BAP05573.1] | 48/67 |
| CTQ60447.1 | 549 | ABC transporter ATP-binding protein | / | / |
| CTQ60448.1 | 544 | ABC transporter ATP-binding protein | / | / |
| CTQ60449.1 | 82 | acyl carrier protein | ChxC [AFO59864.1] | 47/75 |
| CTQ60450.1 | 654 | asparagine synthetase | SmdH [CCC21122.1] | 57/72 |
| CTQ60451.1 | 256 | enoyl-CoA hydratase | BaeH [CAG23955.1] | 52/65 |
| CTQ60452.1 | 422 | beta-ketoacyl synthase | NspH [ADA69244.1] | 44/65 |
| CTQ60453.1 | 85 | acyl carrier protein | CalX [BAP05574.1] | 41/70 |
| CTQ60454.1 | 418 | HMG-CoA synthase | ThaK [ABC34601.1] | 61/76 |
| CTQ60455.1 | 429 | cytochrome P450 | ElaG [AEC04353.1] | 33/52 |
| CTQ60456.1 | 274 | Ser/Thr protein phosphatase | ThaC [ABC35295.1] | 32/45 |
| CTQ60457.1 | 269 | PPTase | BatI [ADD82950.1] | 35/51 |
| CTQ60458.1 | 299 | acyltransferase | OzmM [ABS90474.1] | 37/54 |
| CTQ60459.1 | 412 | cytochrome P450 | BaeS [CAG23962.1] | 33/52 |
| CTQ60460.1 | 425 | cytochrome P450 | ElaG [AEC04353.1] | 29/47 |
| CTQ60461.1 | 433 | cytochrome P450 | BaeS [CAG23962.1] | 34/50 |
| CTQ60462.1 | 341 | 31-O-demethyl-FK506 methyltransferase | CalA [BAP05589.1] | 32/49 |

[a] Number of amino acids

**Table S39.** Predicted functions of ORFs in the AJRJ01000088.1 containing EIM08809.1

| gene | aa[a] | putative function | Protein homologue | %identity/ %similarity |
|---|---|---|---|---|
| EIM08806.1 | 3398 | AT-less type I PKS | BaeJ [CAG23957.2] | 63/77 |
| EIM08807.1 | 248 | enoyl-CoA hydratase | BaeI [CAG23956.1] | 76/88 |
| EIM08808.1 | 255 | enoyl-CoA hydratase | BaeH [CAG23956.1] | 67/83 |
| EIM08809.1 | 420 | HMG-CoA synthase | BaeG [CAG23954.2] | 86/93 |
| EIM08810.1 | 415 | beta-ketoacyl synthase | CalW [BAP05575.1] | 64/80 |
| EIM08811.1 | 82 | acyl carrier protein | AcpK [CAG23953.1] | 72/85 |
| EIM08812.1 | 787 | S-malonyltransferase | BaeE [CAG23952.1] | 66/79 |
| EIM08813.1 | 321 | acyltransferase | BaeD [CAG23951.1] | 58/75 |
| EIM08814.1 | 288 | malonyl-CoA-acyltransferase | BaeC [CAG23950.2] | 76/88 |
| EIM08815.1 | 225 | putative hydrolase | BaeB [CAG23949.2] | 70/80 |
| EIM08816.1 | 196 | putative transcriptional regulator | / | / |

[a] Number of amino acids

**Table S40.** Predicted functions of ORFs in the AMRL01000010.1 containing EKE75767.1

| gene | aa[a] | putative function | Protein homologue | %identity/ %similarity |
|---|---|---|---|---|
| EKE75756.1 | 3800 | hybrid NRPS/polyketide synthase | Ta1 [ABF85931.1] | 39/53 |
| EKE75757.1 | 444 | flavin-containing monooxygenase | PedG [AAS47561.1] | 62/76 |
| EKE75758.1 | 6700 | AT-less type I PKS | Dor5 [ACY01390.1] | 47/58 |
| EKE75759.1 | 768 | AT/Ox | CalY [BAP05573.1] | 54/71 |
| EKE75760.1 | 550 | cyclic peptide transporter | Rhizopodin [CCA89322.1] | 27/47 |
| EKE75761.1 | 549 | cyclic peptide transporter | CalU [BAP05577.1] | 21/39 |
| EKE75762.1 | 84 | acyl carrier protein | SmdG [CCC21121.1] | 55/73 |
| EKE75763.1 | 656 | asparagine synthetase | SmdH [CCC21122.1] | 59/71 |
| EKE75764.1 | 249 | enoyl-CoA hydratase | BatD [ADD82945.1] | 53/68 |
| EKE75765.1 | 400 | beta-ketoacyl synthase | NspH [ADA69244.1] | 46//62 |
| EKE75766.1 | 81 | acyl carrier protein | CalX [BAP05574.1] | 47/74 |
| EKE75767.1 | 422 | HMG-CoA synthase | JamH [AAS98779.1] | 63/79 |
| EKE75768.1 | 415 | cytochrome P450 | BaeS [CAG23962.1] | 34/50 |
| EKE75769.1 | 277 | non-ribosomal peptide synthetase | CalA [BAP05589.1] | 34/53 |

[a] Number of amino acids

**Table S41.** Predicted functions of ORFs in the AQRA01000010.1 containing EZH71961.1

| gene | aa[a] | putative function | Protein homologue | %identity/ %similarity |
|---|---|---|---|---|
| EZH71952.1 | 219 | hypothetical protein ATO12_04855 | MupN [AAM12928.1] | 26/47 |
| EZH71953.1 | 309 | phosphoglyceromutase | / | / |
| EZH71954.1 | 396 | malonyl CoA-ACP transacylase | MisG [AKQ22695.1] | 51/68 |
| EZH71955.1 | 424 | beta-ketoacyl synthase | DipR [AGS06821.1] | 50/71 |
| EZH71956.1 | 86 | acyl carrier protein | DipF [AGS06833.1] | 47/77 |
| EZH71957.1 | 241 | hydroxyacylglutathione hydrolase | BaeB [CAG23949.2] | 42/58 |
| EZH71958.1 | 316 | acyltransferase | BaeD [CAG23951.1] | 35/56 |
| EZH71959.1 | 450 | hypothetical protein | / | / |
| EZH71960.1 | 261 | enoyl-CoA hydratase | BaeH [CAG23956.1] | 57/69 |
| EZH71961.1 | 419 | HMG-CoA synthase | BatC [ADD82944.1] | 70/83 |
| EZH71962.1 | 664 | aminotransferase | BryA [ABM63537.1] | 26/40 |
| EZH71963.1 | 206 | class I SAM-dependent methyltransferase | PedA [AAS47557.1] | 22/39 |
| EZH71964.1 | 298 | cysteine desulfurase | SgvL [AGN74881.1] | 24/47 |
| EZH71965.1 | 273 | hypothetical protein ATO12_04930 | / | / |
| EZH71966.1 | 4622 | AT-less type I PKS | SorH [AND68483.1] | 34/52 |
| EZH71967.1 | 412 | hypothetical protein ATO12_04940 | / | / |
| EZH71968.1 | 4747 | AT-less type I PKS | TaO [ABF92489.1] | 37/55 |
| EZH71969.1 | 3529 | AT-less type I PKS | SorE [AND68480.1] | 43/59 |
| EZH71970.1 | 6114 | AT-less type I PKS | MisF [AKQ22696.1] | 33/52 |

[a] Number of amino acids

**Table S42.** Predicted functions of ORFs in the JGVN01000004.1 containing EZQ03740.1 and EZQ03561.1

| gene | aa[a] | putative function | Protein homologue | %identity/ %similarity |
|---|---|---|---|---|
| EZQ03555.1 | 333 | ABC transporter substrate-binding protein | / | / |
| EZQ03556.1 | 899 | S-malonyltransferase | OocV [AFX60344.1] | 41/59 |
| EZQ03730.1 | 2473 | AT-less type I PKS | ElaK [AEC04357.1] | 38/53 |
| EZQ03731.1 | 130 | hypothetical protein | / | / |
| EZQ03557.1 | 87 | acyl carrier protein | DipF [AGS06833.1] | 30/56 |
| EZQ03558.1 | 260 | enoyl-CoA hydratase | CorF [ADI59528.1] | 40/57 |
| EZQ03732.1 | 338 | hypothetical protein | KirAI [CAN89631.1] | 33/47 |
| EZQ03559.1 | 472 | hypothetical protein | / | / |
| EZQ03560.1 | 429 | monooxygenase | PedG [AAS47561.1] | 47/63 |
| EZQ03733.1 | 5453 | AT-less type I PKS | SorA [ADN68476.1] | 36/50 |
| EZQ03734.1 | 81 | hypothetical protein | / | / |
| EZQ03735.1 | 65 | hypothetical protein | / | / |
| EZQ03736.1 | 91 | hypothetical protein | / | / |
| EZQ03737.1 | 5985 | AT-less type I PKS | MisC [AKQ22699.1] | 38/52 |
| EZQ03738.1 | 4294 | AT-less type I PKS | OzmN [ABS90475.1] | 40/51 |
| EZQ03739.1 | 464 | hypothetical protein | / | / |
| EZQ03740.1 | 601 | HMG-CoA synthase | Rhizopodin [CCA89332.1] | 37/59 |
| EZQ03561.1 | 409 | HMG-CoA synthase | CorE [ADI59527.1] | 54/69 |
| EZQ03741.1 | 869 | hypothetical protein | MisC [AKQ22699.1] | 37/49 |
| EZQ03562.1 | 133 | hypothetical protein | / | / |
| EZQ03563.1 | 222 | PPTase | BatI [ADD82950.1] | 38/54 |
| EZQ03564.1 | 369 | transcriptional regulator | oxazolomycin [ABS90464.1] | 28/41 |
| EZQ03565.1 | 344 | deacylase | / | / |
| EZQ03566.1 | 144 | endoribonuclease L-PSP | / | / |

[a] Number of amino acids

**Table S43.** Predicted functions of ORFs in the BAOS01000010.1 containing GAX60246.1

| gene | aa [a] | putative function | Protein homologue | %identity/ %similarity |
|---|---|---|---|---|
| GAX60242.1 | 893 | beta-ketoacyl synthase | MisF [AKQ22696.1] | 39/57 |
| GAX60243.1 | 731 | hypothetical protein | CylH [ARU81122.1] | 49/66 |
| GAX60244.1 | 254 | enoyl-CoA hydratase | BatE [ADD82946.1] | 57/79 |
| GAX60245.1 | 261 | enoyl-CoA hydratase | CylG [ARU81121.1] | 54/72 |
| GAX60246.1 | 413 | HMG-CoA synthase | CylF [ARU81120.1] | 63/76 |
| GAX60247.1 | 1389 | AT-less type I PKS | MisF [AKQ22696.1] | 48/65 |
| GAX60248.1 | 1659 | AT-less type I PKS | MisF [AKQ22696.1] | 48/65 |
| GAX60249.1 | 2797 | AT-less type I PKS | TaO [ABF92489.1] | 40/58 |
| GAX60250.1 | 3734 | AT-less type I PKS | SorE [ADN68480.1] | 44/60 |
| GAX60251.1 | 2136 | NRPS | DszC [AAY32966.1] | 34/53 |
| GAX60252.1 | 518 | Fe-S oxidoreductase | / | / |
| GAX60253.1 | 320 | acyltransferase | BaeD [CAG23951.1] | 38/56 |
| GAX60254.1 | 736 | cyclic nucleotide-binding protein | / | / |
| GAX60255.1 | 262 | hypothetical protein | / | / |
| GAX60256.1 | 213 | GCN5-related N-acetyltransferase | / | / |
| GAX60257.1 | 206 | N-acetylmuramoyl-L-alanine amidase | / | / |

[a] Number of amino acids

**Table S44.** Predicted functions of ORFs in the KN150850.1 containing KGC14077.1

| gene | aa[a] | putative function | Protein homologue | %identity/ %similarity |
|---|---|---|---|---|
| KGC24078.1 | 178 | ABC transporter substrate-binding protein | / | / |
| KGC24079.1 | 3641 | AT-less type I PKS | MisF [AKQ22696.1] | 35/54 |
| KGC24080.1 | 2358 | AT-less type I PKS | ElaO [AEC04361.1] | 43/57 |
| KGC24074.1 | 2745 | AT-less type I PKS | PedF [AAS47564.1] | 38/52 |
| KGC24076.1 | 1660 | AT-less type I PKS | SorB [ADN68477.1] | 47/59 |
| KGC13006.1 | 486 | polyketide synthase | / | / |
| KGC15096.1 | 1972 | polyketide synthase | / | / |
| KGC13060.1 | 142 | hypothetical protein | / | / |
| KGC12909.1 | 75 | polyketide synthase | / | / |
| KGC12887.1 | 631 | asparagine synthase | Dor4 [ACY01389.1] | 55/67 |
| KGC14223.1 | 298 | acyl transferase | ThaF [ABC34740.1] | 35/51 |
| KGC14725.1 | 88 | acyl carrier protein | smdG [CCC21121.1] | 45/69 |
| KGC12701.1 | 463 | PfaD family | SorN [ADN68488.1] | 69/83 |
| KGC14663.1 | 82 | acyl carrier protein | CalX [BAP05574.1] | 54/78 |
| KGC14795.1 | 409 | beta-ketoacyl synthase | BatB [ADD82943.1] | 60/72 |
| KGC14077.1 | 420 | HMG-CoA synthase | BonG [AFN27479.1] | 70/82 |
| KGC14800.1 | 263 | enoyl-CoA hydratase | BatD [ADD82945.1] | 57/71 |
| KGC14421.1 | 249 | enoyl-CoA isomerase | BaeE [CAG23952.1] | 67/78 |
| KGC14139.1 | 1102 | NACHT domain protein | SorM [ADN68497.1] | 38/51 |
| KGC14204.1 | 120 | hypothetical protein | / | / |
| KGC14532.1 | 467 | amidase family protein | SorP [ADN68490.1] | 46/61 |
| KGC13134.1 | 395 | malonyl CoA-acyl carrier protein | MisG [AKQ22695.1] | 49/64 |
| KGC13462.1 | 285 | PPTase | BatI [ADD82950.1] | 42/56 |
| KGC12823.1 | 325 | acyl transferase | SorO [ADN68489.1] | 45/59 |
| KGC13596.1 | 451 | MATE efflux family protein | SorJ [ADN68485.1] | 31/47 |
| KGC15427.1 | 402 | enoyl-CoA hydratase | / | / |
| KGC14688.1 | 264 | enoyl-CoA hydratase | BaeH [CAG23955.1] | 31/44 |
| KGC12885.1 | 300 | 3-hydroxyisobutyrate dehydrogenase | / | / |
| KGC15246.1 | 509 | methylmalonate-semialdehyde dehydrogenase | pederin [AAS47555.1] | 31/49 |
| KGC13025.1 | 563 | AMP-binding enzyme family | SnaE4 [CBW45740.1] | 29/42 |
| KGC15563.1 | 377 | acyl-CoA dehydrogenase | OzmD [ABA39084.2] | 33/50 |
| KGC15687.1 | 345 | helix-turn-helix domain protein | / | / |

[a] Number of amino acids

**Table S45.** Predicted functions of ORFs in the JSYO01000057.1 containing KIC00730.1

| gene | aa[a] | putative function | Protein homologue | %identity/ %similarity |
|---|---|---|---|---|
| KIC00719.1 | 79 | hypothetical protein | / | / |
| KIC00720.1 | 653 | histidine kinase | / | / |
| KIC00721.1 | 4531 | AT-less type I PKS | BaeN [CAG23960.2] | 37/55 |
| KIC00722.1 | 5426 | AT-less type I PKS | BaeN [CAG23960.2] | 34/52 |
| KIC00723.1 | 3675 | AT-less type I PKS | SorE [ADN68480.1] | 32/51 |
| KIC00724.1 | 62 | hypothetical protein | / | / |
| KIC00725.1 | 600 | cytochrome P450 | Nosperin [ADA69248.1] | 27/43 |
| KIC00726.1 | 436 | hypothetical protein | / | / |
| KIC00727.1 | 756 | S-malonyltransferase | BaeE [CAG23952.1] | 46/65 |
| KIC00728.1 | 439 | hypothetical protein | / | / |
| KIC00729.1 | 77 | acyl carrier protein | AcpK [CAG23953.1] | 46/73 |
| KIC00735.1 | 419 | beta-ketoacyl synthase | JamG [AAS98778.1] | 40/60 |
| KIC00730.1 | 415 | HMG-CoA synthase | BaeG [CAG23954.2] | 43/67 |
| KIC00731.1 | 262 | enoyl-CoA hydratase | JamI [AAS98780.1] | 45/63 |
| KIC00732.1 | 248 | malonyl-CoA-transacylase | BaeE [CAG23952.1] | 52/76 |
| KIC00733.1 | 323 | acyltransferase | BaeD [CAG23951.1] | 34/54 |
| KIC00734.1 | 127 | transposase | / | / |

[a] Number of amino acids

**Table S46.** Predicted functions of ORFs in the JUEB01000029.1 containing KJS64697.1

| gene | aa[a] | putative function | Protein homologue | %identity/ %similarity |
|---|---|---|---|---|
| KJS64692.1 | 423 | glutamate dehydrogenase | / | / |
| KJS64693.1 | 330 | ketol-acid reductoisomerase | Malleilactone [ABC34926.1] | 35/53 |
| KJS64694.1 | 1097 | S-malonyltransferase | BasH [ERM18800.1] | 51/70 |
| KJS64695.1 | 83 | acyl carrier protein | Thailandamide [ABC35804.1] | 41/65 |
| KJS64696.1 | 425 | beta-ketoacyl synthase | DipR [AGS06821.1] | 43/66 |
| KJS64697.1 | 410 | HMG-CoA synthase | MxnE [AGS77285.1] | 60/76 |
| KJS64698.1 | 256 | enoyl-CoA hydratase | JamI [AAS98780.1] | 42/60 |
| KJS64699.1 | 4333 | AT-less type I PKS | RizB [CCA89326.1] | 34/51 |
| KJS64700.1 | 4578 | AT-less type I PKS | DszB [AAY32965.1] | 40/56 |
| KJS64701.1 | 552 | pyridoxal-dependent decarboxylase | JamL [AAS98783.1] | 40/60 |
| KJS64702.1 | 363 | beta-ketoacyl synthase | PsyA [ADA82581.1] | 58/71 |

[a] Number of amino acids

**Table S47.** Predicted functions of ORFs in the JUEB01000006.1 containing KJS69178.1

| gene[a] | aa[b] | putative function | Protein homologue | %identity/ %similarity |
|---|---|---|---|---|
| ORF1 | 1116 | AT-less type I PKS(KS-DH) | TaO [ABF92489.1] | 50/67 |
| KJS69177.1 | 250 | enoyl-CoA hydratase | BaeH [CAG23956.1] | 57/73 |
| KJS69178.1 | 420 | HMG-CoA synthase | BatC [ADD82944.1] | 75/85 |
| KJS69179.1 | 81 | acyl carrier protein | NspG [ADA71314.1] | 55/76 |
| ORF2 | 1116 | AT-less type I PKS(KS-ACP-KS-ACP-ACP) | BaeN [CAG23960.2] | 54/71 |

[a] ORFs are proteins without protein ID provided by NCBI [b] Number of amino acids

**Table S48.** Predicted functions of ORFs in the AMQU01000026.1 containing KKD52735.1

| gene[a] | aa[b] | putative function | Protein homologue | %identity/ %similarity |
|---|---|---|---|---|
| ORF1 | 7518 | hybrid PKS/NRPS (CAL-Acp-C-A-Acp-KS-KR-Acp-KS-Acp-Acp-KS-SH-KR-Acp-KS-DH-KR-Acp-KS) | OnnI [AAV97877.1] | 51/67 |
| KKD52734.1 | 249 | enoyl-CoA hydratase | BaeE [CAG23952.1] | 66/81 |
| KKD53033.1 | 253 | enoyl-CoA hydratase | BaeH [CAG23955.1] | 63/78 |
| KKD52735.1 | 420 | HMG-CoA synthase | BaeG [CAG23954.2] | 78/87 |
| KKD52736.1 | 415 | hypothetical protein | CalW [BAP05575.1] | 63/80 |
| KKD52737.1 | 79 | acyl carrier protein | AcpK [CAG23953.1] | 59/75 |
| KKD52738.1 | 253 | PPTase | Mis12 [AKQ22701.1] | 31/52 |
| KKD52739.1 | 173 | antitermination factor | ElaA [AEC04347.1] | 26/46 |
| KKD52740.1 | 186 | class I SAM-dependent methyltransferase | / | / |
| KKD52741.1 | 90 | hypothetical protein | / | / |

[a] ORFs are proteins without protein ID provided by NCBI [b] Number of amino acids

**Table S49.** Predicted functions of ORFs in the LGKL01000032.1 containing KNA30661.1

| gene | aa[a] | putative function | Protein homologue | %identity/ %similarity |
|---|---|---|---|---|
| KNA30656.1 | 559 | hypothetical protein | / | / |
| KNA30657.1 | 414 | beta-ketoacyl synthase | NspH [ADA69244.1] | 57/73 |
| KNA30658.1 | 79 | acyl carrier protein | ElaE [AEC04351.1] | 56/80 |
| KNA30659.1 | 260 | enoyl-CoA hydratase | BaeE [CAG23952.1] | 61/77 |
| KNA30660.1 | 256 | enoyl-CoA hydratase | OocD [AFX60326.1] | 49/68 |
| KNA30661.1 | 419 | HMG-CoA synthase | ThaK [ABC34601.1] | 70/81 |
| KNA30789.1 | 77 | hypothetical protein | CorK [ADI59533.1] | 38/45 |
| KNA30790.1 | 390 | malonyl CoA-ACP transacylase | fr9O [AIC32701.1] | 61/72 |
| KNA30662.1 | 410 | sodium:proton exchanger | OnnI [AAV97877.1] | 33/44 |
| KNA30663.1 | 4579 | AT-less type I PKS | BasG [ERM18799.1] | 34/50 |
| KNA30664.1 | 5916 | AT-less type I PKS | Ta1 [ABF85931.1] | 41/54 |
| KNA30665.1 | 5041 | AT-less type I PKS | BaeN [CAG23960.2] | 3752 |
| KNA30666.1 | 1564 | AT-less type I PKS | CalE [BAP05593.1] | 50/62 |
| KNA30667.1 | 552 | halogenase | Ena5928 [ABI91473.1] | 24/38 |
| KNA30668.1 | 480 | DSBA oxidoreductase | LnmY [AAN85538.1] | 44/60 |
| KNA30669.1 | 634 | chloride channel protein | / | / |

[a] Number of amino acids

**Table S50.** Predicted functions of ORFs in the JPDT01000569.1 containing KPA17635.1

| gene | aa[a] | putative function | Protein homologue | %identity/ %similarity |
|---|---|---|---|---|
| KPA17623.1 | 155 | hypothetical protein | / | / |
| KPA17624.1 | 71 | short chain dehydrogenase | BatM [ADD82954.1] | 45/66 |
| KPA17625.1 | 603 | hypothetical protein | / | / |
| KPA17626.1 | 57 | hypothetical protein | / | / |
| KPA17627.1 | 675 | phosphohydrolase | / | / |
| KPA17628.1 | 1817 | AT-less type I PKS | DszA [AAY32964.1] | 39/56 |
| KPA17629.1 | 14038 | hybrid PKS/NRPS | CalE [BAP05593.1] | 39/56 |
| KPA17630.1 | 347 | O-methyltransferase | AlbII [CAE52340.1] | 24/43 |
| KPA17631.1 | 1506 | short-chain dehydrogenase | PsyD [ADA82585.1] | 33/51 |
| KPA17632.1 | 561 | phosphotransferase | PsyE [ADA82586.1] | 34/54 |
| KPA17633.1 | 186 | hypothetical protein | / | / |
| KPA17634.1 | 81 | acyl carrier protein | ElaE [AEC04351.1] | 50/70 |
| KPA17635.1 | 419 | HMG-CoA synthase | BaeG [CAG23954.2] | 72/82 |
| KPA17636.1 | 254 | enoyl-CoA hydratase | BaeH [CAG23955.1] | 55/73 |
| KPA17637.1 | 249 | enoyl-CoA hydratase | BaeE [CAG23952.1] | 64/81 |
| KPA17638.1 | 54 | hypothetical protein | / | |
| KPA17639.1 | 178 | GTPase domain-containing protein | / | / |

[a] Number of amino acids

**Table S51.** Predicted functions of ORFs in the LMCY01000011.1 containing KQV49710.1

| gene[a] | aa[b] | putative function | Protein homologue | %identity/ %similarity |
|---|---|---|---|---|
| KQV49706.1 | 185 | hypothetical protein | / | / |
| KQV49707.1 | 94 | hypothetical protein | / | / |
| KQV49708.1 | 249 | enoyl-CoA hydratase | BaeE [CAG23952.1] | 70/85 |
| KQV49709.1 | 263 | enoyl-CoA hydratase | BaeH [CAG23955.1] | 57/71 |
| KQV49710.1 | 420 | HMG-CoA synthase | OnnA [AAV97869.1] | 68/80 |
| KQV49711.1 | 328 | hypothetical protein | / | / |
| KQV49712.1 | 407 | hypothetical protein | / | / |
| KQV49713.1 | 442 | polyketide synthase | Bat3 [ADD82941.1] | 54/68 |
| KQV49714.1 | 296 | hypothetical protein | / | / |
| KQV49715.1 | 828 | hypothetical protein | / | / |
| KQV49716.1 | 4481 | AT-less type I PKS | BaeN [CAG23960.2] | 38/56 |
| KQV49717.1 | 2924 | AT-less type I PKS | pedF [AAS47564.1] | 38/53 |
| KQV49718.1 | 1973 | AT-less type I PKS | OocL [AFX60334.1] | 52/63 |
| ORF1 | 872 | AT-less type I PKS (ECH-ECH-Acp) | OocJ [AFX60332.1] | 38/53 |
| KQV49719.1 | 417 | hypothetical protein | / | / |
| KQV49720.1 | 221 | DUF2306 domain-containing protein | / | / |
| KQV49721.1 | 3689 | AT-less type I PKS | MisF [AKQ22696.1] | 43/58 |
| KQV49722.1 | 247 | hypothetical protein | / | / |
| KQV49723.1 | 217 | hypothetical protein | / | / |
| KQV49724.1 | 435 | hypothetical protein | / | / |
| KQV49725.1 | 317 | hypothetical protein | / | / |
| KQV49726.1 | 2126 | AT-less type I PKS | ElaK [AEC04357.1] | 45/61 |
| KQV49727.1 | 258 | PPTase | BatI [ADD82950.1] | 52/62 |
| KQV49728.1 | 330 | acyltransferase | SorO [ADN68489.1] | 49/66 |
| KQV49729.1 | 372 | S-malonyltransferase | Fr9O [AIC32701.1] | 59/72 |
| KQV49730.1 | 414 | beta-ketoacyl synthase | NspH [ADA69244.1] | 61/77 |
| KQV49731.1 | 79 | acyl carrier protein | FR9M [AIC32699.1] | 59/74 |
| KQV49732.1 | 416 | hypothetical protein | / | / |
| KQV49733.1 | 169 | antitermination factor | ElaA [AEC04347.1] | 30/50 |
| KQV49734.1 | 229 | hypothetical protein | MupR [AAK28504.1] | 33/55 |
| KQV49735.1 | 472 | radical SAM protein | / | / |

[a] ORFs are proteins without protein ID provided by NCBI [b] Number of amino acids

**Table S52.** Predicted functions of ORFs in the LDST01000044.1 containing KTT63895.1

| gene[a] | aa[b] | putative function | Protein homologue | %identity/ %similarity |
|---|---|---|---|---|
| ORF1 | 1394 | AT-less type I PKS (Acp-KS-KR-Acp) | MmpD [AAM12913.1] | 64/73 |
| KTT63947.1 | 403 | NADH oxidase | MupC [AAM12914.1] | 77/86 |
| KTT63889.1 | 92 | acyl carrier protein | MacpA [AAM12915.1] | 77/86 |
| KTT63948.1 | 251 | polyketide synthase | MupD [AAM12916.1] | 64/78 |
| KTT63890.1 | 343 | NADPH:quinone oxidoreductase | MupE [AAM12917.1] | 77/87 |
| KTT63891.1 | 76 | acyl carrier protein | MacpB [AAM12918.1] | 71/81 |
| KTT63892.1 | 335 | short-chain dehydrogenases/reductases | MupF [AAM12919.1] | 68/78 |
| KTT63893.1 | 79 | acyl carrier protein | Macp [AAM12920.1] | 75/86 |
| KTT63894.1 | 412 | beta-ketoacyl synthase | MupG [AAM12921.1] | 80/89 |
| KTT63895.1 | 420 | HMG-CoA synthase | MupH [AAM12922.1] | 84/90 |
| KTT63896.1 | 255 | enoyl-CoA hydratase | MupJ [AAM12923.1] | 73/84 |
| KTT63897.1 | 247 | enoyl-CoA hydratase | MupK [AAM12924.1] | 82/90 |
| KTT63898.1 | 334 | alpha/beta hydrolase | MmpE [AAM12925.1] | 70/83 |
| KTT63899.1 | 1030 | isoleucine--tRNA ligase | MupM [AAM12927.1] | 84/91 |
| KTT63900.1 | 429 | cytochrome P450 | MupO [AAM12929.1] | 82/91 |
| KTT63901.1 | 323 | glyoxalase/bleomycin resistance/dioxygenase | MupP [AAM12930.1] | 58/68 |
| KTT63902.1 | 450 | long-chain fatty acid-CoA ligase | MupQ [AAM12931.1] | 69/80 |
| KTT63903.1 | 253 | SDR family NAD [P]-dependent oxidoreductase | MupS [AAM12932.1] | 77/86 |
| KTT63904.1 | 105 | acyl carrier protein | MacpD [AAM12933.1] | 73/83 |
| KTT63905.1 | 90 | acyl carrier protein | MmpF [AAM12934.1] | 66/76 |
| KTT63949.1 | 107 | Rieske [2Fe-2S] protein | MupT [AAM12936.1] | 77/89 |
| KTT63906.1 | 663 | alpha/beta fold hydrolase | MupV [AAM12938.1] | 77/87 |
| KTT63907.1 | 471 | aromatic ring-hydroxylating dioxygenase subunit | MupW [AAM12939.1] | 84/91 |
| KTT63950.1 | 218 | hypothetical protein | MupR [AAK28504.1] | 68/81 |
| KTT63951.1 | 510 | amidase | MupX [AAM12940.1] | 74/84 |
| KTT63908.1 | 194 | GNAT family N-acetyltransferase | MupI [AAK28505.1] | 71/83 |
| KTT63952.1 | 70 | XRE family transcriptional regulator | / | / |
| KTT63909.1 | 304 | hypothetical protein | / | / |

[a] ORFs are proteins without protein ID provided by NCBI [b] Number of amino acids

**Table S53.** Predicted functions of ORFs in the LQYO01000026.1 containing KYC88112.1

| gene | aa[a] | putative function | Protein homologue | %identity/ %similarity |
|---|---|---|---|---|
| KYC88111.1 | 804 | AT/Ox | DifA [CAG23974.1] | 55/72 |
| KYC88112.1 | 419 | HMG-CoA synthase | BaeG [CAG23954.2] | 47/83 |
| KYC88113.1 | 3057 | AT-less type I PKS | Kirromycin [CAN89656.1] | 35/52 |
| KYC88114.1 | 5945 | AT-less type I PKS | BaeJ [CAG23957.2] | 49/64 |
| KYC88115.1 | 7435 | AT-less type I PKS | NspC [ADA69239.2] | 35/55 |
| KYC88116.1 | 5171 | AT-less type I PKS | TaO [ABF92489.1] | 37/55 |
| KYC88117.1 | 2525 | AT-less type I PKS | Kirromycin [CAN89656.1] | 29/47 |
| KYC88118.1 | 43 | hypothetical protein | / | / |
| KYC88119.1 | 93 | protoheme IX farnesyltransferase | / | / |
| KYC88120.1 | 119 | hypothetical protein | Kirromycin [CAN89610.1] | 41/42 |
| KYC88121.1 | 80 | hypothetical protein | Kirromycin [CAN89610.1] | 26/45 |
| KYC88138.1 | 304 | class A beta-lactamase | / | / |
| KYC88139.1 | 93 | monooxygenase | / | / |
| KYC88140.1 | 470 | MFS transporter | Leinamycin [AAN85487.1] | 24/44 |
| KYC88141.1 | 157 | MarR family transcriptional regulator | / | / |
| KYC88142.1 | 370 | hypothetical protein | / | / |

[a] Number of amino acids

**Table S54.** Predicted functions of ORFs in the MBQV01000032.1 containing OEC77139.1

| gene | aa[a] | putative function | Protein homologue | %identity/ %similarity |
|---|---|---|---|---|
| OEC77135.1 | 239 | DNA-binding transcriptional regulator | / | / |
| OEC77136.1 | 60 | phosphatase RapH inhibitor | / | / |
| OEC77137.1 | 380 | tetratricopeptide repeat protein | / | / |
| OEC77138.1 | 337 | hypothetical protein | / | / |
| OEC77139.1 | 421 | HMG-CoA synthase | BatC [ADD82944.1] | 68/83 |
| OEC77140.1 | 1097 | S-malonyltransferase | BasH [ERM18800.1] | 42/63 |
| OEC77141.1 | 2096 | AT-less type I PKS | ThaO [ABC34675.1] | 38/53 |
| OEC77142.1 | 5906 | AT-less type I PKS | BaeN [CAG23960.2] | 37/56 |
| OEC77143.1 | 1252 | AT-less type I PKS | / | / |

[a] Number of amino acids

**Table S55.** Predicted functions of ORFs in the MKVK01000021.1 containing OJX77765.1

| gene | aa [a] | putative function | Protein homologue | %identity/ %similarity |
|---|---|---|---|---|
| OJX77725.1 | 416 | hypothetical protein | / | / |
| OJX77763.1 | 264 | hypothetical protein | / | / |
| OJX77726.1 | 6228 | AT-less type I PKS | Ta1 [ABF85931.1] | 37/52 |
| OJX77727.1 | 433 | monooxygenase | PedG [AAS47561.1] | 61/77 |
| OJX77728.1 | 6429 | AT-less type I PKS | ChxE [AFO59866.1] | 41/53 |
| OJX77764.1 | 752 | AT/Ox | CalY [BAP05573.1] | 50/65 |
| OJX77729.1 | 551 | ABC transporter | Rhizopodin [CCA89322.1] | 29/52 |
| OJX77730.1 | 544 | ATPase and permease component | CalU [BAP05577.1] | 28/48 |
| OJX77731.1 | 242 | PPTase | BatI [ADD82950.1] | 35/53 |
| OJX77732.1 | 275 | Ser/Thr protein phosphatase | ThaC [ABC35295.1] | 35/53 |
| OJX77733.1 | 267 | hypothetical protein | CalA [BAP05589.1] | 35/53 |
| OJX77734.1 | 400 | cytochrome P450 | BaeS [CAG23962.1] | 35/51 |
| OJX77765.1 | 416 | HMG-CoA synthase | JamH [AAS98779.1] | 60/79 |
| OJX77735.1 | 81 | acyl carrier protein | CalX [BAP05574.1] | 49/73 |
| OJX77736.1 | 405 | beta-ketoacyl synthase | NspH [ADA69244.1] | 45/62 |
| OJX77737.1 | 246 | enoyl-CoA hydratase | BaeH [CAG23955.1] | 48/62 |
| OJX77738.1 | 655 | asparagine synthase | Dor4 [ACY01389.1] | 56/70 |
| OJX77739.1 | 83 | acyl carrier protein | SmdG [CCC21121.1] | 54/71 |
| OJX77740.1 | 411 | hypothetical protein | / | / |
| OJX77741.1 | 289 | hypothetical protein | / | / |
| OJX77742.1 | 285 | alcohol dehydrogenase | / | / |

[a] Number of amino acids

**Table S56.** Predicted functions of ORFs in the NVUB01000125.1 containing PCI83078.1

| gene | aa [a] | putative function | Protein homologue | %identity/ %similarity |
|---|---|---|---|---|
| PCI83073.1 | 530 | phytoene dehydrogenase | / | / |
| PCI83074.1 | 284 | methyltransferase | ChiG [AAY89054.1] | 37/56 |
| PCI83075.1 | 283 | AT/Ox | DifA [CAG23974.1] | 55/75 |
| PCI83076.1 | 372 | acyltransferase | BaeD [CAG23951.1] | 45/60 |
| PCI83077.1 | 473 | 2-nitropropane dioxygenase | DifA [CAG23974.1] | 58/75 |
| PCI83078.1 | 410 | HMG-CoA synthase | CorE [ADI59527.1] | 57/73 |
| PCI83079.1 | 427 | beta-ketoacyl synthase | PedM [AAW33972.1] | 51/67 |
| PCI83080.1 | 83 | acyl carrier protein | PsyL [ADA82592.1] | 49/75 |
| PCI83081.1 | 582 | carbamoyl transferase | albXV [CAE52324.1] | 59/73 |
| ORF1 | 4300 | AT-less type I PKS | DszB [AAY32965.1] | 37/55 |
| PCI83082.1 | 1003 | AT-less type I PKS | bonA [AFN27480.1] | 42/57 |
| PCI83083.1 | 939 | AT-less type I PKS | MupK [AAM12924.1] | 46/64 |

[a] Number of amino acids

**Table S57.** Predicted functions of ORFs in the NVUB01000125.1 containing PCK07191.1

| gene | aa [a] | putative function | Protein homologue | %identity/ %similarity |
|---|---|---|---|---|
| PCK07188.1 | 138 | hypothetical protein | / | / |
| PCK07189.1 | 60 | hypothetical protein | / | / |
| PCK07190.1 | 409 | beta-ketoacyl synthase | TstN [AGN11888.1] | 57/72 |
| PCK07191.1 | 421 | HMG-CoA synthase | BatC [ADD82944.1] | 70/81 |
| PCK07192.1 | 81 | acyl carrier protein | CalX [BAP05574.1] | 44/76 |
| PCK07193.1 | 366 | S-malonyltransferase | TstO [AGN11889.1] | 50/66 |
| PCK07194.1 | 1668 | AT-less type I PKS | SorH [ADN68483.1] | 35/51 |

[a] Number of amino acids

**Table S58.** Predicted functions of ORFs in the NVXL01000012.1 containing PCK09280.1

| gene | aa [a] | putative function | Protein homologue | %identity/ %similarity |
|---|---|---|---|---|
| PCK09264.1 | 6965 | NRPS | AlbI [CAE52339.1] | 29/45 |
| PCK09265.1 | 65 | hypothetical protein | / | / |
| PCK09266.1 | 260 | hypothetical protein | / | / |
| PCK09267.1 | 1430 | hypothetical protein | / | / |
| PCK09278.1 | 457 | permease | / | / |
| PCK09268.1 | 589 | hypothetical protein | / | / |
| PCK09269.1 | 2731 | AT-less type I PKS | BaeL [CAG23958.2] | 33/50 |
| PCK09270.1 | 5756 | AT-less type I PKS | OocN [AFX60336.1] | 41/56 |
| PCK09279.1 | 247 | enoyl-CoA hydratase | BaeI [CAG23956.1] | 62/78 |
| PCK09271.1 | 256 | enoyl-CoA hydratase | BaeH [CAG23955.1] | 55/69 |
| PCK09280.1 | 416 | HMG-CoA synthase | ThaK [ABC34601.1] | 52/67 |
| PCK09272.1 | 84 | acyl carrier protein | JamF [AAS98799.1] | 39/68 |
| PCK09273.1 | 732 | AT-less type I PKS | TaI [ABF89568.1] | 31/49 |
| PCK09274.1 | 363 | S-malonyltransferase | MisG [AKQ22695.1] | 52/70 |
| PCK09275.1 | 135 | glyoxalase | BryB [ABM63527.1] | 28/43 |
| PCK09276.1 | 94 | hypothetical protein | / | / |
| PCK09277.1 | 139 | hypothetical protein | / | / |

[a] Number of amino acids

**Table S59.** Predicted functions of ORFs in the NVWB01000011.1 containing PHS16293.1

| gene | aa [a] | putative function | Protein homologue | %identity/ %similarity |
|---|---|---|---|---|
| PHS16279.1 | 258 | hypothetical protein | / | / |
| PHS16280.1 | 135 | hypothetical protein | / | / |
| PHS16281.1 | 5417 | AT-less type I PKS | BasE [ERM18797.1] | 44/61 |
| PHS16282.1 | 1127 | NRPS | BaeN [CAG23960.2] | 32/57 |
| PHS16283.1 | 2204 | NRPS | SnbDE [CBW45647.1] | 30/47 |
| PHS16284.1 | 1684 | NRPS | JamO [AAS98786.1] | 37/58 |
| PHS16285.1 | 209 | hypothetical protein | / | / |
| PHS16286.1 | 437 | hypothetical protein | / | / |
| PHS16287.1 | 446 | diaminobutyrate--2-oxoglutarate transaminase | BatP [ADD82957.1] | 28/44 |
| PHS16288.1 | 126 | GntR family transcriptional regulator | / | / |
| PHS16289.1 | 834 | hypothetical protein | / | / |
| PHS16290.1 | 222 | phosphonate ABC transporter | LnmR [AAN85531.1] | 28/47 |
| PHS16291.1 | 460 | hypothetical protein | / | / |
| PHS16292.1 | 412 | beta-ketoacyl synthase | BryQ [ABM63532.1 | 50/75 |
| PHS16293.1 | 420 | HMG-CoA synthase | BatC [ADD82944.1] | 66/77 |
| PHS16294.1 | 265 | enoyl-CoA hydratase | NspJ [ADA69246.1] | 40/64 |
| PHS16295.1 | 249 | enoyl-CoA hydratase | BatE [ADD82946.1] | 58/72 |
| PHS16296.1 | 82 | acyl carrier protein | JamF [AAS98799.1] | 49/62 |
| ORF1 | 290 | S-malonyltransferase | BatJ [ADD82951.1] | 52/69 |
| PHS16297.1 | 225 | PPTase | LtmL [ACY01405.1] | 23/46 |
| PHS16298.1 | 54 | hypothetical protein | / | / |
| PHS16299.1 | 147 | hypothetical protein | / | / |
| PHS16300.1 | 919 | hypothetical protein | / | / |

[a] Number of amino acids

**Table S60.** Predicted functions of ORFs in the QHVI01000077.1 containing PYP87646.1

| gene | aa [a] | putative function | Protein homologue | %identity/ %similarity |
|---|---|---|---|---|
| PYP87639.1 | 1851 | NRPS | SgvD4 [AGN74885.1] | 38/52 |
| PYP87640.1 | 242 | hypothetical protein | / | / |
| PYP87641.1 | 133 | glyoxalase | / | / |
| PYP87642.1 | 266 | phosphosulfolactate synthase | / | / |
| PYP87643.1 | 496 | polyketide synthase | MalF [ABC35796.1] | 22/50 |
| PYP87644.1 | 403 | hypothetical protein | / | / |
| PYP87645.1 | 171 | hypothetical protein | / | / |
| PYP87646.1 | 419 | HMG-CoA synthase | BatC [ADD82944.1] | 69/82 |
| PYP87648.1 | 223 | MBL fold metallo-hydrolase | BaeB [CAG23949.2] | 44/62 |
| PYP87647.1 | 4120 | AT-less type I PKS | BaeN [CAG23960.2] | 38/56 |

[a] Number of amino acids

**Table S61.** Predicted functions of ORFs in the QHZP01000155.1 containing PYV42947.1

| gene | aa [a] | putative function | Protein homologue | %identity/ %similarity |
|---|---|---|---|---|
| PYV42944.1 | 1398 | hybrid NRPS/PKS | BaeN [CAG23960.2] | 51/67 |
| PYV42953.1 | 135 | hypothetical protein | PedH [AAS47562.1] | 74/85 |
| PYV42954.1 | 255 | hypothetical protein | TaO [ABF92489.1] | 5/73 |
| PYV42945.1 | 224 | 2-nitropropane dioxygenase | BatK [ADD82952.1] | 60/77 |
| PYV42955.1 | 286 | IS4 family transposase | / | / |
| PYV42946.1 | 68 | hypothetical protein | OocU [AFX60343.1] | 71/85 |
| PYV42947.1 | 419 | HMG-CoA synthase | BatC [ADD82944.1] | 71/82 |
| PYV42948.1 | 262 | enoyl-CoA hydratase | BaeH [CAG23955.1] | 56/70 |
| PYV42949.1 | 248 | enoyl-CoA hydratase | BatE [ADD82946.1] | 67/84 |
| PYV42950.1 | 82 | acyl carrier protein | CalX [BAP05574.1] | 56/81 |
| PYV42951.1 | 417 | beta-ketoacyl synthase | CalW [BAP05575.1] | 64/80 |
| PYV42952.1 | 387 | S-malonyltransferase | TstO [AGN11889.1] | 54/68 |

[a] Number of amino acids

**Table S62.** Predicted functions of ORFs in the QFBR01000009.1 containing RAP32035.1

| gene | aa [a] | putative function | Protein homologue | %identity/ %similarity |
|---|---|---|---|---|
| RAP32028.1 | 370 | AT-less type I PKS | BaeL [CAG23958.2] | 28/44 |
| RAP32029.1 | 4551 | AT-less type I PKS | TaO [ABF92489.1] | 31/49 |
| RAP32030.1 | 1875 | AT-less type I PKS | NspC [ADA69239.2] | 37/57 |
| RAP32031.1 | 755 | AT/Ox | DifA [CAG23974.1] | 49/68 |
| RAP32032.1 | 197 | hypothetical protein | / | / |
| RAP32033.1 | 251 | enoyl-CoA hydratase | CylH [ARU81122.1] | 59/76 |
| RAP32034.1 | 257 | enoyl-CoA hydratase | CylG [ARU81121.1] | 42/64 |
| RAP32035.1 | 420 | HMG-CoA synthase | ThaK [ABC34601.1] | 53/70 |
| RAP32036.1 | 82 | acyl carrier protein | ThaI [ABC35804.1] | 40/58 |
| RAP32037.1 | 703 | SDR family oxidoreductase | BasG [ERM18799.1] | 31/51 |
| RAP32038.1 | 449 | hypothetical protein | / | / |

[a] Number of amino acids

**Table S63.** Predicted functions of ORFs in the OMOH01000005.1 containing SPF68542.1

| gene | aa [a] | putative function | Protein homologue | %identity/ %similarity |
|---|---|---|---|---|
| SPF68531.1 | 404 | Transposase | / | / |
| SPF68532.1 | 171 | hypothetical protein | / | / |
| SPF68533.1 | 332 | acyltransferase | kirCI [CAN89640.1] | 34/49 |
| SPF68534.1 | 229 | metallo-beta-lactamase | BaeB [CAG23949.2] | 40/56 |
| SPF68535.1 | 273 | PPTase | Mis12 [AKQ22701.1] | 32/46 |
| SPF68536.1 | 367 | beta-ketoacyl synthase | CorD [ADI59526.1] | 37/50 |
| SPF68537.1 | 78 | acyl carrier protein | Macp [AAM12920.1] | 41/66 |
| SPF68538.1 | 465 | permease | / | / |
| SPF68539.1 | 420 | permease | / | / |
| SPF68540.1 | 245 | adenosinetriphosphatase | LnmR [AAN85531.1] | 35/51 |
| SPF68541.1 | 735 | AT/Ox | MmpC [AAM12912.1] | 47/63 |
| SPF68542.1 | 412 | HMG-CoA synthase | MxnE [AGS77285.1] | 56/70 |
| SPF68543.1 | 263 | enoyl-CoA hydratase | CorF [ADI59528.1] | 41/57 |
| SPF68544.1 | 3499 | AT-less type I PKS | PedH [AAS47562.1] | 32/48 |
| SPF68545.1 | 1790 | AT-less type I PKS | ElaK [AEC04357.1] | 33/50 |
| SPF68546.1 | 2389 | AT-less type I PKS | Bat2 [ADD82940.1] | 37/51 |
| SPF68547.1 | 682 | AT-less type I PKS | NspA [ADA69237.1] | 38/54 |
| SPF68548.1 | 3385 | AT-less type I PKS | DszB [AAY32965.1] | 33/47 |
| SPF68549.1 | 2664 | AT-less type I PKS | OzmN [ABS90475.1] | 43/54 |
| SPF68550.1 | 2632 | AT-less type I PKS | MisC [AKQ22699.1] | 34/49 |
| SPF68551.1 | 707 | hypothetical protein | / | / |
| SPF68552.1 | 353 | AMP-binding enzyme | MalI [ABC34723.1] | 30/45 |

[a] Number of amino acids

**Table S64.** Predicted functions of ORFs in the NZ_JH791997.1 containing WP_002106872.1

| gene[a] | aa[b] | putative function | Protein homologue | %identity/ %similarity |
|---|---|---|---|---|
| ORF1 | 2665 | AT-less type I PKS | BaeL [CAG23958.2] | 69/80 |
| WP_002106869.1 | 249 | enoyl-CoA hydratase | BatE [ADD82946.1] | 70/86 |
| WP_002106870.1 | 254 | enoyl-CoA hydratase | BaeH [CAG23956.1] | 59/74 |
| WP_002106872.1 | 417 | HMG-CoA synthase | BatC [ADD82944.1] | 71/82 |
| WP_002106873.1 | 411 | beta-ketoacyl synthase | CalW [BAP05575.1] | 61/77 |
| WP_002106874.1 | 82 | acyl carrier protein | AcpK [CAG23953.1] | 60/75 |
| WP_033716972.1 | 1086 | AT-less type I PKS | SorA [ADN68476.1] | 51/67 |
| WP_002106884.1 | 277 | class I SAM-dependent methyltransferase | Dor10 [ACY01395.1] | 29/46 |
| WP_002106885.1 | 268 | alpha/beta hydrolase | OocA [AFX60323.1] | 27/46 |
| WP_002106886.1 | 1113 | S-malonyltransferase | BasH [ERM18800.1] | 53/72 |
| WP_002106887.1 | 225 | MBL fold metallo-hydrolase | BaeB [CAG23949.2] | 56/69 |
| WP_002106888.1 | 406 | NADP/FAD-dependent oxidoreductase | Enacyloxin [ABI91473.1] | 26/41 |
| WP_002106889.1 | 269 | PPTase | Mis12 [AKQ22701.1] | 35/54 |
| WP_002106890.1 | 177 | transcription antiterminator | ElaA [AEC04347.1] | 23/50 |
| WP_002106891.1 | 270 | ndecaprenyl-diphosphate phosphatase | / | / |

[a] ORFs are proteins without protein ID provided by NCBI [b] Number of amino acids

**Table S65.** Predicted functions of ORFs in the NZ_NBTK02000001.1 containing WP_002265857.1

| gene | aa[a] | putative function | Protein homologue | %identity/ %similarity |
|---|---|---|---|---|
| WP_002265846.1 | 372 | ABC transporter permease | / | / |
| WP_002265847.1 | 310 | ABC transporter ATP-binding protein | Rhizopodin [CCA89322.1] | 28/46 |
| WP_002272162.1 | 1701 | alpha/beta fold hydrolase | CorL [ADI59534.1] | 35/54 |
| WP_082999579.1 | 2801 | AT-less type I PKS | DszB [AAY32965.1] | 31/51 |
| WP_002275894.1 | 1588 | AT-less type I PKS | MisF [AKQ22696.1] | 35/56 |
| WP_102990528.1 | 3623 | AT-less type I PKS | BryB [ABM63527.1] | 32/51 |
| WP_082999580.1 | 1228 | AT-less type I PKS | Bat2 [ADD82940.1] | 33/52 |
| WP_082999581.1 | 2110 | AT-less type I PKS | PedH [AAS47562.1] | 31/49 |
| WP_102990529.1 | 3403 | AT-less type I PKS | BryB [ABM63527.1] | 38/57 |
| WP_002272022.1 | 4212 | AT-less type I PKS | MisF [AKQ22696.1] | 35/52 |
| WP_002265856.1 | 270 | enoyl-CoA hydratase | MxnF [AGS77286.1] | 43/62 |
| WP_002265857.1 | 410 | HMG-CoA synthase | MxnE [AGS77285.1] | 58/73 |
| WP_002265858.1 | 423 | beta-ketoacyl synthase | MxnD [AGS77284.1] | 40/62 |
| WP_002265859.1 | 82 | acyl carrier protein | ElaE [AEC04351.1] | 46/70 |
| WP_002265860.1 | 751 | AT/Ox | DifA [CAG23974.1] | 57/74 |
| WP_002265861.1 | 326 | acyltransferase | BaeD [CAG23951.1] | 33/55 |
| WP_024783492.1 | 225 | MBL fold metallo-hydrolase | BaeB [CAG23949.2] | 48/65 |
| WP_002265863.1 | 233 | biosurfactants production protein BBK-1 | MupN [AAM12928.1] | 25/49 |

[a] Number of amino acids

**Table S66.** Predicted functions of ORFs in the NZ_ACXX02000007.1 containing WP_004619350.1

| gene[a] | aa[b] | putative function | Protein homologue | %identity/ %similarity |
|---|---|---|---|---|
| ORF1 | 469 | PPTase | DifL [CAG23983.1] | 31/50 |
| WP_004619350.1 | 417 | HMG-CoA synthase | BatC [ADD82944.1] | 72/84 |
| WP_004619351.1 | 254 | enoyl-CoA hydratase | BaeH [CAG23956.1] | 58/76 |
| WP_004619352.1 | 249 | enoyl-CoA hydratase | BatE [ADD82946.1] | 67/82 |
| WP_004619353.1 | 1780 | AT-less type I PKS | DifJ [CAJ57410.1] | 46/64 |
| WP_004619354.1 | 1439 | AT-less type I PKS | MisF [AKQ22696.1] | 49/66 |
| WP_004619355.1 | 411 | beta-ketoacyl synthase | CalW [BAP05575.1] | 64/79 |
| WP_004619356.1 | 82 | acyl carrier protein | AcpK [CAG23953.1] | 66/79 |
| WP_004619358.1 | 329 | tyrosine recombinase XerC | Marinomycin [BAG50450.1] | 28/61 |
| WP_004619359.1 | 317 | beta-ketoacyl synthase | Bat1 [ADD82939.1] | 26/55 |
| WP_004619360.1 | 180 | NlpC/P60 family protein | MupP [AAM12930.1] | 27/45 |
| WP_004619361.1 | 118 | DUF1634 domain-containing protein | DszB [AAY32965.1] | 36/50 |
| WP_004619362.1 | 277 | sulfite exporter TauE/SafE family | / | / |
| WP_004619363.1 | 315 | hypothetical protein | / | / |
| WP_004619364.1 | 950 | glycoside hydrolase | SgvF [AGN74878.1] | 36/50 |
| WP_051132013.1 | 944 | glycoside hydrolase | DifI [CAJ57409.1] | 30/50 |
| WP_004619366.1 | 592 | spore coat protein CotH | MmpF [AAM12934.1] | 30/50 |
| WP_004619367.1 | 345 | sulfate ABC transporter substrate-binding protein | Basiliskamide [ERM18797.1] | 36/43 |
| WP_004619368.1 | 285 | sulfate ABC transporter permease | / | / |
| WP_004619369.1 | 291 | sulfate ABC transporter permease | / | / |
| WP_004619370.1 | 357 | sulfate ABC transporter ATP-binding protein | LnmR [AAN85531.1] | 36/51 |

[a] Number of amino acids

**Table S67.** Predicted functions of ORFs in the NZ_GG657758.1 containing WP_004936929.1

| gene | aa[a] | putative function | Protein homologue | %identity/ %similarity |
|---|---|---|---|---|
| WP_040908952.1 | 215 | formylglycine-generating enzyme | / | / |
| WP_086014788.1 | 187 | TTPase | LtmL [ACY01405.1] | 59/72 |
| WP_004936894.1 | 409 | cytochrome P450 | BaeS [CAG23962.1] | 41/59 |
| WP_004936897.1 | 127 | DUF4180 domain-containing protein | / | / |
| WP_040908145.1 | 414 | polyketide synthase | BasE [ERM18797.1] | 63//78 |
| WP_084828307.1 | 1030 | AT-less type I PKS | CalG [BAP05595.1] | 45/56 |
| WP_004936916.1 | 3009 | AT-less type I PKS | Ta1 [ABF85931.1] | 42/53 |
| WP_084828308.1 | 205 | polyketide synthase | Streptimidone [ACY01402.1] | 57/66 |
| WP_040908156.1 | 426 | alpha/beta fold hydrolase | RhiF [CAL69894.1] | 41/59 |
| WP_084828391.1 | 466 | PfaD family | Streptimidone [ACY01403.1] | 54/69 |
| WP_040908158.1 | 362 | ACP-S-malonyltransferase | MisG [AKQ22695.1] | 51/66 |
| WP_004936929.1 | 419 | HMG-CoA synthase | BatC [ADD82944.1] | 68/81 |
| WP_004936932.1 | 261 | enoyl-CoA hydratase | BaeH [CAG23956.1] | 51/67 |
| WP_004936934.1 | 250 | enoyl-CoA hydratase | BatE [ADD82946.1] | 64/81 |
| WP_004936936.1 | 81 | acyl carrier protein | NspG [ADA71314.1] | 60/83 |
| WP_106429168.1 | 412 | beta-ketoacyl synthase | NspH [ADA69244.1] | 56/73 |
| WP_004936942.1 | 321 | acyltransferase | SorO [ADN68489.1] | 36/53 |
| WP_004936946.1 | 287 | short chain dehydrogenase | BatM [ADD82954.1] | 37/58 |
| WP_004936953.1 | 1297 | AT-less type I PKS | JamL [AAS98783.1] | 32/48 |
| WP_084828392.1 | 142 | FCD domain-containing protein | / | / |
| WP_004936958.1 | 427 | MFS transporter | SnbR [CBW45761.1] | 30/47 |

[a] Number of amino acids

**Table S68.** Predicted functions of ORFs in the NZ_DS544873.1 containing WP_007093437.1

| gene | aa [a] | putative function | Protein homologue | %identity/ %similarity |
|---|---|---|---|---|
| WP_007093427.1 | 863 | RND superfamily | / | / |
| WP_007093428.1 | 302 | outer membrane lipoprotein | / | / |
| WP_007093429.1 | 429 | GMC family oxidoreductase | / | / |
| WP_007093430.1 | 242 | StAR-related lipid-transfer | / | / |
| WP_007093431.1 | 3908 | AT-less type I PKS | BaeN [CAG23960.2] | 36/55 |
| WP_007093433.1 | 3274 | AT-less type I PKS | OocJ [AFX60332.1] | 43/59 |
| WP_007093434.1 | 2025 | AT-less type I PKS | SorE [ADN68480.1] | 39/57 |
| WP_007093435.1 | 2315 | AT-less type I PKS | BasE [ERM18797.1] | 39/57 |
| WP_083777282.1 | 1371 | AT-less type I PKS | DifF [CAG23977.1] | 36/54 |
| WP_007093437.1 | 419 | HMG-CoA synthase | BatC [ADD82944.1] | 70/80 |
| WP_007093438.1 | 256 | enoyl-CoA hydratase | BaeH [CAG23955.1] | 56/70 |
| WP_007093439.1 | 249 | enoyl-CoA hydratase | BatE [ADD82946.1] | 62/80 |
| WP_007093440.1 | 319 | acyltransferase | BryP [ABM63531.1] | 39/59 |
| WP_007093441.1 | 236 | MBL fold metallo-hydrolase | BaeB [CAG23949.2] | 39/62 |
| WP_007093442.1 | 81 | acyl carrier protein | PedN [AAW33973.1] | 48/75 |
| WP_007093443.1 | 425 | beta-ketoacyl synthase | PedM [AAW33972.1] | 52/72 |
| WP_007093444.1 | 372 | S-malonyltransferase | MisG [AKQ22695.1] | 54/72 |
| WP_007093445.1 | 302 | hypothetical protein | / | / |
| WP_007093446.1 | 223 | PPTase | VirK [BAF50717.1] | 26/42 |
| WP_007093447.1 | 275 | universal stress protein | / | / |
| WP_007093448.1 | 280 | universal stress protein | / | / |

[a] Number of amino acids

**Table S69.** Predicted functions of ORFs in the NZ_CP010978.1 containing WP_007960101.1

| gene | aa[a] | putative function | Protein homologue | %identity/ %similarity |
|---|---|---|---|---|
| WP_007960109.1 | 73 | hypothetical protein | RhiH [CAL69892.1] | 26/40 |
| WP_007960108.1 | 160 | organic solvent tolerance protein | / | / |
| WP_007960107.1 | 437 | lipase family protein | / | / |
| WP_007960104.1 | 381 | thioesterase | RhiF [CAL69894.1] | 45/66 |
| WP_007960103.1 | 1632 | AT-less type I PKS | BaeL [CAG23958.2] | 50/66 |
| WP_007960102.1 | 254 | enoyl-CoA hydratase | BaeH [CAG23955.1] | 61/76 |
| WP_007960101.1 | 420 | HMG-CoA synthase | BatC [ADD82944.1] | 77/86 |
| WP_007960100.1 | 409 | beta-ketoacyl synthase | CalW [BAP05575.1] | 65/81 |
| WP_007960099.1 | 82 | acyl carrier protein | NspG [ADA71314.1] | 59/77 |
| WP_052697343.1 | 2074 | AT-less type I PKS | BaeN [CAG23960.2] | 53/72 |
| WP_007957973.1 | 2510 | AT-less type I PKS | BaeN [CAG23960.2] | 50/68 |
| WP_007957975.1 | 744 | S-malonyltransferase | BaeE [CAG23952.1] | 65/79 |
| WP_007957979.1 | 525 | MFS transporter | / | / |
| WP_007957981.1 | 369 | glycosyl hydrolase | / | / |
| WP_036681384.1 | 313 | LysR family | smdB [CCC21116.1] | 27/44 |
| WP_007957985.1 | 571 | methyl-accepting chemotaxis protein | / | / |

[a] Number of amino acids

**Table S70.** Predicted functions of ORFs in the NZ_AMZN01000028.1 containing WP_009579373.1

| gene[a] | aa[b] | putative function | Protein homologue | %identity/ %similarity |
|---|---|---|---|---|
| WP_040496208.1 | 215 | hypothetical protein | / | / |
| WP_009579361.1 | 192 | antitermination factor | ElaA [AEC04347.1] | 30/52 |
| ORF1 | 7274 | AT-less type I PKS (Acp-KS-DH-KR-Acp-KS-DH-KR-Acp-Acp-KS-DH-KR-Acp-Acp-KS-DH-Acp-TE) | Bat3 [ADD82941.1] | 38/56 |
| WP_040496206.1 | 303 | NAD(P)-dependent alcohol dehydrogenase | TaP [ABF88102.1] | 28/44 |
| WP_009579365.1 | 429 | GMC family oxidoreductase | / | / |
| WP_009579366.1 | 261 | hypothetical protein | / | / |
| WP_040496208.1 | 215 | hypothetical protein | / | / |
| WP_009579368.1 | 871 | NRPS (A-PCP) | CalI [BAP05597.1] | 41/57 |
| WP_009579370.1 | 5613 | AT-less type I PKS | BaeN [CAG23960.2] | 39/58 |
| WP_009579371.1 | 2059 | AT-less type I PKS | SorE [ADN68480.1] | 42/59 |
| WP_009579372.1 | 2433 | AT-less type I PKS | MisF [AKQ22696.1] | 36/54 |
| WP_009579373.1 | 419 | HMG-CoA synthase | BatC [ADD82944.1] | 70/82 |
| WP_009579374.1 | 256 | enoyl-CoA hydratase | BaeH [CAG23956.1] | 59/70 |
| WP_009579375.1 | 248 | enoyl-CoA hydratase | BatE [ADD82946.1] | 65/80 |
| WP_009579376.1 | 228 | DUF4386 domain-containing protein | / | / |

[a] ORFs are proteins without protein ID provided by NCBI [b] Number of amino acids

**Table S71.** Predicted functions of ORFs in the NZ_AEWH01000057.1 containing WP_010097999.1

| gene[a] | aa[b] | putative function | Protein homologue | %identity/ %similarity |
|---|---|---|---|---|
| ORF1 | 1583 | AT-less type I PKS (ACP-KS-DH-KR) | DifI [CAJ57409.1] | 43/62 |
| WP_010097990.1 | 965 | AT-less type I PKS (KS-DH) | SorE [ADN68480.1] | 49/65 |
| WP_010097991.1 | 1625 | AT-less type I PKS (ACP-KR-ACP-KS-ACP) | Misakinolide [AKQ22698.1] | 44/62 |
| WP_029191154.1 | 241 | hypothetical protein | elaK [AEC04357.1] | 34/55 |
| WP_029191155.1 | 1280 | AT-less type I PKS (KS-DH) | MisC [AKQ22699.1] | 37/55 |
| WP_081472478.1 | 919 | AT-less type I PKS (KR-ACP-KS) | BaeL [CAG23958.2] | 51/68 |
| WP_029191157.1 | 3716 | AT-less type I PKS (HD-KR-ACP-KS-DH-KR-ACP-KS-DH) | BaeN [CAG23960.2] | 37/56 |
| WP_029191158.1 | 160 | SDR family oxidoreductase | BasE [ERM18797.1] | 47/76 |
| WP_050801683.1 | 1936 | AT-less type I PKS (ACP-KS-ACP-ACP-KS-ACP) | Thailandamide [ABC34675.1] | 38/53 |
| WP_081472479.1 | 233 | PPTase | MupN [AAM12928.1] | 32/55 |
| WP_010097999.1 | 421 | HMG-CoA synthase | OnnA [AAV97869.1] | 70/83 |
| WP_050801684.1 | 213 | transcription factor FapR | / | / |

[a] ORFs are proteins without protein ID provided by NCBI [b] Number of amino acids

**Table S72.** Predicted functions of ORFs in the NC_010162.1 containing WP_012235812.1

| gene | aa[a] | putative function | Protein homologue | %identity/ %similarity |
|---|---|---|---|---|
| WP_012235804.1 | 251 | S1 RNA-binding domain | / | / |
| WP_044965056.1 | 139 | DUF3037 domain | KirHV [CAN89652.1] | 48/60 |
| WP_012235806.1 | 262 | hypothetical protein | KirHIV [CAN89651.1] | 44/62 |
| WP_080598891.1 | 469 | amidase | SorP [ADN68490.1] | 56/68 |
| WP_012235808.1 | 121 | DUF4180 domain | / | / |
| WP_012235809.1 | 1111 | ATP-binding protein | SorM [ADN68497.1] | 53/66 |
| WP_012235810.1 | 249 | enoyl-CoA hydratase | BonI [AFN27485.1] | 68/83 |
| WP_012235811.1 | 258 | enoyl-CoA hydratase | BatD [ADD82945.1] | 60/77 |
| WP_012235812.1 | 420 | HMG-CoA synthase | BonG [AFN27479.1] | 71/81 |
| WP_012235813.1 | 410 | beta-ketoacyl synthase | BatB [ADD82943.1] | 66/77 |
| WP_012235814.1 | 82 | acyl carrier protein | BatA [ADD82942.1] | 61/74 |
| WP_012235815.1 | 459 | 2-nitropropane dioxygenase | BatK [ADD82952.1] | 68/83 |
| WP_044965058.1 | 68 | hypothetical protein | / | / |
| WP_012235816.1 | 104 | acyl carrier protein | Streptimidone [ACY01398.1] | 40/65 |
| WP_012235817.1 | 305 | ACP S-malonyltransferase | ThaF [ABC34740.1] | 35/50 |
| WP_012235818.1 | 654 | asparagine synthase | SorQ [ADN68491.1] | 52/68 |
| WP_080599263.1 | 380 | SDR family oxidoreductase | Bat2 [ADD82940.1] | 47/64 |
| WP_086018304.1 | 491 | SDR family oxidoreductase | SorA [ADN68476.1] | 53/67 |
| WP_080598892.1 | 3679 | AT-less type I PKS | BasE [ERM18797.1] | 47/63 |
| WP_012235823.1 | 3578 | AT-less type I PKS | MisF [AKQ22696.1] | 35/51 |
| WP_080599265.1 | 374 | alpha/beta hydrolase | ElaR [AEC04364.1] | 30/48 |
| WP_012235804.1 | 251 | S1 RNA-binding domain | / | / |

[a] Number of amino acids

**Table S73.** Predicted functions of ORFs in the NZ_LN681227.1 containing WP_013184044.1

| gene | aa[a] | putative function | Protein homologue | %identity/ %similarity |
|---|---|---|---|---|
| WP_013184029.1 | 69 | hypothetical protein | / | / |
| WP_013184032.1 | 529 | ABC transporter ATP-binding protein | Leinamycin [AAN85547.1] | 31/50 |
| WP_013184033.1 | 2344 | AT-less type I PKS | SgvE2 [AGN74893.1] | 37/50 |
| WP_081480219.1 | 674 | NRPS | SgvE1 [AGN74892.1] | 37/50 |
| WP_013184035.1 | 664 | alpha-keto acid dehydrogenase | SnaF [CBW45750.1] | 49/64 |
| WP_081480176.1 | 232 | thioesterase | VirJ [BAF50718.1] | 43/58 |
| WP_013184037.1 | 400 | MFS transporter | Kirromycin [CAN89618.1] | 26/46 |
| WP_013184038.1 | 286 | S-malonyltransferase | snaM [CBW45739.1] | 48/66 |
| WP_013184039.1 | 2302 | NRPS | SnaD [CBW45640.1] | 27/44 |
| WP_041573665.1 | 2407 | NRPS | SgvE4 [AGN74895.1] | 33/47 |
| WP_013184041.1 | 1980 | AT-less type I PKS | CorL [ADI59534.1] | 34/50 |
| WP_013184042.1 | 250 | enoyl-CoA hydratase | VirE [BAF50723.1] | 50/61 |
| WP_013184043.1 | 262 | enoyl-CoA hydratase | SnaJ [CBW45744.1] | 33/47 |
| WP_013184044.1 | 411 | HMG-CoA synthase | SnaI [CBW45745.1] | 59/72 |
| WP_013184045.1 | 408 | beta-ketoacyl synthase | virB [BAF50726.1] | 42/61 |
| WP_013184046.1 | 81 | acyl carrier protein | DipF [AGS06833.1] | 35/58 |
| WP_013183963.1 | 339 | IS110 family transposase | / | / |

[a] Number of amino acids

**Table S74.** Predicted functions of ORFs in the NC_017093.1 containing WP_014846099.1

| gene | aa[a] | putative function | Protein homologue | %identity/ %similarity |
|---|---|---|---|---|
| WP_014846089.1 | 279 | MerR family transcriptional regulator | / | / |
| WP_014846090.1 | 83 | acyl carrier protein | AcpK [CAG23953.1] | 42/61 |
| WP_041696284.1 | 330 | AT/Ox | MmpC [AAM12912.1] | 37/51 |
| WP_014846092.1 | 419 | beta-ketoacyl synthase | DipR [AGS06821.1] | 26/52 |
| WP_014846093.1 | 240 | MBL fold metallo-hydrolase | BaeB [CAG23949.2] | 41/58 |
| WP_014846094.1 | 372 | O-methyltransferase | AlbII [CAE52340.1] | 27/43 |
| WP_014846095.1 | 769 | AT/Ox | DifA [CAG23974.1] | 49/66 |
| WP_051014945.1 | 278 | hypothetical protein | BatD [ADD82945.1] | 45/62 |
| WP_081490236.1 | 999 | AT-less type I PKS | ElaJ [AEC04356.1] | 39/52 |
| WP_041696287.1 | 2731 | AT-less type I PKS | BryB [ABM63527.1] | 36/54 |
| WP_041696289.1 | 5084 | AT-less type I PKS | LglD [AIU36100.1] | 33/48 |
| WP_081490237.1 | 3741 | AT-less type I PKS | DszB [AAY32965.1] | 36/50 |
| WP_081490238.1 | 1774 | AT-less type I PKS | ThaO [ABC34675.1] | 41/52 |
| WP_014846098.1 | 1536 | AT-less type I PKS | PsyD [ADA82585.1] | 35/50 |
| WP_014846099.1 | 420 | HMG-CoA synthase | BaeG [CAG23954.2] | 64/77 |
| WP_081490381.1 | 253 | enoyl-CoA hydratase | BaeE [CAG23952.1] | 56/68 |
| WP_014846101.1 | 246 | 3-oxoacyl-ACP reductase | DifE [CAG23976.1] | 36/57 |
| WP_081490382.1 | 248 | TTPase | LtmL [ACY01405.1] | 47/60 |
| WP_014846102.1 | 310 | ATP-binding cassette | Rhizopodin [CCA89322.1] | 23/50 |
| WP_041696291.1 | 227 | ABC transporter permease | / | / |

[a] Number of amino acids

**Table S75.** Predicted functions of ORFs in the NZ_LGJF01000001.1 containing WP_016742406.1

| gene | aa[a] | putative function | Protein homologue | %identity/ %similarity |
|---|---|---|---|---|
| WP_016742397.1 | 176 | transcription antiterminator | / | / |
| WP_081603457.1 | 266 | PPTase | Mis12 [AKQ22701.1] | 33/55 |
| WP_016742399.1 | 230 | MBL fold metallo-hydrolase | BaeB [CAG23949.2] | 49/64 |
| WP_016742400.1 | 1097 | malonyl CoA-ACP transacylase | BasH [ERM18800.1] | 48/69 |
| WP_016742401.1 | 1519 | AT-less type I PKS | BaeN [CAG23960.2] | 45/60 |
| WP_081603458.1 | 3259 | AT-less type I PKS | BryB [ABM63527.1] | 42/59 |
| WP_020372034.1 | 2632 | AT-less type I PKS | BryB [ABM63527.1] | 40/56 |
| WP_016742403.1 | 1841 | AT-less type I PKS | OnnI [AAV97877.1] | 46/61 |
| WP_016742405.1 | 412 | beta-ketoacyl synthase | CalW [BAP05575.1] | 42/69 |
| WP_016742406.1 | 418 | HMG-CoA synthase | BaeG [CAG23954.2] | 69/80 |
| WP_016742407.1 | 254 | enoyl-CoA hydratase | BaeH [CAG23955.1] | 53/70 |
| WP_016742408.1 | 248 | enoyl-CoA hydratase | ElaN [AEC04360.1] | 68/80 |
| WP_049684135.1 | 1949 | AT-less type I PKS | TaO [ABF92489.1] | 51/68 |
| WP_081603459.1 | 3467 | AT-less type I PKS | SorA [ADN68476.1] | 47/62 |
| WP_016742410.1 | 4771 | AT-less type I PKS | MisF [AKQ22696.1] | 39/56 |
| WP_016742411.1 | 70 | hypothetical protein | / | / |
| WP_016742412.1 | 306 | RluA family pseudouridine synthase | / | / |

[a] Number of amino acids

**Table S76.** Predicted functions of ORFs in the NZ_JH806633.1 containing WP_017178648.1

| gene | aa[a] | putative function | Protein homologue | %identity/ %similarity |
|---|---|---|---|---|
| WP_017178642.1 | 288 | polyketide synthase | / | / |
| WP_051000626.1 | 234 | PPTase | LtmL [ACY01405.1] | 55/65 |
| WP_017178644.1 | 281 | Ser/Thr protein phosphatase | ThaC [ABC35295.1] | 31/41 |
| WP_017178645.1 | 238 | oxidoreductase | DifE [CAG23976.1] | 35/57 |
| WP_017178646.1 | 82 | acyl carrier protein | ThaI [ABC35804.1] | 41/58 |
| WP_017178647.1 | 253 | enoyl-CoA hydratase | BaeE [CAG23952.1] | 52/67 |
| WP_017178648.1 | 416 | HMG-CoA synthase | ThaK [ABC34601.1] | 64/79 |
| WP_017178649.1 | 1517 | AT-less type I PKS | JamL [AAS98783.1] | 40/56 |
| WP_017178650.1 | 2634 | AT-less type I PKS | LglD [AIU36100.1] | 34/49 |
| WP_020372764.1 | 3690 | AT-less type I PKS | SorE [ADN68480.1] | 36/49 |
| WP_082181219.1 | 179 | polyketide synthase | Ta1 [ABF85931.1] | 37/57 |
| WP_082181191.1 | 244 | polyketide synthase | ElaQ [AEC04363.1] | 36/48 |
| WP_043508391.1 | 1381 | AT-less type I PKS | SorA [ADN68476.1] | 34/50 |
| WP_017178652.1 | 215 | BC-2 transporter permease | / | / |
| WP_017178653.1 | 291 | ABC transporter ATP-binding protein | Rhizopodin [CCA89322.1] | 31/47 |
| WP_082181192.1 | 962 | AT-less type I PKS | ElaJ [AEC04356.1] | 35/53 |
| WP_082181193.1 | 282 | enoyl-CoA hydratase | ThaL [ABC35267.1] | 45/57 |
| WP_017178656.1 | 730 | AT/Ox | DifA [CAG23974.1] | 44/62 |
| WP_017178657.1 | 410 | beta-ketoacyl synthase | JamG [AAS98778.1] | 32/50 |
| WP_017178658.1 | 313 | S-malonyltransferase | MxnM [AGS77293.1] | 34/48 |
| WP_082181194.1 | 251 | DNA-binding response regulator | smdD [CCC21118.1] | 35/50 |
| WP_017178660.1 | 333 | ATP-binding cassette domain | Rhizopodin [CCA89322.1] | 31/45 |
| WP_017178661.1 | 259 | ABC transporter | / | / |
| WP_017178662.1 | 311 | sensor histidine kinase | smdC [CCC21117.1] | 33/47 |

[a] Number of amino acids

**Table S77.** Predicted functions of ORFs in the NZ_AUUC01000043.1 containing WP_018596752.1

| protein_id | aa[a] | putative function | protein homologue | %identity/ %similarity |
|---|---|---|---|---|
| WP_018596747.1 | 120 | IS66 family insertion sequence | CalC [BAP05591.1] | 34/46 |
| WP_018596749.1 | 755 | ACP-S-malonyltransferase | BaeE [CAG23952.1] | 41/63 |
| WP_026255616.1 | 84 | acyl carrier protein | OocG [AFX60329.1] | 41/61 |
| WP_018596751.1 | 408 | beta-ketoacyl synthase | DipR [AGS06821.1] | 38/59 |
| WP_018596752.1 | 410 | HMG-CoA synthase | MxnE [AGS77285.1] | 54/70 |
| WP_018596753.1 | 270 | enoyl-CoA hydratase | CorF [ADI59528.1] | 37/58 |
| WP_018596754.1 | 3339 | AT-less type I PKS | CorK [ADI59533.1] | 29/49 |
| WP_081624831.1 | 908 | AT-less type I PKS | DifL [CAG23983.1] | 44/61 |
| WP_018596757.1 | 1750 | AT-less type I PKS | OzmN [ABS90475.1] | 33/49 |
| WP_081624832.1 | 2495 | AT-less type I PKS | LglD [AIU36100.1] | 37/55 |

[a] Number of amino acids

**Table S78.** Predicted functions of ORFs in the NC_021658.1 containing WP_020732789.1

| gene[a] | aa[b] | putative function | Protein homologue | %identity/ %similarity |
|---|---|---|---|---|
| WP_044987500.1 | 476 | F0F1 ATP synthase subunit beta | / | / |
| ORF1 | 4901 | AT-less type I PKS (KS-DH-KR-MT-Acp-ER-KS-PS-KR-Acp-KS) | ElaK [AEC04357.1] | 39/54 |
| ORF2 | 5997 | AT-less type I PKS (Acp-KS-DH-MT-Acp-KS-KR-Acp-KS-ECH-Acp -KS-DH-KR-MT-Acp) | BasE [ERM18797.1] | 35/52 |
| ORF3 | 6735 | AT-less type I PKS (KS-KS-DH-Acp-KS-DH-KR-Acp-KS-KR-Acp-KS-oMT-Acp-KS-KR-MT-Acp) | MisC [AKQ22699.1] | 37/53 |
| ORF4 | 4430 | AT-less type I PKS (KS-KR-Acp-KS-PS-KR-Acp-KS-KR-Acp-TE) | DszB [AAY32965.1] | 39/52 |
| WP_080681951.1 | 1007 | Ser/Thr kinase | Rhizopodin [CCA89332.1] | 37/58 |
| WP_080681952.1 | 478 | 2-polyprenyl-6-methoxyphenol hydroxylase | / | / |
| WP_020732780.1 | 120 | hypothetical protein | SnaE1 [CBW45749.1] | 37/58 |
| WP_020732781.1 | 933 | ACP-S-malonyltransferase | OocV [AFX60344.1] | 40/58 |
| WP_020732782.1 | 84 | hypothetical protein | BonL [AFN27475.1] | 25/45 |
| WP_020732783.1 | 59 | hypothetical protein | BaeJ [CAG23957.2] | 45/54 |
| WP_020732784.1 | 339 | NADP-dependent oxidoreductase | Dor9 [ACY01394.1] | 41/55 |
| WP_080681953.1 | 146 | acyl carrier protein | PsyL [ADA82592.1] | 43/69 |
| WP_020732786.1 | 269 | enoyl-CoA hydratase | MxnF [AGS77286.1] | 45/59 |
| WP_080681954.1 | 331 | hypothetical protein | KirAI [CAN89631.1] | 34/46 |
| WP_020732788.1 | 393 | enoyl-CoA hydratase | BonI [AFN27485.1] | 43/58 |
| WP_020732789.1 | 413 | HMG-CoA synthase | MxnE [AGS77285.1] | 54/69 |
| WP_044985488.1 | 133 | lysine cyclodeaminase | PipA [CBW45757.1] | 38/56 |
| WP_049949408.1 | 125 | transposase | SorR [ADN68492.1] | 41/54 |
| WP_020732790.1 | 199 | hypothetical protein | RizD [CCA89328.1] | 35/45 |
| WP_020732791.1 | 427 | MFS transporter | Leinamycin [AAN85500.1] | 27/45 |
| WP_020732793.1 | 305 | glutamyl-Q tRNA (Asp) synthetase | SnbC [CBW45648.1] | 32/44 |
| WP_020732794.1 | 599 | glutamate--tRNA ligase | ChiB [AAY89049.1] | 29/42 |

[a] ORFs are proteins without protein ID provided by NCBI [b] Number of amino acids

**Table S79.** Predicted functions of ORFs in the NZ_CP011966.2 containing WP_023973751.1 and WP_023973754.1

| gene | aa[a] | putative function | Protein homologue | %identity/ %similarity |
|---|---|---|---|---|
| WP_031275628.1 | 290 | IS91 family transposase | / | / |
| WP_054249082.1 | 520 | benzoate-CoA ligase | AlbVII [CAE52336.1] | 26/46 |
| WP_023973737.1 | 80 | hypothetical protein | BryB [ABM63527.1] | 27/49 |
| WP_023973738.1 | 250 | lpha/beta hydrolase | / | / |
| WP_023973739.1 | 235 | PPTase | MupN [AAM12928.1] | 28/50 |
| WP_023973740.1 | 313 | alpha/beta hydrolase | Oxazolomycin [ABS90461.1] | 32/45 |
| WP_054249081.1 | 322 | hypothetical protein | Bat2 [ADD82940.1] | 33/54 |
| WP_054249080.1 | 687 | AT-less type I PKS | ChxE [AFO59866.1] | 46/65 |
| WP_023973742.1 | 81 | acyl carrier protein | SmdG [CCC21121.1] | 49/75 |
| WP_023973743.1 | 256 | class I SAM-dependent methyltransferase | OnnD [AAV97872.1] | 26/45 |
| WP_023973744.1 | 470 | AMP-dependent synthetase | Kirromycin [CAN89663.1] | 21/38 |
| WP_023973745.1 | 235 | beta-ketoacyl synthase | BatB [ADD82943.1] | 57/77 |
| WP_054249079.1 | 1544 | canonical type I PKS | PsyD [ADA82585.1] | 34/54 |
| WP_054249078.1 | 1820 | AT-less type I PKS | LglD [AIU36100.1] | 41/57 |
| WP_023973750.1 | 249 | enoyl-CoA hydratase | BatE [ADD82946.1] | 65/83 |
| WP_023973751.1 | 419 | HMG-CoA synthase | BatC [ADD82944.1] | 62/80 |
| WP_023973752.1 | 84 | ACP | AcpK [CAG23953.1] | 49/77 |
| WP_023973753.1 | 254 | enoyl-CoA hydratase | BaeH [CAG23956.1] | 47/70 |
| WP_023973754.1 | 419 | HMG-CoA synthase | BatC [ADD82944.1] | 76/87 |
| WP_031275632.1 | 412 | beta-ketoacyl synthase | / | / |
| WP_023973756.1 | 711 | malonyl CoA-ACP transacylase | BasH [ERM18800.1] | 46/66 |
| WP_023973757.1 | 239 | MBL fold metallo-hydrolase | BaeB [CAG23949.2] | 54/71 |
| WP_023973758.1 | 170 | transcription antiterminator | ElaA [AEC04347.1] | 26/49 |

[a] Number of amino acids

**Table S80.** Predicted functions of ORFs in the NZ_JAGE01000002.1 containing WP_024834393.1

| gene | aaa | putative function | Protein homologue | %identity/ %similarity |
|---|---|---|---|---|
| WP_024834386.1 | 487 | hypothetical protein | / | / |
| WP_024834387.1 | 317 | hypothetical protein | / | / |
| WP_024834388.1 | 772 | AT/Ox | DifA [CAG23974.1] | 61/77 |
| WP_024834389.1 | 269 | alpha/beta hydrolase | / | / |
| WP_024834390.1 | 273 | methyltransferase | TaQ [ABF89350.1] | 23/44 |
| WP_024834391.1 | 3515 | AT-less type I PKS | BaeN [CAG23960.2] | 41/60 |
| WP_024834392.1 | 4235 | AT-less type I PKS | BryB [ABM63527.1] | 47/64 |
| WP_024834393.1 | 420 | HMG-CoA synthase | BatC [ADD82944.1] | 74/84 |
| WP_024834394.1 | 249 | enoyl-CoA hydratase | BatE [ADD82946.1] | 69/84 |
| WP_024834395.1 | 2954 | AT-less type I PKS | Bat2 [ADD82940.1] | 50/65 |
| WP_024834396.1 | 2368 | AT-less type I PKS | SorB [ADN68477.1] | 42/58 |
| WP_024834397.1 | 1617 | AT-less type I PKS | NspC [ADA69239.2] | 40/58 |
| WP_024834398.1 | 329 | aldo/keto reductase | / | / |
| WP_024834399.1 | 234 | HAD family phosphatase | SgvC [AGN74907.1] | 41/60 |
| WP_024834400.1 | 223 | hypothetical protein | / | / |
| WP_024834401.1 | 463 | MATE family efflux transporter [ | SorJ [ADN68485.1] | 23/46 |

[a] Number of amino acids

**Table S81.** Predicted functions of ORFs in the FR902444.1 containing WP_024834510.1

| gene | aa [a] | putative function | Protein homologue | %identity/ %similarity |
|---|---|---|---|---|
| WP_024834502.1 | 252 | sugar-binding protein | / | / |
| WP_024834503.1 | 63 | hypothetical protein | / | / |
| WP_024834504.1 | 82 | acyl carrier protein | OocG [AFX60329.1] | 51/68 |
| WP_024834505.1 | 424 | hypothetical protein | PedM [AAW33972.1] | 51/68 |
| WP_024834506.1 | 784 | AT/Ox | DifA [CAG23974.1] | 58/75 |
| WP_024834507.1 | 3405 | AT-less type I PKS | ElaP [AEC04362.1] | 40/59 |
| WP_024834508.1 | 3626 | AT-less type I PKS | BaeL [CAG23958.2] | 49/66 |
| WP_024834509.1 | 2904 | AT-less type I PKS | BaeL [CAG23958.2] | 44/62 |
| WP_024834510.1 | 419 | HMG-CoA synthase | BaeG [CAG23954.2] | 78//85 |
| WP_034848345.1 | 249 | enoyl-CoA hydratase | BaeH [CAG23955.1] | 56/73 |
| WP_051461068.1 | 2981 | AT-less type I PKS | TaO [ABF92489.1] | 48/66 |
| WP_024834511.1 | 1251 | hypothetical protein | / | / |
| WP_081741830.1 | 413 | methyltransferase | DszA [AAY32964.1] | 38/58 |
| WP_024834513.1 | 182 | DUF3795 domain-containing protein | / | / |
| WP_024834514.1 | 196 | cysteine hydrolase | leinamycin [AAN85507.1] | 45/62 |
| WP_024834515.1 | 336 | hydrogenase | leinamycin [AAN85508.1] | 60/77 |
| WP_024834516.1 | 770 | carbamoyltransferase | leinamycin [AAN85509.1] | 48/67 |
| WP_024834517.1 | 249 | enoyl-CoA hydratase | BatE [ADD82946.1] | 62/78 |
| WP_024834518.1 | 345 | DUF362 domain-containing protein [ | / | / |
| WP_024834519.1 | 300 | permease | / | / |

[a] Number of amino acids

**Table S82.** Predicted functions of ORFs in the NZ_ASSC01000679.1 containing WP_025701966.1

| gene[a] | aa[b] | putative function | Protein homologue | %identity/ %similarity |
|---|---|---|---|---|
| ORF1 | 3145 | AT-less type I PKS (MT-Acp-KS-DH-KR-Acp-KS-KR-Acp) | DifI [CAJ57409.1] | 48/63 |
| WP_025701964.1 | 1939 | AT-less type I PKS (ER-KS-DH-KR-Acp) | Myxovirescin [ABF92489.1] | 54/70 |
| WP_084463298.1 | 506 | enoyl-CoA hydratase | BatE [ADD82946.1] | 68/83 |
| WP_025701966.1 | 419 | HMG-CoA synthase | BaeG [CAG23954.2] | 74/83 |
| WP_025701967.1 | 82 | acyl carrier protein | NspG [ADA71314.1] | 59/79 |
| ORF2 | 936 | AT-less type I PKS (KS-Acp-Acp) | ElaK [AEC04357.1] | 47/60 |

[a] ORFs are proteins without protein ID provided by NCBI [b] Number of amino acids

**Table S83.** Predicted functions of ORFs in the NZ_BARF01000050.1 containing WP_078622965.1 and WP_026150782.1

| gene | aa[a] | putative function | Protein homologue | %identity/ %similarity |
|---|---|---|---|---|
| WP_026150777.1 | 409 | cytochrome P450 | Dor11 [ACY01396.1] | 60/67 |
| WP_026150779.1 | 250 | thioesterase | LnmN [AAN85527.1] | 45/56 |
| WP_078622964.1 | 964 | hypothetical protein | Streptimidone [ACY01400.1] | 51/63 |
| WP_019059253.1 | 85 | acyl carrier protein | SmdG [CCC21121.1] | 39/67 |
| WP_019059254.1 | 265 | SAM-dependent methyltransferase | Dor10 [ACY01395.1] | 29/35 |
| WP_019059255.1 | 479 | hypothetical protein | ChiD [AAY89051.1] | 27/41 |
| WP_052347028.1 | 1541 | AT-less type I PKS | PsyD [ADA82585.1] | 38/53 |
| WP_019059257.1 | 1843 | AT-less type I PKS | RizB [CCA89326.1] | 45/58 |
| WP_052347029.1 | 2564 | AT-less type I PKS | BasE [ERM18797.1] | 40/63 |
| WP_019059258.1 | 2522 | AT-less type I PKS | ElaP [AEC04362.1] | 36/53 |
| WP_019059259.1 | 249 | malonyl-CoA-transacylase | BaeE [CAG23952.1] | 64/79 |
| WP_026150780.1 | 254 | enoyl-CoA hydratase | BatD [ADD82945.1] | 56/70 |
| WP_078622965.1 | 420 | HMG-CoA synthase | BaeG [CAG23954.2] | 65/79 |
| WP_078622966.1 | 413 | beta-ketoacyl synthase | BatB [ADD82943.1] | 60/74 |
| WP_019059263.1 | 85 | acyl carrier protein | acpK [CAG23953.1] | 65/79 |
| WP_026150782.1 | 419 | HMG-CoA synthase | BaeG [CAG23954.2] | 71/81 |
| WP_026150783.1 | 286 | S-malonyltransferase | BaeC [CAG23950.2] | 50/68 |
| WP_019059266.1 | 404 | cytochrome P450 | ElaG [AEC04353.1] | 36/55 |

[a] Number of amino acids

**Table S84.** Predicted functions of ORFs in the NZ_AUCH01000004.1 containing WP_026686156.1

| gene[a] | aa[b] | putative function | Protein homologue | %identity/ %similarity |
|---|---|---|---|---|
| WP_051281936.1 | 305 | ADP-ribosylglycohydrolase | / | / |
| WP_026686150.1 | 292 | alpha/beta fold hydrolase | PtzD [AHC73995.1] | 30/45 |
| WP_051281938.1 | 284 | PPTase | BatI [ADD82950.1] | 27/44 |
| WP_026686151.1 | 779 | AT/Ox | CalY [BAP05573.1] | 35/69 |
| ORF1 | 7010 | AT-less type I PKS | Nosperin [ADA69237.1] | 37/53 |
| WP_026686153.1 | 433 | monooxygenase | PedG [AAS47561.1] | 61/77 |
| ORF2 | 5978 | AT-less type I PKS | Ta1 [ABF85931.1] | 39/53 |
| WP_084513605.1 | 294 | alpha/beta fold hydrolase | PedH [AAS47562.1] | 38/50 |
| WP_026686155.1 | 268 | FkbM family methyltransferase | CalA [BAP05589.1] | 38/51 |
| WP_051281940.1 | 413 | cytochrome P450 | ElaG [AEC04353.1] | 35/50 |
| WP_026686156.1 | 421 | HMG-CoA synthase | JamH [AAS98779.1] | 61/78 |
| WP_026686157.1 | 86 | acyl carrier protein | CalX [BAP05574.1] | 41/70 |
| WP_051281942.1 | 399 | beta-ketoacyl synthase | NspH [ADA69244.1] | 47/63 |
| WP_084513585.1 | 180 | DUF3916 domain | / | / |

[a] ORFs are proteins without protein ID provided by NCBI [b] Number of amino acids

**Table S85.** Predicted functions of ORFs in the NZ_AUKW01000003.1 containing WP_028430629.1

| gene | aa[a] | putative function | Protein homologue | %identity/ %similarity |
|---|---|---|---|---|
| WP_078505134.1 | 436 | Aminotransferase | SgvL [AGN74881.1] | 65/72 |
| WP_078505136.1 | 567 | 2,3-dihydroxybenzoate-AMP ligase | SnbA [CBW45758.1] | 67/75 |
| WP_051264698.1 | 362 | lysine cyclodeaminase | PipA [CBW45757.1] | 69/76 |
| WP_051264787.1 | 386 | cytochrome P450 | SnbF [CBW45756.1] | 67/81 |
| WP_078505219.1 | 636 | alpha-keto acid dehydrogenase | SnaF [CBW45750.1] | 75/83 |
| WP_028430626.1 | 5068 | NRPS | VirA [BAF50727.1] | 61/69 |
| WP_028430627.1 | 2467 | AT-less type I PKS | VirA [BAF50727.1] | 64/71 |
| WP_028430628.1 | 79 | acyl carrier protein | SnaG [CBW45747.1] | 63/75 |
| WP_078505138.1 | 460 | beta-ketoacyl synthase | VirB [BAF50726.1] | 73/79 |
| WP_028430629.1 | 417 | HMG-CoA synthase | VirC [BAF50725.1] | 86/90 |
| WP_051264700.1 | 257 | enoyl-CoA hydratase | VirD [BAF50724.1] | 70/74 |
| WP_051264788.1 | 248 | enoyl-CoA hydratase | VirE [BAF50723.1] | 77/85 |
| WP_028430630.1 | 2066 | AT-less type I PKS | SnaE3 [CBW45741.1] | 60/68 |
| WP_028430631.1 | 2664 | NRPS | VirH [BAF50720.1] | 64/70 |
| WP_051264702.1 | 295 | S-malonyltransferase | VirI [BAF50719.1] | 75/82 |
| WP_078505140.1 | 271 | thioesterase | VirJ [BAF50718.1] | 67/73 |
| WP_078505141.1 | 305 | PPTase | VirK [BAF50717.1] | 59/69 |
| WP_051264704.1 | 565 | ABC transporter ATP-binding protein | VirL [BAF50716.1] | 69/78 |
| WP_078505143.1 | 350 | regulatory protein | VmsS [BAF50715.1] | 64/72 |
| WP_028430633.1 | 385 | N-methyl-L-tryptophan oxidase | VirM [BAF50714.1] | 71/81 |
| WP_051264706.1 | 368 | LLM class flavin-dependent oxidoreductase | VirN [BAF50713.1] | 77/87 |
| WP_078505145.1 | 114 | acyl carrier protein | SnaX [CBW45732.1] | 46/67 |
| WP_051264710.1 | 280 | helix-turn-helix transcriptional regulator | VmsT [BAF50712.1] | 47/55 |
| WP_028430634.1 | 2647 | NRPS | VisE [BAF50711.1] | 66/73 |
| WP_028430635.1 | 2382 | NRPS | SnbDE [CBW45647.1] | 57/67 |
| WP_051264712.1 | 505 | enoyl-CoA hydratase | PglA [CBW45646.1] | 64/72 |
| WP_028430636.1 | 361 | pyruvate dehydrogenase | PglB [CBW45645.1] | 69/78 |
| WP_051264792.1 | 323 | lpha-ketoacid dehydrogenase | PglC [CBW45644.1] | 83/86 |
| WP_028430638.1 | 71 | protein mbtH | MbtY [CBW45642.1] | 72/84 |
| WP_078505150.1 | 465 | PLP-dependent aminotransferase | PglE [CBW45641.1] | 72/81 |
| WP_028430640.1 | 2473 | NRPS | PglE [CBW45641.1] | 65/72 |
| WP_037902734.1 | 436 | FMN-dependent monooxygenase | SnaA [CBW45639.1] | 77/85 |
| WP_051264714.1 | 311 | LLM class flavin-dependent oxidoreductase | SnaB [CBW45637.1] | 49/58 |
| WP_078505152.1 | 269 | NAD [P]H-dependent oxidoreductase | / | / |

[a] Number of amino acids

**Table S86.** Predicted functions of ORFs in the NZ_KE384307.1 containing WP_028545681.1

| gene | aa[a] | putative function | Protein homologue | %identity/ %similarity |
|---|---|---|---|---|
| WP_028545674.1 | 531 | NRPS | SnbDE [CBW45647.1] | 33/47 |
| WP_084159208.1 | 185 | hypothetical protein | / | / |
| WP_051287160.1 | 306 | PPTase | Mis12 [AKQ22701.1] | 37/55 |
| WP_028545676.1 | 1127 | S-malonyltransferase | BasH [ERM18800.1] | 48/66 |
| WP_028545677.1 | 355 | histidinol-phosphate transaminase | / | / |
| WP_028545678.1 | 2079 | AT-less type I PKS | BasE [ERM18797.1] | 51/68 |
| WP_051287163.1 | 3777 | AT-less type I PKS | BaeN [CAG23960.2] | 44/61 |
| WP_028545679.1 | 82 | acyl carrier protein | CalX [BAP05574.1] | 57/79 |
| WP_028545680.1 | 408 | beta-ketoacyl synthase | CalW [BAP05575.1] | 59/75 |
| WP_028545681.1 | 421 | HMG-CoA synthase | BaeG [CAG23954.2] | 71/83 |
| WP_028545682.1 | 256 | enoyl-CoA hydratase | BaeH [CAG23955.1] | 52/68 |
| WP_028545683.1 | 249 | enoyl-CoA hydratase | BaeE [CAG23952.1] | 58/73 |
| WP_028545684.1 | 897 | beta-ketoacyl synthase | PsyA [ADA82581.1] | 53/66 |
| WP_028545685.1 | 1872 | NRPS | OzmH [ABS90470.1] | 35/49 |
| WP_051287164.1 | 334 | 2,3-diaminopropionate | Kirromycin [CAN89659.1] | 33/52 |
| WP_051287166.1 | 349 | 2,3-diaminopropionate | / | / |
| WP_028545688.1 | 234 | fibronectin type III domain | / | / |

[a] Number of amino acids

**Table S87.** Predicted functions of ORFs in the NZ_BBDI01000007.1 containing WP_082390521.1 and WP_028559341.1

| gene | aa[a] | putative function | Protein homologue | %identity/ %similarity |
|---|---|---|---|---|
| WP_028559336.1 | 196 | hypothetical protein | / | / |
| WP_054706182.1 | 240 | PPTase | Mis12 [AKQ22701.1] | 35/52 |
| WP_054706184.1 | 313 | acyltransferase | BaeD [CAG23951.1] | 47/65 |
| WP_054706187.1 | 399 | polyketide synthase | Thailandamide [ABC34832.1] | 59/76 |
| WP_054706189.1 | 942 | AT-less type I PKS | BasE [ERM18797.1] | 44/62 |
| WP_082390521.1 | 413 | HMG-CoA synthase | BasE [ERM18797.1] | 46/63 |
| WP_082390522.1 | 287 | polyketide synthase | BasE [ERM18797.1] | 74/88 |
| WP_054706195.1 | 147 | hypothetical protein | DifJ [CAJ57410.1] | 46/67 |
| WP_054706197.1 | 949 | AT-less type I PKS | DifJ [CAJ57410.1] | 40/55 |
| WP_082390523.1 | 180 | polyketide synthase | ElaP [AEC04362.1] | 78/88 |
| WP_082390537.1 | 108 | polyketide synthase | ElaP [AEC04362.1] | 74/85 |
| WP_054706202.1 | 846 | AT-less type I PKS | BaeL [CAG23958.2] | 34/52 |
| WP_054706204.1 | 571 | AT-less type I PKS | BaeN [CAG23960.2] | 43/59 |
| WP_082390524.1 | 753 | beta-ketoacyl synthase | DifI [CAJ57409.1] | 56/71 |
| WP_028559341.1 | 421 | HMG-CoA synthase | BatC [ADD82944.1] | 69/82 |
| WP_054706210.1 | 375 | beta-ketoacyl synthase | PsyA [ADA82581.1] | 64/76 |
| WP_054706212.1 | 498 | AT-less type I PKS | Bat3 [ADD82941.1] | 44/58 |
| WP_082390526.1 | 697 | AT-less type I PKS | JamO [AAS98786.1] | 35/54 |
| WP_054706215.1 | 354 | AT-less type I PKS | RhiB [CAL69889.1] | 43/61 |
| WP_082390527.1 | 199 | AT-less type I PKS | TmpD [CBK62733.1] | 34/47 |
| WP_082390528.1 | 521 | AT-less type I PKS | CorK [ADI59533.1] | 41/57 |
| WP_036708441.1 | 347 | 2,3-diaminopropionate | / | / |
| WP_028559347.1 | 615 | alpha/beta fold hydrolase | MupV [AAM12938.1] | 23/41 |
| WP_082390529.1 | 200 | TetR/AcrR transcriptional regulator | SgvB [AGN74906.1] | 24/54 |
| WP_081666975.1 | 88 | DUF2642 domain | / | / |

[a] Number of amino acids

**Table S88.** Predicted functions of ORFs in NZ_AXBA01000001.1 containing WP_029002374.1

| gene[a] | aa[b] | putative function | Protein homologue | %identity/ %similarity |
|---|---|---|---|---|
| WP_029002372.1 | 249 | thioesterase | CorM [ADI59536.1] | 39/57 |
| WP_051356311.1 | 260 | PPTase | TmlN [CBK62709.1] | 34/52 |
| WP_051356312.1 | 415 | cytochrome P450 | ElaG [AEC04353.1] | 33/53 |
| WP_029002374.1 | 416 | HMG-CoA synthase | JamH [AAS98779.1] | 61/77 |
| WP_051356313.1 | 94 | acyl carrier protein | CalX [BAP05574.1] | 42/72 |
| WP_029002376.1 | 414 | beta-ketoacyl synthase | NspH [ADA69244.1] | 50/67 |
| WP_029002377.1 | 262 | enoyl-CoA hydratase | BaeH [CAG23955.1] | 45/60 |
| WP_051356315.1 | 652 | asparagine synthase | Dor4 [ACY01389.1] | 58/70 |
| WP_035713272.1 | 86 | acyl carrier protein | ChxC [AFO59864.1] | 50/65 |
| WP_029002379.1 | 551 | cyclic peptide export ABC transporter | CalU [BAP05577.1] | 23/41 |
| WP_029002380.1 | 550 | cyclic peptide export ABC transporter | LnmR [AAN85531.1] | 30/48 |
| WP_029002381.1 | 788 | AT/Ox | CalY [BAP05573.1] | 50/66 |
| ORF1 | 6815 | AT-less type I PKS (KS-DH-KR-Acp-KS-B-Acp-KS-Acp-MT-Acp-KS-ECH-Acp-Acp-KS) | Dor5 [ACY01390.1] | 46/57 |
| WP_035713094.1 | 441 | flavin-dependent monooxygenase | OocK [AFX60333.1] | 58/74 |
| ORF2 | 6516 | AT-less type I PKS (Acp-KS-DH-KR-Acp-KS-Acp-KS-DH-Acp-KS-KR-Acp-KS-Acp-C-A-PCP-TE) | Ta1 [ABF85931.1] | 38/51 |
| WP_029002384.1 | 274 | FkbM family methyltransferase | CalA [BAP05589.1] | 34/49 |
| WP_084539553.1 | 431 | cytochrome P450 | BaeS [CAG23962.1] | 33/49 |

[a] ORFs are proteins without protein ID provided by NCBI [b] Number of amino acids

**Table S89.** Predicted functions of ORFs in the NZ_JOID01000025.1 containing WP_030547332.1 and WP_078843784.1

| gene | aa[a] | putative function | Protein homologue | %identity/ %similarity |
|---|---|---|---|---|
| WP_030547330.1 | 288 | S-malonyltransferase | BaeC [CAG23950.2] | 53/69 |
| WP_030547332.1 | 419 | HMG-CoA synthase | BaeG [CAG23954.2] | 70/82 |
| WP_030547334.1 | 81 | acyl carrier protein | AcpK [CAG23953.1] | 56/75 |
| WP_030547335.1 | 406 | beta-ketoacyl synthase | BatB [ADD82943.1] | 60/73 |
| WP_078843784.1 | 412 | HMG-CoA synthase | ADA69245.1 [nosperin] | 59/73 |
| WP_030547339.1 | 255 | enoyl-CoA hydratase | Nosperin [ADA69246.1] | 51/68 |
| WP_030547341.1 | 249 | enoyl-CoA hydratase | CalR [BAP05580.1] | 64/79 |
| WP_030547342.1 | 2531 | AT-less type I PKS | ElaP [AEC04362.1] | 37/53 |
| WP_051723246.1 | 2548 | AT-less type I PKS | BasE [ERM18797.1] | 41/57 |
| WP_051723248.1 | 1850 | AT-less type I PKS | RizB [CCA89326.1] | 45/56 |
| WP_051723250.1 | 1514 | AT-less type I PKS | PsyD [ADA82585.1] | 39/54 |
| WP_030547350.1 | 262 | hypothetical protein | / | / |
| WP_030547352.1 | 467 | adenylation domain | Leinamycin [AAN85501.1] | 31/42 |
| WP_030547354.1 | 261 | SAM-dependent methyltransferase | TaQ [ABF89350.1] | 31/45 |
| WP_030547356.1 | 86 | acyl carrier protein | SmdG [CCC21121.1] | 40/66 |
| WP_078843785.1 | 925 | beta-ketoacyl synthase | SmdI [CCC21123.1] | 53/64 |
| WP_030547357.1 | 338 | acyltransferase | OzmM [ABS90474.1] | 49/61 |
| WP_030547358.1 | 225 | TTPase | LtmL [ACY01405.1] | 49/58 |
| WP_078843786.1 | 300 | MBL fold metallo-hydrolase | / | / |

[a] Number of amino acids

**Table S90.** Predicted functions of ORFs in the NZ_JNFS01000003.1 containing WP_031415145.1

| gene | aa[a] | putative function | Protein homologue | %identity/ %similarity |
|---|---|---|---|---|
| WP_031415118.1 | 4765 | NRPS | AlbI [CAE52339.1] | 31/48 |
| WP_031415119.1 | 2551 | NRPS | VisE [BAF50711.1] | 31/47 |
| WP_031415120.1 | 2549 | NRPS | SnbDE [CBW45647.1] | 35/53 |
| WP_031415121.1 | 3901 | NRPS | SnbDE [CBW45647.1] | 34/51 |
| ORF1 | 4048 | AT-less type I PKS (Acp-Acp-KS-DH-KR-Acp-KS-DH-KR -Acp-KS-Acp) | BaeN [CAG23960.2] | 36/53 |
| ORF2 | 7442 | AT-less type I PKS/NRPS (C-A-PCP-KS-DH-KR-MT-Acp-Acp-KS-C-A-PCP-KS-KR-Acp-KS) | Bat2 [ADD82940.1] | 40/57 |
| ORF3 | 5998 | AT-less type I PKS (GNAT-Acp-KS-KR-Acp-KS-DH-Acp-KS-KR-Acp-KS-DH-Acp-KS) | BaeJ [CAG23957.2] | 48/64 |
| WP_031415140.1 | 294 | pentapeptide repeat-containing protein | / | / |
| WP_035293158.1 | 3050 | NRPS (C-A-PCP-C-A-MT-PCP-C) | SgvD4 [AGN74885.1] | 33/49 |
| WP_031415143.1 | 261 | enoyl-CoA hydratase | BaeH [CAG23956.1] | 56/73 |
| WP_031415145.1 | 418 | HMG-CoA synthase | BaeG [CAG23954.2] | 75/87 |
| WP_031415147.1 | 409 | beta-ketoacyl synthase | CalW [BAP05575.1] | 63/81 |
| WP_031415148.1 | 82 | acyl carrier protein | AcpK [CAG23953.1] | 62/77 |
| WP_081873208.1 | 1135 | S-malonyltransferase | BasH [ERM18800.1] | 49/69 |
| WP_031415152.1 | 222 | MBL fold metallo-hydrolase | BaeB [CAG23949.2] | 50/64 |
| WP_051870661.1 | 248 | PPTase | Mis12 [AKQ22701.1] | 35/50 |
| WP_031415155.1 | 196 | transcription antiterminator | ElaA [AEC04347.1] | 22/40 |
| WP_031415157.1 | 213 | ATP-binding cassette domain | LnmR [AAN85531.1] | 32/49 |
| WP_031415160.1 | 763 | NRPS | CalE [BAP05593.1] | 36/53 |
| WP_051870662.1 | 2875 | NRPS | SnbDE [CBW45647.1] | 30/48 |
| WP_031415163.1 | 1760 | hybrid NRPS/PKS | JamJ [AAS98781.1] | 43/61 |
| WP_035293161.1 | 4533 | hybrid NRPS/PKS | JamP [AAS98787.1] | 44/64 |
| WP_031415167.1 | 274 | GNAT family N-acetyltransferase | DipC [AGS06836.1] | 38/53 |
| WP_035293164.1 | 283 | 3-hydroxyacyl-CoA dehydrogenase | OzmG [ABS90469.1] | 34/56 |
| WP_031415170.1 | 1487 | NRPS (C-A-PCP-E) | JamO [AAS98786.1] | 31/52 |
| WP_035293167.1 | 331 | hypothetical protein | Bat1 [ADD82939.1] | 35/53 |
| WP_031415171.1 | 379 | acyl-CoA dehydrogenase | OzmD [ABA39084.2] | 28/48 |
| WP_031415172.1 | 85 | acyl carrier protein | OzmE [ABS90467.1] | 31/63 |
| WP_035293169.1 | 347 | glyceroyl transferase/phosphatase | OzmB [ABA39082.2] | 44/60 |
| WP_031415174.1 | 1032 | cyclic peptide export ABC transporter | / | / |
| WP_031415175.1 | 322 | 2,3-diaminopropionate | Kirromycin [CAN89659.1] | 32/53 |

[a] ORFs are proteins without protein ID provided by NCBI [b] Number of amino acids

**Table S91.** Predicted functions of ORFs in the NZ_JNLT01000002.1 containing WP_032762518.1

| gene | aa[a] | putative function | Protein homologue | %identity/ %similarity |
|---|---|---|---|---|
| WP_032762507.1 | 410 | cytochrome P450 | BaeS [CAG23962.1] | 43/58 |
| WP_050486702.1 | 301 | short chain dehydrogenase | BatM [ADD82954.1] | 38/57 |
| WP_032762510.1 | 321 | acyltransferase | SorO [ADN68489.1] | 36/55 |
| WP_032762512.1 | 410 | beta-ketoacyl synthase | NspH [ADA69244.1] | 58/75 |
| WP_032762514.1 | 81 | acyl carrier protein | fr9M [AIC32699.1] | 60/80 |
| WP_032762516.1 | 250 | enoyl-CoA hydratase | BatE [ADD82946.1] | 65/80 |
| WP_032762517.1 | 260 | enoyl-CoA hydratase | BaeH [CAG23956.1] | 52/69 |
| WP_032762518.1 | 419 | HMG-CoA synthase | BatC [ADD82944.1] | 69/82 |
| WP_032762523.1 | 376 | ACP-S-malonyltransferase | BonK [AFN27477.1] | 49/62 |
| WP_032762524.1 | 451 | flavin-dependent nitroreductase | OocU [AFX60343.1] | 51/70 |
| WP_050486703.1 | 3113 | AT-less type I PKS | MisF [AKQ22696.1] | 39/56 |
| WP_050486704.1 | 3137 | AT-less type I PKS | BaeL [CAG23958.2] | 41/56 |
| WP_078566461.1 | 3084 | AT-less type I PKS | SorA [ADN68476.1] | 45/58 |
| WP_050486706.1 | 4665 | AT-less type I PKS | TaO [ABF92489.1] | 40/54 |
| WP_106969539.1 | 7961 | AT-less type I PKS | ElaQ [AEC04363.1] | 41/56 |
| WP_050486707.1 | 100 | AT-less type I PKS | DifI [CAJ57409.1] | 29/50 |
| WP_050486708.1 | 1432 | AT-less type I PKS | DifL [CAG23983.1] | 34/58 |
| WP_078566463.1 | 575 | IS1182 family transposase | / | / |
| WP_032762528.1 | 134 | hypothetical protein | / | / |
| WP_106969540.1 | 269 | IS5 family transposase | RhiE [CAL69893.1] | 30/51 |
| WP_032762532.1 | 250 | DUF899 domain-containing protein | DifF [CAG23977.1] | 37/55 |
| WP_032762534.1 | 156 | hypothetical protein | Pederin [AAS47555.1] | 37/51 |
| WP_106969541.1 | 290 | Chitinase | / | / |

[a] Number of amino acids

**Table S92.** Predicted functions of ORFs in the NZ_JPST01000022.1 containing WP_033101211.1

| gene | aa[a] | putative function | Protein homologue | %identity/ %similarity |
|---|---|---|---|---|
| WP_052154233.1 | 281 | PPTase | Mis12 [AKQ22701.1] | 33/46 |
| WP_033101254.1 | 771 | AT/Ox | DifA [CAG23974.1] | 67/77 |
| WP_033101208.1 | 81 | acyl carrier protein | NspG [ADA71314.1] | 62/82 |
| WP_033101209.1 | 222 | MBL fold metallo-hydrolase | BaeB [CAG23949.2] | 52/69 |
| WP_033101255.1 | 260 | alpha/beta hydrolase | MupL [AAM12926.1] | 25/42 |
| WP_052154234.1 | 3370 | AT-less type I PKS | Misakinolide [AKQ22698.1] | 44/60 |
| WP_033101210.1 | 408 | beta-ketoacyl synthase | CalW [BAP05575.1] | 64/80 |
| WP_033101211.1 | 419 | HMG-CoA synthase | BaeG [CAG23954.2] | 79/87 |
| WP_033101258.1 | 254 | enoyl-CoA hydratase | BaeH [CAG23956.1] | 57/74 |
| WP_033101212.1 | 248 | enoyl-CoA hydratase | BatE [ADD82946.1] | 68/84 |
| WP_081943876.1 | 288 | sugar phosphate isomerase | Tartrolon [ACR13997.1] | 35/54 |
| WP_081943886.1 | 211 | 3-deoxy-D-manno-octulosonic-acid transferase | Marinomycin [BAG50469.1] | 21/41 |
| WP_081943877.1 | 68 | hypothetical protein | / | / |
| WP_033101214.1 | 154 | hypothetical protein | / | / |
| WP_033101215.1 | 478 | hypothetical protein | cycloheximide [CCC21128.1] | 52/59 |
| WP_081943878.1 | 467 | MATE family efflux transporter | SorJ [ADN68485.1] | 23/45 |
| WP_052154236.1 | 444 | MATE family efflux transporter | SorJ [ADN68485.1] | 32/50 |
| WP_081943887.1 | 358 | acyltransferase | BaeD [CAG23951.1] | 42/59 |
| WP_033101217.1 | 272 | methyltransferase | Dor10 [ACY01395.1] | 29/48 |
| WP_033101218.1 | 299 | glucose-1-phosphate thymidylyltransferase | Marinomycin [BAG50457.1] | 67/83 |
| WP_033101219.1 | 412 | DUF4309 domain | CorI [ADI59531.1] | 26/43 |
| WP_081943879.1 | 53 | DUF2292 domain | Misakinolide [AKQ22698.1] | 35/52 |
| WP_033101220.1 | 339 | sulfate ABC transporter | cycloheximide [CCC21126.1] | 38/47 |
| WP_033101221.1 | 278 | sulfate ABC transporter permease | / | / |
| WP_033101222.1 | 280 | sulfate ABC transporter permease | / | / |
| WP_081943880.1 | 381 | sulfate ABC transporter | LnmR [AAN85531.1] | 33/48 |

[a] Number of amino acids

**Table S93.** Predicted functions of ORFs in the NZ_JNLE01000003.1 containing WP_033165478.1

| gene | aa[a] | putative function | Protein homologue | %identity/ %similarity |
|---|---|---|---|---|
| WP_051685112.1 | 266 | acetyl-CoA carboxylase | / | / |
| WP_081848768.1 | 275 | acetyl-CoA carboxylase carboxyl transferase | ThaB [ABC35022.1] | 44/62 |
| WP_051685114.1 | 164 | acetyl-CoA carboxylase | / | / |
| WP_033165466.1 | 457 | ATP-grasp domain-containing protein | / | / |
| WP_033165467.1 | 249 | PPTase | Mis12 [AKQ22701.1] | 29/52 |
| WP_033165468.1 | 774 | AT/Ox | DifA [CAG23974.1] | 59/75 |
| WP_033165469.1 | 81 | acyl carrier protein | PsyL [ADA82592.1] | 46/69 |
| WP_051685115.1 | 437 | beta-ketoacyl synthase | DipR [AGS06821.1] | 44/65 |
| WP_033165470.1 | 1733 | AT-less type I PKS | CorK [ADI59533.1] | 32/48 |
| WP_051685116.1 | 1533 | AT-less type I PKS | BaeN [CAG23960.2] | 4057 |
| WP_051685117.1 | 2930 | AT-less type I PKS | MisD [AKQ22698.1] | 34/53 |
| WP_051685118.1 | 1827 | AT-less type I PKS | pks2G | 39/59 |
| WP_033165471.1 | 3480 | AT-less type I PKS | RizB [CCA89326.1] | 36/53 |
| WP_051685119.1 | 1034 | AT-less type I PKS | MisD [AKQ22698.1] | 46/64 |
| WP_033165472.1 | 2981 | AT-less type I PKS | ChiC [AAY89050.1] | 35/52 |
| WP_033165473.1 | 297 | hypothetical protein | / | / |
| WP_033165474.1 | 418 | hypothetical protein | / | / |
| WP_033165475.1 | 216 | HAD family phosphatase | OzmW [ABS90484.1] | 25/42 |
| WP_033165476.1 | 430 | glycosyltransferase | SorF [ADN68481.1] | 35/50 |
| WP_033165477.1 | 253 | enoyl-CoA hydratase | Nosperin [ADA69246.1] | 53/73 |
| WP_033165478.1 | 419 | HMG-CoA synthase | BaeG [CAG23954.2] | 69/82 |
| WP_033165479.1 | 154 | hypothetical protein | / | / |

[a] Number of amino acids

**Table S94.** Predicted functions of ORFs in the NZ_BAZT01000003.1 containing WP_036646965.1

| gene[a] | aa[b] | putative function | Protein homologue | %identity/ %similarity |
|---|---|---|---|---|
| WP_036646958.1 | 125 | ArsR family transcriptional regulator | Cycloheximide [CCC21113.1] | 28/47 |
| WP_036646959.1 | 332 | NADP-dependent oxidoreductase | CalB [BAP05590.1] | 29/47 |
| WP_036646960.1 | 224 | MBL fold metallo-hydrolase | BaeB [CAG23949.2] | 61/72 |
| WP_052020073.1 | 253 | PPTase | Mis12 [AKQ22701.1] | 33/58 |
| WP_036646961.1 | 288 | S-malonyltransferase | BaeC [CAG23950.2] | 62/77 |
| WP_036646962.1 | 768 | S-malonyltransferase | BaeE [CAG23952.1] | 64/77 |
| WP_036646963.1 | 82 | acyl carrier protein | AcpK [CAG23953.1] | 63/79 |
| WP_036646964.1 | 408 | beta-ketoacyl synthase | CalW [BAP05575.1] | 59/77 |
| WP_036646965.1 | 417 | HMG-CoA synthase | BaeG [CAG23954.2] | 78/87 |
| WP_036647161.1 | 255 | enoyl-CoA hydratase | BaeH [CAG23955.1] | 58/71 |
| WP_036646966.1 | 249 | enoyl-CoA hydratase | BaeE [CAG23952.1] | 62/80 |
| ORF1 | 3325 | NRPS | BaeJ [CAG23957.2] | 53/69 |
| ORF2 | 207 | KS | BaeJ [CAG23957.2] | 76/89 |
| ORF3 | 587 | KS | BaeJ [CAG23957.2] | 56/72 |
| ORF4 | 908 | AT-less type I PKS (KR-Acp-KS) | BaeJ [CAG23957.2] | 63/75 |
| ORF5 | 2056 | AT-less type I PKS (DH-Acp-KS-DH-KR-Acp) | BaeL [CAG23958.2] | 55/69 |
| ORF6 | 700 | AT-less type I PKS (KR-Acp) | BaeL [CAG23958.2] | 62/73 |
| ORF7 | 1608 | AT-less type I PKS (Acp-KS-KR-Acp) | BaeL [CAG23958.2] | 54/69 |
| ORF8 | 299 | KS | BaeJ [CAG23957.2] | 67/81 |
| ORF9 | 2402 | AT-less type I PKS (DH-Acp-KS-DH-KR-MT-Acp) | BaeM [CAG23959.2] | 52/67 |
| ORF10 | 426 | KS | BaeM [CAG23959.2] | 74/85 |
| ORF11 | 1523 | AT-less type I PKS (KR-Acp-KS-Acp) | OocS [AFX60341.1] | 41/57 |
| ORF12 | 3455 | hybrid NRPS/PKS | BaeN [CAG23960.2] | 56/72 |
| ORF13 | 603 | KR | BaeN [CAG23960.2] | 47/62 |
| ORF14 | 1080 | AT-less type I PKS (KS-DH) | BaeN [CAG23960.2] | 57/71 |
| ORF15 | 411 | KR | BaeN [CAG23960.2] | 56/69 |
| WP_052020074.1 | 1050 | AT-less type I PKS (MT-Acp-KS) | BaeR [CAG23961.2] | 57/71 |
| WP_052020075.1 | 1428 | AT-less type I PKS (DH-Acp-KS-TE) | BaeR [CAG23961.2] | 33/50 |
| WP_036646969.1 | 404 | cytochrome P450 | BaeS [CAG23962.1] | 63/79 |
| WP_036646970.1 | 1189 | hypothetical protein | / | / |

[a] ORFs are proteins without protein ID provided by NCBI [b] Number of amino acids

**Table S95.** Predicted functions of ORFs in the NZ_KE384560.1 containing WP_037576445.1

| gene | aa[a] | putative function | Protein homologue | %identity/ %similarity |
|---|---|---|---|---|
| WP_028981300.1 | 310 | AT-less type I PKS | BryB [ABM63527.1] | 38/62 |
| WP_051313368.1 | 3587 | AT-less type I PKS | MisF [AKQ22696.1] | 43/60 |
| WP_028981301.1 | 2079 | AT-less type I PKS | OnnB [AAV97870.1] | 41/56 |
| WP_081671125.1 | 289 | AT-less type I PKS | BasE [ERM18797.1] | 53/71 |
| WP_028981302.1 | 2178 | AT-less type I PKS | SorE [ADN68480.1] | 42/59 |
| WP_037576442.1 | 2415 | AT-less type I PKS | DszA [AAY32964.1] | 34/52 |
| WP_028981304.1 | 312 | alcohol dehydrogenase | Enacyloxin [ABI91459.1] | 27/46 |
| WP_028981305.1 | 324 | hypothetical protein | Marinomycin [BAG50483.1] | 26/47 |
| WP_037576445.1 | 419 | HMG-CoA synthase | BatC [ADD82944.1] | 71/84 |
| WP_028981306.1 | 263 | enoyl-CoA hydratase | BaeH [CAG23956.1] | 58/71 |
| WP_028981307.1 | 249 | enoyl-CoA hydratase | BatE [ADD82946.1] | 65/85 |
| WP_028981308.1 | 863 | hypothetical protein | / | / |
| WP_028981309.1 | 292 | outer membrane protein | CalG [BAP05595.1] | 33/57 |
| WP_028981310.1 | 444 | hypothetical protein | / | / |
| WP_051313372.1 | 250 | DUF4386 domain | ElaQ [AEC04363.1] | 23/48 |
| WP_028981311.1 | 231 | hypothetical protein | PedF [AAS47564.1] | 34/50 |
| WP_028981312.1 | 327 | alcohol dehydrogenase | DifL [CAG23983.1] | 30/48 |
| WP_081671075.1 | 321 | acyltransferase | BaeD [CAG23951.1] | 35/58 |
| WP_051313374.1 | 234 | MBL fold metallo-hydrolase | BaeB [CAG23949.2] | 41/62 |
| WP_028981315.1 | 85 | acyl carrier protein | DipF [AGS06833.1] | 45/74 |
| WP_028981316.1 | 424 | beta-ketoacyl synthase | DipR [AGS06821.1] | 52/70 |
| WP_028981317.1 | 388 | ACP-S-malonyltransferase | MisG [AKQ22695.1] | 54/69 |
| WP_028981318.1 | 223 | PPTase | VirK [BAF50717.1] | 23/41 |
| WP_028981319.1 | 178 | hypothetical protein | BonB [AFN27481.1] | 33/51 |

[a] Number of amino acids

**Table S96.** Predicted functions of ORFs in the NZ_APMV01000005.1 containing WP_038924941.1

| gene | aa[a] | putative function | Protein homologue | %identity/ %similarity |
|---|---|---|---|---|
| WP_016940753.1 | 202 | hypothetical protein | OocB [AFX60324.1] | 72/84 |
| WP_016940752.1 | 249 | enoyl-CoA hydratase | OocC [AFX60325.1] | 86/91 |
| WP_029456274.1 | 262 | enoyl-CoA hydratase | OocC [AFX60325.1] | 84/94 |
| WP_038924941.1 | 420 | HMG-CoA synthase | OocE [AFX60327.1] | 89/93 |
| WP_033581351.1 | 418 | beta-ketoacyl synthase | OocF [AFX60328.1] | 81/89 |
| WP_016940748.1 | 83 | acyl carrier protein | OocG [AFX60329.1] | 76/86 |
| WP_038924942.1 | 3832 | FkbM family methyltransferase | OocJ [AFX60332.1] | 69/77 |
| WP_016940746.1 | 434 | flavin-dependent monooxygenase | OocK [AFX60333.1] | 91/ |
| WP_038924943.1 | 2523 | AT-less type I PKS | OocL [AFX60334.1] | 67/76 |
| WP_016940744.1 | 377 | LLM class flavin-dependent oxidoreductase | OocM [AFX60335.1] | 89/92 |
| WP_016940742.1 | 89 | acyl carrier protein | OocO [AFX60337.1] | 88/92 |
| WP_016940741.1 | 388 | hypothetical protein | OocP [AFX60338.1] | 91/95 |
| WP_020477011.1 | 121 | hypothetical protein | OocQ [AFX60339.1] | 79/88 |
| WP_038924946.1 | 2354 | AT-less type I PKS | OocR [AFX60340.1] | 73/81 |
| WP_038924947.1 | 5547 | AT-less type I PKS | OocS [AFX60341.1] | 68/77 |
| WP_016940737.1 | 276 | hypothetical protein | OocT [AFX60342.1] | 73/82 |
| WP_038924948.1 | 481 | flavin-dependent nitroreductase | OocU [AFX60343.1] | 80/89 |
| WP_071526186.1 | 639 | S-malonyltransferase | OocV [AFX60344.1] | 75/84 |
| WP_016940734.1 | 379 | S-malonyltransferase | OocW [AFX60345.1] | 73/82 |

[a] Number of amino acids

**Table S97.** Predicted functions of ORFs in the NZ_JSZF01000085.1 containing WP_039412188.1

| gene | aa[a] | putative function | Protein homologue | %identity/ %similarity |
|---|---|---|---|---|
| WP_082002640.1 | 1849 | AT-less type I PKS | FR9I [AIC32695.1] | 65/72 |
| WP_052210040.1 | 305 | S-malonyltransferase | FR9J [AIC32696.1] | 71/78 |
| WP_081419541.1 | 154 | cyclase | FR9Q [AIC32703.1] | 30/41 |
| WP_039412188.1 | 419 | HMG-CoA synthase | FR9K [AIC32697.1] | 87/91 |
| WP_039412190.1 | 261 | enoyl-CoA hydratase | BaeH [CAG23955.1] | 53/66 |
| WP_039412192.1 | 79 | acyl carrier protein | FR9M [AIC32699.1] | 66/81 |
| WP_039412195.1 | 426 | beta-ketoacyl synthase | FR9P [AIC32700.1] | 75/81 |
| WP_039412197.1 | 377 | S-malonyltransferase | fr9O [AIC32701.1] | 71/82 |
| WP_081419540.1 | 323 | acyltransferase | SorO [ADN68489.1] | 49/65 |
| WP_081419539.1 | 266 | PPTase | BatI [ADD82950.1] | 41/60 |

[a] Number of amino acids

**Table S98.** Predicted functions of ORFs in the NZ_CP009280.1 containing WP_042135866.1 and WP_042135870.1

| gene[a] | aa[b] | putative function | protein homologue | %identity/ %similarity |
|---|---|---|---|---|
| WP_042135863.1 | 120 | DUF3221 domain-containing protein | MxnJ [AGS77290.1] | 41/48 |
| WP_081969930.1 | 220 | hypothetical protein | TaA [ABF91060.1] | 29/44 |
| WP_042135866.1 | 419 | HMG-CoA synthase | BaeG [CAG23954.2] | 72/84 |
| WP_042135868.1 | 255 | enoyl-CoA hydratase | BaeH [CAG23956.1] | 57/72 |
| WP_052421794.1 | 1810 | AT-less type I PKS | BasG [ERM18799.1] | 46/64 |
| WP_042135870.1 | 420 | HMG-CoA synthase | BatC [ADD82944.1] | 69/82 |
| WP_042141021.1 | 249 | enoyl-CoA hydratase | BatE [ADD82946.1] | 64/79 |
| ORF1 | 1755 | AT-less type I PKS | BaeL [CAG23958.2] | 53/66 |
| WP_081969934.1 | 2257 | AT-less type I PKS | DifH [CAJ57408.1] | 47/63 |
| ORF2 | 6126 | AT-less type I PKS | BaeM [CAG23959.2] | 46/62 |
| WP_081970479.1 | 587 | methyltransferase domain | CalF [BAP05594.1] | 45/58 |
| WP_052421797.1 | 4448 | AT-less type I PKS | MmpD [AAM12913.1] | 38/52 |
| WP_042135874.1 | 559 | NRPS | DifL [CAG23983.1] | 38/55 |
| WP_081969935.1 | 403 | NADH:flavin oxidoreductase | MupC [AAM12914.1] | 31/50 |
| WP_042135876.1 | 162 | hypothetical protein | Oxazolomycin [ABS90486.1] | 18/36 |
| WP_042135878.1 | 116 | hypothetical protein | MmpA [AAM12909.2] | 34/53 |
| WP_042135879.1 | 403 | MFS transporter | SgvT1 [AGN74873.1] | 21/45 |

[a] ORFs are proteins without protein ID provided by NCBI [b] Number of amino acids

**Table S99.** Predicted functions of ORFs in the NZ_CP009288.1 containing WP_042206744.1

| gene[a] | aa[b] | putative function | protein homologue | %identity/ %similarity |
|---|---|---|---|---|
| WP_042206736.1 | 149 | MarR family transcriptional regulator | / | / |
| WP_042206736.1 | 335 | AraC family transcriptional regulator | / | / |
| WP_042206738.1 | 264 | oxidoreductase | ChxH [AFO59869.1] | 26/40 |
| WP_042206739.1 | 178 | transcription antiterminator | TaA [ABF91060.1] | 23/43 |
| WP_052410216.1 | 264 | PPTase | Mis12 [AKQ22701.1] | 31/50 |
| WP_042206740.1 | 225 | MBL fold metallo-hydrolase | BaeB [CAG23949.2] | 53/70 |
| WP_042206741.1 | 320 | acyltransferase | BaeD [CAG23951.1] | 45/62 |
| WP_042206742.1 | 787 | AT/Ox | DifA [CAG23974.1] | 50/69 |
| WP_081949519.1 | 88 | acyl carrier protein | OocG [AFX60329.1] | 43/65 |
| WP_052410217.1 | 425 | beta-ketoacyl synthase | CorD [ADI59526.1] | 44/63 |
| WP_042206744.1 | 410 | HMG-CoA synthase | MxnE [AGS77285.1] | 57/74 |
| WP_042206745.1 | 256 | enoyl-CoA hydratase | MxnF [AGS77286.1] | 40/63 |
| WP_042206746.1 | 226 | ABC transporter ATP-binding protein | LnmR [AAN85531.1] | 32/51 |
| WP_042206747.1 | 413 | ABC transporter permease | / | / |
| WP_042206748.1 | 440 | ABC transporter permease | / | / |
| WP_052410218.1 | 1569 | AT-less type I PKS | BaeJ [CAG23957.2] | 44/59 |
| WP_052410219.1 | 2047 | AT-less type I PKS | DszB [AAY32965.1] | 37/52 |
| WP_042206749.1 | 3363 | AT-less type I PKS | RizD [CCA89328.1] | 39/55 |
| WP_081949520.1 | 2107 | AT-less type I PKS | DszB [AAY32965.1] | 35/51 |
| ORF1 | 5042 | AT-less type I PKS | DszB [AAY32965.1] | 38/54 |
| WP_081949868.1 | 303 | AT-less type I PKS | MisF [AKQ22696.1] | 39/54 |
| WP_042206752.1 | 3936 | AT-less type I PKS | DszB [AAY32965.1] | 36/52 |
| WP_042206753.1 | 1670 | AT-less type I PKS | DszA [AAY32964.1] | 38/57 |
| WP_052410220.1 | 1802 | AT-less type I PKS | ChiF [AAY89053.1] | 43/58 |
| WP_052410221.1 | 2021 | AT-less type I PKS | ChiE [AAY89052.1] | 38/55 |
| WP_052410222.1 | 2533 | AT-less type I PKS | ChiF [AAY89053.1] | 44/57 |
| WP_042206754.1 | 749 | AT-less type I PKS | ChiF [AAY89053.1] | 32/50 |
| WP_042206755.1 | 189 | dihydrofolate reductase | Leinamycin [AAN85498.1] | 25/38 |
| WP_042206756.1 | 257 | sugar phosphate isomerase | / | / |
| WP_042206757.1 | 111 | EthD family reductase | OtnB [BAG50482.1] | 27/53 |
| WP_042209355.1 | 323 | esterase | oxazolomycin [ABS90461.1] | 24/41 |
| WP_042206758.1 | 90 | hypothetical protein | DifG [CAG23978.1] | 38/58 |
| WP_042209356.1 | 322 | AraC family transcriptional regulator | / | / |
| WP_042206759.1 | 468 | MFS transporter | LnmY [AAN85538.1] | 35/51 |
| WP_042206760.1 | 255 | ABC transporter permease | / | / |
| WP_042206761.1 | 256 | glycosyl transferase family 8 | lkcJ [BAC76467.1] | 29/46 |
| WP_042206762.1 | 288 | pentapeptide repeat protein | AlbXIX [CAE52332.1] | 21/38 |
| WP_042206763.1 | 180 | TetR/AcrR family | Leinamycin [AAN85544.1] | 29/51 |
| WP_042206764.1 | 751 | excinuclease ABC subunit UvrA | SgvT2 [AGN74890.1] | 26/40 |

[a] ORFs are proteins without protein ID provided by NCBI [b] Number of amino acids

**Table S100.** Predicted functions of ORFs in the NZ_CP009282.1 containing WP_042236664.1

| gene | aa[a] | putative function | Protein homologue | %identity/ %similarity |
|---|---|---|---|---|
| WP_042236661.1 | 170 | hypothetical protein | / | / |
| WP_081956550.1 | 273 | PPTase | Mis12 [AKQ22701.1] | 29/51 |
| WP_052416441.1 | 263 | enoyl-CoA hydratase | JamI [AAS98780.1] | 36/58 |
| WP_042236664.1 | 410 | HMG-CoA synthase | CorE [ADI59527.1] | 58/73 |
| WP_042236665.1 | 434 | beta-ketoacyl synthase | CorD [ADI59526.1] | 44/62 |
| WP_042236666.1 | 79 | acyl carrier protein | PsyL [ADA82592.1] | 46/71 |
| WP_042236667.1 | 1132 | S-malonyltransferase | BasH [ERM18800.1] | 42/61 |
| WP_081956551.1 | 235 | MBL fold metallo-hydrolase | BaeB [CAG23949.2] | 48/66 |
| WP_042236669.1 | 852 | alpha/beta fold hydrolase | JamJ [AAS98781.1] | 32/51 |
| WP_042236671.1 | 2993 | AT-less type I PKS | PedH [AAS47562.1] | 35/52 |
| WP_042236672.1 | 3080 | AT-less type I PKS | OzmN [ABS90475.1] | 38/51 |
| WP_052416442.1 | 1275 | AT-less type I PKS | DszB [AAY32965.1] | 38/56 |
| WP_052416443.1 | 4145 | AT-less type I PKS | OzmN [ABS90475.1] | 35/49 |
| WP_052416445.1 | 712 | AT-less type I PKS | OocJ [AFX60332.1] | 42/59 |
| WP_081956552.1 | 2137 | AT-less type I PKS | ChiF [AAY89053.1] | 38/54 |
| WP_052416448.1 | 2102 | AT-less type I PKS | DifI [CAJ57409.1] | 32/48 |
| WP_052416450.1 | 2818 | AT-less type I PKS | DszA [AAY32964.1] | 34/50 |
| WP_081956553.1 | 206 | hypothetical protein | / | / |
| WP_042236677.1 | 582 | ferredoxin | ThaR [ABC36202.1] | 21/35 |

[a] Number of amino acids

**Table S101.** Predicted functions of ORFs in the NZ_KN039946.1 containing WP_043371337.1

| gene | aa[a] | putative function | Protein homologue | %identity/ %similarity |
|---|---|---|---|---|
| WP_043371335.1 | 4117 | AT-less type I PKS | DszB [AAY32965.1] | 40/52 |
| WP_052411981.1 | 1637 | AT-less type I PKS | JamL [AAS98783.1] | 32/49 |
| WP_043371337.1 | 421 | HMG-CoA synthase | ThaK [ABC34601.1] | 71/82 |
| WP_043371340.1 | 252 | enoyl-CoA hydratase | BaeE [CAG23952.1] | 58/72 |
| WP_043371343.1 | 80 | acyl carrier protein | OocG [AFX60329.1] | 39/66 |
| WP_043376750.1 | 245 | oxidoreductase | DifE [CAG23976.1] | 40/59 |
| WP_043371346.1 | 523 | acyl-CoA carboxylase | SnbS [CBW45762.1] | 91/94 |
| WP_043371347.1 | 68 | hypothetical protein | / | / |
| WP_052411982.1 | 382 | hypothetical protein | / | / |
| WP_052412144.1 | 324 | ABC transporter ATP-binding protein | LnmR [AAN85531.1] | 34/47 |
| WP_043371348.1 | 250 | ABC transporter permease | / | / |

[a] Number of amino acids

**Table S102.** Predicted functions of ORFs in the NZ_BACU01000416.1 containing WP_044562062.1

| gene | aa[a] | putative function | Protein homologue | %identity/ %similarity |
|---|---|---|---|---|
| WP_044562076.1 | 464 | PfaD family polyunsaturated fatty acid | dipL [AGS06827.1] | 45/67 |
| WP_083901224.1 | 438 | cytochrome P450 | CorO [ADI59538.1] | 34/51 |
| WP_044562057.1 | 447 | cytochrome P450 | CorO [ADI59538.1] | 33/49 |
| WP_085940054.1 | 266 | Ser/Thr protein phosphatase | ThaC [ABC35295.1] | 29/41 |
| WP_044562059.1 | 78 | acyl carrier protein | / | / |
| WP_044562060.1 | 451 | hypothetical protein | / | / |
| WP_044562061.1 | 249 | enoyl-CoA hydratase | BatE [ADD82946.1] | 50/68 |
| WP_083901226.1 | 259 | enoyl-CoA hydratase | CylG [ARU81121.1] | 33/57 |
| WP_044562062.1 | 407 | HMG-CoA synthase | DifN [CAG23985.1] | 50/64 |
| WP_049974260.1 | 411 | beta-ketoacyl synthase | CorD [ADI59526.1] | 41/53 |
| WP_083901230.1 | 77 | acyl carrier protein | PtzD [AHC73995.1] | 59/75 |
| WP_044562064.1 | 268 | SDR family oxidoreductase | DifE [CAG23976.1] | 27/43 |
| WP_083901231.1 | 203 | hypothetical protein | TmpD [CBK62733.1] | 32/55 |
| WP_049974261.1 | 1147 | oxidoreductase | PtzB [AHC73997.1] | 62/74 |
| WP_083901232.1 | 200 | acyl carrier protein | CalE [BAP05593.1] | 26/40 |
| WP_085940052.1 | 3150 | AT-less type I PKS | CorL [ADI59534.1] | 35/45 |
| WP_044562068.1 | 120 | hypothetical protein | / | / |
| WP_044562069.1 | 89 | hypothetical protein | / | / |
| WP_083901227.1 | 679 | AarF/ABC1/UbiB kinase | Rhizopodin [CCA89332.1] | 38/54 |
| WP_044562071.1 | 292 | S-malonyltransferase | RhiG [CAL69887.1] | 55/71 |
| WP_044562072.1 | 314 | hypothetical protein | PedC [AAS47559.1] | 31/51 |
| WP_085940053.1 | 477 | SagB/ThcOx family dehydrogenase | / | / |
| WP_083901228.1 | 1695 | AT-less type I PKS | Ena5925 [ABI91470.1] | 37/50 |
| WP_044562074.1 | 69 | hypothetical protein | / | / |

[a] Number of amino acids

**Table S103.** Predicted functions of ORFs in the NZ_CP007142.1 containing WP_044617470.1

| gene[a] | aa[b] | putative function | protein homologue | %identity/ %similarity |
|---|---|---|---|---|
| WP_044617467.1 | 61 | hypothetical protein | / | / |
| WP_044617468.1 | 252 | enoyl-CoA hydratase | CalR [BAP05580.1] | 62/78 |
| WP_044617469.1 | 261 | enoyl-CoA hydratase | OocD [AFX60326.1] | 58/77 |
| WP_044617470.1 | 420 | HMG-CoA synthase | BatC [ADD82944.1] | 69/84 |
| ORF1 | 6819 | AT-less type I PKS | OocJ [AFX60332.1] | 46/60 |
| ORF2 | 5472 | AT-less type I PKS | RhiC [CAL69890.1] | 43/57 |
| ORF3 | 380 | AT-less type I PKS | OocM [AFX60335.1] | 56/70 |
| ORF4 | 4258 | AT-less type I PKS | OocN [AFX60336.1] | 45/58 |
| ORF5 | 6350 | AT-less type I PKS | MisF [AKQ22696.1] | 40/56 |
| WP_082070961.1 | 448 | AT-less type I PKS | OocS [AFX60341.1] | 41/58 |
| WP_044617475.1 | 493 | apocarotenoid-15,15'-oxygenase | / | / |
| WP_082070718.1 | 809 | hypothetical protein | / | / |
| WP_044617477.1 | 412 | cytochrome P450 | Streptimidone [ACY01404.1] | 33/51 |
| WP_044617478.1 | 79 | acyl carrier protein | Fr9M [AIC32699.1] | 51/79 |
| WP_044617479.1 | 409 | beta-ketoacyl synthase | NspH [ADA69244.1] | 55/73 |
| WP_044617480.1 | 774 | hypothetical protein | / | / |
| WP_044617481.1 | 178 | hypothetical protein | / | / |

[a] ORFs are proteins without protein ID provided by NCBI [b] Number of amino acids

**Table S104.** Predicted functions of ORFs in the NZ_LANC01000010.1 containing WP_045825686.1

| gene | aa[a] | putative function | Protein homologue | %identity/ %similarity |
|---|---|---|---|---|
| WP_045825684.1 | 4354 | AT-less type I PKS | ElaQ [AEC04363.1] | 37/55 |
| WP_082075875.1 | 455 | hypothetical protein | MxnI [AGS77289.1] | 29/48 |
| WP_045826001.1 | 452 | PfaD family | Myxovirescins [ABF87992.1] | 54/72 |
| WP_045825686.1 | 419 | HMG-CoA synthase | BonG [AFN27479.1] | 69/82 |
| WP_045825687.1 | 272 | enoyl-CoA hydratase | BaeH [CAG23955.1] | 52/72 |
| WP_045825688.1 | 249 | enoyl-CoA hydratase | BaeE [CAG23952.1] | 59/73 |
| WP_045825689.1 | 230 | MBL fold metallo-hydrolase | BaeB [CAG23949.2] | 42/60 |
| WP_045825690.1 | 83 | acyl carrier protein | NspG [ADA71314.1] | 54/74 |
| WP_045825691.1 | 416 | beta-ketoacyl synthase | BonF [AFN27478.1] | 55/72 |
| WP_045825692.1 | 383 | S-malonyltransferase | Fr9O [AIC32701.1] | 52/70 |
| WP_045825693.1 | 314 | acyltransferase | SorO [ADN68489.1] | 36/57 |
| WP_045825694.1 | 398 | AraC family transcriptional regulator | / | / |

[a] Number of amino acids

**Table S105.** Predicted functions of ORFs in the NZ_JZJL01000040.1 containing WP_046158218.1

| gene | aa[a] | putative function | protein homologue | %identity/ %similarity |
|---|---|---|---|---|
| WP_052729380.1 | 4438 | AT-less type I PKS | BaeN [CAG23960.2] | 38/54 |
| WP_052729381.1 | 233 | MBL fold metallo-hydrolase | BaeB [CAG23949.2] | 42/58 |
| WP_046158218.1 | 419 | HMG-CoA synthase | Fr9K [AIC32697.1] | 72/84 |
| WP_046158219.1 | 262 | enoyl-CoA hydratase/isomerase | Fr9L [AIC32698.1] | 61/76 |
| WP_046158220.1 | 249 | enoyl-CoA hydratase | BatE [ADD82946.1] | 63/75 |
| WP_021476122.1 | 79 | acyl carrier protein | Fr9M [AIC32699.1] | 60/76 |
| WP_046158221.1 | 411 | beta-ketoacyl synthase | BonF [AFN27478.1] | 61/72 |
| WP_046158222.1 | 376 | ACP-S-malonyltransferase | fr9O [AIC32701.1] | 60/73 |
| WP_046158223.1 | 312 | acyltransferase domain | BatH [ADD82949.1] | 52/65 |
| WP_052717349.1 | 111 | hypothetical protein | BryA [ABM63537.1] | 29/44 |
| WP_046158224.1 | 229 | NAD-dependent epimerase | / | / |

[a] Number of amino acids

**Table S106.** Predicted functions of ORFs in the NZ_CP023738.1 containing WP_051418953.1

| gene [a] | aa[b] | putative function | Protein homologue | %identity/ %similarity |
|---|---|---|---|---|
| WP_003612726.1 | 3236 | NRPS/PKS | AlbI [CAE52339.1] | 32/46 |
| WP_003612728.1 | 3114 | AT-less type I PKS | kirAIV [CAN89634.1] | 39/50 |
| WP_099831921.1 | 2547 | AT-less type I PKS | BryB [ABM63527.1] | 34/47 |
| WP_099831922.1 | 1919 | AT-less type I PKS (KR-MT-Acp-KS) | CorL [ADI59534.1] | 37/49 |
| WP_099831923.1 | 514 | AT-less type I PKS (KR-Acp) | CorL [ADI59534.1] | 33/45 |
| WP_003614160.1 | 1250 | AT-less type I PKS (KS-oMT) | PtzB [AHC73997.1] | 45/59 |
| ORF1 | 2857 | AT-less type I PKS (KS-DH-KR-Acp-KS-DH-DH-KR-MT) | OzmN [ABS90475.1] | 37/47 |
| WP_099831924.1 | 2951 | AT-less type I PKS | OzmN [ABS90475.1] | 37/48 |
| WP_099831925.1 | 219 | acyl carrier protein | Fr9DEF [AIC32693.1] | 38/60 |
| WP_024750044.1 | 268 | KR | DifE [CAG23976.1] | 27/47 |
| WP_099831926.1 | 3641 | AT-less type I PKS | PtzD [AHC73995.1] | 41/56 |
| WP_024750042.1 | 401 | beta-ketoacyl synthase | MxnD [AGS77284.1] | 37/52 |
| WP_051418953.1 | 406 | HMG-CoA synthase | JamH [AAS98779.1] | 49/68 |
| WP_003614670.1 | 257 | enoyl-CoA hydratase | CylG [ARU81121.1] | 37/57 |
| WP_003614677.1 | 248 | enoyl-CoA hydratase | BatE [ADD82946.1] | 51/65 |
| WP_003614678.1 | 84 | type II toxin-antitoxin system | / | / |

[a] ORFs are proteins without protein ID provided by NCBI [b] Number of amino acids

**Table S107.** Predicted functions of ORFs in the NZ_BBCG01000005.1 containing WP_054739556.1

| gene | aa[a] | putative function | Protein homologue | %identity/ %similarity |
|---|---|---|---|---|
| WP_054739540.1 | 391 | biotin/lipoyl-binding protein | / | / |
| WP_054739542.1 | 423 | beta-ketoacyl synthase | DipR [AGS06821.1] | 43/66 |
| WP_083461231.1 | 93 | acyl carrier protein | AcpK [CAG23953.1] | 44/67 |
| WP_083461232.1 | 550 | NRPS | AlbI [CAE52339.1] | 23/45 |
| WP_054739548.1 | 650 | NPRS | Ta1 [ABF85931.1] | 34/53 |
| WP_054739550.1 | 444 | glycoside hydrolase family | / | / |
| WP_054739552.1 | 152 | Raf kinase inhibitor | / | / |
| WP_083461233.1 | 582 | enoyl-CoA hydratase | BatE [ADD82946.1] | 47/67 |
| WP_083461234.1 | 262 | enoyl-CoA hydratase | BaeH [CAG23956.1] | 47/61 |
| WP_054739556.1 | 416 | HMG-CoA synthase | ElaL [AEC04358.1] | 67/81 |
| WP_083461238.1 | 61 | acyl carrier protein | DifL [CAG23983.1] | 50/76 |
| WP_054739558.1 | 212 | SDR family oxidoreductase | BasF [ERM18798.1] | 53/70 |
| WP_083461235.1 | 527 | SDR family oxidoreductase | BasG [ERM18799.1] | 32/58 |
| WP_054739561.1 | 1138 | methyltransferase | BasE [ERM18797.1] | 48/65 |
| WP_054739563.1 | 807 | AT-less type I PKS | SorE [ADN68480.1] | 47/65 |
| WP_054739565.1 | 1984 | AT-less type I PKS | SorB [ADN68477.1] | 37/56 |
| WP_083461236.1 | 3802 | AT-less type I PKS | BaeN [CAG23960.2] | 36/55 |
| WP_054739568.1 | 1449 | AT-less type I PKS | DifH [CAJ57408.1] | 33/50 |
| WP_054739570.1 | 858 | AT-less type I PKS | BasD [ERM18796.1] | 36/58 |
| WP_054739572.1 | 536 | AT-less type I PKS | BasE [ERM18797.1] | 58/75 |
| WP_054739574.1 | 557 | AT-less type I PKS | ElaO [AEC04361.1] | 52/70 |
| WP_054739576.1 | 395 | AT-less type I PKS | BasE [ERM18797.1] | 42/64 |
| WP_054739579.1 | 1251 | AT-less type I PKS | MisC [AKQ22699.1] | 31/49 |
| WP_054739581.1 | 770 | AT/Ox | DifA [CAG23974.1] | 52/69 |
| WP_054739583.1 | 253 | Ser/Thr protein phosphatase | ThaC [ABC35295.1] | 41/59 |
| WP_083461237.1 | 230 | IS91 family transposase | / | / |

[a] Number of amino acids

**Table S108.** Predicted functions of ORFs in the NZ_LJWU01000040.1 containing WP_055129622.1

| gene | aa[a] | putative function | Protein homologue | %identity/ %similarity |
|---|---|---|---|---|
| WP_055129613.1 | 382 | catechol 1,2-dioxygenase | / | / |
| WP_055129614.1 | 255 | enoyl-CoA hydratase | CorF [ADI59528.1] | 38/53 |
| WP_082460511.1 | 1253 | AMP-dependent ligase | JamA [AAS98774.1] | 39/59 |
| WP_055129616.1 | 1258 | AT-less type I PKS | PedH [AAS47562.1] | 40/55 |
| WP_055129617.1 | 1152 | AT-less type I PKS | FR9GH [AIC32694.1] | 35/50 |
| WP_082460512.1 | 2655 | AT-less type I PKS | LglD [AIU36100.1] | 34/53 |
| WP_055129619.1 | 357 | flavin-dependent oxidoreductase | Myxovirescin [ABF90561.1] | 46/65 |
| WP_055129620.1 | 920 | AT-less type I PKS | CalE [BAP05593.1] | 38/56 |
| WP_082460513.1 | 456 | APC family permease | / | / |
| WP_055129622.1 | 409 | HMG-CoA synthase | MxnE [AGS77285.1] | 52/66 |
| WP_082460514.1 | 416 | beta-ketoacyl synthase | DipR [AGS06821.1] | 33/54 |
| WP_055129624.1 | 92 | acyl carrier protein | / | / |
| WP_082460515.1 | 296 | PPTase | SnaN [CBW45738.1] | 28/42 |
| WP_055129626.1 | 760 | S-malonyltransferase | BaeE [CAG23952.1] | 45/61 |
| WP_055129627.1 | 386 | epoxide hydrolase | / | / |

[a] Number of amino acids

**Table S109.** Predicted functions of ORFs in the NZ_FCOK02000016.1 containing WP_062085598.1

| gene | aa[a] | putative function | Protein homologue | %identity/ %similarity |
|---|---|---|---|---|
| WP_063977806.1 | 247 | short-chain dehydrogenase | / | / |
| WP_062085589.1 | 316 | hypothetical protein | / | / |
| WP_082913389.1 | 272 | PPTase | BatI [ADD82950.1] | 50/65 |
| WP_062085591.1 | 382 | ACP-S-malonyltransferase | fr9O [AIC32701.1] | 61/74 |
| WP_062085593.1 | 415 | beta-ketoacyl synthase | fr9N [AIC32700.1] | 68/78 |
| WP_062085595.1 | 79 | acyl carrier protein | fr9M [AIC32699.1] | 59/78 |
| WP_062085683.1 | 249 | enoyl-CoA hydratase | BatE [ADD82946.1] | 63/79 |
| WP_062085596.1 | 261 | enoyl-CoA hydratase | Fr9L [AIC32698.1] | 62/74 |
| WP_062085598.1 | 419 | HMG-CoA synthase | fr9K [AIC32697.1] | 73/85 |
| WP_062085600.1 | 233 | MBL fold metallo-hydrolase | BaeB [CAG23949.2] | 43/58 |
| WP_082913396.1 | 454 | hypothetical protein | BaeN [CAG23960.2] | 35/53 |
| WP_062085604.1 | 4759 | AT-less type I PKS | Bat2 [ADD82940.1] | 39/54 |
| WP_082913397.1 | 447 | hypothetical protein | MisF [AKQ22696.1] | 37/58 |
| WP_062085608.1 | 88 | acyl carrier protein | DifC [CAJ57406.1] | 38/65 |
| WP_062085610.1 | 245 | SDR family oxidoreductase | DifE [CAG23976.1] | 48/72 |
| WP_062085612.1 | 481 | long-chain fatty acid--CoA ligase | DifD [CAJ57407.1] | 45/65 |
| WP_062085614.1 | 265 | hypothetical protein | / | / |

[a] Number of amino acids

**Table S110.** Predicted functions of ORFs in the NZ_FKCG01000006.1 containing WP_063925844.1

| ene | aa [a] | putative function | Protein homologue | %identity/ %similarity |
|---|---|---|---|---|
| WP_080472785.1 | 1056 | AT-less type I PKS | MxnK [AGS77291.1] | 38/56 |
| WP_063925840.1 | 1010 | AT-less type I PKS | SorA [ADN68476.1] | 32/50 |
| WP_063925841.1 | 1202 | AT-less type I PKS | Ta1 [ABF85931.1] | 32/48 |
| WP_063925842.1 | 855 | AT-less type I PKS | MxnK [AGS77291.1] | 34/50 |
| WP_080472786.1 | 258 | enoyl-CoA hydratase | MxnG [AGS77287.1] | 53/67 |
| ORF1 | 126 | enoyl-CoA hydratase | CylG [ARU81121.1] | 36/53 |
| WP_063925844.1 | 409 | HMG-CoA synthase | MxnE [AGS77285.1] | 53/70 |
| WP_080472787.1 | 447 | beta-ketoacyl synthase | PedM [AAW33972.1] | 38/57 |
| WP_063925846.1 | 81 | acyl carrier protein | ThaI [ABC35804.1] | 42/60 |
| WP_063925847.1 | 276 | S-malonyltransferase | BaeC [CAG23950.2] | 46/59 |
| WP_063925848.1 | 391 | hypothetical protein | / | / |
| WP_063925849.1 | 233 | PPTase | dipB [AGS06837.1] | 25/52 |
| WP_063925850.1 | 234 | IS6 family transposase | / | / |

[a] Number of amino acids

**Table S111.** Predicted functions of ORFs in the NZ_MAIS01000009.1 containing WP_065290801.1

| gene | aa [a] | putative function | Protein homologue | %identity/ %similarity |
|---|---|---|---|---|
| WP_065290784.1 | 282 | dTDP-4-dehydrorhamnose reductase | kirromycin [CAN89624.1] | 27/40 |
| WP_065290785.1 | 406 | glycosyltransferase | SorF [ADN68481.1] | 34/49 |
| WP_065290786.1 | 301 | alpha/beta hydrolase | / | / |
| WP_065290787.1 | 1888 | AT-less type I PKS | OocN [AFX60336.1] | 37/54 |
| WP_065290788.1 | 979 | AT-less type I PKS | Bat2 [ADD82940.1] | 40/57 |
| WP_065290789.1 | 3002 | AT-less type I PKS | MisC [AKQ22699.1] | 43/60 |
| WP_083189086.1 | 1001 | AT-less type I PKS | ChiB [AAY89049.1] | 42/60 |
| WP_065290791.1 | 1724 | AT-less type I PKS | MisD [AKQ22698.1] | 37/54 |
| WP_083189087.1 | 2396 | AT-less type I PKS | MisD [AKQ22698.1] | 38/54 |
| WP_065290793.1 | 2822 | AT-less type I PKS | DszB [AAY32965.1] | 35/52 |
| WP_065290794.1 | 2730 | AT-less type I PKS | PedF [AAS47564.1] | 36/54 |
| WP_065290795.1 | 1858 | AT-less type I PKS | ChiE [AAY89052.1] | 39/54 |
| WP_065290796.1 | 2547 | AT-less type I PKS | ChiF [AAY89053.1] | 43/58 |
| WP_065290797.1 | 1761 | AT-less type I PKS | MisF [AKQ22696.1] | 39/51 |
| WP_065290798.1 | 480 | AT-less type I PKS | MisC [AKQ22699.1] | 55/71 |
| WP_065290799.1 | 1562 | AT-less type I PKS | MxnK [AGS77291.1] | 40/56 |
| WP_065290800.1 | 268 | enoyl-CoA hydratase | MxnF [AGS77286.1] | 38/58 |
| WP_065290801.1 | 410 | HMG-CoA synthase | MxnE [AGS77285.1] | 56/72 |
| WP_065290802.1 | 422 | beta-ketoacyl synthase | PedM [AAW33972.1] | 49/65 |
| WP_065290803.1 | 79 | acyl carrier protein | ElaE [AEC04351.1] | 47/68 |
| ORF1 | 315 | acyltransferase | BaeD [CAG23951.1] | 40/60 |
| WP_065290805.1 | 225 | MBL fold metallo-hydrolase | BaeB [CAG23949.2] | 48/63 |
| WP_083189088.1 | 265 | PPTase | Mis12 [AKQ22701.1] | 33/52 |
| WP_083189089.1 | 189 | antiterminator LoaP | / | / |

[a] Number of amino acids

**Table S112.** Predicted functions of ORFs in the NZ_MAUJ01000012.1 containing WP_065792709.1

| gene | aa [a] | putative function | Protein homologue | %identity/ %similarity |
|---|---|---|---|---|
| WP_065792699.1 | 400 | hypothetical protein | | |
| WP_081310686.1 | 362 | hypothetical protein | thiomarinol [CBK62711.1] | 54/72 |
| WP_065792701.1 | 1079 | hypothetical protein | SorM [ADN68497.1] | 25/42 |
| WP_065792702.1 | 321 | acyltransferase | BryP [ABM63531.1] | 38/52 |
| WP_065792703.1 | 385 | S-malonyltransferase | TstO [AGN11889.1] | 53/69 |
| WP_065792704.1 | 416 | beta-ketoacyl synthase | bonF [AFN27478.1] | 55/71 |
| WP_065792705.1 | 80 | acyl carrier protein | TstM [AGN11887.1] | 56/77 |
| WP_065792706.1 | 223 | MBL fold metallo-hydrolase | BaeB [CAG23949.2] | 39/56 |
| WP_065792707.1 | 249 | enoyl-CoA hydratase | BatE [ADD82946.1] | 56/72 |
| WP_065792708.1 | 271 | enoyl-CoA hydratase | BaeH [CAG23955.1] | 51/69 |
| WP_065792709.1 | 419 | HMG-CoA synthase | BonG [AFN27479.1] | 70/81 |
| WP_081310687.1 | 477 | 2-nitropropane dioxygenase | BatK [ADD82952.1] | 55/71 |
| WP_081310691.1 | 396 | hypothetical protein | CalB [BAP05590.1] | 28/46 |
| ORF1 | 4613 | AT-less type I PKS | SorH [ADN68483.1] | 32/50 |
| ORF2 | 4702 | AT-less type I PKS | SorD [ADN68479.1] | 36/51 |
| ORF3 | 6758 | AT-less type I PKS | SorH [ADN68483.1] | 42/58 |
| ORF4 | 1569 | AT-less type I PKS | PedI [AAR19304.1] | 43/59 |

[a] Number of amino acids

**Table S113.** Predicted functions of ORFs in the NZ_LYOY01000076.1 containing WP_067167622.1

| gene | aa [a] | putative function | Protein homologue | %identity/ %similarity |
|---|---|---|---|---|
| WP_067167614.1 | 2486 | AT-less type I PKS | PedH [AAS47562.1] | 42/57 |
| WP_067167615.1 | 2523 | AT-less type I PKS | SgvD4 [AGN74885.1] | 34/46 |
| WP_067167617.1 | 3060 | AT-less type I PKS | SnbDE [CBW45647.1] | 34/47 |
| WP_067167619.1 | 83 | acyl carrier protein | CalX [BAP05574.1] | 50/69 |
| WP_079126532.1 | 402 | beta-ketoacyl synthase | TstN [AGN11888.1] | 62/72 |
| WP_067167622.1 | 418 | HMG-CoA synthase | BaeG [CAG23954.2] | 67/81 |

[a] Number of amino acids

**Table S114.** Predicted functions of ORFs in the NZ_LOJP01000001.1 containing WP_067499190.1

| gene | aa [a] | putative function | Protein homologue | %identity/ %similarity |
|---|---|---|---|---|
| WP_067499179.1 | 323 | dTDP-glucose 4,6-dehydratase | marinomycin [BAG50456.1] | 46/59 |
| WP_067499180.1 | 125 | DUF4180 domain-containing protein | / | / |
| WP_082771995.1 | 426 | cytochrome P450 | ElaG [AEC04353.1] | 43/60 |
| WP_067499182.1 | 288 | SDR family oxidoreductase | WP_067499182.1 | 40/55 |
| WP_067499183.1 | 316 | acyltransferase | SorO [ADN68489.1] | 37/54 |
| WP_067499184.1 | 407 | beta-ketoacyl synthase | NspH [ADA69244.1] | 58/75 |
| WP_067499185.1 | 81 | acyl carrier protein | ElaE [AEC04351.1] | 59/78 |
| WP_067499187.1 | 252 | enoyl-CoA hydratase | BaeI [CAG23956.1] | 61/75 |
| WP_067499188.1 | 263 | enoyl-CoA hydratase | BaeH [CAG23955.1] | 51/66 |
| WP_067499190.1 | 419 | HMG-CoA synthase | BatC [ADD82944.1] | 66/81 |
| WP_082771996.1 | 495 | S-malonyltransferase | TstO [AGN11889.1] | 53/61 |
| WP_067499193.1 | 456 | flavin-dependent nitroreductase | OocU [AFX60343.1] | 5269 |
| WP_067499195.1 | 2920 | AT-less type I PKS | MisF [AKQ22696.1] | 37/54 |
| ORF1 | 4096 | AT-less type I PKS | SorB [ADN68477.1] | 43/56 |
| ORF2 | 549 | AT-less type I PKS | BasE [ERM18797.1] | 54/70 |
| WP_067499200.1 | 503 | TldD/PmbA family protein | / | / |
| WP_067499202.1 | 288 | homocysteine S-methyltransferase | / | / |

[a] Number of amino acids

**Table S115.** Predicted functions of ORFs in the NZ_PSWT01000067.1 containing WP_068251875.1

| gene | aa [a] | putative function | Protein homologue | %identity/ %similarity |
|---|---|---|---|---|
| WP_084415807.1 | 90 | hypothetical protein | / | / |
| WP_084415806.1 | 177 | IS3 family transposase | / | / |
| WP_068251884.1 | 82 | LysR family transcriptional regulator | / | / |
| WP_068251880.1 | 59 | hypothetical protein | / | / |
| WP_084415805.1 | 638 | S-malonyltransferase | BryP [ABM63531.1] | 36/58 |
| WP_068251877.1 | 730 | flavin-dependent nitroreductase | OocU [AFX60343.1] | 46/63 |
| WP_068251875.1 | 418 | HMG-CoA synthase | DifN [CAG23985.1] | 59/71 |
| WP_068251872.1 | 428 | beta-ketoacyl synthase | BryQ [ABM63532.1] | 49/68 |
| WP_068251870.1 | 87 | acyl carrier protein | BatA [ADD82942.1] | 46/68 |
| WP_068251867.1 | 3071 | NRPS | SgvD4 [AGN74885.1] | 35/46 |
| WP_068251865.1 | 2557 | hybrid NRPS/PKS | kirromycin [CAN89656.1] | 33/44 |
| WP_104261928.1 | 4431 | AT-less type I PKS | SorB [ADN68477.1] | 43/55 |
| WP_068251859.1 | 6430 | hybrid NRPS/PKS | Bat2 [ADD82940.1] | 35/49 |
| WP_084415803.1 | 3740 | AT-less type I PKS | SorH [ADN68483.1] | 38/51 |
| WP_068251857.1 | 255 | thioesterase | LnmN [AAN85527.1] | 44/55 |
| WP_068256981.1 | 217 | PPTase | LtmL [ACY01405.1] | 53/65 |
| WP_068251856.1 | 280 | Ser/Thr protein phosphatase | ThaC [ABC35295.1] | 28/40 |

[a] Number of amino acids

**Table S116.** Predicted functions of ORFs in the NZ_MCGG01000009.1 containing WP_069956867.1

| gene | aa [a] | putative function | Protein homologue | %identity/ %similarity |
|---|---|---|---|---|
| ORF1 | 6560 | AT-less type I PKS | Ta1 [ABF85931.1] | 37/51 |
| ORF2 | 442 | AT-less type I PKS | dipN [AGS06825.1] | 60/76 |
| ORF3 | 6898 | AT-less type I PKS | Dor5 [ACY01390.1] | 44/58 |
| ORF4 | 754 | AT/Ox | CalY [BAP05573.1] | 49/68 |
| WP_069956864.1 | 258 | PPTase | TmlN [CBK62709.1] | 35/55 |
| WP_069956865.1 | 274 | Ser/Thr protein phosphatase | ThaC [ABC35295.1] | 29/45 |
| WP_084004943.1 | 369 | linear amide C-N hydrolase | / | / |
| WP_069956866.1 | 415 | cytochrome P450 | BaeS [CAG23962.1] | 32/50 |
| WP_069956867.1 | 419 | HMG-CoA synthase | CylF [ARU81120.1] | 58/76 |
| WP_069956930.1 | 81 | acyl carrier protein | CalX [BAP05574.1] | 49/73 |
| WP_069956868.1 | 414 | beta-ketoacyl synthase | ElaF [AEC04352.1] | 48/63 |
| WP_069956869.1 | 253 | enoyl-CoA hydratase | BaeH [CAG23955.1] | 52/66 |
| WP_069956870.1 | 660 | asparagine synthase | ChxD [AFO59865.1] | 57/72 |
| WP_069956871.1 | 81 | acyl carrier protein | ChxC [AFO59864.1] | 51/71 |
| WP_069956872.1 | 355 | acyltransferase | BaeD [CAG23951.1] | 35/53 |
| WP_084004944.1 | 270 | cupin domain-containing protein | CalA [BAP05589.1] | 32/50 |
| WP_069956875.1 | 410 | 6-phosphofructokinase | / | / |

[a] Number of amino acids

**Table S117.** Predicted functions of ORFs in the NZ_LJGU01000152.1 containing WP_070198817.1

| gene | aa [a] | putative function | Protein homologue | %identity/ %similarity |
|---|---|---|---|---|
| WP_079166934.1 | 371 | NAD-dependent epimerase | / | / |
| WP_070198811.1 | 302 | methyltransferase | / | / |
| WP_070198812.1 | 422 | glycosyltransferase | / | / |
| WP_107402085.1 | 406 | cytochrome P450 | ElaG [AEC04353.1] | 43/60 |
| WP_070198813.1 | 319 | acyltransferase | BaeD [CAG23951.1] | 36/53 |
| WP_070198814.1 | 414 | beta-ketoacyl synthase | NspH [ADA69244.1] | 59/76 |
| WP_070198815.1 | 81 | acyl carrier protein | TstM [AGN11887.1] | 64/77 |
| WP_107402086.1 | 250 | enoyl-CoA hydratase | BatE [ADD82946.1] | 63/80 |
| WP_070198816.1 | 261 | enoyl-CoA hydratase | BaeH [CAG23955.1] | 52/69 |
| WP_070198817.1 | 420 | HMG-CoA synthase | BatC [ADD82944.1] | 65/80 |
| WP_079166935.1 | 443 | S-malonyltransferase | TstO [AGN11889.1] | 54/64 |
| WP_070198818.1 | 3071 | AT-less type I PKS | MisF [AKQ22696.1] | 39/57 |
| WP_107402087.1 | 6128 | AT-less type I PKS | BaeL [CAG23958.2] | 41/56 |
| WP_070198820.1 | 2902 | AT-less type I PKS | BaeL [CAG23958.2] | 48/63 |

[a] Number of amino acids

**Table S118.** Predicted functions of ORFs in the NZ_FPIS01000005.1 containing WP_072324073.1

| gene | aa [a] | putative function | Protein homologue | %identity/ %similarity |
|---|---|---|---|---|
| WP_083527570.1 | 254 | ABC transporter ATP-binding protein | LnmR [AAN85531.1] | 33/46 |
| WP_072324066.1 | 415 | HlyD family efflux transporter | / | / |
| WP_072324067.1 | 1129 | NRPS | CalI [BAP05597.1] | 37/54 |
| WP_072324069.1 | 240 | hypothetical protein | / | / |
| WP_083527571.1 | 847 | FtsX-like permease family | / | / |
| WP_072324072.1 | 376 | S-malonyltransferase | fr9O [AIC32701.1] | 53/66 |
| WP_072324078.1 | 84 | acyl carrier protein | TstM [AGN11887.1] | 55/77 |
| WP_072324073.1 | 418 | HMG-CoA synthase | BatC [ADD82944.1] | 67/78 |
| WP_083527572.1 | 238 | MBL fold metallo-hydrolase | BaeB [CAG23949.2] | 41/58 |
| WP_072324076.1 | 615 | asparagine synthase | BasB [ERM18801.1] | 42/59 |
| ORF1 | 2997 | AT-less type I PKS | MisF [AKQ22696.1] | 33/48 |

[a] Number of amino acids

**Table S119.** Predicted functions of ORFs in the NZ_BBYF01000002.1 containing WP_072728425.1

| gene | aa [a] | putative function | Protein homologue | %identity/ %similarity |
|---|---|---|---|---|
| WP_083545044.1 | 211 | polyketide synthase | ElaO [AEC04361.1] | 71/84 |
| WP_072728420.1 | 3478 | AT-less type I PKS | DszB [AAY32965.1] | 38/55 |
| WP_083545045.1 | 493 | AT-less type I PKS | DszA [AAY32964.1] | 33/53 |
| WP_072728423.1 | 4353 | AT-less type I PKS | DszB [AAY32965.1] | 35/52 |
| WP_083545009.1 | 4050 | AT-less type I PKS | DszB [AAY32965.1] | 40/56 |
| WP_072728425.1 | 419 | HMG-CoA synthase | BaeG [CAG23954.2] | 78/86 |
| WP_072728426.1 | 254 | enoyl-CoA hydratase | BaeH [CAG23955.1] | 57/72 |
| WP_083545012.1 | 303 | Ser/Thr protein phosphatase | ThaC [ABC35295.1] | 42/62 |
| WP_072728428.1 | 259 | PPTase | Mis12 [AKQ22701.1] | 37/53 |
| WP_065291672.1 | 264 | thymidylate synthase | / | / |

[a] Number of amino acids

**Table S120.** Predicted functions of ORFs in the NZ_FNWO01000008.1 containing WP_074768427.1

| gene | aa [a] | putative function | Protein homologue | %identity/ %similarity |
|---|---|---|---|---|
| WP_083386743.1 | 342 | sulfate exporter family transporter | / | / |
| WP_074768405.1 | 229 | class I SAM-dependent methyltransferase | KirM [CAN89643.1] | 38/54 |
| WP_074768407.1 | 764 | AT/Ox | CalY [BAP05573.1] | 51/68 |
| ORF1 | 6482 | AT-less type I PKS | ChxE [AFO59866.1] | 42/53 |
| WP_074768411.1 | 433 | monooxygenase | PedG [AAS47561.1] | 62/77 |
| ORF2 | 6327 | AT-less type I PKS | Ta1 [ABF85931.1] | 38/52 |
| WP_083386750.1 | 287 | hypothetical protein | JamP [AAS98787.1] | 31/47 |
| WP_074768415.1 | 316 | acyltransferase | BaeD [CAG23951.1] | 39/56 |
| WP_074768417.1 | 82 | acyl carrier protein | SmdG [CCC21121.1] | 48/67 |
| WP_074768419.1 | 656 | asparagine synthase | SorQ [ADN68491.1] | 55/70 |
| WP_074768421.1 | 251 | enoyl-CoA hydratase | BaeH [CAG23955.1] | 47/63 |
| WP_074768423.1 | 405 | beta-ketoacyl synthase | NspH [ADA69244.1] | 46/66 |
| WP_074768425.1 | 82 | acyl carrier protein | CalX [BAP05574.1] | 45/67 |
| WP_074768427.1 | 418 | HMG-CoA synthase | ThaK [ABC34601.1] | 62/78 |
| WP_083386744.1 | 416 | cytochrome P450 | BaeS [CAG23962.1] | 34/49 |
| WP_074768431.1 | 272 | FkbM family methyltransferase | CalA [BAP05589.1] | 34/48 |
| WP_074768433.1 | 247 | PPTase | BatI [ADD82950.1] | 35/51 |
| WP_074768435.1 | 686 | heavy metal translocating P-type ATPase | PsyN [ADA82595.1] | 42/63 |
| WP_083386745.1 | 245 | DNA-binding response regulator | smdD [CCC21118.1] | 3758 |
| WP_083386746.1 | 689 | methyl-accepting chemotaxis protein | / | / |

[a] Number of amino acids

**Table S121.** Predicted functions of ORFs in the NZ_AP014940.1 containing WP_074868379.1

| gene | aa [a] | putative function | Protein homologue | %identity/ %similarity |
|---|---|---|---|---|
| WP_096378561.1 | 164 | hypothetical protein | / | / |
| WP_074868352.1 | 167 | UpxY family transcription antiterminator | TaA [ABF91060.1] | 31/48 |
| WP_096378564.1 | 317 | acyltransferase | BatH [ADD82949.1] | 53/66 |
| WP_096378566.1 | 281 | PPTase | BatI [ADD82950.1] | 50/66 |
| WP_096378569.1 | 253 | Ser/Thr protein phosphatase | ThaC [ABC35295.1] | 46/62 |
| WP_096378572.1 | 373 | S-malonyltransferase | TstO [AGN11889.1] | 59/71 |
| WP_096378575.1 | 411 | beta-ketoacyl synthase | TstN [AGN11888.1] | 67/79 |
| WP_074868371.1 | 84 | acyl carrier protein | TstM [AGN11887.1] | 59/71 |
| WP_096378578.1 | 260 | enoyl-CoA hydratase | BatE [ADD82946.1] | 67/79 |
| WP_003806645.1 | 260 | enoyl-CoA hydratase | BaeH [CAG23955.1] | 55/69 |
| WP_074868379.1 | 418 | HMG-CoA synthase | fr9K [AIC32697.1] | 73/85 |
| WP_096378581.1 | 238 | MBL fold metallo-hydrolase | BaeB [CAG23949.2] | 38/54 |
| WP_096378584.1 | 4522 | AT-less type I PKS | SorH [ADN68483.1] | 38/52 |
| WP_096378587.1 | 4796 | AT-less type I PKS | MisD [AKQ22698.1] | 39/56 |
| WP_096378590.1 | 5483 | AT-less type I PKS | BaeL [CAG23958.2] | 40/56 |
| WP_096378593.1 | 88 | acyl carrier protein | MacpD [AAM12933.1] | 52/62 |
| WP_083382655.1 | 243 | SDR family oxidoreductase | DifE [CAG23976.1] | 50/71 |
| WP_074868403.1 | 479 | long-chain fatty acid--CoA ligase | DifD [CAJ57407.1] | 46/66 |
| WP_096378596.1 | 257 | hypothetical protein | / | / |
| WP_096378599.1 | 259 | hypothetical protein | / | / |

[a] Number of amino acids

**Table S122.** Predicted functions of ORFs in the NZ_MRUG01000025.1 containing WP_075185089.1

| gene | aa [a] | putative function | Protein homologue | %identity/ %similarity |
|---|---|---|---|---|
| WP_075185087.1 | 383 | IS4 family transposase | / | / |
| WP_083607969.1 | 217 | hypothetical protein | BaeB [CAG23949.2] | 35/55 |
| WP_075185089.1 | 418 | HMG-CoA synthase | BatC [ADD82944.1] | 68/81 |
| WP_075185090.1 | 261 | enoyl-CoA hydratase | BaeH [CAG23955.1] | 55/71 |
| WP_075185091.1 | 249 | enoyl-CoA hydratase | BatE [ADD82946.1] | 57/78 |
| WP_075185092.1 | 79 | acyl carrier protein | ElaE [AEC04351.1] | 48/75 |
| WP_075185093.1 | 418 | beta-ketoacyl synthase | TstN [AGN11888.1] | 59/73 |
| WP_075185094.1 | 371 | S-malonyltransferase | TstO [AGN11889.1] | 52/67 |
| WP_075185095.1 | 323 | acyltransferase | SorO [ADN68489.1] | 38/59 |
| WP_075185096.1 | 1181 | hypothetical protein | SorM [ADN68497.1] | 27/44 |
| WP_075185097.1 | 235 | PPTase | LtmL [ACY01405.1] | 33/47 |
| WP_075185099.1 | 180 | hypothetical protein | / | / |
| WP_075185100.1 | 141 | hypothetical protein | / | / |
| WP_075185101.1 | 207 | glutathione S-transferase | / | / |
| WP_075185102.1 | 366 | S-malonyltransferase | TstO [AGN11889.1] | 51/66 |
| WP_075185103.1 | 402 | hypothetical protein | / | / |
| WP_075185104.1 | 1126 | hypothetical protein | DifL [CAG23983.1] | 31/49 |
| ORF1 | 4096 | AT-less type I PKS | BasG [ERM18799.1] | 34/52 |
| ORF2 | 6766 | AT-less type I PKS | CalC [BAP05591.1] | 45/60 |
| WP_075185107.1 | 857 | hypothetical protein | / | / |

[a] Number of amino acids

**Table S123.** Predicted functions of ORFs in the NZ_MKZS01000001.1 containing WP_075900452.1

| gene | aa [a] | putative function | Protein homologue | %identity/ %similarity |
|---|---|---|---|---|
| WP_075906166.1 | 564 | response regulator | / | / |
| WP_075900442.1 | 530 | Na+/H+ antiporter | / | / |
| WP_075900444.1 | 97 | Dabb family protein | / | / |
| WP_075900446.1 | 1674 | AT-less type I PKS | JamJ [AAS98781.1] | 62/76 |
| WP_075900448.1 | 1176 | NRPS | ChiD [AAY89051.1] | 34/53 |
| WP_081431224.1 | 2942 | AT-less type I PKS | JamJ [AAS98781.1] | 67/79 |
| WP_081431225.1 | 258 | enoyl-CoA hydratase | CylG [ARU81121.1] | 84/91 |
| WP_075900452.1 | 419 | HMG-CoA synthase | CylF [ARU81120.1] | 84/91 |
| WP_075900454.1 | 407 | beta-ketoacyl synthase | CylE [ARU81119.1] | 76/87 |
| WP_075900456.1 | 83 | acyl carrier protein | JamF [AAS98799.1] | 76//89 |
| WP_075900458.1 | 2828 | AT-less type I PKS | JamK [AAS98782.1] | 57/72 |
| WP_075900460.1 | 1138 | methyltransferase | BryX [ABM63529.1] | 41//58 |
| WP_075900462.1 | 84 | hypothetical protein | BryX [ABM63529.1] | 52/72 |
| WP_075900464.1 | 60 | hypothetical protein | / | / |
| WP_075900466.1 | 84 | hypothetical protein | / | / |
| WP_075900468.1 | 66 | hypothetical protein | / | / |

[a] Number of amino acids

**Table S124.** Predicted functions of ORFs in the NZ_MVBD01000017.1 containing WP_077412983.1

| gene | aa [a] | putative function | Protein homologue | %identity/ %similarity |
|---|---|---|---|---|
| WP_077412974.1 | 415 | HlyD family efflux transporter | / | / |
| WP_077412975.1 | 1115 | hypothetical protein | SorM [ADN68497.1] | 29/48 |
| WP_077412976.1 | 274 | PPTase | TmlN [CBK62709.1] | 35/60 |
| WP_077412977.1 | 327 | acyltransferase | MxnM [AGS77293.1] | 36/60 |
| WP_077412978.1 | 395 | S-malonyltransferas | TstO [AGN11889.1] | 48/63 |
| WP_077412979.1 | 414 | beta-ketoacyl synthase | NspH [ADA69244.1] | 55/72 |
| WP_077412980.1 | 79 | acyl carrier protein | NspG [ADA71314.1] | 61/77 |
| WP_077412981.1 | 264 | enoyl-CoA hydratase | BaeH [CAG23955.1] | 51/70 |
| WP_077412983.1 | 419 | HMG-CoA synthase | BatC [ADD82944.1] | 68/81 |
| WP_077412984.1 | 235 | MBL fold metallo-hydrolase | BaeB [CAG23949.2] | 42/61 |
| WP_077412992.1 | 474 | PfaD family polyunsaturated fatty acid | BaeN [CAG23960.2] | 35/52 |
| WP_077412986.1 | 5086 | AT-less type I PKS | BryB [ABM63527.1] | 32/50 |
| WP_077412987.1 | 374 | LLM class flavin-dependent oxidoreductase | OocM [AFX60335.1] | 61/76 |
| WP_077412988.1 | 2531 | AT-less type I PKS | BaeN [CAG23960.2] | 35/52 |
| WP_077412989.1 | 59 | hypothetical protein | / | / |

[a] Number of amino acids

**Table S125.** Predicted functions of ORFs in the NZ_BAMR01000013.1 containing WP_081611316.1

| gene | aa [a] | putative function | Protein homologue | %identity/ %similarity |
|---|---|---|---|---|
| WP_025897181.1 | 389 | hypothetical protein | | |
| WP_025897183.1 | 59 | hypothetical protein | | |
| WP_081786585.1 | 411 | hypothetical protein | leinamycin [AAN85512.1] | 24/41 |
| WP_081786586.1 | 142 | hypothetical protein | RhiB [CAL69889.1] | 44/61 |
| WP_081786587.1 | 135 | hypothetical protein | DszC [AAY32966.1] | 45/65 |
| WP_081786588.1 | 1348 | AT-less type I PKS | OocJ [AFX60332.1] | 51/69 |
| WP_081786589.1 | 328 | AT-less type I PKS | LglD [AIU36100.1] | 36/53 |
| WP_025897196.1 | 423 | NRPS | Ta1 [ABF85931.1] | 42/57 |
| WP_025897198.1 | 138 | AT-less type I PKS | CalI [BAP05597.1] | 25/45 |
| WP_081786590.1 | 585 | AT-less type I PKS | OzmH [ABS90470.1] | 28/56 |
| WP_025897202.1 | 2064 | AT-less type I PKS | DifI [CAJ57409.1] | 31/46 |
| WP_081786591.1 | 279 | AT-less type I PKS | SorG [ADN68482.1] | 57/72 |
| WP_025897206.1 | 144 | hypothetical protein | ChiB [AAY89049.1] | 61/75 |
| WP_025897207.1 | 1307 | AT-less type I PKS | ChiB [AAY89049.1] | 53/71 |
| WP_081786592.1 | 2015 | AT-less type I PKS | OocJ [AFX60332.1] | 37/53 |
| WP_081786593.1 | 1067 | AT-less type I PKS | OzmN [ABS90475.1] | 45/57 |
| WP_025897219.1 | 415 | AT-less type I PKS | SgvE1 [AGN74892.1] | 31/46 |
| WP_025897221.1 | 1595 | AT-less type I PKS | MxnK [AGS77291.1] | 44/58 |
| WP_020398753.1 | 121 | hypothetical protein | / | / |
| WP_020398754.1 | 258 | enoyl-CoA hydratase | CylG [ARU81121.1] | 32/51 |
| WP_081611316.1 | 439 | HMG-CoA synthase | PyxM [ASA76639.1] | 56/68 |
| WP_081786594.1 | 90 | hypothetical protein | TmlG [CBK62726.1] | 40/60 |
| WP_081786595.1 | 361 | beta-ketoacyl synthase | DipR [AGS06821.1] | 27/46 |
| WP_020398757.1 | 778 | AT/Ox | DifA [CAG23974.1] | 46/62 |
| WP_020398758.1 | 281 | PPTase | Mis12 [AKQ22701.1] | 33/46 |
| WP_020398759.1 | 715 | ABC transporter ATP-binding protein | CalU [BAP05577.1] | 37/57 |
| WP_081611318.1 | 94 | hypothetical protein | / | / |
| WP_020398761.1 | 125 | hypothetical protein | / | / |

[a] Number of amino acids

**Table S126.** Predicted functions of ORFs in the NZ_FXAM01000002.1 containing WP_085216372.1

| gene | aa [a] | putative function | Protein homologue | %identity/ %similarity |
|---|---|---|---|---|
| WP_085216362.1 | 521 | efflux transporter outer membrane subunit | / | / |
| WP_085216363.1 | 106 | hypothetical protein | / | / |
| WP_085216364.1 | 87 | hypothetical protein | / | / |
| WP_085216365.1 | 291 | S-malonyltransferase | RhiG [CAL69887.1] | 48/69 |
| WP_085216366.1 | 319 | acyltransferase | BaeD [CAG23951.1] | 35/51 |
| WP_085216367.1 | 267 | PPTase | MupN [AAM12928.1] | 32/49 |
| WP_085216368.1 | 598 | AarF/ABC1/UbiB kinase family protein | Rhizopodin [CCA89332.1] | 36/54 |
| WP_085216369.1 | 450 | PfaD family polyunsaturated fatty acid | DipL [AGS06827.1] | 44/64 |
| WP_085216370.1 | 481 | cytochrome P450 | CorO [ADI59538.1] | 38/54 |
| WP_085216371.1 | 428 | beta-ketoacyl synthase | CorD [ADI59526.1] | 35/46 |
| WP_085216372.1 | 409 | HMG-CoA synthase | CylF [ARU81120.1] | 46/64 |
| WP_085216373.1 | 262 | enoyl-CoA hydratase | MxnF [AGS77286.1] | 41/53 |
| WP_085216374.1 | 248 | enoyl-CoA hydratase | BatE [ADD82946.1] | 53/69 |
| WP_085216375.1 | 450 | hypothetical protein | / | / |
| WP_085216376.1 | 80 | hypothetical protein | / | / |
| WP_085216377.1 | 1775 | PAS domain S-box protein | / | / |

[a] Number of amino acids

**Table S127.** Predicted functions of ORFs in the NZ_NEHY01000040.1 containing WP_085653298.1

| gene | aa [a] | putative function | Protein homologue | %identity/ %similarity |
|---|---|---|---|---|
| WP_085600740.1 | 449 | AT-less type I PKS | BonH [AFN27484.1] | 69/81 |
| WP_085653293.1 | 1945 | AT-less type I PKS | CalG [BAP05595.1] | 36/50 |
| WP_085653295.1 | 1272 | AT-less type I PKS | bonA [AFN27480.1] | 53/64 |
| WP_085653298.1 | 420 | HMG-CoA synthase | BonG [AFN27479.1] | 77/86 |
| WP_085653303.1 | 410 | beta-ketoacyl synthase | BonF [AFN27478.1] | 69/78 |
| WP_085653300.1 | 367 | S-malonyltransferase | BonK [AFN27477.1] | 58/71 |

[a] Number of amino acids

**Table S128.** Predicted functions of ORFs in the NZ_CP021780.1 containing WP_087913544.1

| gene | aa [a] | putative function | Protein homologue | %identity/ %similarity |
|---|---|---|---|---|
| WP_087913485.1 | 412 | group II intron reverse transcriptase | / | / |
| WP_087913532.1 | 190 | antiterminator LoaP | / | / |
| WP_087913533.1 | 257 | PPTase | Mis12 [AKQ22701.1] | 37/55 |
| WP_087913534.1 | 790 | S-malonyltransferase | BaeE [CAG23952.1] | 58/74 |
| WP_087913535.1 | 2663 | AT-less type I PKS | BaeM [CAG23959.2] | 34/51 |
| WP_087913536.1 | 3826 | AT-less type I PKS | BaeN [CAG23960.2] | 41/58 |
| WP_087913537.1 | 4812 | AT-less type I PKS | RhiB [CAL69889.1] | 41/57 |
| WP_087913538.1 | 1917 | AT-less type I PKS | RhiA [CAL69888.1] | 39/54 |
| WP_087913539.1 | 5233 | NRPS | CalC [BAP05591.1] | 43/59 |
| WP_087913540.1 | 1209 | AT-less type I PKS | JamP [AAS98787.1] | 52/69 |
| WP_087913541.1 | 474 | MATE family efflux transporter | / | / |
| WP_087913542.1 | 82 | acyl carrier protein | AcpK [CAG23953.1] | 57/75 |
| WP_087913543.1 | 412 | beta-ketoacyl synthase | CalW [BAP05575.1] | 59/78 |
| WP_087913544.1 | 419 | HMG-CoA synthase | BaeG [CAG23954.2] | 75/84 |
| WP_087913545.1 | 254 | enoyl-CoA hydratase | BaeH [CAG23955.1] | 55/69 |
| WP_087913546.1 | 252 | enoyl-CoA hydratase | BatE [ADD82946.1] | 68/83 |
| WP_087913547.1 | 324 | acyltransferase | BaeD [CAG23951.1] | 44/62 |
| WP_087913548.1 | 129 | hypothetical protein | / | / |
| WP_087913549.1 | 250 | hypothetical protein | / | / |

[a] Number of amino acids

**Table S129.** Predicted functions of ORFs in the NZ_FYEP01000021.1 containing WP_088833751.1

| gene | aa [a] | putative function | Protein homologue | %identity/ %similarity |
|---|---|---|---|---|
| WP_088833745.1 | 196 | hypothetical protein | / | / |
| WP_088833746.1 | 140 | hypothetical protein | / | / |
| WP_036691017.1 | 141 | 3-hydroxyacyl dehydratase | / | / |
| WP_088833747.1 | 2592 | AT-less type I PKS | DifI [CAJ57409.1] | 39/54 |
| WP_088833748.1 | 162 | hypothetical protein | MisE [AKQ22697.1] | 54/79 |
| WP_088833749.1 | 3845 | AT-less type I PKS | BryB [ABM63527.1] | 42/58 |
| WP_036691012.1 | 249 | enoyl-CoA hydratase | BatE [ADD82946.1] | 67/84 |
| WP_088833750.1 | 257 | enoyl-CoA hydratase | BaeH [CAG23955.1] | 62/76 |
| WP_088833751.1 | 419 | HMG-CoA synthase | BatC [ADD82944.1] | 76/86 |
| WP_088833752.1 | 408 | beta-ketoacyl synthase | CalW [BAP05575.1] | 63/79 |
| WP_088833753.1 | 82 | acyl carrier protein | NspG [ADA71314.1] | 58//77 |
| WP_088833754.1 | 5451 | AT-less type I PKS | SorB [ADN68477.1] | 48/62 |
| WP_088833755.1 | 2765 | AT-less type I PKS | MmpD [AAM12913.1] | 41/58 |
| WP_088833756.1 | 5176 | AT-less type I PKS | DifF [CAG23977.1] | 44//61 |
| WP_088833757.1 | 3838 | AT-less type I PKS | MisF [AKQ22696.1] | 36//53 |
| WP_088833758.1 | 1133 | S-malonyltransferase | BasH [ERM18800.1] | 50/69 |
| WP_088833759.1 | 226 | MBL fold metallo-hydrolase | BaeB [CAG23949.2] | 54/68 |
| WP_088833760.1 | 253 | PPTase | Mis12 [AKQ22701.1] | 38/56 |
| WP_036690984.1 | 184 | antiterminator LoaP | / | / |

[a] Number of amino acids

**Table S130.** Predicted functions of ORFs in the NZ_NAPR01000012.1 containing WP_089156252.1

| gene | aa [a] | putative function | Protein homologue | %identity/ %similarity |
|---|---|---|---|---|
| WP_089156243.1 | 364 | LLM class flavin-dependent oxidoreductase | virN [BAF50713.1] | 77/85 |
| WP_089156244.1 | 385 | N-methyl-L-tryptophan oxidase | virM [BAF50714.1] | 74/84 |
| WP_089156245.1 | 266 | thioesterase | VirJ [BAF50718.1] | 67/76 |
| WP_089156246.1 | 296 | S-malonyltransferase | VirI [BAF50719.1] | 75/83 |
| WP_089156247.1 | 2013 | NRPS | SgvE4 [AGN74895.1] | 47/55 |
| WP_089156248.1 | 465 | hypothetical protein | VirH [BAF50720.1] | 62/69 |
| WP_089156249.1 | 2000 | AT-less type I PKS | SnaE3 [CBW45741.1] | 62/69 |
| WP_089156250.1 | 261 | enoyl-CoA hydratase | VirE [BAF50723.1] | 79/86 |
| WP_089156251.1 | 259 | enoyl-CoA hydratase | VirD [BAF50724.1] | 70/75 |
| WP_089156252.1 | 416 | HMG-CoA synthase | VirC [BAF50725.1] | 85/91 |
| WP_089156253.1 | 450 | beta-ketoacyl synthase | VirB [BAF50726.1] | 68/74 |
| WP_089156254.1 | 80 | acyl carrier protein | SnaG [CBW45747.1] | 59/77 |
| WP_089156255.1 | 1223 | SDR family oxidoreductase | VirA [BAF50727.1] | 62/70 |
| WP_089156256.1 | 131 | hypothetical protein | VirA [BAF50727.1] | 68/78 |
| WP_089156257.1 | 1659 | NRPS | VirA [BAF50727.1] | 60/68 |
| WP_089156398.1 | 635 | alpha-keto acid dehydrogenase | SnaF [CBW45750.1] | 73/81 |
| WP_089156258.1 | 402 | cytochrome P450 | SnbF [CBW45756.1] | 64//77 |
| WP_089156259.1 | 252 | tetratricopeptide repeat protein | / | / |
| WP_089156260.1 | 62 | tetratricopeptide repeat protein | / | / |

[a] Number of amino acids

**Table S131.** Predicted functions of ORFs in the NZ_FOMK01000001.1 containing WP_090100317.1

| gene | aa [a] | putative function | Protein homologue | %identity/ %similarity |
|---|---|---|---|---|
| WP_090100308.1 | 771 | TonB-dependent receptor | / | / |
| WP_090100310.1 | 83 | ATP-binding cassette domain | / | / |
| WP_090100312.1 | 335 | acyltransferase | BaeD [CAG23951.1] | 38/54 |
| WP_090100314.1 | 248 | enoyl-CoA hydratase | BatE [ADD82946.1] | 54/76 |
| WP_090100315.1 | 265 | enoyl-CoA hydratase | CylG [ARU81121.1] | 54/67 |
| WP_090100317.1 | 414 | HMG-CoA synthase | CylF [ARU81120.1] | 64/78 |
| WP_090100319.1 | 415 | beta-ketoacyl synthase | CylE [ARU81119.1] | 44/61 |
| WP_090100321.1 | 80 | acyl carrier protein | NspG [ADA71314.1] | 38/65 |
| WP_090100323.1 | 742 | AT/Ox | DifA [CAG23974.1] | 53/72 |
| WP_090100325.1 | 7288 | NRPS | TaO [ABF92489.1] | 38/55 |
| WP_090100327.1 | 5911 | AT-less type I PKS | BaeL [CAG23958.2] | 40/57 |
| WP_090100329.1 | 2557 | AT-less type I PKS | BaeN [CAG23960.2] | 39/55 |
| WP_090100331.1 | 353 | GHKL domain-containing protein | / | / |
| WP_090100333.1 | 240 | DNA-binding response regulator | / | / |

[a] Number of amino acids

**Table S132.** Predicted functions of ORFs in the NZ_BARF01000050.1 containing WP_090739057.1

| gene | aaa | putative function | Protein homologue | %identity/ %similarity |
|---|---|---|---|---|
| WP_090739048.1 | 178 | hypothetical protein | / | / |
| WP_090739049.1 | 310 | cysteine synthase A | Kirromycin [CAN89659.1] | 33/52 |
| WP_090739050.1 | 226 | hypothetical protein | SgvC [AGN74907.1] | 41/56 |
| WP_090739051.1 | 1294 | AT-less type I PKS | OocR [AFX60340.1] | 56/67 |
| WP_090739052.1 | 843 | AT-less type I PKS | DifI [CAJ57409.1] | 38/54 |
| WP_090739053.1 | 2328 | AT-less type I PKS | SorB [ADN68477.1] | 48/61 |
| WP_090739054.1 | 3456 | AT-less type I PKS | SorE [ADN68480.1] | 44/58 |
| WP_090739055.1 | 249 | enoyl-CoA hydratase | BaeE [CAG23952.1] | 66/81 |
| WP_090739056.1 | 254 | enoyl-CoA hydratase | BaeH [CAG23955.1] | 56/70 |
| WP_090739057.1 | 420 | HMG-CoA synthase | BaeG [CAG23954.2] | 72/84 |
| WP_090739192.1 | 405 | beta-ketoacyl synthase | CalW [BAP05575.1] | 61/75 |
| WP_090739058.1 | 82 | acyl carrier protein | NspG [ADA71314.1] | 51/77 |
| WP_090739059.1 | 4342 | NRPS | BryB [ABM63527.1] | 44/59 |
| WP_090739060.1 | 258 | hypothetical protein | CalB [BAP05590.1] | 41/52 |
| WP_090739061.1 | 1583 | AT-less type I PKS | RhiA [CAL69888.1] | 39/53 |
| WP_090739062.1 | 1552 | AT-less type I PKS | BaeN [CAG23960.2] | 41/56 |
| WP_090739063.1 | 271 | methyltransferase domain | Dor10 [ACY01395.1] | 31/46 |
| WP_090739064.1 | 265 | alpha/beta hydrolase | / | / |
| WP_090739065.1 | 1181 | S-malonyltransferase | BasH [ERM18800.1] | 49/67 |
| WP_090739066.1 | 225 | MBL fold metallo-hydrolase | BaeB [CAG23949.2] | 50/71 |
| WP_090739067.1 | 546 | MFS transporter | SnbR [CBW45761.1] | 25/44 |
| WP_090739068.1 | 198 | transcription antiterminator | / | / |

[a] Number of amino acids

**Table S133.** Predicted functions of ORFs in the NZ_FPAA01000002.1 containing WP_091833656.1

| gene | aa [a] | putative function | Protein homologue | %identity/ %similarity |
|---|---|---|---|---|
| WP_091834437.1 | 115 | YkgJ family cysteine | / | / |
| WP_091834439.1 | 460 | MATE family efflux transporter | SorJ [ADN68485.1] | 23/48 |
| WP_091833648.1 | 610 | enoyl-CoA hydratase | BatE [ADD82946.1] | 54/73 |
| WP_091833651.1 | 249 | enoyl-CoA hydratase | BatE [ADD82946.1] | 64/78 |
| WP_091833653.1 | 255 | enoyl-CoA hydratase | BaeH [CAG23955.1] | 56/73 |
| WP_091833656.1 | 419 | HMG-CoA synthase | BatC [ADD82944.1] | 74/86 |
| WP_091833659.1 | 2761 | AT-less type I PKS | bonA [AFN27480.1] | 43/59 |
| WP_091833662.1 | 5791 | AT-less type I PKS | MisF [AKQ22696.1] | 41/59 |
| WP_091833664.1 | 1677 | AT-less type I PKS | MisF [AKQ22696.1] | 44/62 |
| WP_091833667.1 | 2328 | AT-less type I PKS | Ta1 [ABF85931.1] | 40/56 |
| WP_091833670.1 | 5877 | AT-less type I PKS | MisD [AKQ22698.1] | 41/58 |
| WP_091833673.1 | 273 | methyltransferase | Dor10 [ACY01395.1] | 31/47 |
| WP_091833676.1 | 270 | alpha/beta hydrolase | OocA [AFX60323.1] | 31/50 |
| WP_091833679.1 | 758 | AT/Ox | DifA [CAG23974.1] | 60/80 |
| WP_091833682.1 | 207 | hypothetical protein | / | / |
| WP_091833685.1 | 319 | acyltransferase | BaeD [CAG23951.1] | 43/60 |
| WP_091833688.1 | 407 | cytochrome P450 | DifM [CAG23984.1] | 37/58 |
| WP_091833691.1 | 178 | antiterminator LoaP | / | / |

[a] Number of amino acids

**Table S134.** Predicted functions of ORFs in the NZ_FNHB01000001.1 containing WP_092069005.1

| gene | aa [a] | putative function | Protein homologue | %identity/ %similarity |
|---|---|---|---|---|
| WP_092068982.1 | 344 | NAD-dependent epimerase | / | / |
| WP_092068985.1 | 461 | aminotransferase class I | / | / |
| WP_092068988.1 | 8085 | hybrid PKS/NRPS | BaeN [CAG23960.2] | 41/59 |
| WP_092068991.1 | 6050 | AT-less type I PKS | MisF [AKQ22696.1] | 44/60 |
| WP_092068994.1 | 4582 | AT-less type I PKS | BaeL [CAG23958.2] | 47/64 |
| WP_092068997.1 | 173 | antitermination protein NusG | TaA [ABF91060.1] | 27/44 |
| WP_092068999.1 | 249 | enoyl-CoA hydratase | BatE [ADD82946.1] | 68//82 |
| WP_092069002.1 | 254 | enoyl-CoA hydratase | BaeH [CAG23955.1] | 61/76 |
| WP_092069005.1 | 420 | HMG-CoA synthase | BaeG [CAG23954.2] | 65/76 |
| WP_092069008.1 | 289 | S-malonyltransferase | BaeC [CAG23950.2] | 66/78 |
| WP_092069011.1 | 312 | acyltransferase | BaeD [CAG23951.1] | 50/65 |
| WP_092069014.1 | 225 | MBL fold metallo-hydrolase | BaeB [CAG23949.2] | 50/67 |
| WP_092069017.1 | 578 | ABC transporter ATP-binding protein | CalU [BAP05577.1] | 28/48 |
| WP_092069020.1 | 596 | ABC transporter ATP-binding protein | CalU [BAP05577.1] | 28/48 |
| WP_092069024.1 | 318 | AraC family transcriptional regulator | / | / |

[a] Number of amino acids

**Table S135.** Predicted functions of ORFs in the NZ_FOYZ01000004.1 containing WP_092559882.1

| gene | aa [a] | putative function | Protein homologue | %identity/ %similarity |
|---|---|---|---|---|
| WP_092559872.1 | 365 | 4Fe-4S dicluster | / | / |
| WP_092559873.1 | 1023 | NRPS | NspC [ADA69239.2] | 32/51 |
| WP_092559874.1 | 458 | MATE family efflux transporter | SorJ [ADN68485.1] | 33/56 |
| WP_092559875.1 | 2153 | NRPS | BaeJ [CAG23957.2] | 30/50 |
| WP_092559876.1 | 253 | Ser/Thr protein phosphatase | ThaC [ABC35295.1] | 35/57 |
| WP_092559877.1 | 754 | AT/Ox | DifA [CAG23974.1] | 56/74 |
| WP_092559878.1 | 2121 | AT-less type I PKS | BasE [ERM18797.1] | 56/71 |
| WP_092559879.1 | 4872 | AT-less type I PKS | BaeN [CAG23960.2] | 34/53 |
| WP_092559880.1 | 1495 | AT-less type I PKS | SorA [ADN68476.1] | 40/59 |
| WP_092559881.1 | 2039 | AT-less type I PKS | BasE [ERM18797.1] | 40/57 |
| WP_092559882.1 | 419 | HMG-CoA synthase | BaeG [CAG23954.2] | 67/80 |
| WP_092559883.1 | 249 | enoyl-CoA hydratase | BaeH [CAG23955.1] | 50/68 |
| WP_092559884.1 | 618 | enoyl-CoA hydratase | ElaN [AEC04360.1] | 47/66 |
| WP_092559885.1 | 436 | glycoside hydrolase | / | / |
| WP_092559886.1 | 189 | hypothetical protein | / | / |

[a] Number of amino acids

**Table S136.** Predicted functions of ORFs in the NZ_FMUV01000003.1 containing WP_092974861.1

| gene | aa [a] | putative function | Protein homologue | %identity/ %similarity |
|---|---|---|---|---|
| WP_092974809.1 | 6641 | AT-less type I PKS | BaeL [CAG23958.2] | 45/62 |
| WP_092974812.1 | 1987 | AT-less type I PKS | DifI [CAJ57409.1] | 46/63 |
| WP_092974815.1 | 2043 | AT-less type I PKS | DifF [CAG23977.1] | 33/52 |
| WP_092974818.1 | 286 | AT-less type I PKS | DifL [CAG23983.1] | 43/58 |
| WP_092974821.1 | 5556 | AT-less type I PKS | TaO [ABF92489.1] | 32/51 |
| WP_092974824.1 | 2278 | AT-less type I PKS | ThaO [ABC34675.1] | 37/52 |
| WP_092974827.1 | 1558 | AT-less type I PKS | PsyD [ADA82585.1] | 31/50 |
| WP_092974829.1 | 308 | ATP-binding cassette domain | SgvT2 [AGN74890.1] | 25/42 |
| WP_092974832.1 | 265 | hypothetical protein | / | / |
| WP_092974835.1 | 270 | hypothetical protein | / | / |
| WP_092974838.1 | 70 | hypothetical protein | / | / |
| WP_092974842.1 | 242 | ABC transporter ATP-binding protein | LnmR [AAN85531.1] | 26/43 |
| WP_092974846.1 | 244 | ABC transporter permease | / | / |
| WP_092974849.1 | 241 | PPTase | Mis12 [AKQ22701.1] | 28/53 |
| WP_092974852.1 | 225 | MBL fold metallo-hydrolase | BaeB [CAG23949.2] | 46/63 |
| WP_092974855.1 | 318 | acyltransferase | BaeD [CAG23951.1] | 39/60 |
| WP_092974858.1 | 758 | AT/Ox | DifA [CAG23974.1] | 50/72 |
| WP_092974861.1 | 421 | HMG-CoA synthase | ThaK [ABC34601.1] | 62/79 |
| WP_092974864.1 | 252 | enoyl-CoA hydratase | BaeH [CAG23955.1] | 47/65 |
| WP_092974867.1 | 254 | enoyl-CoA hydratase | BatE [ADD82946.1] | 54/74 |
| WP_092974870.1 | 81 | acyl carrier protein | ElaE [AEC04351.1] | 51/72 |
| WP_092974873.1 | 429 | beta-ketoacyl synthase | CorD [ADI59526.1] | 40/59 |
| WP_092974876.1 | 245 | 3-oxoacyl-ACP reductase FabG | TmlS [CBK62720.1] | 35/55 |
| WP_092974879.1 | 343 | FAD:protein FMN transferase | / | / |
| WP_092974882.1 | 230 | ABC transporter ATP-binding protein | LnmR [AAN85531.1] | 33/53 |
| WP_092974885.1 | 193 | TetR/AcrR family transcriptional regulator | / | / |
| WP_092974888.1 | 155 | terminase small subunit | / | / |

[a] Number of amino acids

**Table S137.** Predicted functions of ORFs in the NZ_NHTR01000001.1 containing WP_094099311.1

| gene | aa [a] | putative function | Protein homologue | %identity/ %similarity |
|---|---|---|---|---|
| WP_065547235.1 | 166 | acetyl-CoA carboxylase biotin carboxyl carrier | / | / |
| WP_065547234.1 | 453 | ATP-grasp domain-containing protein | / | / |
| WP_065547233.1 | 230 | PPTase | MupN [AAM12928.1] | 25/47 |
| WP_065547232.1 | 174 | antiterminator LoaP | TaA [ABF91060.1] | 24/39 |
| WP_065547231.1 | 314 | acyltransferase | BaeD [CAG23951.1] | 34/53 |
| WP_065547230.1 | 82 | acyl carrier protein | PsyL [ADA82592.1] | 33/65 |
| WP_065547229.1 | 750 | AT/Ox | DifA [CAG23974.1] | 51/70 |
| WP_065547228.1 | 430 | beta-ketoacyl synthase | DipR [AGS06821.1] | 41/62 |
| WP_065547227.1 | 253 | enoyl-CoA hydratase | MxnG [AGS77287.1] | 54/68 |
| WP_065547226.1 | 89 | hypothetical protein | / | / |
| WP_065547225.1 | 593 | carbamoyl transferase | albXV [CAE52324.1] | 50/70 |
| WP_065547224.1 | 200 | hypothetical protein | / | / |
| WP_094099311.1 | 411 | HMG-CoA synthase | MxnE [AGS77285.1] | 55/72 |
| WP_065547222.1 | 253 | enoyl-CoA hydratase | MxnF [AGS77286.1] | 45/61 |
| WP_084414505.1 | 585 | AT-less type I PKS | BaeJ [CAG23957.2] | 32/49 |
| WP_065547220.1 | 6509 | AT-less type I PKS | MisF [AKQ22696.1] | 33/50 |
| WP_065547219.1 | 3338 | AT-less type I PKS | CalF [BAP05594.1] | 36/52 |
| WP_065547218.1 | 4052 | AT-less type I PKS | Ta1 [ABF85931.1] | 34/50 |
| WP_065547217.1 | 3331 | AT-less type I PKS | Ta1 [ABF85931.1] | 34/50 |
| WP_065547216.1 | 103 | hypothetical protein | / | / |
| WP_065547214.1 | 323 | radical SAM protein | / | / |

[a] Number of amino acids

**Table S138.** Predicted functions of ORFs in the NZ_LGSU01001496.1 containing WP_094678774.1

| gene | aa [a] | putative function | Protein homologue | %identity/ %similarity |
|---|---|---|---|---|
| WP_094678762.1 | 98 | hypothetical protein | / | / |
| WP_094678764.1 | 77 | hypothetical protein | / | / |
| WP_094678766.1 | 397 | glycosyltransferase | / | / |
| WP_094678768.1 | 1621 | AT-less type I PKS | JamE [AAS98777.1] | 65/77 |
| WP_094678770.1 | 79 | acyl carrier protein | JamF [AAS98799.1] | 91/96 |
| WP_094678772.1 | 475 | beta-ketoacyl synthase | JamG [AAS98778.1] | 86/93 |
| WP_094678774.1 | 419 | HMG-CoA synthase | JamH [AAS98779.1] | 88/94 |
| WP_094678776.1 | 251 | enoyl-CoA hydratase | JamI [AAS98780.1] | 75/85 |
| WP_094678778.1 | 577 | thioesterase | CylH [ARU81122.1] | 60/71 |
| WP_094678780.1 | 862 | AT-less type I PKS | JamL [AAS98783.1] | 63/78 |
| WP_094678782.1 | 1523 | AT-less type I PKS | OzmL [ABS90473.1] | 27/49 |

[a] Number of amino acids

**Table S139.** Predicted functions of ORFs in the NZ_MRYI01000022.1 containing WP_094712021.1

| gene | aa [a] | putative function | Protein homologue | %identity/ %similarity |
|---|---|---|---|---|
| WP_094712008.1 | 284 | hypothetical protein | / | / |
| WP_094712009.1 | 138 | carboxymuconolactone decarboxylase | / | / |
| WP_094712010.1 | 2153 | NRPS | SnbDE [CBW45647.1] | 28/43 |
| WP_094712011.1 | 1659 | NRPS | leinamycin [AAN85512.1] | 30/46 |
| WP_094712012.1 | 433 | hypothetical protein | / | / |
| WP_094712013.1 | 410 | flavin-dependent oxidoreductase | tartrolon [ACR11438.1] | 44/63 |
| WP_094712014.1 | 1102 | type I PKS | NspD [ADA69241.1] | 41/57 |
| WP_094712015.1 | 263 | PPTase | kirP [CAN89630.1] | 40/55 |
| WP_094712016.1 | 256 | enoyl-CoA hydratase | OocD [AFX60326.1] | 53/70 |
| WP_094712017.1 | 378 | S-malonyltransferase | PedD [AAS47563.1] | 55/71 |
| WP_094712018.1 | 427 | beta-ketoacyl synthase | PedM [AAW33972.1] | 45/63 |
| WP_094712019.1 | 83 | acyl carrier protein | PedN [AAW33973.1] | 45/65 |
| WP_094712020.1 | 132 | acyl-CoA thioesterase | / | / |
| WP_094712021.1 | 409 | HMG-CoA synthase | MxnE [AGS77285.1] | 56/72 |
| WP_094712022.1 | 2550 | AT-less type I PKS | LglD [AIU36100.1] | 31/48 |
| WP_094712023.1 | 2019 | AT-less type I PKS | OocJ [AFX60332.1] | 36/52 |
| WP_094712024.1 | 5552 | AT-less type I PKS | CalB [BAP05590.1] | 34/50 |
| WP_094712025.1 | 224 | hypothetical protein | / | / |
| WP_094712026.1 | 459 | hypothetical protein | / | / |

[a] Number of amino acids

**Table S140.** Predicted functions of ORFs in the NZ_CP023254.1 containing WP_095839501.1

| gene | aa [a] | putative function | Protein homologue | %identity/ %similarity |
|---|---|---|---|---|
| WP_095839487.1 | 286 | AraC family transcriptional regulator | / | / |
| WP_095839488.1 | 335 | alcohol dehydrogenase | MupE [AAM12917.1] | 26/45 |
| WP_095839489.1 | 70 | hypothetical protein | / | / |
| WP_095839490.1 | 174 | antitermination factor | ElaA [AEC04347.1] | 55/75 |
| WP_095839491.1 | 791 | S-malonyltransferase | ElaB [AEC04348.1] | 59/73 |
| WP_095839492.1 | 324 | acyltransferase | ElaC [AEC04349.1] | 45/62 |
| WP_095839493.1 | 94 | hypothetical protein | ElaD [AEC04350.1] | 57/69 |
| WP_095839494.1 | 11059 | AT-less type I PKS | TaO [ABF92489.1] | 44/60 |
| WP_095839495.1 | 1782 | AT-less type I PKS | MisD [AKQ22698.1] | 41/57 |
| WP_095839496.1 | 10493 | AT-less type I PKS | BasE [ERM18797.1] | 43/59 |
| WP_095839497.1 | 3019 | AT-less type I PKS | MisD [AKQ22698.1] | 45/61 |
| WP_095839498.1 | 13566 | AT-less type I PKS | Bat2 [ADD82940.1] | 39/54 |
| WP_095839499.1 | 80 | acyl carrier protein | ElaE [AEC04351.1] | 65/86 |
| WP_095839500.1 | 412 | beta-ketoacyl synthase | CalW [BAP05575.1] | 64/79 |
| WP_095839501.1 | 420 | HMG-CoA synthase | BatC [ADD82944.1] | 74/87 |
| WP_095839502.1 | 254 | enoyl-CoA hydratase | BatD [ADD82945.1] | 62/76 |
| WP_095839503.1 | 249 | enoyl-CoA hydratase | BatE [ADD82946.1] | 70/86 |
| WP_095839504.1 | 1756 | AT-less type I PKS | BaeL [CAG23958.2] | 54/67 |
| WP_095839505.1 | 1442 | AT-less type I PKS | ElaR [AEC04364.1] | 44/59 |
| WP_095839506.1 | 536 | carbamoyltransferase | BatF [ADD82947.1] | 49/66 |
| WP_095839507.1 | 76 | hypothetical protein | BatF [ADD82947.1] | 71/85 |
| WP_095839508.1 | 592 | carbamoyl transferase | albXV [CAE52324.1] | 49/65 |
| WP_095839509.1 | 590 | carbamoyl transferase | albXV [CAE52324.1] | 51/66 |
| WP_095839510.1 | 246 | PPTase | Mis12 [AKQ22701.1] | 37/58 |
| WP_095839511.1 | 96 | hypothetical protein | ElaD [AEC04350.1] | 39/58 |
| WP_095839512.1 | 243 | pyridoxal phosphate-dependent enzyme | / | / |
| WP_095839513.1 | 295 | mechanosensitive ion channel protein | / | / |

[a] Number of amino acids

**Table S141.** Predicted functions of ORFs in the NZ_FXXN01000026.1 containing WP_096703393.1

| gene | aa [a] | putative function | Protein homologue | %identity/ %similarity |
|---|---|---|---|---|
| WP_096703386.1 | 143 | PIN domain-containing protein | / | / |
| WP_096703387.1 | 80 | type II toxin-antitoxin | / | / |
| WP_096703388.1 | 228 | DUF4102 domain-containing protein | / | / |
| WP_096703389.1 | 6777 | AT-less type I PKS | DszB [AAY32965.1] | 36/48 |
| WP_096703390.1 | 5862 | NRPS | TaO [ABF92489.1] | 40/53 |
| WP_096703391.1 | 113 | hypothetical protein | / | / |
| WP_096703392.1 | 269 | enoyl-CoA hydratase | CorF [ADI59528.1] | 38/56 |
| WP_096703393.1 | 415 | HMG-CoA synthase | MxnE [AGS77285.1] | 57/71 |
| WP_096703394.1 | 426 | beta-ketoacyl synthase | MxnD [AGS77284.1] | 45/61 |
| WP_096703395.1 | 87 | acyl carrier protein | AcpK [CAG23953.1] | 46/62 |
| WP_096703396.1 | 761 | AT/ER | MxnA [AGS77281.1] | 50/66 |
| WP_096703397.1 | 268 | PPTase | Mis12 [AKQ22701.1] | 35/49 |
| WP_096703398.1 | 698 | ABC transporter ATP-binding protein | CalU [BAP05577.1] | 40/60 |
| WP_096703399.1 | 79 | hypothetical protein | / | / |
| WP_096703300.1 | 204 | hypothetical protein | / | / |

[a] Number of amino acids

**Table S142.** Predicted functions of ORFs in the NZ_BAOS01000001.1 containing WP_096892270.1

| gene | aa [a] | putative function | Protein homologue | %identity/ %similarity |
|---|---|---|---|---|
| WP_096892266.1 | 75 | hypothetical protein | / | / |
| WP_096892267.1 | 486 | hypothetical protein | / | / |
| WP_096892268.1 | 250 | PPTase | Mis12 [AKQ22701.1] | 34/54 |
| WP_096892269.1 | 920 | hypothetical protein | / | / |
| WP_096892270.1 | 410 | HMG-CoA synthase | CylF [ARU81120.1] | 56/72 |
| WP_096892271.1 | 62 | enoyl-CoA hydratase | MxnG [AGS77287.1] | 47/68 |
| WP_096892272.1 | 258 | polyketide synthase | NspC [ADA69239.2] | 53/70 |
| WP_096892273.1 | 1205 | AT-less type I PKS | DszB [AAY32965.1] | 34/52 |
| WP_096892274.1 | 810 | AT-less type I PKS | BaeL [CAG23958.2] | 50/67 |
| WP_096892275.1 | 84 | acyl carrier protein | PtzD [AHC73995.1] | 28/50 |
| WP_096892276.1 | 297 | 2-hydroxyglutaryl-CoA dehydratase | / | / |
| WP_096892277.1 | 348 | 2-hydroxyacyl-CoA dehydratase | / | / |
| WP_096892278.1 | 416 | 2-hydroxyacyl-CoA dehydratase | / | / |
| WP_096892279.1 | 4670 | AT-less type I PKS | CorL [ADI59534.1] | 34/52 |
| WP_096892280.1 | 3703 | AT-less type I PKS | DszB [AAY32965.1] | 39/55 |
| WP_096892281.1 | 2589 | AT-less type I PKS | DszB [AAY32965.1] | 34/51 |
| WP_096892282.1 | 2305 | AT-less type I PKS | MisC [AKQ22699.1] | 37/56 |
| WP_096892283.1 | 445 | radical SAM protein | / | / |
| WP_096892284.1 | 247 | class I SAM-dependent methyltransferase | TaQ [ABF89350.1] | 42/56 |
| WP_096892285.1 | 453 | radical SAM protein | / | / |

[a] Number of amino acids

**Table S143.** Predicted functions of ORFs in the NZ_OBML01000008.1 containing WP_097175566.1

| gene | aa [a] | putative function | Protein homologue | %identity/ %similarity |
|---|---|---|---|---|
| WP_097175612.1 | 430 | TRAP transporter small permease | / | / |
| WP_097175560.1 | 149 | 3-dehydroquinate dehydratase | / | / |
| WP_097175561.1 | 321 | acyltransferase | BaeD [CAG23951.1] | 38/57 |
| WP_097175562.1 | 274 | PPTase | MupN [AAM12928.1] | 38/53 |
| WP_097175563.1 | 288 | Ser/Thr protein phosphatase | ThaC [ABC35295.1] | 33/43 |
| WP_097175564.1 | 316 | FkbM family methyltransferase | CalA [BAP05589.1] | 35/53 |
| WP_097175565.1 | 418 | cytochrome P450 | ElaG [AEC04353.1] | 31/51 |
| WP_097175566.1 | 425 | HMG-CoA synthase | ThaK [ABC34601.1] | 62/77 |
| WP_067221771.1 | 81 | acyl carrier protein | CalX [BAP05574.1] | 50/73 |
| WP_097175567.1 | 406 | beta-ketoacyl synthase | TstN [AGN11888.1] | 52/65 |
| WP_097175568.1 | 257 | enoyl-CoA hydratase | NspJ [ADA69246.1] | 46/64 |
| WP_097175569.1 | 669 | asparagine synthase | SmdH [CCC21122.1] | 57/72 |
| WP_067221777.1 | 82 | acyl carrier protein | ChxC [AFO59864.1] | 52/73 |
| WP_097175570.1 | 545 | cyclic peptide export ABC | / | / |
| WP_097175571.1 | 549 | cyclic peptide export ABC transporter | / | / |
| WP_097175572.1 | 787 | AT/Ox | CalY [BAP05573.1] | 53/70 |
| WP_097175573.1 | 6900 | AT-less type I PKS | ChxE [AFO59866.1] | 46/57 |
| WP_097175574.1 | 445 | monooxygenase | PedG [AAS47561.1] | 62/75 |
| WP_097175575.1 | 6761 | NRPS | TaO [ABF92489.1] | 42/55 |
| WP_067221789.1 | 288 | prohibitin family protein | / | / |

[a] Number of amino acids

**Table S144.** Predicted functions of ORFs in the NZ_OCNE01000015.1 containing WP_097232703.1

| gene | aa [a] | putative function | Protein homologue | %identity/ %similarity |
|---|---|---|---|---|
| WP_097232696.1 | 832 | Hsp70 family protein | / | / |
| WP_097232697.1 | 323 | tetratricopeptide repeat protein | / | / |
| WP_097232698.1 | 191 | DUF98 domain-containing protein | / | / |
| WP_097232699.1 | 308 | PPTase | Mis12 [AKQ22701.1] | 31/48 |
| WP_097232700.1 | 632 | S-malonyltransferase | BasH [ERM18800.1] | 39/59 |
| WP_097232701.1 | 83 | acyl carrier protein | PsyL [ADA82592.1] | 41/64 |
| WP_097232702.1 | 459 | beta-ketoacyl synthase | CorD [ADI59526.1] | 43/54 |
| WP_097232703.1 | 419 | HMG-CoA synthase | BatC [ADD82944.1] | 68/80 |
| WP_097232704.1 | 260 | enoyl-CoA hydratase | ElaM [AEC04359.1] | 50/66 |
| WP_097232705.1 | 263 | enoyl-CoA hydratase | BatE [ADD82946.1] | 59/75 |
| WP_097232706.1 | 3784 | AT-less type I PKS | bonA [AFN27480.1] | 42/53 |

[a] Number of amino acids

**Table S145.** Predicted functions of ORFs in the NZ_NVNE01000022.1 containing WP_097812164.1 and WP_097812167.1

| gene | aa [a] | putative function | Protein homologue | %identity/ %similarity |
|---|---|---|---|---|
| WP_097812173.1 | 186 | L,D-transpeptidase | / | / |
| WP_097812172.1 | 228 | GIY-YIG nuclease family protein | / | / |
| WP_097812171.1 | 177 | antiterminator LoaP | / | / |
| WP_097812184.1 | 225 | MBL fold metallo-hydrolase | BaeB [CAG23949.2] | 55/69 |
| WP_097812170.1 | 1085 | S-malonyltransferase | BasH [ERM18800.1] | 51/71 |
| WP_097812169.1 | 267 | PPTase | Mis12 [AKQ22701.1] | 35/56 |
| WP_097812168.1 | 298 | S-malonyltransferase | BaeC [CAG23950.2] | 58/75 |
| WP_097812167.1 | 419 | HMG-CoA synthase | BatC [ADD82944.1] | 75/86 |
| WP_097812166.1 | 85 | acyl carrier protein | AcpK [CAG23953.1] | 61/78 |
| WP_097812165.1 | 412 | beta-ketoacyl synthase | CalW [BAP05575.1] | 62/77 |
| WP_097812164.1 | 420 | HMG-CoA synthase | BatC [ADD82944.1] | 71/82 |
| WP_097812163.1 | 254 | enoyl-CoA hydratase | BaeH [CAG23955.1] | 59/74 |
| WP_097812162.1 | 250 | enoyl-CoA hydratase | BatE [ADD82946.1] | 69//85 |
| WP_097812161.1 | 2474 | AT-less type I PKS | ElaP [AEC04362.1] | 38/57 |
| WP_097812160.1 | 3104 | AT-less type I PKS | MisF [AKQ22696.1] | 45/64 |
| WP_097812159.1 | 1917 | AT-less type I PKS | Bat3 [ADD82941.1] | 43/60 |
| WP_097826716.1 | 1549 | AT-less type I PKS | PsyD [ADA82585.1] | 37/57 |
| WP_097812157.1 | 263 | hypothetical protein | / | / |
| WP_097812156.1 | 394 | hypothetical protein | / | / |
| WP_097812155.1 | 472 | AMP-dependent synthetase | MalI [ABC34723.1] | 26//44 |
| WP_097812154.1 | 255 | class I SAM-dependent methyltransferase | / | / |
| WP_097812153.1 | 82 | acyl carrier protein | SmdG [CCC21121.1] | 49/72 |
| WP_097812152.1 | 1383 | AT-less type I PKS | DifH [CAJ57408.1] | 38/56 |
| WP_097812151.1 | 303 | nucleoside hydrolase | / | / |
| WP_097812150.1 | 99 | hypothetical protein | / | / |

[a] Number of amino acids

**Table S146.** Predicted functions of ORFs in the NZ_NUCA01000042.1 containing WP_098202504.1

| gene | aa [a] | putative function | Protein homologue | %identity/ %similarity |
|---|---|---|---|---|
| PEP90341.1 | 682 | AT-less type I PKS | TaO [ABF92489.1] | 60/73 |
| WP_098202502.1 | 249 | enoyl-CoA hydratase | BatE [ADD82946.1] | 70/84 |
| WP_098202503.1 | 254 | enoyl-CoA hydratase | BaeH [CAG23955.1] | 63/77 |
| WP_098202504.1 | 419 | HMG-CoA synthase | BaeG [CAG23954.2] | 77/86 |
| WP_098202505.1 | 408 | beta-ketoacyl synthase | CalW [BAP05575.1] | 62/78 |
| WP_098202506.1 | 82 | acyl carrier protein | AcpK [CAG23953.1] | 63/78 |

[a] Number of amino acids

**Table S147.** Predicted functions of ORFs in the NZ_CP024608.1 containing WP_099878367.1

| gene | aa [a] | putative function | Protein homologue | %identity/ %similarity |
|---|---|---|---|---|
| WP_099878336.1 | 426 | GMC family oxidoreductase | / | / |
| WP_099878338.1 | 244 | hypothetical protein | / | / |
| WP_099878341.1 | 259 | hypothetical protein | / | / |
| WP_099878344.1 | 463 | long-chain fatty acid--CoA ligase | DifD [CAJ57407.1] | 47/65 |
| WP_099878347.1 | 245 | SDR family oxidoreductase | DifE [CAG23976.1] | 48/69 |
| WP_099878350.1 | 89 | acyl carrier protein | DifC [CAJ57406.1] | 45//68 |
| WP_099878353.1 | 5011 | AT-less type I PKS | Bat2 [ADD82940.1] | 42/57 |
| WP_099878356.1 | 4721 | AT-less type I PKS | DifF [CAG23977.1] | 40/59 |
| WP_099878360.1 | 4335 | AT-less type I PKS | BaeN [CAG23960.2] | 39/56 |
| WP_099878363.1 | 235 | MBL fold metallo-hydrolase | BaeB [CAG23949.2] | 44/57 |
| WP_099878367.1 | 419 | HMG-CoA synthase | fr9K [AIC32697.1] | 73/85 |
| WP_099878370.1 | 263 | enoyl-CoA hydratase | TstL [AGN11886.1] | 61/75 |
| WP_099878371.1 | 249 | enoyl-CoA hydratase | BatE [ADD82946.1] | 64/78 |
| WP_099878374.1 | 79 | acyl carrier protein | NspG [ADA71314.1] | 56/76 |
| WP_099878377.1 | 417 | beta-ketoacyl synthase | TstN [AGN11888.1] | 67/80 |
| WP_099878380.1 | 377 | S-malonyltransferase | TstO [AGN11889.1] | 57/70 |
| WP_099878383.1 | 97 | hypothetical protein | / | / |
| WP_099878385.1 | 323 | acyltransferase | TstB [AGN11890.1] | 48/64 |
| WP_099878388.1 | 839 | methyltransferase | BatL [ADD82953.1] | 59/73 |
| WP_099878390.1 | 277 | PPTase | BatI [ADD82950.1] | 52/66 |
| WP_099878393.1 | 816 | apolipoprotein N-acyltransferase | / | / |
| WP_099882621.1 | 201 | hypothetical protein | / | / |
| WP_099878395.1 | 362 | hypothetical protein | / | / |
| WP_099878398.1 | 764 | TonB-dependent siderophore receptor | / | / |

[a] Number of amino acids

**Table S148.** Predicted functions of ORFs in the NZ_NNBN01000011.1 containing WP_100564352.1

| gene | aa [a] | putative function | Protein homologue | %identity/ %similarity |
|---|---|---|---|---|
| WP_100564314.1 | 4128 | NRPS | LnmI [AAN85522.1] | 50/59 |
| WP_100564316.1 | 315 | regulatory protein | leinamycin [AAN85494.1] | 40/55 |
| WP_100564318.1 | 306 | kinase | kirromycin [CAN89658.1] | 54/62 |
| WP_100564320.1 | 1059 | NRPS | CalI [BAP05597.1] | 33/48 |
| WP_100564322.1 | 402 | ATP-grasp domain | kirromycin [CAN89659.1] | 42/53 |
| WP_100564324.1 | 487 | argininosuccinate lyase | kirromycin [CAN89657.1] | 39/52 |
| WP_100564326.1 | 352 | cysteine synthase | kirromycin [CAN89659.1] | 45/56 |
| WP_100564328.1 | 160 | acyl carrier protein | kirromycin [CAN89663.1] | 41/50 |
| WP_100564330.1 | 317 | chlorinating enzyme | / | / |
| WP_100564332.1 | 301 | alpha/beta fold hydrolase | AlbXI [CAE52328.1] | 27/42 |
| WP_100564570.1 | 6931 | AT-less type I PKS | SorA [ADN68476.1] | 35/47 |
| WP_100564334.1 | 299 | AT/DC | LnmK [AAN85524.1] | 54/67 |
| WP_100564336.1 | 86 | acyl carrier protein | LnmL [AAN85525.1] | 58/68 |
| WP_100564338.1 | 406 | HMG-CoA synthase | LnmL [AAN85525.1] | 66/77 |
| WP_100564340.1 | 79 | acyl carrier protein | PsyL [ADA82592.1] | 35/70 |
| WP_100564342.1 | 418 | beta-ketoacyl synthase | MxnD [AGS77284.1] | 45/58 |
| WP_100564344.1 | 266 | thioesterase | LnmN [AAN85527.1] | 53/64 |
| WP_100564346.1 | 210 | Raf kinase inhibitor | / | / |
| WP_100564348.1 | 127 | DUF3224 domain | LnmZ' [AAN85540.1] | 37/55 |
| WP_100564350.1 | 261 | enoyl-CoA hydratase | CylG [ARU81121.1] | 41/60 |
| WP_100564352.1 | 415 | HMG-CoA synthase | PyxM [ASA76639.1] | 58/69 |
| WP_100564572.1 | 451 | cation/H(+) antiporter | / | / |
| WP_100564354.1 | 244 | Crp/Fnr family transcriptional regulator | LnmO [AAN85528.1] | 43/61 |
| WP_100564356.1 | 134 | DUF3224 domain | LnmZ' [AAN85540.1] | 39/55 |
| WP_100564358.1 | 314 | hypothetical protein | LnmH [AAN85521.1] | 44/59 |
| WP_100564360.1 | 514 | long-chain fatty acid--CoA ligase | LnmW [AAN85536.1] | 42/55 |
| WP_100564362.1 | 236 | N-acetylglucosaminyl deacetylase | LnmX [AAN85537.1] | 49/62 |
| WP_100564364.1 | 294 | hypothetical protein | LnmE [AAN85518.1] | 47/65 |
| WP_100564574.1 | 370 | acyl carrier protein | / | / |
| WP_100564366.1 | 500 | long-chain fatty acid--CoA ligase | LnmW [AAN85536.1] | 43/57 |
| WP_100564576.1 | 451 | serine hydroxymethyltransferase | / | / |
| WP_107528456.1 | 883 | NRPS | JamO [AAS98786.1] | 34/48 |
| WP_100564370.1 | 118 | DUF427 domain-containing protein | / | / |
| WP_100564372.1 | 362 | hypothetical protein | / | / |

[a] Number of amino acids

**Table S149.** Predicted functions of ORFs in the NZ_PHSU01000014.1 containing WP_100941015.1

| gene | aa [a] | putative function | Protein homologue | %identity/ %similarity |
|---|---|---|---|---|
| WP_100940998.1 | 201 | threonine transporter RhtB | oocB [AFX60324.1] | 33/53 |
| WP_100940999.1 | 272 | LuxR family transcriptional regulator | MupR [AAK28504.1] | 27/44 |
| WP_100941000.1 | 137 | hypothetical protein | MupZ [AFD33556.1] | 76/83 |
| WP_100941001.1 | 369 | LLM class flavin-dependent oxidoreductase | MupA [AAK28503.1] | 80/87 |
| WP_100941002.1 | 2877 | AT-less type I PKS | MmpA [AAM12909.2] | 66/75 |
| WP_100941003.1 | 304 | condensing domain | MupB1 [AAM12910.1] | 68/78 |
| WP_100941004.1 | 2000 | AT-less type I PKS | MmpB [AAM12911.1] | 62/73 |
| WP_100941005.1 | 1103 | AT/Ox | MmpC [AAM12912.1] | 74/82 |
| WP_100941006.1 | 6651 | AT-less type I PKS | MmpD [AAM12913.1] | 63/73 |
| WP_100941007.1 | 434 | NADH: flavin oxidoreductase | MupC [AAM12914.1] | 80/85 |
| WP_100941008.1 | 92 | acyl carrier protein | MacpA [AAM12915.1] | 79/91 |
| WP_100941009.1 | 249 | SDR family oxidoreductase | MupD [AAM12916.1] | 69//78 |
| WP_100941010.1 | 340 | NADPH: quinone oxidoreductase | MupE [AAM12917.1] | 83/90 |
| WP_100941011.1 | 85 | acyl carrier protein | MacpB [AAM12918.1] | 73/81 |
| WP_100941012.1 | 332 | short-chain dehydrogenases/reductases | MupF [AAM12919.1] | 72/81 |
| WP_100941013.1 | 79 | acyl carrier protein | Macp [AAM12920.1] | 68/82 |
| WP_100941014.1 | 411 | beta-ketoacyl synthase | MupG [AAM12921.1] | 81/90 |
| WP_100941015.1 | 421 | HMG-CoA synthase | MupH [AAM12922.1] | 88/95 |
| WP_100941016.1 | 255 | enoyl-CoA hydratase | MupJ [AAM12923.1] | 79/88 |
| WP_100941017.1 | 248 | enoyl-CoA hydratase | MupK [AAM12924.1] | 85/91 |
| WP_100941018.1 | 1198 | FAD-binding protein | MmpE [AAM12925.1] | 67/76 |
| WP_100941019.1 | 285 | alpha/beta hydrolase | MupL [AAM12926.1] | 72/83 |
| WP_100941020.1 | 1030 | isoleucine--tRNA ligase | MupM [AAM12927.1] | 89/93 |
| WP_100941021.1 | 288 | PPTase | MupN [AAM12928.1] | 56/69 |
| WP_100941022.1 | 454 | cytochrome P450 | MupO [AAM12929.1] | 84/89 |
| WP_100941023.1 | 312 | hypothetical protein | MupP [AAM12930.1] | 60/71 |
| WP_100941024.1 | 451 | long-chain fatty acid--CoA ligase | MupQ [AAM12931.1] | 74/83 |
| WP_100941025.1 | 255 | SDR family oxidoreductase | MupS [AAM12932.1] | 80/88 |
| WP_100941026.1 | 862 | polyketide synthase | MmpF [AAM12934.1] | 64/74 |
| WP_100941027.1 | 80 | hypothetical protein | MacpE [AAM12935.1] | 74/86 |
| WP_100941028.1 | 146 | Rieske (2Fe-2S) protein | MupT [AAM12936.1] | 64/81 |
| WP_100941029.1 | 528 | hypothetical protein | MupU [AAM12937.1] | 68/79 |
| WP_100941030.1 | 661 | alpha/beta fold hydrolase | MupV [AAM12938.1] | 79/87 |
| WP_100941031.1 | 472 | aromatic ring-hydroxylating dioxygenase | MupW [AAM12939.1] | 79/85 |
| WP_100941032.1 | 233 | hypothetical protein | MupR [AAK28504.1] | 70/82 |
| WP_100941033.1 | 511 | amidase | MupX [AAM12940.1] | 75/85 |
| WP_100941034.1 | 191 | acyl-homoserine-lactone synthase | MupI [AAK28505.1] | 75/84 |
| WP_100941035.1 | 69 | XRE family transcriptional regulator | / | / |
| WP_100941036.1 | 345 | isopenicillin N synthase family oxygenase | / | / |

[a] Number of amino acids

**Table S150.** Predicted functions of ORFs in the NZ_CP025429.1 containing WP_101709341.1

| gene | aa [a] | putative function | Protein homologue | %identity/ %similarity |
|---|---|---|---|---|
| WP_101709324.1 | 227 | cytidylyltransferase | / | / |
| WP_101709325.1 | 237 | DNA polymerase III subunit epsilon | / | / |
| WP_101709326.1 | 553 | hypothetical protein | corallopyronin [ADI59535.1] | 27/42 |
| WP_101709327.1 | 416 | beta-ketoacyl synthase | MxnD [AGS77284.1] | 46/61 |
| WP_101709328.1 | 83 | acyl carrier protein | OocG [AFX60329.1] | 47/68 |
| WP_101709329.1 | 518 | MFS transporter | leinamycin [AAN85487.1] | 31/47 |
| WP_101709330.1 | 449 | 2-nitropropane dioxygenase | BatK [ADD82952.1] | 56/75 |
| WP_101709331.1 | 211 | PPTase | LtmL [ACY01405.1] | 34/47 |
| WP_101709332.1 | 357 | acyltransferase | BaeD [CAG23951.1] | 41/55 |
| WP_101709333.1 | 286 | S-malonyltransferase | BaeC [CAG23950.2] | 50/69 |
| WP_101709334.1 | 259 | enoyl-CoA hydratase | MxnF [AGS77286.1] | 45/61 |
| WP_101709335.1 | 256 | enoyl-CoA hydratase | BatE [ADD82946.1] | 50/66 |
| WP_101709336.1 | 1126 | AT-less type I PKS | LglD [AIU36100.1] | 36/53 |
| WP_101709337.1 | 1510 | AT-less type I PKS | RhiE [CAL69893.1] | 44/59 |
| WP_101709338.1 | 2272 | AT-less type I PKS | MisC [AKQ22699.1] | 35/51 |
| WP_101709339.1 | 3204 | AT-less type I PKS | MisC [AKQ22699.1] | 34/49 |
| WP_101709340.1 | 928 | AT-less type I PKS | MxnI [AGS77289.1] | 30/44 |
| WP_101709341.1 | 410 | HMG-CoA synthase | MxnE [AGS77285.1] | 57/72 |
| WP_101709342.1 | 147 | ribonuclease HI | / | / |
| WP_101709343.1 | 254 | methyltransferase | / | / |
| WP_101710734.1 | 252 | hydroxyacylglutathione hydrolase | BaeB [CAG23949.2] | 32/49 |
| WP_101709344.1 | 631 | LysM peptidoglycan-binding domain | / | / |
| WP_101709345.1 | 347 | hypothetical protein | / | / |

[a] Number of amino acids

**Table S151.** Predicted functions of ORFs in the NZ_QAPB01000044.1 containing WP_103621801.1

| gene | aa [a] | putative function | Protein homologue | %identity/ %similarity |
|---|---|---|---|---|
| ORF1 | 704 | AT-less type I PKS | BaeL [CAG23958.2] | 59/73 |
| WP_103621799.1 | 248 | enoyl-CoA hydratase | BatE [ADD82946.1] | 72/83 |
| WP_103621800.1 | 258 | enoyl-CoA hydratase | BaeH [CAG23955.1] | 55/71 |
| WP_103621801.1 | 419 | HMG-CoA synthase | BaeG [CAG23954.2] | 75/86 |
| WP_103621803.1 | 411 | beta-ketoacyl synthase | TstN [AGN11888.1] | 61/75 |
| WP_103621802.1 | 82 | acyl carrier protein | TaB [ABF90753.1] | 62/78 |
| ORF2 | 2549 | AT-less type I PKS | MisE [AKQ22697.1] | 49/66 |

[a] Number of amino acids

**Table S152.** Predicted functions of ORFs in the NZ_QAPB01000044.1 containing WP_103865517.1

| gene | aa [a] | putative function | Protein homologue | %identity/ %similarity |
|---|---|---|---|---|
| WP_103865509.1 | 174 | antitermination factor | ElaA [AEC04347.1] | 33/53 |
| WP_103865510.1 | 276 | alpha/beta hydrolase | / | / |
| WP_103865511.1 | 3996 | AT-less type I PKS | BaeJ [CAG23957.2] | 36/55 |
| WP_103865512.1 | 2035 | AT-less type I PKS | DifF [CAG23977.1] | 37/56 |
| WP_103865513.1 | 4516 | AT-less type I PKS | BaeN [CAG23960.2] | 36/55 |
| WP_103865519.1 | 763 | S-malonyltransferase | Dor8 [ACY01393.1] | 48/67 |
| WP_103865514.1 | 79 | acyl carrier protein | AcpK [CAG23953.1] | 41/65 |
| WP_103865515.1 | 81 | acyl carrier protein | JamF [AAS98799.1] | 42/60 |
| WP_103865516.1 | 420 | beta-ketoacyl synthase | TmlG [CBK62726.1] | 39/61 |
| WP_103865517.1 | 414 | HMG-CoA synthase | BatC [ADD82944.1] | 65/79 |
| WP_103865518.1 | 256 | enoyl-CoA hydratase | CylG [ARU81121.1] | 51/69 |

[a] Number of amino acids

**Table S153.** Predicted functions of ORFs in the NZ_PRKR01000001.1 containing WP_104148568.1

| gene | aa [a] | putative function | Protein homologue | %identity/ %similarity |
|---|---|---|---|---|
| WP_104148558.1 | 193 | antiterminator LoaP | / | / |
| WP_104148559.1 | 225 | MBL fold metallo-hydrolase | BaeB [CAG23949.2] | 51/66 |
| WP_104148560.1 | 1101 | S-malonyltransferase | BasH [ERM18800.1] | 52/71 |
| WP_104148561.1 | 1555 | AT-less type I PKS | BasG [ERM18799.1] | 46/65 |
| WP_104148562.1 | 3605 | AT-less type I PKS | BryB [ABM63527.1] | 46/62 |
| WP_104148563.1 | 2664 | AT-less type I PKS | BryB [ABM63527.1] | 40/59 |
| ORF1 | 5861 | AT-less type I PKS | ElaK [AEC04357.1] | 44/62 |
| WP_104148564.1 | 2146 | AT-less type I PKS | OnnI [AAV97877.1] | 49/66 |
| WP_104148565.1 | 4537 | AT-less type I PKS | BaeN [CAG23960.2] | 39/58 |
| WP_104148566.1 | 82 | acyl carrier protein | NspG [ADA71314.1] | 51/76 |
| WP_104148567.1 | 411 | beta-ketoacyl synthase | CalW [BAP05575.1] | 63/81 |
| WP_104148568.1 | 419 | HMG-CoA synthase | BaeG [CAG23954.2] | 76/85 |
| WP_104148569.1 | 254 | enoyl-CoA hydratase | BaeH [CAG23955.1] | 55/72 |
| WP_104148570.1 | 247 | enoyl-CoA hydratase | BatE [ADD82946.1] | 69/83 |
| WP_104148571.1 | 1925 | AT-less type I PKS | TaO [ABF92489.1] | 52/71 |
| WP_104148572.1 | 3783 | AT-less type I PKS | DifI [CAJ57409.1] | 49/66 |
| WP_104148573.1 | 4369 | AT-less type I PKS | ElaK [AEC04357.1] | 46/63 |

[a] Number of amino acids

**Table S154.** Predicted functions of ORFs in the NZ_PTIY01000008.1 containing WP_104424066.1

| gene | aa [a] | putative function | Protein homologue | %identity/ %similarity |
|---|---|---|---|---|
| WP_104424063.1 | 480 | ferrochelatase | / | / |
| WP_104424064.1 | 236 | hypothetical protein | / | / |
| WP_104424065.1 | 226 | PPTase | dipB [AGS06837.1] | 46/69 |
| WP_104424091.1 | 126 | GxxExxY protein | / | / |
| WP_104424066.1 | 426 | HMG-CoA synthase | PedP [AAW33975.1] | 75/85 |
| WP_104424067.1 | 84 | acyl carrier protein | PedN [AAW33973.1] | 59/79 |
| WP_104424068.1 | 437 | beta-ketoacyl synthase | PedM [AAW33972.1] | 65/78 |
| WP_104424069.1 | 411 | enoyl-CoA hydratase | BaeH [CAG23955.1] | 57/72 |
| WP_104424070.1 | 568 | carboxylesterase family protein | / | / |
| WP_104424071.1 | 6176 | hybrid NRPS/PKS | PedH [AAS47562.1] | 58/71 |
| WP_104424072.1 | 324 | hypothetical protein | / | / |
| WP_104424073.1 | 111 | hypothetical protein | / | / |

[a] Number of amino acids

**Table S155.** Predicted functions of ORFs in the NZ_CP012673.1 containing WP_104987424.1

| gene | aa [a] | putative function | Protein homologue | %identity/ %similarity |
|---|---|---|---|---|
| WP_104982245.1 | 516 | serine/threonine protein kinase | Disorazols [AAY32975.1] | 39/55 |
| WP_104982246.1 | 329 | hypothetical protein | / | / |
| WP_104982247.1 | 234 | hypothetical protein | / | / |
| WP_104987424.1 | 413 | HMG-CoA synthase | MxnE [AGS77285.1] | 53/69 |
| WP_104982248.1 | 320 | enoyl-CoA hydratase | BonI [AFN27485.1] | 44/60 |
| WP_104982249.1 | 336 | hypothetical protein | KirAI [CAN89631.1] | 33/45 |
| WP_104982250.1 | 269 | enoyl-CoA hydratase | MxnF [AGS77286.1] | 44/59 |
| WP_104982251.1 | 81 | acyl carrier protein | NspG [ADA71314.1] | 42/58 |
| WP_104982252.1 | 112 | hypothetical protein | / | / |
| WP_104982253.1 | 85 | hypothetical protein | / | / |
| WP_104982254.1 | 975 | S-malonyltransferase | BryP [ABM63531.1] | 34/54 |
| WP_104982255.1 | 120 | hypothetical protein | / | / |
| WP_104982256.1 | 118 | hypothetical protein | / | / |
| WP_104982257.1 | 478 | oxidoreductase | / | / |
| WP_104982258.1 | 100 | hypothetical protein | Rhizopodin [CCA89332.1] | 40/63 |
| WP_104982259.1 | 4513 | AT-less type I PKS | DszA [AAY32964.1] | 42/55 |
| WP_104982260.1 | 685 | AT-less type I PKS | RizD [CCA89328.1] | 41/54 |
| WP_104982261.1 | 6166 | AT-less type I PKS | ChiE [AAY89052.1] | 43/55 |
| WP_104982262.1 | 4952 | AT-less type I PKS | BonD [AFN27483.1] | 42/57 |
| WP_104982263.1 | 401 | efflux RND transporter periplasmic adaptor | / | / |
| WP_104982264.1 | 1052 | multidrug efflux RND transporter permease | / | / |

[a] Number of amino acids

**Table S156.** Predicted functions of ORFs in the NZ_CP023666.1 containing WP_105979318.1

| gene [a] | aa [b] | putative function | Protein homologue | %identity/ %similarity |
|---|---|---|---|---|
| WP_003178866.1 | 247 | ABC transporter ATP-binding protein | LnmR [AAN85531.1] | 23/53 |
| WP_035337254.1 | 154 | hypothetical protein | / | / |
| WP_105979306.1 | 213 | TetR/AcrR family transcriptional regulator | / | / |
| WP_105979307.1 | 370 | NADH:flavin oxidoreductase | ChxG [AFO59868.1] | 53/65 |
| WP_105979308.1 | 336 | aldo/keto reductase | CylM [ARU81127.1] | 28/50 |
| WP_105979309.1 | 405 | cytochrome P450 | DifM [CAG23984.1] | 38/59 |
| WP_105979310.1 | 516 | NRPS | kirromycin [CAN89656.1] | 47/63 |
| WP_105979311.1 | 404 | MFS transporter | / | / |
| WP_105979312.1 | 1233 | AT-less type I PKS | NspD [ADA69241.1] | 58/73 |
| WP_105979313.1 | 1816 | AT-less type I PKS | BasE [ERM18797.1] | 52/68 |
| WP_105979314.1 | 2848 | AT-less type I PKS | BaeN [CAG23960.2] | 44/62 |
| WP_105979315.1 | 286 | PPTase | / | / |
| ORF1 | 3242 | AT-less type I PKS | MisF [AKQ22696.1] | 45/63 |
| ORF2 | 1055 | AT-less type I PKS | SorB [ADN68477.1] | 45/63 |
| ORF3 | 1671 | AT-less type I PKS | Ta1 [ABF85931.1] | 42/58 |
| WP_105979316.1 | 249 | enoyl-CoA hydratase | ElaN [AEC04360.1] | 66/81 |
| WP_105979317.1 | 256 | enoyl-CoA hydratase | BaeH [CAG23955.1] | 57/73 |
| WP_105979318.1 | 420 | HMG-CoA synthase | BatC [ADD82944.1] | 71/84 |
| WP_105979319.1 | 411 | beta-ketoacyl synthase | CalW [BAP05575.1] | 62/78 |
| WP_105979320.1 | 82 | acyl carrier protein | BatA [ADD82942.1] | 51/70 |
| WP_105979321.1 | 3102 | AT-less type I PKS | MisE [AKQ22697.1] | 49/66 |
| WP_105979322.1 | 1351 | AT-less type I PKS | SorE [ADN68480.1] | 48/64 |
| WP_105979323.1 | 4601 | AT-less type I PKS | BaeN [CAG23960.2] | 39/58 |
| WP_105979324.1 | 276 | SAM-dependent methyltransferase | NspF [ADA69243.1] | 27/48 |
| WP_105979325.1 | 259 | alpha/beta hydrolase | / | / |

[a] ORFs are proteins without protein ID provided by NCBI [b] Number of amino acids

**Table S157.** Predicted functions of ORFs in the NZ_CP027526.1 containing WP_106002065.1

| gene | aa [a] | putative function | Protein homologue | %identity/ %similarity |
|---|---|---|---|---|
| WP_106002049.1 | 1117 | response regulator | smdC [CCC21117.1] | 30/49 |
| WP_106002050.1 | 543 | cyclic peptide export ABC transporter | Rhizopodin [CCA89322.1] | 26/46 |
| WP_106002051.1 | 550 | cyclic peptide export ABC transporter | Rhizopodin [CCA89322.1] | 32/48 |
| WP_106002052.1 | 753 | AT/Ox | CalY [BAP05573.1] | 49/67 |
| WP_106002053.1 | 6342 | AT-less type I PKS | ChxE [AFO59866.1] | 43/54 |
| WP_106002054.1 | 433 | monooxygenase | PedG [AAS47561.1] | 62/76 |
| WP_106002055.1 | 6237 | NRPS | PedH [AAS47562.1] | 37/52 |
| WP_106002056.1 | 304 | prohibitin family protein | / | / |
| WP_106002057.1 | 437 | hyporthetical protein | / | / |
| WP_106002058.1 | 206 | DUF697 domain-containing protein | / | / |
| WP_106003749.1 | 442 | acyltransferase | BaeD [CAG23951.1] | 37/56 |
| WP_106003750.1 | 235 | amino acid ABC transporter substrate | | |
| WP_106002059.1 | 472 | hyporthetical protein | | |
| WP_106002060.1 | 81 | acyl carrier protein | SmdG [CCC21121.1] | 53/69 |
| WP_106002061.1 | 657 | asparagine synthase | SmdH [CCC21122.1] | 55/69 |
| WP_106002062.1 | 251 | enoyl-CoA hydratase | NspJ [ADA69246.1] | 48/63 |
| WP_106002063.1 | 407 | beta-ketoacyl synthase | NspH [ADA69244.1] | 44/63 |
| WP_106002064.1 | 82 | acyl carrier protein | CalX [BAP05574.1] | 42/66 |
| WP_106002065.1 | 418 | HMG-CoA synthase | ThaK [ABC34601.1] | 64/78 |
| WP_106002066.1 | 416 | cytochrome P450 | ElaG [AEC04353.1] | 31/51 |
| WP_106002067.1 | 311 | FkbM family methyltransferase | CalA [BAP05589.1] | 35/48 |
| WP_106002068.1 | 285 | Ser/Thr protein phosphatase | ThaC [ABC35295.1] | 29/41 |
| WP_106002069.1 | 317 | lipid A deacylase LpxR | / | / |
| WP_106002070.1 | 99 | PepSY domain-containing protein | / | / |

[a] Number of amino acids

**Table S158.** Predicted functions of ORFs in the NZ_PYAW01000002.1 containing WP_106528595.1

| gene | aa [a] | putative function | Protein homologue | %identity/ %similarity |
|---|---|---|---|---|
| WP_106528579.1 | 159 | plasmid maintenance system antidote protein | / | / |
| WP_106528580.1 | 87 | hypothetical protein | / | / |
| WP_106528581.1 | 232 | PPTase | Mis12 [AKQ22701.1] | 33/51 |
| WP_106528582.1 | 449 | glycosyltransferase | CylN [ARU81128.1] | 27/46 |
| WP_106528583.1 | 2518 | AT-less type I PKS | BaeJ [CAG23957.2] | 43/57 |
| WP_106528584.1 | 6116 | AT-less type I PKS | BaeM [CAG23959.2] | 43/60 |
| WP_106528585.1 | 3106 | AT-less type I PKS | SorA [ADN68476.1] | 41/57 |
| WP_106528586.1 | 4198 | AT-less type I PKS | MisF [AKQ22696.1] | 40/56 |
| WP_106528587.1 | 5976 | AT-less type I PKS | BryA [ABM63537.1] | 44/59 |
| WP_106528588.1 | 1733 | AT-less type I PKS | BaeL [CAG23958.2] | 53/67 |
| WP_106528589.1 | 3613 | AT-less type I PKS | Bat2 [ADD82940.1] | 44/59 |
| WP_106528590.1 | 1429 | AT-less type I PKS | RhiF [CAL69894.1] | 44/63 |
| WP_106528828.1 | 170 | antitermination factor | ElaA [AEC04347.1] | 52/76 |
| WP_106528829.1 | 796 | S-malonyltransferase | ElaB [AEC04348.1] | 56/70 |
| WP_106528591.1 | 321 | acyltransferase | ElaC [AEC04349.1] | 47/64 |
| WP_106528592.1 | 96 | hypothetical protein | ElaD [AEC04350.1] | 55/69 |
| WP_106528593.1 | 79 | acyl carrier protein | ElaE [AEC04351.1] | 61/78 |
| WP_106528594.1 | 408 | beta-ketoacyl synthase | TstN [AGN11888.1] | 60/74 |
| WP_106528595.1 | 421 | HMG-CoA synthase | BatC [ADD82944.1] | 73/87 |
| WP_106528596.1 | 249 | enoyl-CoA hydratase | BaeH [CAG23955.1] | 40/72 |
| WP_106528597.1 | 249 | enoyl-CoA hydratase | ElaN [AEC04360.1] | 65/80 |
| WP_106528598.1 | 281 | methyltransferase | Dor10 [ACY01395.1] | 34/48 |
| WP_106528599.1 | 345 | cysteine S-methyltransferase | / | / |
| WP_106528600.1 | 346 | AraC family transcriptional regulator | / | / |

[a] Number of amino acids

**Table S159.** Predicted functions of ORFs in the NZ_QICS01000001.1 containing WP_110290069.1

| gene | aa [a] | putative function | Protein homologue | %identity/ %similarity |
|---|---|---|---|---|
| WP_110290054.1 | 265 | acetyl-CoA carboxylase carboxyltransferase | / | / |
| WP_110290055.1 | 281 | acetyl-CoA carboxylase carboxyl transferase | ThaB [ABC35022.1] | 48/64 |
| WP_110290056.1 | 175 | hypothetical protein | / | / |
| WP_110290057.1 | 448 | ATP-grasp domain-containing protein | / | / |
| WP_110290058.1 | 757 | AT/Ox | DifA [CAG23974.1] | 56/74 |
| WP_094376764.1 | 80 | acyl carrier protein | ElaE [AEC04351.1] | 40/62 |
| WP_110290059.1 | 437 | beta-ketoacyl synthase | MxnD [AGS77284.1] | 45/63 |
| WP_110290060.1 | 3375 | AT-less type I PKS | BaeN [CAG23960.2] | 42/61 |
| WP_110290061.1 | 2871 | AT-less type I PKS | RhiC [CAL69890.1] | 35/51 |
| WP_110290062.1 | 1682 | AT-less type I PKS | DszB [AAY32965.1] | 35/54 |
| WP_110290063.1 | 3288 | AT-less type I PKS | RizB [CCA89326.1] | 34/54 |
| WP_110290064.1 | 3081 | AT-less type I PKS | DszA [AAY32964.1] | 37/56 |
| WP_110290065.1 | 285 | hypothetical protein | / | / |
| WP_094376772.1 | 418 | hypothetical protein | / | / |
| WP_094376773.1 | 215 | HAD family phosphatase | OzmW [ABS90484.1] | 24/44 |
| WP_110290066.1 | 324 | acyltransferase | BaeD [CAG23951.1] | 38/55 |
| WP_094376775.1 | 140 | hypothetical protein | / | / |
| WP_094376776.1 | 66 | hypothetical protein | / | / |
| WP_094376777.1 | 192 | etR/AcrR family transcriptional regulator | / | / |
| WP_110290067.1 | 302 | NAD-dependent epimerase | / | / |
| WP_110290068.1 | 418 | glycosyltransferase | SorF [ADN68481.1] | 36/51 |
| WP_094376780.1 | 254 | enoyl-CoA hydratase | BaeH [CAG23955.1] | 52/69 |
| WP_110290069.1 | 419 | HMG-CoA synthase | CylF [ARU81120.1] | 68/83 |
| WP_094376782.1 | 199 | hypothetical protein | / | / |
| WP_110290070.1 | 248 | DNA-binding response regulator | / | / |
| WP_094376784.1 | 437 | ATP-binding protein | / | / |

[a] Number of amino acids

**Table S160.** Predicted functions of ORFs in the NZ_QICS01000001.1 containing WP_111432520.1

| gene | aa [a] | putative function | Protein homologue | %identity/ %similarity |
|---|---|---|---|---|
| WP_111432507.1 | 396 | CoA transferase | enacyloxin [ABI91461.1] | 40/55 |
| WP_111432508.1 | 442 | MFS transporter | Disorazols [AAY32956.1] | 52/85 |
| WP_111432509.1 | 102 | hypothetical protein | / | / |
| WP_111432510.1 | 268 | alpha/beta hydrolase | / | / |
| WP_111432511.1 | 428 | MFS transporter | LnmY [AAN85538.1] | 25/40 |
| WP_111432512.1 | 376 | hypothetical protein | / | / |
| WP_111432513.1 | 644 | transcriptional regulator | Disorazols [AAY32957.1] | 47/62 |
| WP_111432514.1 | 481 | bifunctional enoyl-CoA hydratase /phosphate acetyltransferase | / | / |
| WP_111432515.1 | 438 | acetate/propionate family kinase | / | / |
| WP_111432516.1 | 364 | acyltransferase | BaeD [CAG23951.1] | 35/56 |
| WP_111432643.1 | 271 | PPTase | TmlN [CBK62709.1] | 35/55 |
| WP_111432517.1 | 286 | Ser/Thr protein phosphatase | ThaC [ABC35295.1] | 32/42 |
| WP_111432518.1 | 310 | FkbM family methyltransferase | CalA [BAP05589.1] | 35/53 |
| WP_111432519.1 | 415 | cytochrome P450 | ElaG [AEC04353.1] | 32/49 |
| WP_111432520.1 | 418 | HMG-CoA synthase | ThaK [ABC34601.1] | 61/74 |
| WP_111432521.1 | 81 | acyl carrier protein | CalX [BAP05574.1] | 53/71 |
| WP_111432522.1 | 410 | beta-ketoacyl synthase | TstN [AGN11888.1] | 52/66 |
| WP_111432523.1 | 252 | enoyl-CoA hydratas | BaeH [CAG23955.1] | 51/65 |
| WP_111432524.1 | 664 | asparagine synthase | SmdH [CCC21122.1] | 59/72 |
| WP_111432525.1 | 84 | acyl carrier protein | ChxC [AFO59864.1] | 50/75 |
| WP_111432526.1 | 543 | yclic peptide export ABC transporter | / | / |
| WP_111432527.1 | 547 | ATP-binding cassette domain | / | / |
| WP_111432644.1 | 809 | AT/Ox | CalY [BAP05573.1] | 49/66 |
| WP_111432528.1 | 6852 | AT-less type I PKS | Dor5 [ACY01390.1] | 45/56 |
| WP_111432529.1 | 438 | monooxygenase | PedG [AAS47561.1] | 63/77 |
| WP_111432530.1 | 6732 | hybrid PKS/NRPS | TaO [ABF92489.1] | 40/54 |
| WP_111432531.1 | 290 | prohibitin family protein | / | / |
| WP_111432532.1 | 462 | hypothetical protein | / | / |

[a] Number of amino acids

**Table S161.** Predicted functions of ORFs in the NZ_QMIG01000028.1 containing WP_112260017.1

| gene | aa [a] | putative function | Protein homologue | %identity/ %similarity |
|---|---|---|---|---|
| WP_112260008.1 | 145 | deazaflavin-dependent oxidoreductase | / | / |
| WP_112260009.1 | 275 | exodeoxyribonuclease III | / | / |
| WP_112260010.1 | 174 | DUF98 domain-containing protein | ElaI [AEC04355.1] | 24/43 |
| WP_112260011.1 | 225 | DNA-binding response regulator | smdD [CCC21118.1] | 33/50 |
| WP_112260025.1 | 278 | Ser/Thr protein phosphatase | ThaC [ABC35295.1] | 27/41 |
| WP_112260012.1 | 262 | PPTase | Mis12 [AKQ22701.1] | 34/50 |
| WP_112260013.1 | 334 | acyltransferase | BaeD [CAG23951.1] | 37/54 |
| WP_112260014.1 | 287 | S-malonyltransferase | BaeE [CAG23952.1] | 53/71 |
| WP_112260015.1 | 84 | acyl carrier protein | PsyL [ADA82592.1] | 40/69 |
| WP_112260016.1 | 434 | beta-ketoacyl synthase | CorD [ADI59526.1] | 48/60 |
| WP_112260017.1 | 419 | HMG-CoA synthase | BatC [ADD82944.1] | 68/81 |
| WP_112260018.1 | 261 | enoyl-CoA hydratase | BaeH [CAG23955.1] | 55/68 |
| WP_112260019.1 | 254 | enoyl-CoA hydratase | BatE [ADD82946.1] | 57/74 |
| WP_112260020.1 | 3985 | AT-less type I PKS | RhiE [CAL69893.1] | 38/53 |

[a] Number of amino acids

**Table S162.** Predicted functions of ORFs in the NZ_QFFJ01000002.1 containing WP_113618604.1

| gene | aa [a] | putative function | Protein homologue | %identity/ %similarity |
|---|---|---|---|---|
| WP_113618595.1 | 231 | type 1 glutamine amidotransferase | / | / |
| WP_113618596.1 | 312 | NAD-dependent epimerase | / | / |
| WP_113618597.1 | 243 | PPTase | Mis12 [AKQ22701.1] | 36/56 |
| WP_113618598.1 | 5719 | AT-less type I PKS | BaeN [CAG23960.2] | 35/53 |
| WP_113618599.1 | 5673 | AT-less type I PKS | BaeL [CAG23958.2] | 43/59 |
| WP_113618600.1 | 6211 | NRPS | DifI [CAJ57409.1] | 33/51 |
| WP_113618601.1 | 758 | AT/Ox | DifA [CAG23974.1] | 52/72 |
| WP_113618602.1 | 78 | acyl carrier protein | AcpK [CAG23953.1] | 41/65 |
| WP_113618603.1 | 420 | beta-ketoacyl synthase | CylE [ARU81119.1] | 42/59 |
| WP_113618604.1 | 415 | HMG-CoA synthase | JamH [AAS98779.1] | 66/77 |
| WP_113618605.1 | 264 | enoyl-CoA hydratase | CylG [ARU81121.1] | 51/68 |
| WP_113618606.1 | 248 | enoyl-CoA hydratase | BatE [ADD82946.1] | 59/77 |
| WP_113618607.1 | 317 | acyltransferase | BaeD [CAG23951.1] | 34/54 |
| WP_113618608.1 | 275 | alpha/beta hydrolase | OocA [AFX60323.1] | 32/50 |
| WP_113618609.1 | 113 | type III effector protein | / | / |
| WP_113618610.1 | 1043 | AsmA family protein | / | / |

[a] Number of amino acids

**Table S163.** Predicted functions of ORFs in the NZ_QFFJ01000002.1 containing WP_113672097.1 and WP_113672099.1

| gene | aa [a] | putative function | Protein homologue | %identity/ %similarity |
|---|---|---|---|---|
| WP_113672087.1 | 390 | iron-containing alcohol dehydrogenase | / | / |
| WP_113672088.1 | 151 | peptide deformylase | / | / |
| WP_113672089.1 | 971 | hypothetical protein | BaeL [CAG23958.2] | 37/56 |
| WP_113672090.1 | 80 | acyl carrier protein | SmdG [CCC21121.1] | 41/65 |
| WP_113672091.1 | 255 | methyltransferase | / | / |
| WP_113672092.1 | 472 | AMP-dependent synthetase | MalI [ABC34723.1] | 22/43 |
| WP_113672093.1 | 3569 | AT-less type I PKS | MisF [AKQ22696.1] | 42/60 |
| WP_113672094.1 | 1587 | AT-less type I PKS | BasE [ERM18797.1] | 51/71 |
| WP_113672095.1 | 3666 | AT-less type I PKS | BaeM [CAG23959.2] | 42/66 |
| WP_113674435.1 | 249 | enoyl-CoA hydratase | BatE [ADD82946.1] | 67/81 |
| WP_113672096.1 | 254 | enoyl-CoA hydratase | BaeH [CAG23955.1] | 57/77 |
| WP_113672097.1 | 419 | HMG-CoA synthase | BaeG [CAG23954.2] | 75/84 |
| WP_113672098.1 | 411 | beta-ketoacyl synthase | BatB [ADD82943.1] | 61/77 |
| WP_113672099.1 | 417 | HMG-CoA synthase | BatC [ADD82944.1] | 66/80 |
| WP_113672100.1 | 82 | acyl carrier protein | BatA [ADD82942.1] | 54/76 |
| WP_113672101.1 | 613 | S-malonyltransferase | BasH [ERM18800.1] | 46/68 |
| WP_113672102.1 | 248 | PPTase | Mis12 [AKQ22701.1] | 33/45 |
| WP_113672103.1 | 225 | MBL fold metallo-hydrolase | BaeB [CAG23949.2] | 47/66 |
| WP_113672104.1 | 274 | acetyl-CoA carboxylase carboxyltransferase | ThaB [ABC35022.1] | 48/64 |
| WP_113672105.1 | 283 | acetyl-CoA carboxylase carboxyltransferase | SnbS [CBW45762.1] | 26/46 |
| WP_113672106.1 | 442 | acetyl-CoA carboxylase biotin carboxylase | / | / |
| WP_113672107.1 | 152 | hypothetical protein | / | / |

[a] Number of amino acids

**Table S164.** Predicted functions of ORFs in the NZ_QLKQ01000044.1 containing WP_114859988.1

| gene [a] | aa [b] | putative function | Protein homologue | %identity/ %similarity |
|---|---|---|---|---|
| WP_114859977.1 | 4037 | AT-less type I PKS | PedH [AAS47562.1] | 52/65 |
| WP_114859978.1 | 433 | monooxygenase | PedG [AAS47561.1] | 76/87 |
| ORF1 | 8485 | AT-less type I PKS | pedF [AAS47564.1] | 53/66 |
| WP_114859979.1 | 267 | methyltransferase | pedE [AAS47560.1] | 60/76 |
| WP_114859980.1 | 207 | hypothetical protein | / | / |
| WP_114859981.1 | 232 | PPTase | dipB [AGS06837.1] | 36/55 |
| WP_114859988.1 | 426 | HMG-CoA synthase | PedP [AAW33975.1] | 70/82 |
| WP_114859982.1 | 588 | cyclic peptide export ABC transporter | / | / |
| WP_114859983.1 | 548 | cyclic peptide export ABC transporter | lkcJ [BAC76467.1] | 30/45 |
| WP_114859984.1 | 4933 | AT-less type I PKS | PedI [AAR19304.1] | 50/63 |
| WP_114859985.1 | 369 | LLM class flavin-dependent oxidoreductase | PedJ [AAR19305.1] | 70/83 |
| WP_114859986.1 | 76 | hypothetical protein | MisG [AKQ22695.1] | 64/74 |

[a] ORFs are proteins without protein ID provided by NCBI [b] Number of amino acids

**Table S165.** Predicted functions of ORFs in the NZ_QTSU01000001.1 containing WP_115857952.1

| gene | aa [a] | putative function | Protein homologue | %identity/ %similarity |
|---|---|---|---|---|
| WP_115857944.1 | 301 | hypothetical protein | / | / |
| WP_115857945.1 | 410 | hypothetical protein | / | / |
| WP_115857946.1 | 253 | Ser/Thr protein phosphatase | ThaC [ABC35295.1] | 4463 |
| WP_115857947.1 | 293 | PPTase | BatI [ADD82950.1] | 52/64 |
| WP_115857948.1 | 3522 | AT-less type I PKS | OocJ [AFX60332.1] | 42/54 |
| WP_115857949.1 | 3199 | AT-less type I PKS | OocJ [AFX60332.1] | 43/56 |
| WP_115857950.1 | 4292 | AT-less type I PKS | DszA [AAY32964.1] | 35/49 |
| WP_115857951.1 | 1359 | AT-less type I PKS | DifF [CAG23977.1] | 37/52 |
| WP_115857952.1 | 419 | HMG-CoA synthase | BatC [ADD82944.1] | 70/81 |
| WP_115857953.1 | 264 | enoyl-CoA hydratase | OocD [AFX60326.1] | 52/71 |
| WP_115857954.1 | 250 | enoyl-CoA hydratase | BatE [ADD82946.1] | 61/75 |
| WP_115857955.1 | 79 | acyl carrier protein | CalX [BAP05574.1] | 52/77 |
| WP_115857956.1 | 412 | beta-ketoacyl synthase | TstN [AGN11888.1] | 65/76 |
| WP_115857957.1 | 403 | S-malonyltransferase | TstO [AGN11889.1] | 51/64 |
| WP_115857958.1 | 344 | acyltransferase | BaeD [CAG23951.1] | 35/53 |
| WP_115857959.1 | 141 | hotdog fold thioesterase | / | / |
| WP_115857960.1 | 364 | NAD(P)-dependent alcohol dehydrogenase | enacyloxin [ABI91459.1] | 32/48 |
| WP_115857961.1 | 3499 | AT-less type I PKS | Bat3 [ADD82941.1] | 39/55 |
| WP_115857962.1 | 469 | long-chain fatty acid--CoA ligase | DifD [CAJ57407.1] | 48/66 |
| WP_115857963.1 | 251 | SDR family oxidoreductase | DifE [CAG23976.1] | 47/68 |
| WP_115857964.1 | 81 | hypothetical protein | DifC [CAJ57406.1] | 41/66 |
| WP_115857965.1 | 1403 | AT-less type I PKS | ElaQ [AEC04363.1] | 37/54 |
| WP_115857966.1 | 527 | hypothetical protein | / | / |
| WP_115857967.1 | 582 | aminotransferase | BatP [ADD82957.1] | 27/43 |
| WP_115857968.1 | 1253 | AT-less type I PKS | SorA [ADN68476.1] | 31/48 |
| WP_115857969.1 | 824 | hypothetical protein | / | / |

[a] Number of amino acids

**Table S166.** Predicted functions of ORFs in the NZ_QNVT01000021.1 containing WP_115972399.1

| gene | aa [a] | putative function | Protein homologue | %identity/ %similarity |
|---|---|---|---|---|
| WP_115972392.1 | 64 | cold shock domain-containing protein | / | / |
| WP_115972393.1 | 153 | MaoC family dehydratase | / | / |
| WP_115972394.1 | 190 | DUF892 family protein | / | / |
| WP_115972395.1 | 259 | PPTase | Mis12 [AKQ22701.1] | 31/52 |
| WP_115972396.1 | 331 | FAA hydrolase family protein | / | / |
| WP_115972397.1 | 274 | alpha/beta fold hydrolase | / | / |
| WP_115972398.1 | 417 | monooxygenase | PedG [AAS47561.1] | 30/51 |
| WP_115972415.1 | 262 | enoyl-CoA hydratase | CylG [ARU81121.1] | 49//62 |
| WP_115972399.1 | 414 | HMG-CoA synthase | JamH [AAS98779.1] | 64/78 |
| WP_115972400.1 | 423 | beta-ketoacyl synthase | CylE [ARU81119.1] | 38/58 |
| WP_115972401.1 | 82 | acyl carrier protein | TaB [ABF90753.1] | 44/71 |
| WP_115972402.1 | 744 | AT/Ox | DifA [CAG23974.1] | 52/70 |
| WP_115972403.1 | 3121 | hybrid PKS/NRPS | DifI [CAJ57409.1] | 42/56 |
| WP_115972404.1 | 3847 | AT-less type I PKS | ElaQ [AEC04363.1] | 33/51 |
| WP_115972405.1 | 3071 | AT-less type I PKS | MisF [AKQ22696.1] | 34/51 |
| WP_115972406.1 | 429 | flavin-dependent monooxygenase | OocK [AFX60333.1] | 56/73 |
| WP_115972407.1 | 1259 | AT-less type I PKS | OnnB [AAV97870.1] | 43/59 |
| WP_115972408.1 | 109 | DUF4280 domain-containing protein | / | / |
| WP_115972409.1 | 214 | hypothetical protein | / | / |
| WP_115972410.1 | 349 | hypothetical protein | / | / |

[a] Number of amino acids

**Table S167.** Predicted functions of ORFs in the NZ_LSRW01000045.1 containing WP_116811145.1

| gene | aa [a] | putative function | Protein homologue | %identity/ %similarity |
|---|---|---|---|---|
| WP_116811132.1 | 1460 | AT-less type I PKS | MisF [AKQ22696.1] | 46/63 |
| WP_116811133.1 | 407 | hypothetical protein | / | / |
| WP_116811134.1 | 271 | hypothetical protein | / | / |
| WP_116811135.1 | 4497 | AT-less type I PKS | MisF [AKQ22696.1] | 43/58 |
| WP_116811136.1 | 591 | AT-less type I PKS | MisF [AKQ22696.1] | 39/58 |
| WP_116811137.1 | 4917 | AT-less type I PKS | BaeM [CAG23959.2] | 35/52 |
| WP_116811138.1 | 3910 | AT-less type I PKS | MisF [AKQ22696.1] | 38/54 |
| WP_116811139.1 | 856 | hypothetical protein | / | / |
| WP_116811140.1 | 294 | outer membrane lipoprotein | / | / |
| WP_116811141.1 | 443 | hypothetical protein | / | / |
| WP_116811202.1 | 323 | NAD(P)-dependent alcohol dehydrogenase | kirromycin [CAN89616.1] | 31/45 |
| WP_116811142.1 | 423 | hypothetical protein | / | / |
| WP_116811143.1 | 342 | hypothetical protein | / | / |
| WP_116811144.1 | 380 | S-malonyltransferase | TstO [AGN11889.1] | 58/71 |
| WP_116811145.1 | 420 | HMG-CoA synthase | BatC [ADD82944.1] | 68/81 |
| WP_116811146.1 | 264 | enoyl-CoA hydratase | BaeH [CAG23955.1] | 57/70 |
| WP_116811147.1 | 248 | enoyl-CoA hydratase | BatE [ADD82946.1] | 66/82 |
| WP_116811148.1 | 79 | acyl carrier protein | TaB [ABF90753.1] | 57/69 |
| WP_116811149.1 | 414 | beta-ketoacyl synthase | NspH [ADA69244.1] | 62/76 |
| WP_116811150.1 | 178 | UpxY family transcription antiterminator | TaA [ABF91060.1] | 33/51 |
| WP_116811151.1 | 209 | GNAT family N-acetyltransferase | MupI [AAK28505.1] | 34/54 |
| WP_116811152.1 | 131 | DUF4902 domain-containing protein | / | / |
| WP_116811153.1 | 123 | hypothetical protein | enacyloxin [ABI91455.1] | 30/45 |
| WP_116811154.1 | 124 | LuxR family transcriptional regulator | enacyloxin [ABI91455.1] | 39/55 |
| WP_116811155.1 | 463 | cysteine--tRNA ligase | / | / |
| WP_116811156.1 | 306 | lytic murein transglycosylase | / | / |

[a] Number of amino acids

**Table S168.** Predicted functions of ORFs in the NZ_LSRW01000045.1 containing WP_117189355.1

| gene | aa [a] | putative function | Protein homologue | %identity/ %similarity |
|---|---|---|---|---|
| WP_117189350.1 | 192 | hypothetical protein | / | / |
| WP_117189351.1 | 459 | PfaD family polyunsaturated fatty acid | CorA [ADI59523.1] | 46/63 |
| WP_117189352.1 | 250 | PPTase | LtmL [ACY01405.1] | 48/56 |
| WP_117189353.1 | 241 | SDR family oxidoreductase | DifE [CAG23976.1] | 39/60 |
| WP_117189354.1 | 110 | acyl carrier protein | JamF [AAS98799.1] | 37/64 |
| WP_117189371.1 | 234 | enoyl-CoA hydratase | BatE [ADD82946.1] | 52/68 |
| WP_117189355.1 | 421 | HMG-CoA synthase | ThaK [ABC34601.1] | 64/78 |
| WP_117189356.1 | 1526 | AT-less type I PKS | JamL [AAS98783.1] | 41/56 |
| WP_117189357.1 | 2572 | AT-less type I PKS | kirAIV [CAN89634.1] | 41/53 |
| WP_117189358.1 | 3663 | AT-less type I PKS | BaeN [CAG23960.2] | 30/47 |
| WP_117189359.1 | 3984 | AT-less type I PKS | LglD [AIU36100.1] | 30/47 |

[a] Number of amino acids

**Table S169.** Predicted functions of ORFs in the NZ_CP031841.1 containing WP_117412785.1

| gene | aa [a] | putative function | Protein homologue | %identity/ %similarity |
|---|---|---|---|---|
| WP_117412777.1 | 486 | prohibitin family protein | / | / |
| WP_117412778.1 | 568 | ATP-binding cassette domain | / | / |
| WP_117412779.1 | 150 | hypothetical protein | CalV [BAP05576.1] | 39/51 |
| WP_117412780.1 | 723 | ATP-binding cassette domain | CalU [BAP05577.1] | 38/57 |
| WP_117412781.1 | 142 | PPTase | / | / |
| WP_117412782.1 | 804 | AT/Ox | DifA [CAG23974.1] | 47/65 |
| WP_117412783.1 | 86 | acyl carrier protein | DipF [AGS06833.1] | 36/64 |
| WP_117412784.1 | 413 | beta-ketoacyl synthase | MxnD [AGS77284.1] | 39/54 |
| WP_117412785.1 | 411 | HMG-CoA synthase | MxnE [AGS77285.1] | 52/66 |
| WP_117412786.1 | 254 | enoyl-CoA hydratase | MxnF [AGS77286.1] | 36/49 |
| WP_117412787.1 | 120 | hypothetical protein | / | / |
| WP_117412788.1 | 896 | polyketide synthase | MisC [AKQ22699.1] | 33/46 |
| WP_117412789.1 | 81 | hypothetical protein | / | / |
| WP_117412790.1 | 675 | hypothetical protein | CorK [ADI59533.1] | 50/60 |
| WP_117412791.1 | 885 | AT-less type I PKS | OzmN [ABS90475.1] | 33/43 |
| WP_117412792.1 | 2745 | AT-less type I PKS | TaO [ABF92489.1] | 37/51 |
| WP_117412793.1 | 1116 | AT-less type I PKS | RizB [CCA89326.1] | 34/47 |
| WP_117412794.1 | 1286 | AT-less type I PKS | BonD [AFN27483.1] | 3748 |
| WP_117412795.1 | 2978 | hybrid PKS/NRPS | OzmH [ABS90470.1] | 39/51 |
| WP_117412796.1 | 1142 | NRPS | CalI [BAP05597.1] | 40/54 |
| WP_117412797.1 | 315 | hypothetical protein | / | / |
| WP_117412798.1 | 272 | hypothetical protein | / | / |

[a] Number of amino acids

**Table S170.** Predicted functions of ORFs in the NZ_KZ984561.1 containing WP_119294050.1

| gene | aa [a] | putative function | Protein homologue | %identity/ %similarity |
|---|---|---|---|---|
| WP_119294038.1 | 598 | acyl-CoA dehydrogenase | / | / |
| WP_119294039.1 | 318 | cyclase | / | / |
| WP_119294040.1 | 95 | acyl carrier protein | snaX [CBW45732.1] | 69/89 |
| WP_119294041.1 | 243 | alpha/beta fold hydrolase | snaP [CBW45734.1] | 78/87 |
| WP_119294042.1 | 331 | hypothetical protein | PapR2 [CBW45736.1] | 79/85 |
| WP_119294043.1 | 173 | flavin reductase | snaC [CBW45737.1] | 62/74 |
| WP_119294044.1 | 285 | PPTase | snaN [CBW45738.1] | 75/83 |
| WP_119294045.1 | 296 | S-malonyltransferase | SnaM [CBW45739.1] | 79/84 |
| WP_119294046.1 | 2549 | NRPS | VirH [BAF50720.1] | 64/71 |
| WP_119294047.1 | 2069 | AT-less type I PKS | MisF [AKQ22696.1] | 35/50 |
| WP_119294048.1 | 258 | enoyl-CoA hydratase | snaK [CBW45743.1] | 81/90 |
| WP_119294049.1 | 244 | enoyl-CoA hydratase | SnaJ [CBW45744.1] | 76/80 |
| WP_119294050.1 | 416 | HMG-CoA synthase | SnaI [CBW45745.1] | 88/93 |
| WP_119294051.1 | 422 | beta-ketoacyl synthase | SnaH [CBW45746.1] | 81/85 |
| WP_119294052.1 | 83 | acyl carrier protein | SnaG [CBW45747.1] | 71/83 |
| WP_119294053.1 | 2417 | AT-less type I PKS | SnaE2 [CBW45748.1] | 68/74 |
| WP_119294054.1 | 5151 | hybrid PKS/NRPS | VirA [BAF50727.1] | 58/65 |
| WP_119294055.1 | 663 | alpha-keto acid dehydrogenase | SnaF [CBW45750.1] | 78/85 |
| WP_119294056.1 | 317 | HemK family protein methyltransferase | papM [CBW45752.1] | 71/77 |
| WP_119294057.1 | 113 | chorismate mutase | papB [CBW45753.1] | 74/81 |
| WP_119294058.1 | 288 | prephenate dehydrogenase | papC [CBW45754.1] | 62/70 |
| WP_119294059.1 | 713 | aminodeoxychorismate synthase | papA [CBW45755.1] | 75/81 |
| WP_119294112.1 | 341 | lysine cyclodeaminase | PipA [CBW45757.1] | 78/84 |
| WP_119294060.1 | 550 | 2,3-dihydroxybenzoate-AMP ligase | SnbA [CBW45758.1] | 71/81 |
| WP_119294061.1 | 514 | MFS transporter | SnbR [CBW45761.1] | 80/86 |
| WP_119294062.1 | 229 | TetR family transcriptional regulator | leinamycin [AAN85544.1] | 71/79 |
| WP_119294063.1 | 545 | ATP-binding cassette domain | leinamycin [AAN85545.1] | 87/92 |
| WP_119294064.1 | 173 | hypothetical protein | / | / |
| WP_119294065.1 | 427 | FAD-dependent oxidoreductase | / | / |

[a] Number of amino acids

**Table S171.** Predicted functions of ORFs in the CP001344.1 containing ACL45227.1

| gene | aa[a] | putative function | Protein homologue | %identity/ %similarity |
|---|---|---|---|---|
| ACL45212.1 | 211 | Crp/Fnr family | / | / |
| ACL45213.1 | 467 | tetratricopeptide repeat protein | / | / |
| ACL45214.1 | 173 | phycocyanin, β subunit | / | / |
| ACL45215.1 | 163 | phycocyanin, alpha subunit | / | / |
| ACL45216.1 | 327 | rhodanese domain | / | / |
| ACL45217.1 | 170 | hypothetical protein | / | / |
| ACL45218.1 | 305 | fatty acid desaturase | JamB [AAS98775.1] | 45/64 |
| ACL45219.1 | 586 | AMP-dependent synthetase | JamA [AAS98774.1] | 45/64 |
| ACL45220.1 | 90 | phosphopantetheine-binding | JamC [AAS98798.1] | 31/62 |
| ACL45221.1 | 286 | fatty acid desaturase | / | / |
| ACL45222.1 | 376 | fatty acid desaturase | BatZ [ADD82967.1] | 19/38 |
| ACL45223.1 | 312 | stearoyl-CoA 9-desaturase | JamB [AAS98775.1] | 52/70 |
| ACL45224.1 | 609 | flavin-containing monooxygenase | OocK [AFX60333.1] | 27/41 |
| ACL45225.1 | 2462 | canonical type I PKS | JamJ [AAS98781.1] | 48/64 |
| ACL45226.1 | 258 | enoyl-CoA hydratase | JamI [AAS98780.1] | 57/74 |
| ACL45227.1 | 419 | HMG-CoA synthase | JamH [AAS98779.1] | 71/84 |
| ACL45228.1 | 415 | beta-ketoacyl synthase | JamG [AAS98778.1] | 60/75 |
| ACL45229.1 | 85 | acyl carrier protein | acpK [CAG23953.1] | 51/77 |
| ACL45230.1 | 1208 | canonical type I PKS | JamE [AAS98777.1] | 44/58 |
| ACL45231.1 | 34 | hypothetical protein | / | / |
| ACL45232.1 | 496 | protoporphyrinogen oxidase | / | / |
| ACL45233.1 | 399 | ABC exporter membrane fusion protein | / | / |
| ACL45234.1 | 395 | DevC protein | LnmR [AAN85531.1] | 34/50 |
| ACL45235.1 | 453 | phytochrome sensor protein | / | / |
| ACL45236.1 | 189 | calcium-binding EF-hand-containing protein | / | / |
| ACL45237.1 | 356 | integrase domain protein SAM | / | / |

[a] Number of amino acids

**Table S172.** Predicted functions of ORFs in the CP003987.1 containing AJC59428.1

| gene | aa[a] | putative function | Protein homologue | %identity/ %similarity |
|---|---|---|---|---|
| AJC59413.1 | 263 | oxidoreductase | leinamycin [AAN85505.1] | 30/51 |
| AJC59414.1 | 215 | regulatory protein C TetR | / | / |
| AJC59415.1 | 298 | integral membrane protein | / | / |
| AJC59416.1 | 234 | ArsR family transcriptional regulator | / | / |
| AJC59417.1 | 428 | major facilitator transporter | kirromycin [CAN89665.1] | 42/53 |
| AJC59418.1 | 251 | phosphate regulator | / | / |
| AJC59419.1 | 135 | hypothetical protein | / | / |
| AJC59420.1 | 476 | cytochrome P450 | nosperin [ADA69248.1] | 29/46 |
| AJC59421.1 | 554 | ABC transporter ATP-binding protein | leinamycin [AAN85547.1] | 70/80 |
| AJC59422.1 | 225 | haloacid dehalogenase | / | / |
| AJC59423.1 | 81 | hypothetical protein | / | / |
| AJC59425.1 | 585 | hybrid polyketide synthase/NRPS | VirA [BAF50727.1] | 53/66 |
| AJC59426.1 | 83 | acyl carrier protein | PsyL [ADA82592.1] | 40/68 |
| AJC59427.1 | 420 | beta-ketoacyl synthase | VirB [BAF50726.1] | 51/63 |
| AJC59428.1 | 411 | HMG-CoA synthase | SnaI [CBW45745.1] | 57/72 |
| AJC59429.1 | 251 | enoyl-CoA hydratase | VirD [BAF50724.1] | 45/54 |
| AJC59430.1 | 262 | enoyl-CoA hydratase | VirE [BAF50723.1] | 46/57 |
| AJC59431.1 | 275 | acyltransferase | VirI [BAF50719.1] | 54/63 |
| AJC59432.1 | 2101 | canonical type I PKS | JamL [AAS98783.1] | 33/49 |
| AJC59433.1 | 212 | carboxymuconolactone decarboxylase | / | / |
| AJC59434.1 | 1340 | canonical type I PKS | kirAVI [CAN89637.1] | 48/60 |
| AJC59435.1 | 170 | hypothetical protein | / | / |

[a] Number of amino acids

**Table S173.** Predicted functions of ORFs in the NC_017093.1 containing WP_014445147.1

| gene | aa[a] | putative function | Protein homologue | %identity/ %similarity |
|---|---|---|---|---|
| WP_014445137.1 | 184 | TetR/AcrR transcriptional regulator | / | / |
| WP_014445138.1 | 465 | MFS transporter | SnbR [CBW45761.1] | 37/55 |
| WP_014445139.1 | 166 | hypothetical protein | / | / |
| WP_014445140.1 | 642 | membrane protein | / | / |
| WP_014445141.1 | 161 | hypothetical protein | / | / |
| WP_014445142.1 | 256 | LLM class oxidoreductase | Cycloheximide [CCC21141.1] | 38/59 |
| WP_083888691.1 | 153 | DUF385 domain | / | / |
| WP_014445144.1 | 593 | polyketide synthase | VirA [BAF50727.1] | 43/57 |
| WP_014445145.1 | 258 | enoyl-CoA hydratase | VirE [BAF50723.1] | 44/59 |
| WP_051042126.1 | 248 | enoyl-CoA hydratase | VirD [BAF50724.1] | 49/56 |
| WP_014445147.1 | 407 | HMG-CoA synthase | LnmM [AAN85526.1] | 55/71 |
| WP_014445148.1 | 81 | acyl carrier protein | LnmL [AAN85525.1] | 50/63 |
| WP_014445149.1 | 315 | AT/DC | LnmK [AAN85524.1] | 45/62 |
| WP_014445150.1 | 95 | acyl carrier protein | VirA [BAF50727.1] | 49/59 |
| WP_014445151.1 | 403 | ketosynthase chain-length factor | MxnD [AGS77284.1] | 28/46 |
| WP_014445152.1 | 422 | beta-ketoacyl synthase | MxnD [AGS77284.1] | 32/46 |
| WP_014445153.1 | 78 | acyl carrier protein | ChiB [AAY89049.1] | 37/48 |
| WP_014445154.1 | 374 | 3-oxoacyl-ACP synthase | / | / |
| WP_014445155.1 | 246 | oxidoreductase | DifE [CAG23976.1] | 34/51 |
| WP_041831255.1 | 345 | luciferase-like monooxygenase | Cycloheximide [CCC21141.1] | 34/46 |
| WP_014445157.1 | 171 | hemerythrin domain | / | / |
| WP_014445158.1 | 381 | hydrolase | / | / |
| WP_014445159.1 | 283 | regulatory protein | papR1 [CBW45751.1] | 48/56 |
| WP_014445160.1 | 307 | MBL fold metallo-hydrolase | Psymberin [ADA82578.1] | 58/69 |
| WP_014445161.1 | 149 | hypothetical protein | / | / |
| WP_014445162.1 | 303 | nitroreductase family | / | / |
| WP_014445163.1 | 402 | acyltransferase | / | / |
| WP_014445164.1 | 399 | cytochrome P450 | / | / |

[a] Number of amino acids

**Table S174.** Predicted functions of ORFs in the NZ_CP011340.1 containing WP_005321723.1

| gene | aa[a] | putative function | Protein homologue | %identity/ %similarity |
|---|---|---|---|---|
| WP_014445137.1 | 184 | TetR/AcrR transcriptional regulator | / | / |
| WP_014445138.1 | 465 | MFS transporter | SnbR [CBW45761.1] | 37/55 |
| WP_014445139.1 | 166 | hypothetical protein | / | / |
| WP_014445140.1 | 642 | membrane protein | / | / |
| WP_014445141.1 | 161 | hypothetical protein | / | / |
| WP_014445142.1 | 256 | LLM class oxidoreductase | Cycloheximide [CCC21141.1] | 38/59 |
| WP_083888691.1 | 153 | DUF385 domain | / | / |
| WP_014445144.1 | 593 | polyketide synthase | VirA [BAF50727.1] | 43/57 |
| WP_014445145.1 | 258 | enoyl-CoA hydratase | VirE [BAF50723.1] | 44/59 |
| WP_051042126.1 | 248 | enoyl-CoA hydratase | VirD [BAF50724.1] | 49/56 |
| WP_014445147.1 | 407 | HMG-CoA synthase | LnmM [AAN85526.1] | 55/71 |
| WP_014445148.1 | 81 | acyl carrier protein | LnmL [AAN85525.1] | 50/63 |
| WP_014445149.1 | 315 | AT/DC | LnmK [AAN85524.1] | 45/62 |
| WP_014445150.1 | 95 | acyl carrier protein | VirA [BAF50727.1] | 49/59 |
| WP_014445151.1 | 403 | ketosynthase chain-length factor | MxnD [AGS77284.1] | 28/46 |
| WP_014445152.1 | 422 | beta-ketoacyl synthase | MxnD [AGS77284.1] | 32/46 |
| WP_014445153.1 | 78 | acyl carrier protein | ChiB [AAY89049.1] | 37/48 |
| WP_014445154.1 | 374 | 3-oxoacyl-ACP synthase | / | / |
| WP_014445155.1 | 246 | oxidoreductase | DifE [CAG23976.1] | 34/51 |
| WP_041831255.1 | 345 | luciferase-like monooxygenase | Cycloheximide [CCC21141.1] | 34/46 |
| WP_014445157.1 | 171 | hemerythrin domain | / | / |
| WP_014445158.1 | 381 | hydrolase | / | / |
| WP_014445159.1 | 283 | regulatory protein | papR1 [CBW45751.1] | 48/56 |
| WP_014445160.1 | 307 | MBL fold metallo-hydrolase | Psymberin [ADA82578.1] | 58/69 |
| WP_014445161.1 | 149 | hypothetical protein | / | / |
| WP_014445162.1 | 303 | nitroreductase family | / | / |
| WP_014445163.1 | 402 | acyltransferase | / | / |
| WP_014445164.1 | 399 | cytochrome P450 | / | / |

[a] Number of amino acids

**Table S175.** Predicted functions of ORFs in the NZ_KB913032.1 containing WP_020634864.1

| gene | aa[a] | putative function | Protein homologue | %identity/ %similarity |
|---|---|---|---|---|
| WP_051137561.1 | 998 | NRPS | JamO [AAS98786.1] | 33/53 |
| WP_026467562.1 | 158 | hypothetical protein | / | / |
| WP_039794296.1 | 64 | hypothetical protein | / | / |
| WP_084702170.1 | 286 | hypothetical protein | / | / |
| WP_020634855.1 | 1007 | NRPS | Ta1 [ABF85931.1] | 32/49 |
| WP_051137714.1 | 222 | hypothetical protein | / | / |
| WP_020634857.1 | 407 | beta-ketoacyl synthase | JamG [AAS98778.1] | 34/50 |
| WP_084702171.1 | 151 | acyl carrier protein | / | / |
| WP_020634859.1 | 407 | beta-ketoacyl synthase | SnaH [CBW45746.1] | 44/57 |
| WP_084702335.1 | 423 | aminotransferase | / | / |
| WP_020634861.1 | 322 | Zn-dependent oxidoreductase | MupE [AAM12917.1] | 31/43 |
| WP_020634862.1 | 253 | hypothetical protein | VirE [BAF50723.1] | 50/65 |
| WP_020634863.1 | 247 | hypothetical protein | VirD [BAF50724.1] | 44/52 |
| WP_020634864.1 | 404 | HMG-CoA synthase | LnmM [AAN85526.1] | 56/69 |
| WP_020634865.1 | 81 | acyl carrier protein | LnmL [AAN85525.1] | 46/62 |
| WP_020634866.1 | 314 | AT/DC | LnmK [AAN85524.1] | 48/64 |
| WP_020634867.1 | 90 | acyl carrier protein | SnaE2 [CBW45748.1] | 36/61 |
| WP_020634868.1 | 348 | maleylacetate reductase | / | / |
| WP_020634869.1 | 288 | hypothetical protein | / | / |

[a] Number of amino acids

**Table S176.** Predicted functions of ORFs in the NZ_FNOK01000001.1 containing WP_093260285.1

| gene | aa [a] | putative function | Protein homologue | %identity/ %similarity |
|---|---|---|---|---|
| WP_093260273.1 | 564 | GGDEF domain-containing protein | / | / |
| WP_093260275.1 | 446 | hypothetical protein | Trs2 [CBW45638.1] | 32/43 |
| WP_093260277.1 | 173 | hypothetical protein | / | / |
| WP_093260519.1 | 391 | esterase | / | / |
| WP_093260279.1 | 570 | polyketide synthase | VirA [BAF50727.1] | 41/55 |
| WP_093260281.1 | 270 | enoyl-CoA hydratase | VirE [BAF50723.1] | 47/61 |
| WP_093260283.1 | 273 | enoyl-CoA hydratase | SnaJ [CBW45744.1] | 47/54 |
| WP_093260285.1 | 409 | HMG-CoA synthase | LnmM [AAN85526.1] | 55/72 |
| WP_093260287.1 | 78 | acyl carrier protein | LnmL [AAN85525.1] | 47/63 |
| WP_093260289.1 | 295 | AT/DC | LnmK [AAN85524.1] | 59/68 |
| WP_093260291.1 | 91 | acyl carrier protein | SnaE2 [CBW45748.1] | 45/64 |
| WP_093260293.1 | 402 | ketosynthase chain-length factor | MxnD [AGS77284.1] | 29/44 |
| WP_093260295.1 | 426 | beta-ketoacyl synthase | MxnD [AGS77284.1] | 33/46 |
| WP_093260297.1 | 78 | acyl carrier protein | / | / |
| WP_093260299.1 | 368 | 3-oxoacyl-ACP synthase | / | / |
| WP_093260301.1 | 245 | SDR family oxidoreductase | ChxH [AFO59869.1] | 36/52 |
| WP_093260303.1 | 347 | LLM class flavin-dependent oxidoreductase | / | / |
| WP_093260521.1 | 315 | hypothetical protein | / | / |

[a] Number of amino acids

**Table S177.** Predicted functions of ORFs in the NZ_PVLV01000053.1 containing WP_105867428.1

| gene | aa [a] | putative function | Protein homologue | %identity/ %similarity |
|---|---|---|---|---|
| WP_105867456.1 | 147 | nuclear transport factor 2 | / | / |
| WP_105867418.1 | 347 | LLM class flavin-dependent oxidoreductase | / | / |
| WP_105867419.1 | 245 | SDR family oxidoreductase | ChxH [AFO59869.1] | 39/52 |
| WP_105867420.1 | 341 | acyltransferase | NspD [ADA69241.1] | 33/51 |
| WP_105867421.1 | 341 | 3-oxoacyl-ACP synthase | / | / |
| WP_105867422.1 | 91 | actinorhodin polyketide synthase | / | / |
| WP_105867423.1 | 422 | beta-ketoacyl synthase | MxnD [AGS77284.1] | 32/46 |
| WP_105867424.1 | 402 | ketosynthase chain-length factor | MxnD [AGS77284.1] | 29/45 |
| WP_105867425.1 | 86 | acyl carrier protein | SnaE2 [CBW45748.1] | 44/62 |
| WP_105867426.1 | 315 | AT/DC | LnmK [AAN85524.1] | 54/68 |
| WP_105867427.1 | 81 | acyl carrier protein | LnmL [AAN85525.1] | 49/66 |
| WP_105867428.1 | 409 | HMG-CoA synthase | LnmM [AAN85526.1] | 57/72 |
| WP_105867429.1 | 246 | enoyl-CoA hydratase | VirD [BAF50724.1] | 44/52 |
| WP_105867430.1 | 270 | crotonase | VirE [BAF50723.1] | 49/63 |
| WP_105867431.1 | 572 | polyketide synthase | VirA [BAF50727.1] | 44/55 |
| WP_105867457.1 | 314 | DUF1205 domain-containing protein | / | / |
| WP_105867432.1 | 392 | esterase | / | / |
| WP_105867433.1 | 453 | NDP-hexose 2,3-dehydratase | / | / |

[a] Number of amino acids

**Table S178.** Predicted functions of ORFs in the NZ_PQHZ01000005.1 containing WP_116041223.1

| gene | aa [a] | putative function | Protein homologue | %identity/ %similarity |
|---|---|---|---|---|
| WP_116041214.1 | 171 | hemerythrin domain-containing protein | / | / |
| WP_116041215.1 | 390 | beta-lactamase-related serine hydrolase | / | / |
| WP_116041216.1 | 562 | polyketide synthase | SnaE1 [CBW45749.1] | 43/55 |
| WP_116041218.1 | 284 | acyltransferase | SgvQ [AGN74897.1] | 48/61 |
| WP_116041219.1 | 244 | enoyl-CoA hydratase | snaK [CBW45743.1] | 43/59 |
| WP_116041221.1 | 243 | enoyl-CoA hydratase | MxnF [AGS77286.1] | 38/55 |
| WP_116041223.1 | 410 | HMG-CoA synthase | LnmM [AAN85526.1] | 53/71 |
| WP_116041225.1 | 86 | acyl carrier protein | LnmL [AAN85525.1] | 46/69 |
| WP_116041450.1 | 287 | AT/DC | LnmK [AAN85524.1] | 51/63 |
| WP_116041227.1 | 86 | acyl carrier protein | SnaE2 [CBW45748.1] | 47/66 |
| WP_116041229.1 | 401 | ketosynthase chain-length factor | PedM [AAW33972.1] | 25/43 |
| WP_116041231.1 | 422 | beta-ketoacyl synthase II | MxnD [AGS77284.1] | 30/45 |
| WP_116041233.1 | 78 | acyl carrier protein | ChiB [AAY89049.1] | 36/44 |
| WP_116041452.1 | 341 | 3-oxoacyl-ACP synthase | / | / |
| WP_116041235.1 | 332 | acyltransferase | JamP [AAS98787.1] | 35/51 |
| WP_116041237.1 | 245 | SDR family oxidoreductase | ChxH [AFO59869.1] | 34/51 |
| WP_116041239.1 | 344 | LLM class flavin-dependent oxidoreductase | / | / |
| WP_116041454.1 | 142 | nuclear transport factor 2 | / | / |

[a] Number of amino acids

**Table S179.** Predicted functions of ORFs in the LMNW01000012.1 containing KQQ40262.1

| gene | aa[a] | putative function | Protein homologue | %identity/ %similarity |
|---|---|---|---|---|
| KQQ40260.1 | 535 | diguanylate cyclase | / | / |
| KQQ40261.1 | 253 | 2-nitropropane dioxygenase | MupS [AAM12932.1] | 31/50 |
| KQQ40529.1 | 448 | 2-nitropropane dioxygenase | BatK [ADD82952.1] | 55/74 |
| KQQ40262.1 | 416 | HMG-CoA synthase | BonG [AFN27479.1] | 68/80 |
| KQQ40263.1 | 260 | enoyl-CoA hydratase | BaeH [CAG23955.1] | 48/64 |
| KQQ40264.1 | 353 | malonyl CoA-ACP transacylase | Fr9O [AIC32701.1] | 50/63 |
| KQQ40265.1 | 386 | polyketide synthase | ThaG [ABC34832.1] | 53/70 |
| KQQ40266.1 | 87 | acyl carrier protein | CalX [BAP05574.1] | 49/69 |
| KQQ40267.1 | 419 | beta-ketoacyl synthase | NspH [ADA69244.1] | 56/72 |
| KQQ40268.1 | 179 | hypothetical protein | / | / |
| KQQ40269.1 | 301 | alpha/beta hydrolase | / | / |
| KQQ40270.1 | 326 | acyltransferase | BaeD [CAG23951.1] | 34/50 |
| KQQ40271.1 | 514 | MFS transporter | LnmY [AAN85538.1] | 28/41 |
| KQQ40272.1 | 363 | transporter | / | / |
| KQQ40273.1 | 202 | hypothetical protein | / | / |
| KQQ40274.1 | 165 | antitermination factor | ElaA [AEC04347.1] | 25/41 |
| KQQ40275.1 | 407 | sodium:proton exchanger | / | / |
| KQQ40276.1 | 324 | PPTase | VirK [BAF50717.1] | 36/43 |
| KQQ40277.1 | 447 | sensor histidine kinase | Disorazols [AAY32961.1] | 31/46 |
| KQQ40278.1 | 192 | hypothetical protein | / | / |
| KQQ40279.1 | 231 | hypothetical protein | / | / |

[a] Number of amino acids

**Table S180.** Predicted functions of ORFs in the NZ_LN929761.1 containing WP_059006297.1

| gene | aa[a] | putative function | Protein homologue | %identity/ %similarity |
|---|---|---|---|---|
| WP_079031795.1 | 150 | DUF3224 domain | LnmZ' [AAN85540.1]' | 38/57 |
| WP_059006294.1 | 238 | N-acetylglucosaminyl deacetylase | LnmX [AAN85537.1] | 54/65 |
| WP_079031796.1 | 285 | thioesterase | LnmN [AAN85527.1] | 50/59 |
| WP_059006295.1 | 422 | beta-ketoacyl synthase | CorD [ADI59526.1] | 47/58 |
| WP_079031797.1 | 88 | acyl carrier protein | SnaG [CBW45747.1] | 45/60 |
| WP_059006297.1 | 410 | HMG-CoA synthase | LnmM [AAN85526.1] | 64/75 |
| WP_104530923.1 | 76 | acyl carrier protein | LnmL [AAN85525.1] | 55/72 |
| WP_104530924.1 | 315 | AT/DC | LnmK [AAN85524.1] | 49/60 |
| WP_059006300.1 | 1350 | alpha/beta fold hydrolase | LnmJ [AAN85523.1] | 51/65 |

[a] Number of amino acids

**Table S181.** Predicted functions of ORFs in the LAJX01000157.1 containing KJV05900.1

| gene | aa[a] | putative function | Protein homologue | %identity/ %similarity |
|---|---|---|---|---|
| KJV05897.1 | 199 | NAD(P)/FAD-dependent oxidoreductase | / | / |
| KJV05898.1 | 273 | short-chain dehydrogenase | Leinamycin [AAN85505.1] | 31/49 |
| KJV05899.1 | 280 | hypothetical protein VZ94_14860 | OnnF [AAV97874.1] | 59/76 |
| KJV05900.1 | 426 | HMG-CoA synthase | BaeG [CAG23954.2] | 71/82 |

[a] Number of amino acids

**Table S182.** Predicted functions of ORFs in the QHXD01000660.1 containing PYS20272.1

| gene | aa [a] | putative function | Protein homologue | %identity/ %similarity |
|---|---|---|---|---|
| PYS20270.1 | 830 | malonyl CoA-ACP transacylase | MisF [AKQ22696.1] | 41/58 |
| PYS20271.1 | 221 | MBL fold metallo-hydrolase | BaeB [CAG23949.2] | 44/62 |
| PYS20272.1 | 366 | HMG-CoA synthase | BatC [ADD82944.1] | 72/83 |

[a] Number of amino acids

**Table S183.** Predicted functions of ORFs in the NZ_JOJM01000004.1 containing WP_030670438.1

| gene[a] | aa[b] | putative function | Protein homologue | %identity/ %similarity |
|---|---|---|---|---|
| ORF1 | 2011 | NRPS | SnbDE [CBW45647.1] | 35/47 |
| WP_030670431.1 | 84 | acyl carrier protein | CalX [BAP05574.1] | 53/68 |
| WP_030670435.1 | 407 | beta-ketoacyl synthase | NspH [ADA69244.1] | 56/74 |
| WP_030670438.1 | 430 | HMG-CoA synthase | BaeG [CAG23954.2] | 66/78 |
| WP_030670441.1 | 798 | 2-nitropropane dioxygenase | BatK [ADD82952.1] | 54/72 |
| WP_078868641.1 | 644 | S-malonyltransferase | OocV [AFX60344.1] | 47/63 |
| WP_030670448.1 | 411 | cytochrome P450 | BaeS [CAG23962.1] | 37/55 |
| WP_030670451.1 | 275 | pentapeptide repeat-containing protein | / | / |

[a] ORFs are proteins without protein ID provided by NCBI [b] Number of amino acids

**Table S184.** Predicted functions of ORFs in the NZ_JTJG01000021.1 containing WP_047478363.1

| gene | aa [a] | putative function | Protein homologue | %identity/ %similarity |
|---|---|---|---|---|
| WP_047478360.1 | 405 | beta-ketoacyl synthase | NspH [ADA69244.1] | 57/77 |
| WP_047478363.1 | 419 | HMG-CoA synthase | BaeG [CAG23954.2] | 75/85 |
| WP_047478364.1 | 253 | enoyl-CoA hydratase | BaeH [CAG23955.1] | 58/75 |
| WP_047478367.1 | 249 | enoyl-CoA hydratase | BatE [ADD82946.1] | 66/85 |
| WP_047478370.1 | 279 | sugar phosphate isomerase | / | / |
| WP_047478374.1 | 400 | glycosyltransferase | CylN [ARU81128.1] | 24/42 |
| WP_047478379.1 | 155 | hypothetical protein | / | / |
| WP_047478383.1 | 478 | hypothetical protein | / | / |

[a] Number of amino acids

**Table S185.** Predicted functions of ORFs in the NZ_LABX01000062.1 containing WP_048463408.1

| gene | aa[a] | putative function | Protein homologue | %identity/ %similarity |
|---|---|---|---|---|
| WP_048463402.1 | 369 | hypothetical protein | / | / |
| WP_048463403.1 | 278 | enoyl-CoA hydratase | CorF [ADI59528.1] | 31/49 |
| WP_048463404.1 | 458 | 2-nitropropane dioxygenase | BatK [ADD82952.1] | 51/69 |
| WP_048463405.1 | 294 | S-malonyltransferase | RizF [CCA89330.1] | 51/63 |
| WP_048463406.1 | 249 | enoyl-CoA hydratase | BonI [AFN27485.1] | 55/70 |
| WP_048463407.1 | 264 | enoyl-CoA hydratase | CorF [ADI59528.1] | 39/51 |
| WP_048463408.1 | 411 | HMG-CoA synthase | BonG [AFN27479.1] | 51/65 |

[a] Number of amino acids

**Table S186.** Predicted functions of ORFs in the NZ_LN929762.1 containing WP_059006317.1

| gene | aa[a] | putative function | Protein homologue | %identity/ %similarity |
|---|---|---|---|---|
| WP_059006309.1 | 393 | alpha/beta fold hydrolase | MupV [AAM12938.1] | 29/39 |
| WP_079031800.1 | 321 | chlorinating enzyme | / | / |
| WP_059006311.1 | 600 | NRPS | Kirromycin [CAN89656.1] | 50/60 |
| WP_059006361.1 | 328 | hypothetical protein | LnmH [AAN85521.1] | 42/58 |
| WP_079031801.1 | 767 | AT/Ox | DifA [CAG23974.1] | 50/68 |
| WP_059006316.1 | 261 | enoyl-CoA hydratase | MxnF [AGS77286.1] | 41/53 |
| WP_059006317.1 | 410 | HMG-CoA synthase | JamH [AAS98779.1] | 49/64 |
| WP_059006319.1 | 299 | hypothetical protein | LnmE [AAN85518.1] | 42/58 |
| WP_079031802.1 | 249 | Crp/Fnr family transcriptional regulator | LnmO [AAN85528.1] | 46/65 |
| WP_104530925.1 | 431 | cytochrome P450 | ElaG [AEC04353.1] | 37/55 |
| WP_079031803.1 | 243 | Crp/Fnr family transcriptional regulator | LnmO [AAN85528.1] | 42/60 |
| WP_059006327.1 | 184 | DUF1697 domain | Leinamycin [AAN85541.1] | 33/49 |
| WP_059006363.1 | 1168 | hypothetical protein | / | / |

[a] Number of amino acids

**Table S187.** Predicted functions of ORFs in the NZ_FZOD01000025.1 containing WP_089209624.1

| gene | aa [a] | putative function | Protein homologue | %identity/ %similarity |
|---|---|---|---|---|
| WP_089209621.1 | 189 | carbonic anhydrase | / | / |
| WP_089209622.1 | 188 | HIT domain-containing protein | misakinolide [AKQ22692.1] | 36/50 |
| WP_089209623.1 | 315 | PfaD family polyunsaturated fatty acid | LnmG [AAN85520.1] | 67/78 |
| WP_089209624.1 | 410 | HMG-CoA synthase | MxnE [AGS77285.1] | 55/70 |
| WP_089209625.1 | 424 | cation/H(+) antiporter | / | / |
| WP_089209626.1 | 501 | hypothetical protein | / | / |
| WP_089209627.1 | 217 | hypothetical protein | / | / |

[a] Number of amino acids

**Table S188.** Predicted functions of ORFs in the NZ_OMOF01000372.1 containing WP_106800321.1

| gene | aa [a] | putative function | Protein homologue | %identity/ %similarity |
|---|---|---|---|---|
| WP_106800320.1 | 234 | hypothetical protein | / | / |
| WP_106800321.1 | 421 | HMG-CoA synthase | BatC [ADD82944.1] | 76/88 |
| WP_106800322.1 | 253 | enoyl-CoA hydratase | BaeH [CAG23955.1] | 62/78 |
| WP_106800323.1 | 484 | radical SAM protein | / | / |

[a] Number of amino acids

**Table S189.** Predicted functions of ORFs in the NZ_NOLN01000021.1 containing WP_011997326.1

| gene | aa[a] | putative function | Protein homologue | %identity/ %similarity |
|---|---|---|---|---|
| WP_005772946.1 | 252 | ATP-binding cassette domain | LnmR [AAN85531.1] | 32/52 |
| WP_011997319.1 | 526 | ABC transporter permease | / | / |
| WP_011997320.1 | 395 | serine protease | / | / |
| WP_005772950.1 | 246 | serine/threonine protein kinase | / | / |
| WP_011997321.1 | 149 | hypothetical protein | / | / |
| WP_011997322.1 | 150 | hypothetical protein | / | / |
| WP_043880970.1 | 248 | hypothetical protein | / | / |
| WP_011997324.1 | 92 | acyl carrier protein | ThaI [ABC35804.1] | 41/62 |
| WP_080512905.1 | 204 | enoyl-CoA hydratase | CorF [ADI59528.1] | 41//59 |
| WP_011997326.1 | 412 | HMG-CoA synthase | MxnE [AGS77285.1] | 53/72 |
| WP_011997327.1 | 411 | beta-ketoacyl synthase | DipR [AGS06821.1] | 40/60 |
| WP_011997328.1 | 591 | polyketide synthase | LglD [AIU36100.1] | 46/64 |
| WP_011997329.1 | 481 | thioesterase | LnmN [AAN85527.1] | 31/47 |
| WP_011997330.1 | 1280 | NRPS | JamO [AAS98786.1] | 34/52 |
| WP_011997331.1 | 336 | hypothetical protein | / | / |

[a] Number of amino acids

**Table S190.** Domain and module organization of the *S.* sp. CB01881 AT-less PKS gene cluster

| gene[a] | aa[b] | putative Function | Accession number [Origin] | Identity(%)/ Similarity (%) |
|---|---|---|---|---|
| orf -1 | 392 | hypothetical protein | KDN85868.1 [*Kitasatospora*] | 67/75 |
| OrfA | 432 | cytochrome P450 | PBC70420.1 [*S. sp.* TLI_235] | 77/85 |
| OrfB | 206 | translation factor | BAU87570.1 [*S. laurentii*] | 85/93 |
| OrfC | 245 | PPTase | CCA59555.1 [*S. venezuelae* ] | 66/76 |
| OrfD | 443 | MDO-like protein | OGN43113.1 [*Caulobacterales*] | 40/52 |
| OrfE | 302 | short-chain dehydrogenase | OHE18601.1 [*Syntrophobacterales*] | 60/77 |
| OrfF | 321 | acyltransferase | SMF97326.1 [*Methylomagnum ishizawai*] | 36/48 |
| OrfG | 419 | malonyl-ACP decarboxylase | SDL84277.1 [*Dendrosporobacter*] | 59/73 |
| OrfH | 82 | acyl carrier protein | EIP85586.1 [*Burkholderia thailandensis*] | 61/85 |
| OrfI | 247 | enoyl-CoA hydratase | ATY28343.1 [*Bacillus velezensis*] | 64/76 |
| OrfJ | 260 | enoyl-CoA hydratase | ARL19220.1 [*Burkholderia pseudomallei*] | 57/75 |
| OrfK | 419 | HMG-CoA synthase | AMH40430.1 [*Leptolyngbya sp.* ] | 72/83 |
| OrfLa | 414 | malonyl CoA-ACP transacylase | OEU94861.1 [*S.oceani*] | 70/79 |
| OrfM | 451 | PfaD family protein, partial | EFL43494.1 [*S.* griseoflavus Tu4000] | 80/88 |
| OrfN | 3122 | AT-less type I PKS | KQS09640.1 [*Brevibacillus sp.* Leaf182] | 46/62 |
| OrfO | 3245 | AT-less type I PKS | OEU94845.1 [*S.* oceani] | 61/70 |
| OrfP | 2902 | AT-less type I PKS | OEU94845.1 [*S.* oceani] | 63/72 |
| OrfQ | 4675 | AT-less type I PKS | OEU95469.1 [*S.* oceani] | 58/68 |
| OrfR | 5713 | AT-less type I PKS | EFL43485.1 [*S. griseoflavus* Tu4000] | 68/76 |
| OrfS | 2613 | AT-less type I PKS | EFL43484.1 [*S. griseoflavus* Tu4000] | 66/75 |
| OrfT | 533 | methylmalonyl-CoA carboxyltransferase | PJN24676.1 [*Kitasatospora sp.* CB02891] | 95/96 |
| OrfU | 67 | hypothetical protein | CCH32753.1 [*Saccharothrix espanaensis*] | 58/66 |
| OrfV | 204 | TetR transcriptional regulator | KJS52667.1 [*S.* rubellomurinus] | 84/89 |
| orf +1 | 64 | hypothetical protein | AUG81647.1 [*Kitasatospora*] | 79/83 |

[a]orf(-1) and orf(+1) are predicted to represent the upstream and downstream boundaries of the 1881 AT-less gene cluster. [b]Number of amino acids.

**Table S191.** Annotation of *S.* sp. CB01881 AT-less gene clusters in comparison of characterized AT-less PKS gene cluster

| gene[a] | aa[b] | putative Function | protein homologue | Identity(%)/ Similarity (%) |
|---|---|---|---|---|
| orf -1 | 392 | / | / | |
| OrfA | 432 | cytochrome P450 | BaeS [ABS74066.1] | 32/47 |
| OrfB | 206 | / | / | |
| OrfC | 245 | / | / | 66/76 |
| OrfD | 443 | / | / | |
| OrfE | 302 | short-chain dehydrogenase | BatT [ADD82961.1] | 29/46 |
| OrfF | 321 | acyltransferase | BryP [ABM63531.1] | 33/55 |
| OrfG | 419 | beta-ketoacyl synthase | TstN [AGN11888.1] | 61/74 |
| OrfH | 82 | acyl carrier protein | CalX [BAP05574.1] | 58/81 |
| OrfI | 247 | enoyl-CoA hydratase | BatE [ADD82946.1] | 63/80 |
| OrfJ | 260 | enoyl-CoA hydratase | CalS [BAP05579.1] | 56/71 |
| OrfK | 419 | HMG-CoA synthase | BatC [ADD82944.1] | 69/83 |
| OrfLa | 414 | acyl transferase II | BonK [AFN27477.1] | 53/68 |
| OrfM | 451 | PfaD family protein, partial | Myxovirescin [ABF87992.1] | 55/71 |
| OrfN | 3122 | AT-less type I PKS | ElaK [ABM63527.1] | 52/57 |
| OrfO | 3245 | AT-less type I PKS | EtnF [CAN93348.1] | 45/51 |
| OrfP | 2902 | AT-less type I PKS | SorA [ADN68476.1] | 43/50 |
| OrfQ | 4675 | AT-less type I PKS | SorB [ADN68477.1] | 45/48 |
| OrfR | 5713 | AT-less type I PKS | OnnB [AAV97870.1] | 41/46 |
| OrfS | 2613 | AT-less type I PKS | CorL [ADI59534.1] | 34/47 |
| OrfT | 533 | methylmalonyl-CoA decarboxylase | MmdA [CBW45762.1] | 87/93 |
| OrfU | 67 | Null | / | / |
| OrfV | 204 | Null | / | / |
| orf +1 | 64 | hypothetical protein | Streptomycin [CAH94355.1] | 53/57 |

[a]orf(-1) and orf(+1) are predicted to represent the upstream and downstream boundaries of the *S.* sp. CB01881 AT-less gene cluster. [b]Number of amino acids.

**Table S192.** Soil samples location data. The oblique line in the table represents no corresponding data

| Sample ID | LONG | LAT | ALT | Country | Province | City/Area |
|---|---|---|---|---|---|---|
| S1 | 104.668725 | 26.738486 | 1765.01 | China | Guizhou | Liupanshui |
| S2 | 104.753513 | 26.710197 | 1954.77 | China | Guizhou | Liupanshui |
| S3 | 105.890373 | 32.374193 | / | China | Sichuan | Guangyuan |
| S4 | 105.877026 | 32.374015 | / | China | Sichuan | Guangyuan |
| S5 | 121.510592 | 31.304189 | / | China | Shanghai | Yangpu |
| S6 | / | / | / | China | Guangxi | Guilin |
| S7 | / | / | / | China | Sinkiang | Korla |
| S8 | 116.402869 | 39.927599 | / | China | Beijing | Dongcheng |
| S9 | / | / | / | China | Hunan | Zhangjiajie |
| S10 | 104.757722 | 26.772305 | / | China | Guizhou | Liupanshui |
| S11 | 104.759211 | 26.777525 | 2254.3 | China | Guizhou | Liupanshui |
| S12 | 104.717069 | 26.698508 | 1756 | China | Guizhou | Liupanshui |
| S13 | 104.702102 | 26.678705 | 1883 | China | Guizhou | Liupanshui |
| S14 | 104.690761 | 26.770943 | 2035.99 | China | Guizhou | Liupanshui |
| S15 | 104.674377 | 26.736605 | 1775 | China | Guizhou | Liupanshui |
| S16 | 104.790611 | 26.697722 | 1885 | China | Guizhou | Liupanshui |
| S17 | 104.723333 | 26.69675 | 1824 | China | Guizhou | Liupanshui |
| S18 | 104.727369 | 26.695041 | 1868 | China | Guizhou | Liupanshui |
